# High-Throughput Synthesis and Screening of a Cyanimide Library Identifies Selective Inhibitors of ISG15-specific Protease USP18

**DOI:** 10.1101/2025.04.18.649523

**Authors:** Raymond Kooij, Vito Pol, Jin Gan, Bjorn R. van Doodewaerd, Aysegul Sapmaz, Günter Fritz, Klaus-Peter Knobeloch, Paul P. Geurink

## Abstract

High-throughput screening (HTS) of (large) compound collections is a critical early step in many drug discovery programs, enabling the rapid identification of lead molecules with desirable biological activity. The success of HTS depends heavily on the quality of the compound libraries used, and as such, the development of targeted libraries has emerged as a promising approach to enhance the effectiveness of HTS efforts. However, the acquisition and synthesis of such libraries remain costly and labor-intensive, often yielding compound quantities far exceeding the small amounts required for screening. To address these challenges, we present a high-throughput synthesis-to-screening method for the efficient in-plate generation and immediate HTS of a deubiquitinase (DUB)-focused compound library. Central to our approach is the use of an Echo acoustic liquid handler, which enables precise nanoliter-scale transfers of DMSO-based solutions, facilitating efficient and miniaturized synthesis directly in 1,536-well plates. Using this platform, we constructed a library of 7,536 compounds featuring a DUB-privileged cyanimide warhead and screened it against a panel of twelve DUBs and ubiquitin-like proteases. This identified two structurally related molecules with selective inhibitory activity against the interferon-stimulated gene 15 (ISG15) protease USP18, which we further developed into a first-in-class USP18 inhibitor with 35 nM potency. This compound, BB07CA902, demonstrated exceptional specificity for USP18 across a panel of 41 DUBs and effectively increased ISGylation levels in cells by inhibiting USP18 activity. With our technology, we enabled the efficient preparation of large DUB-targeted cyanimide-based libraries, which will accelerate future DUB inhibitor development.

## Introduction

Despite the modern-day availability of various virtual methods, many drug discovery projects often commence with the physical high-throughput screening (HTS) of large compound libraries to identify lead compounds.^[1]^ In addition to a robust screening assay, the success of a screening campaign depends heavily on the availability of high-quality libraries of drug-like molecules. Poor-quality or poorly designed libraries can lead to false positives, delaying the identification of viable drug candidates.^[2]^ Compound libraries come in different flavours, such as large, diverse compound libraries, composed of a wide range of chemical scaffolds and functional groups, fragment libraries^[3]^ containing small, low molecular weight compounds that serve as the building blocks for larger drug-like molecules, and focused or targeted libraries,^[4]^ which are curated collections that target specific proteins or biological pathways, offering a more streamlined approach to drug discovery but with a potentially narrower scope. The production of high-quality screening libraries remains a significant challenge. Traditional methods for synthesizing compound libraries are often costly, labour-intensive, and time-consuming, requiring extensive synthetic chemistry expertise and infrastructure. Therefore, a number of strategies have been developed for the rapid and efficient generation of screening libraries, such as diversity-oriented synthesis (DOS),^[5]^ DNA-encoded library (DEL) technology,^[6]^ and parallel and combinatorial synthesis.^[7]^ In recent years, advances in automated synthesis platforms and miniaturized chemistry have further accelerated the generation of screening libraries.^[8]^

A major technological advancement in screening sensitivity and throughput was introduced with the development of acoustic dispensing (i.e. tipless dispensing using soundwaves), which opened novel avenues for parallel reagent delivery, analysis and in-situ testing.^[9]^ The Echo acoustic liquid handler, which accurately transfers DMSO or aqueous liquids in droplets of 2.5 nL or 25 nL from a source plate to an inverted destination plate positioned above the source plate by using ultrasonic acoustic energy, was initially developed for improved compound handling in drug discovery screens, but its applications have expanded into the fields of sequencing, genomics, protein crystallography and microarrays. Recently, some groups have started to explore the possibilities of acoustic liquid transfer in synthetic organic chemistry.^[10-15]^ Upon recognizing its great potential in terms of the rapid miniaturized synthesis of large compound sets (even possible in 1,536-well format), only requiring microliter reaction volumes, and the fact that the reaction plates can be directly used in HTS, again using acoustic dispensing, we opted to investigate whether we could implement Echo-mediated in-plate synthesis to boost our ongoing efforts of developing inhibitors for the highly challenging class of deubiquitinating enzymes (DUBs) and related proteases of ubiquitin-like proteins (UbL).

There are about 100 DUBs in human, which reverse ubiquitination, a post-translational modification that involves the instalment of (chains of) the 8.5 kDa protein ubiquitin (Ub) and that plays a major role in almost all cellular processes.^[16]^ The DUBs are classified into several families, like the USPs (ubiquitin-specific proteases), UCHs (ubiquitin carboxy-terminal hydrolases), MJDs (Machado–Josephin domain-containing proteases), OTUs (ovarian tumour proteases), or JAMMs (JAB1, MPN, MOV34 family). While most DUBs are cysteine peptidases, the JAMMs are zinc metallopeptidases. DUBs effectively erase the signal given by a certain polyUb chain, thereby regulating the stability, localization and function of substrate proteins.^[17]^ For many DUBs, it has been shown that their malfunctioning contributes to a wide variety of diseases, including cancer^[18]^ and neurodegenerative disorders,^[19]^ which has initiated several drug discovery programs aimed at developing selective DUB inhibitors. Despite several reports on DUB inhibitors, it remains highly challenging to obtain compounds that potently target specific members of the DUB family.^[20]^

The Ub-like protein ISG15, Interferon Stimulated Gene of 15 kDa, is strongly induced by Type-I interferons as part of the innate immune response upon viral and bacterial infections.^[21]^ Similar to ubiquitination, ISGylation is a reversible process. Ubiquitin-Specific Protease 18 (USP18) is a member of the USP subfamily of DUBs but lacks deubiquitinase activity and is the major ISG15-specific deconjugating enzyme.^[22]^ USP18 also functions as a feedback regulator of type I interferon signalling by binding to the interferon receptor 2 (IFNAR2) subunit, thereby preventing dimerization and subsequent signalling. This activity is vital but independent of USP18’s enzymatic activity.^[23,24]^ Interestingly, USP18-deficient mice suffer from brain injuries and premature death, whereas transgenic mice expressing catalytically inactive USP18 are healthy, have increased levels of ISG15 conjugates and are resistant to vaccinia virus, influenza B virus, and coxsackievirus infections.^[25,26]^ As such, an effective USP18 inhibitor may serve as a suitable lead for the development of antiviral agents, but to date, no inhibitors specifically and effectively targeting USP18 have been reported.

An interesting compound class reported to act as (semi)-reversible covalent DUB inhibitors are compounds equipped with a cyanimide moiety. This group of compounds was initially identified as targeting cathepsins, where they react with an active site cysteine residue, resulting in the formation of an isothiourea adduct. Cyanimides are remarkably stable under physiological conditions; they do not easily react with alcohols and amines,^[27]^ and they are stabile towards free cysteine and glutathione,^[28]^ indicating minimal change in unspecific reactions in a cellular environment. In particular, compounds having the cyanopyrrolidine scaffold or derivatives thereof have been identified as potent DUB inhibitors. For example, compounds **1-4** (Figure 1) were reported as nanomolar inhibitors of USP30,^[29]^ and compound **5** (Figure 1) was reported as a UCHL1 inhibitor.^[30]^ This compound class has now been designated as ‘DUB-privileged’, which is underscored by the reporting of several cyanimide inhibitors for different DUBs, including UCHL1,^[31-34]^ USP7,^[35]^ Cezanne,^[36]^ JOSD1,^[37]^ and some that display broad-spectrum DUB inhibition.^[38,39]^ Because the DUBs targeted by these cyanimides are structurally diverse and belong to different subfamilies, this compound class offers great potential for inhibiting many other DUBs. We reasoned that the large-scale production of thousands of compounds equipped with cyanimide-based warheads, in combination with DUB activity screens, would yield novel cyanimide DUB inhibitors. This provided us with a unique opportunity for our envisioned high-throughput in-plate synthesis coupled to HTS strategy. We here present the Echo-mediated miniaturized synthesis of a library containing 7,536 cyanimides in 1,536 well plates. The subsequent HTS on a panel of 12 DUBs and UbL proteases identified several hits for different DUBs. Re-synthesis, validation and chemical optimization of two of these hits resulted in the development of a first-in-class inhibitor for ISG15 protease USP18 with 35 nM potency *in vitro*. This compound, BB07CA902, showed full USP18 specificity amongst a panel of 41 DUBs, and we showed this inhibitor to be active in cells, as it increased cellular protein ISGylation by inhibiting USP18. With our technology, we enabled the efficient preparation of large DUB-targeted cyanimide-based libraries, which will accelerate future DUB inhibitor development.

**Figure 1.**
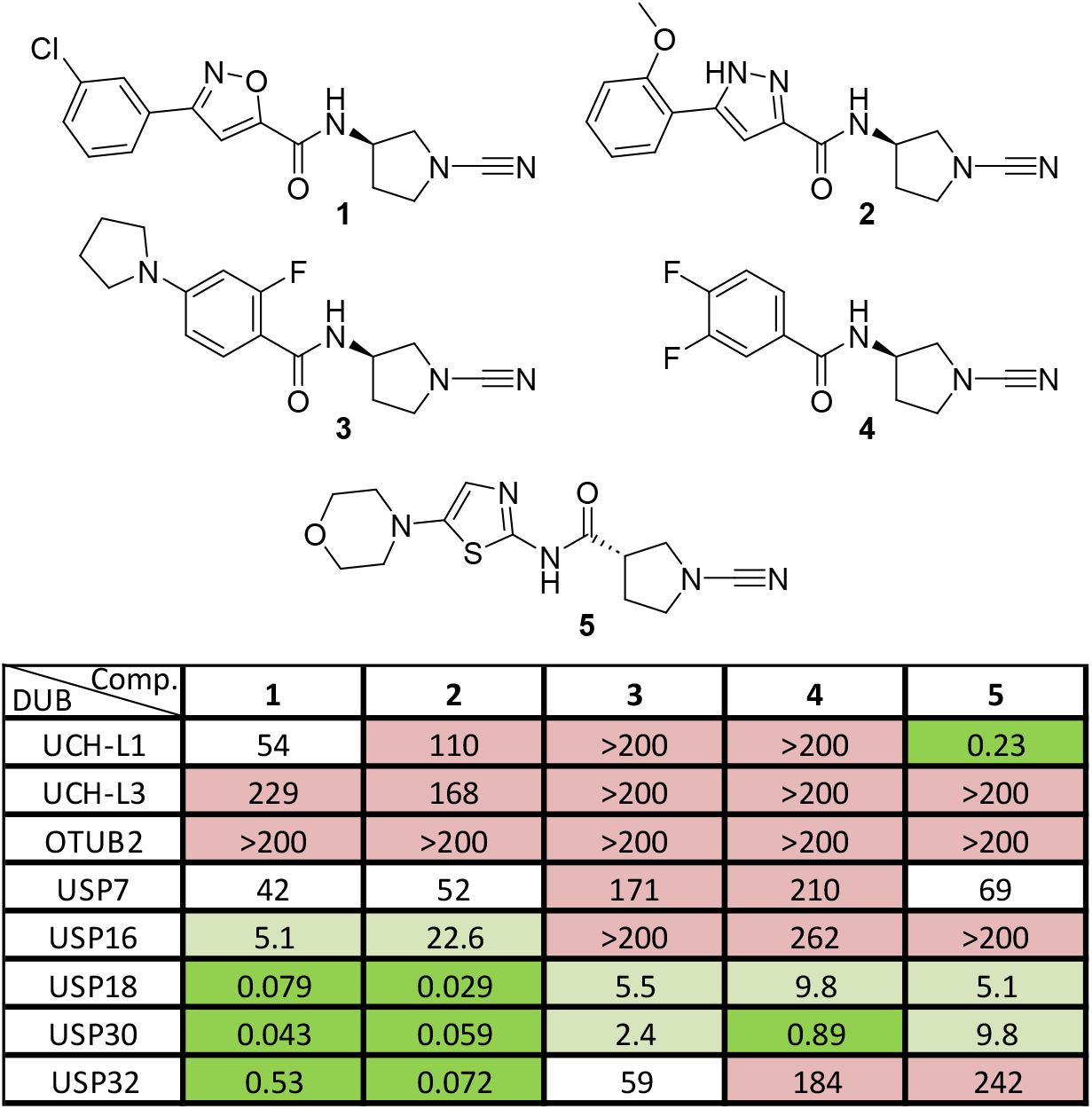
Structures and inhibition data (IC_50_ (µM) values) of cyanimides **1-5** on a panel of DUBs. Colours indicate inhibitory potency: green <1 µM; light green 1-20 µM; white 20-100 µM; pink >100 µM.

## Results and Discussion

Although compounds **1-4** were reported as potent inhibitors of USP30 and compound **5** to target UCHL1 (Figure 1), no data on cross-reactivity with other DUBs were disclosed.^[29-30]^ To gain insights into the selectivity and inhibitory potency of compounds **1-5** towards DUBs other than their respective targets, the compounds were synthesized, purified and subjected to IC_50_ determination on a small panel of DUBs from different DUB families (Figure 1, Figure S1). This panel included five proteases from the USP family, two from the UCH family, and one OTU. The IC_50_ values were determined in a well-established biochemical activity assay using the fluorogenic substrate Ub-Rho-morpholine.^[32]^ For USP18, which is unreactive to Ub but specific for ISG15, we used the ISG15_CT_-Rho-morpholine (consisting of the C-terminal domain of ISG15) substrate to monitor its activity (see Supporting Information for preparation of this substrate). Reported UCHL1 inhibitor **5**, with an inverted amide bond compared to the other compounds, indeed potently inhibited UCHL1 and showed about 20-40 fold less potency towards USP18 and USP30 (Figure 1). Other DUBs were not significantly inhibited by compound **5**. Compounds **1** and **2** not only showed nanomolar potency towards their target USP30, but also towards USP18 and USP32, while leaving the activity of other DUBs largely unchanged. Interestingly, compounds **3** and **4**, comprising an identical (*R*)-3-amino-1-cyanopyrrolidine moiety, proved much less potent, with only compound **4** showing potent USP30 inhibition. These findings highlight that small modifications on these types of cyanimide inhibitors, such as altering the amide orientation or carboxylate side, can significantly affect the potency and selectivity of DUB inhibition. We therefore opted to exploit this observation to develop more potent and DUB-selective inhibitors.

Rather than going through a laborious, time-consuming endeavour of synthesizing, purifying and testing individual compounds, we adopted a more efficient, high-throughput-based synthesis strategy, which encompassed reaction miniaturization by in-plate synthesis, as outlined in Figure 2A. The key component of this strategy is the Echo acoustic liquid dispenser. The strategy was as follows: DMSO solutions of reagents and reactants are transferred from Echo-compatible source plates and combined in new plates (the destination plates), where the product-forming reactions take place. These plates then serve as the compound library plates for HTS. The resulting crude mixtures of the newly formed compounds are then transferred to multiple copies of assay plates for DUB inhibition screening. To ensure effectiveness, this strategy needs to meet certain criteria: the chemical reaction should work in DMSO, be efficient, high yielding, proceed without significant side-product formation, all while avoiding reagents that might interfere in subsequent (biochemical) assays. Retro-synthetically, the target cyanimide inhibitors can be formed through an amide coupling between a cyanimide moiety-containing amine and a carboxylic acid (Figure 2B). This well-established, often high-yielding amidation reaction can be executed in DMSO, making it well-suited for the in-plate synthesis strategy. To optimize the amidation procedure, we tested several reaction conditions for synthesizing model compound **3** in DMSO, by varying different coupling reagents, bases, and the molar ratios between reactants and reagents (Figure 2C and Supporting Information). Reactions were monitored by LC-MS analysis, and the optimal reaction conditions were chosen based on the degree of consumed amine, optimal product formation and the least amount of undesired side product formation. The best results were achieved using 1 equivalent amine, 5 equivalents carboxylic acid and 5 equivalents of a combination of *N,N*’-diisopropylcarbodiimide (DIC) and 1-hydroxybenzotriazole (HOBt) (Figure 2D, Figure S2).

**Figure 2.**
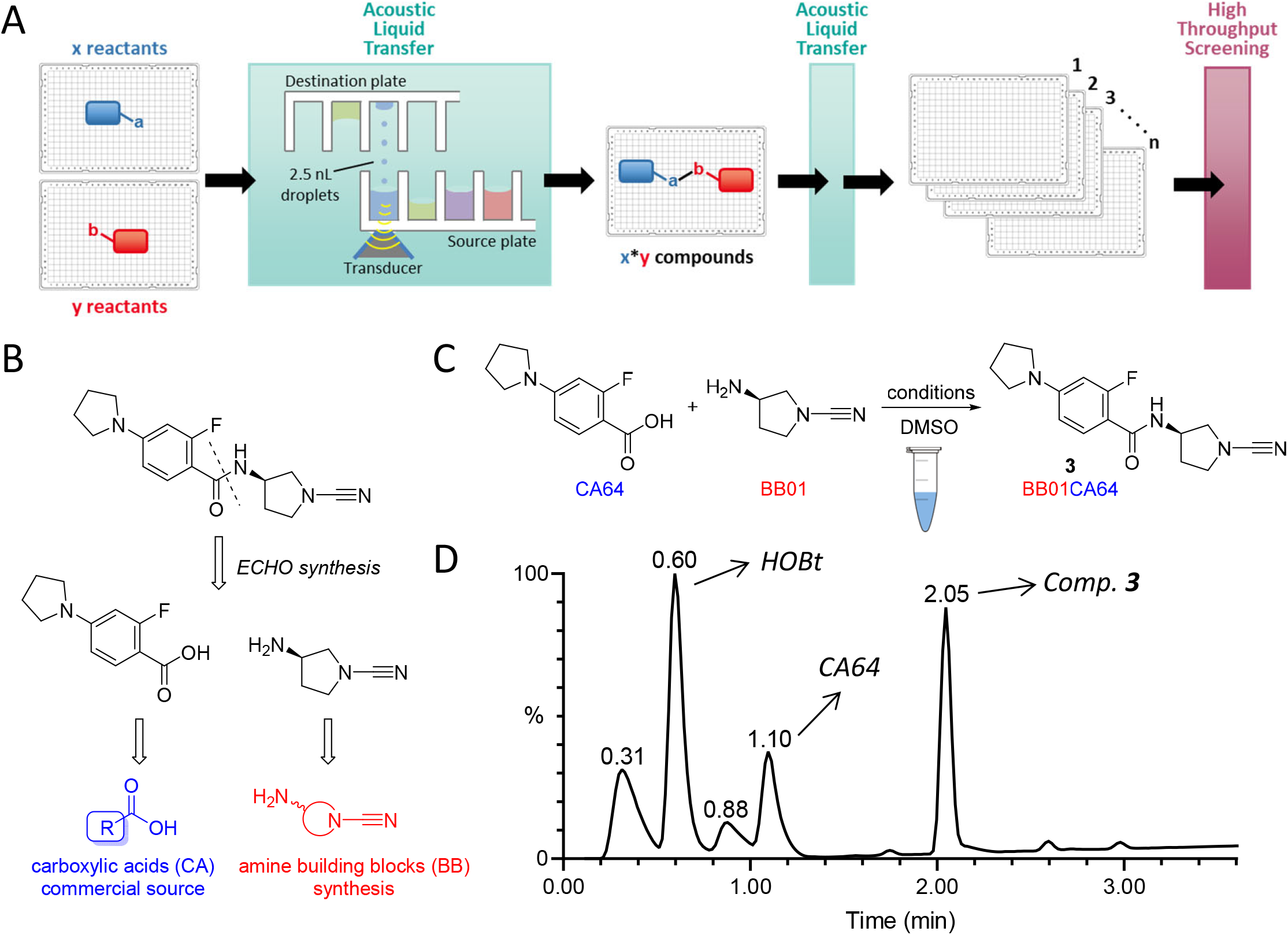
Echo synthesis set-up and reaction conditions optimizations. A) Schematic representation of in-plate Echo synthesis coupled to high-throughput screening strategy. Reactants are transferred from “Echo source” to “Echo destination” plates by acoustic liquid transfer. Formed crude products are then transferred by acoustic liquid transfer to assay plates for high-throughput screening. B) Retro-synthesis of a representative target compound. The compounds are formed through an amide coupling between a carboxylic acid (CA) and the cyanimide-containing amine building block (BB). C) Model reaction used to optimize reaction conditions and explanation of compound numbering: Compound ID’s are unique combinations of the amine (BB number) and carboxylic acid (CA number). D) HPLC analysis chromatogram of the model reaction shown in panel C under optimized conditions (BB 1 eq., CA 5 eq., DIC 5 eq., HOBt 5 eq., 10 mM (BB) in DMSO, room temperature, overnight).

After optimizing synthesis conditions, we proceed with a first in-plate synthesis (Figure 3A). We selected 89 carboxylic acids from our in-house collection, ensuring that they lacked any moiety that could interfere with the amide coupling reaction. Additionally, 16 cyclic cyanimide amine building blocks were synthesized in-house from readily available cyclic diamines (for details, see Supporting Information). This panel (Figure 3B) was designed to cover a broad range of cyanimide reactivity by for example varying the ring size (BB01, BB04, BB05) and amine location (BB02, BB04, BB11), by including electron-withdrawing moieties (BB13, BB15), and by using multicyclic systems (BB06, BB07, BB12, BB14). For the initial test, we combined the 89 carboxylic acids with seven cyanimide building blocks (BB1-4 and BB6-8) in two 384-well low-dead-volume (LDV) Echo plates (final volume was 10 µL per well), pre-charged with DIC/HOBt in the above-described ratios and using the plate layout as shown in Figure 3C and Figure S3. The success rate of the in-plate synthesis was assessed by LC-MS analysis. Out of 623 potential newly formed compounds, 274 compounds (44%) were evaluated by LC-MS analysis and labelled according to the quality of product formation as depicted in Figure 3C and Figure S3. Product formation was confirmed for 232 compounds, resulting in a success rate of ∼85%. Encouraged by these results, we aimed to expand the library and concomitantly reduce the reaction volume. As such, we purchased additional 382 carboxylic acids, taking into account the previously set requirements (see Supporting Information for a detailed description of the selection criteria), bringing the total of 471 carboxylic acids (CA1-471) and combined them with the full panel of 16 cyanimide amines (BB1-16) in a 1,536-well format with a total volume of 2.5 µL per well. This resulted in the theoretical formation of 7,536 unique compounds distributed over five 1,536-well plates as outlined in Supporting Information Figure S4. This time, the product formation assessment by LC-MS analysis was performed using a so-called hockey stick or checkmark approach (going diagonally through the plate to cover all rows and columns), which again indicated an overall success rate of ∼85% (Figure 3D, Figures S5, Figure S6).

**Figure 3.**
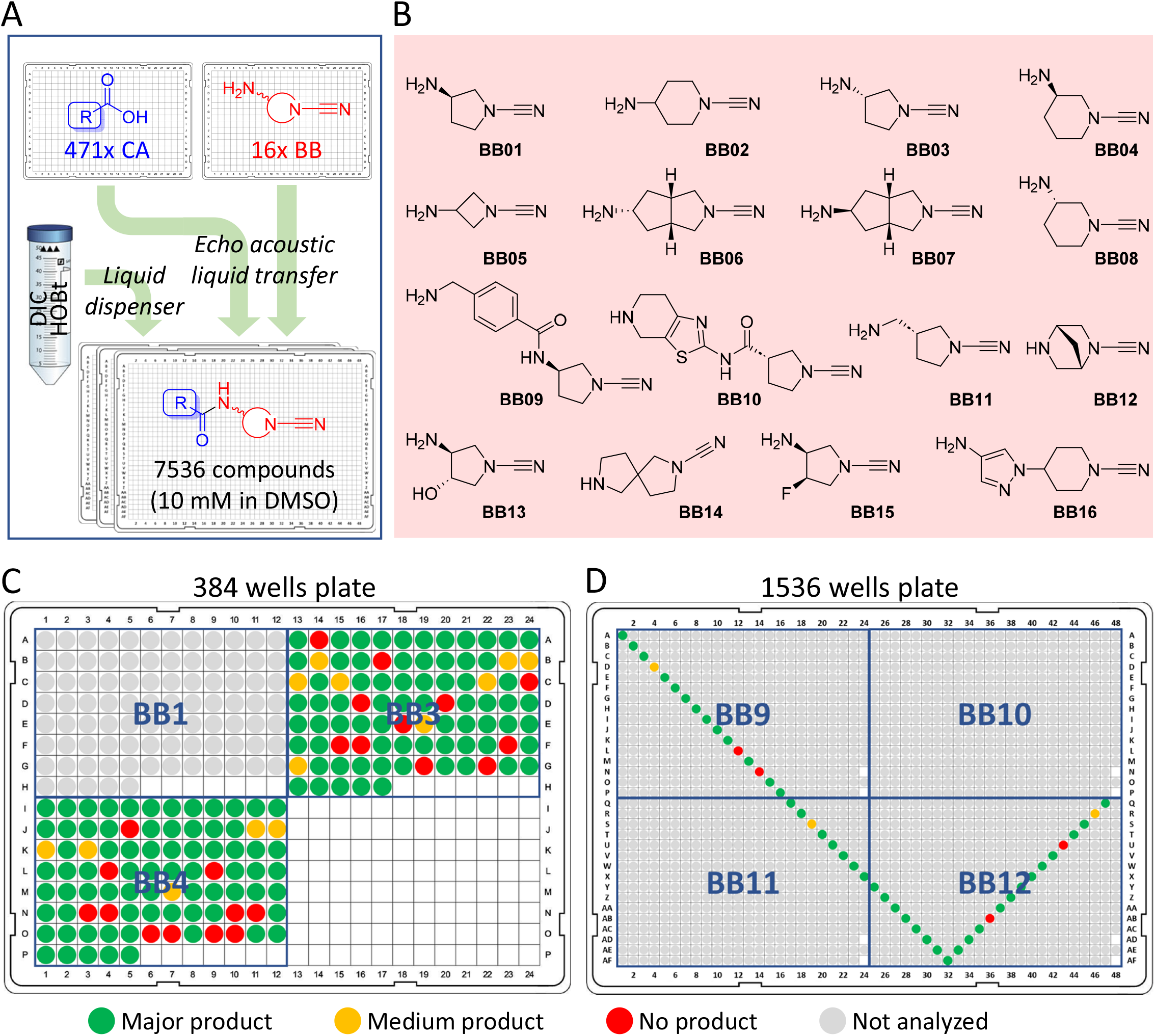
In-plate Echo synthesis of a library of 7,536 cyanimide screening compounds. A) DMSO solutions of 471 carboxylic acids (CA) and 16 amines (BB) were transferred from 384-well Echo plates and combined and treated with DIC/HOBt coupling reagent in DMSO in Echo plates, resulting in a compound library of 7,536 compounds. B) Overview of the in-house prepared 16 cyanimide-containing amines with their corresponding BB identification number. C) LC-MS analysis of the first-generation compound library in 384-well format. All compounds from two quadrants in the plate were analyzed as indicated. D) LC-MS analysis of the second-generation compound library in 1,536-well format. A representative number of compounds covering all rows and columns in the plate were analyzed as indicated. C,D) Colours represent product quality. Green: >75% product; yellow 25-75% product; red: 0-25% product; grey: not analyzed.

Since the in-plate synthesis was conducted in 1,536-well Echo plates, the compounds could readily be transferred from these plates into 1,536-well assay plates using acoustic liquid dispensing (Figure 2A) and subjected to HTS. We screened the whole 7,536 compound library against a panel of twelve enzymes, including three DUBs from the UCH family, two DUBs from the OTU family, five DUBs from the USP family, plus ISG15 protease USP18, and SUMO protease SENP1. Each compound was tested at a single dose of 1.25 µM, assuming full conversion (10 mM in DMSO of the formed products) during the in-plate synthesis, using the fluorogenic assay described above. For each compound, the percentage inhibition was calculated after normalizing the data to the positive (10 mM NEM, 100% inhibition) and negative (DMSO, 0% inhibition) controls. The heatmap in Figure 4A shows the percentage inhibition of each compound for each of the tested enzymes. The compounds are sorted by the amine building blocks (BB1-16), and within each BB, sorted by carboxylic acid (CA1-471), thereby providing a comprehensive and systematic overview of the overall inhibition by each BB and CA combination. Some interesting patterns can be obtained from this map. DUBs from the UCH family are hardly inhibited by these compounds, with the only exception of the BB10-derived compounds efficiently targeting UCHL1. Indeed, BB10 is the privileged scaffold for UCHL1 as it originates from UCHL1 inhibitor **5** (Figure 1). Other DUBs, like OTUB2, USP8, USP16, USP24, and SUMO protease SENP1, were generally not inhibited, although a few compounds showed activity against some of these enzymes. On the other hand, OTUD1 and USP32 show a much higher hit rate, and this is even more pronounced for USP30 and ISG15 protease USP18, each showing many active compounds distributed over several BB’s. Interestingly, BB1 (derived from compounds **1-4**, Figure 1), but not its enantiomer BB3, yielded many inhibitors for USP18, USP30 and USP32, indicating a stereochemical preference for the (*R*) configuration of the 3-aminopyrrolidine moiety. USP30 showed an elevated preference for BB9, which can be considered as an extended version of BB1, as nearly all compounds derived from this BB acted as USP30 inhibitors.

**Figure 4.**
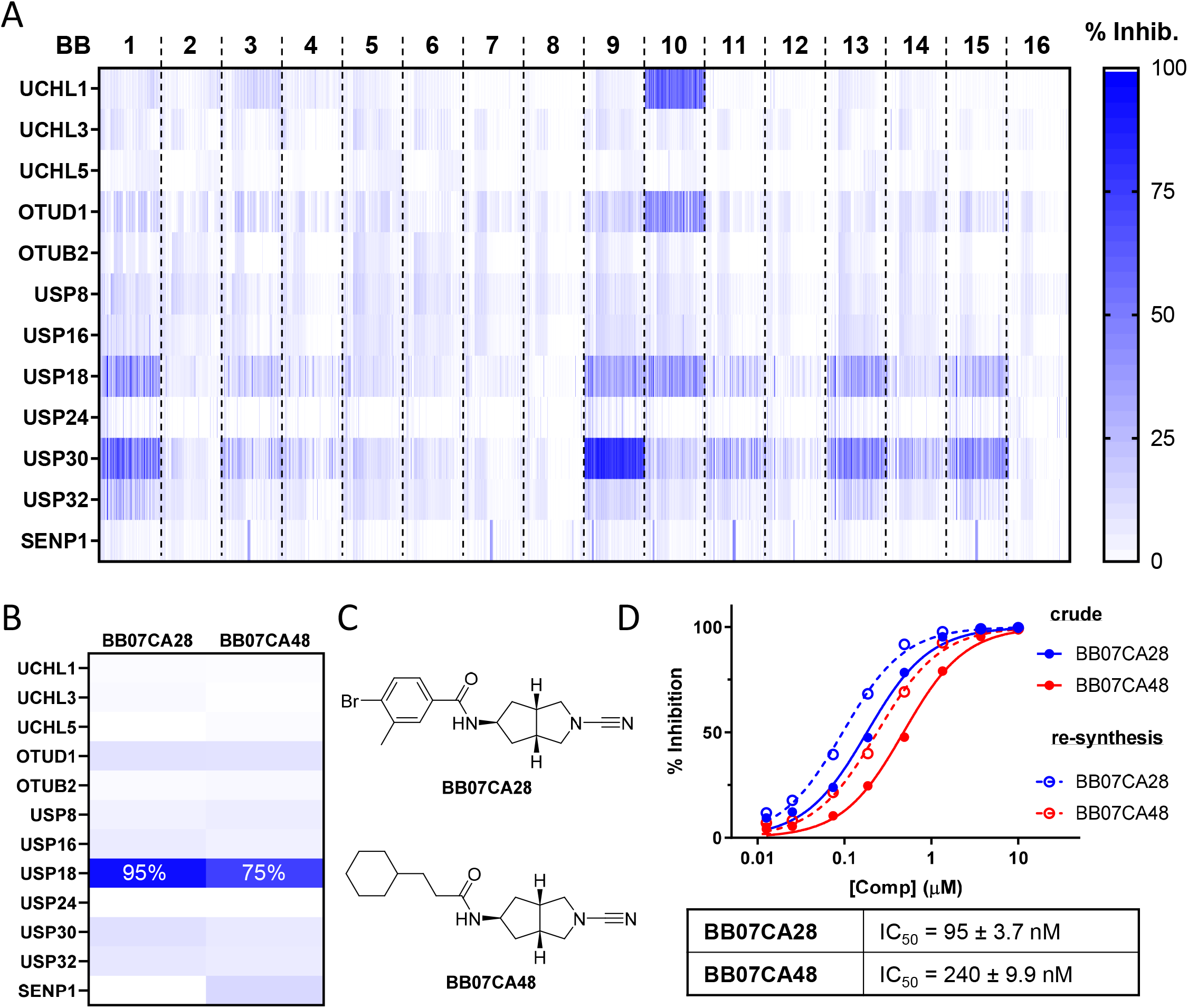
High-throughput screening results and USP18 hits validations. A) Heatmap representing the percentage inhibition of each of the 7,536 compounds (1.25 µM) on 12 Ub(-like) proteases. B) Percentage inhibition of two USP18 screening hits. C) Structures of the USP18 screening hits. D) IC_50_ curves showing the dose-dependent inhibition of USP18 by the two screening hits. Hits were cherry-picked directly from the library plate (crude, solid lines) and re-synthesized and purified (re-synthesis, dashed lines) for testing. IC_50_ values shown below the curves were obtained with the pure compounds.

To validate our method, we included the components forming compound **1** (Figure 1), here BB01 and CA066, in the high-throughput synthesis. The inhibition data for this crude compound, BB01CA066, (e.g. ∼90% inhibition of USP30 and USP18) were in line with the inhibition obtained for the pure compound **1** (Figure 1), indicating effective formation of the compound by the in-plate synthesis.

Instead of focusing solely on potency, we sought enzyme-specific inhibitors. In this regard, our attention was drawn towards two compounds that specifically inhibited ISG15 protease USP18, both derived from bicyclic cyanimide BB7, a BB that hardly yielded any actives overall. USP18 counteracts the host’s viral defence by deconjugating ISGylated proteins. As such, a USP18 inhibitor may serve as a starting point for the development of antiviral agents. The crude mixtures of BB07CA28 and BB07CA48 showed 95% and 75% inhibition of USP18 activity, respectively, and a negligible inhibition of all other enzymes tested (Figure 4B,C). To confirm that the inhibition could indeed be attributed to the compounds, we synthesized, purified and characterized both compounds, and subjected them to IC^50^ determinations, along with their corresponding in-plate prepared crudes. This showed both compounds to be potent USP18 inhibitors (Figure 4D) with IC_50_ values in the nanomolar range, and confirmed their selectivity over USP30 (IC_50_ of 13-17 µM, Figure S7). As expected, the pure compounds were more active than the crude ones, most likely because the actual concentration of the in-plate prepared compounds was lower than the assumed concentration (e.g. 10 mM stocks based on 100% conversion). Together, these results demonstrate the feasibility of our high-throughput synthesis-coupled-to-screening strategy.

We further explored the potential of our USP18 hits. With BB07CA28 being the more potent inhibitor, we initially conducted a small chemical optimization of this compound with respect to the substituents on the aromatic moiety. This yielded compound BB07CA902, having the bromide in BB07CA28 replaced by an isopropyl group, as the most potent USP18 inhibitor with an IC_50_ value of 35 nM (Figure 5A,B). Its stereoisomer, BB06CA902, showed a >300-fold reduction in potency, making it an appropriate negative control compound for future biological experiments. Based on previous data, these cyanimides likely bind to USP18 in a covalent manner. Indeed, intact protein mass spectrometry analysis confirmed the formation of a covalent stoichiometric USP18-BB07CA902 complex, showing a mass difference of 311 Da, which corresponds to the addition of exactly one molecule of BB07CA902 (Figure 5C, Figure S9). Enzyme kinetics measurements showed that BB07CA902 covalently inhibits USP18 with an enzyme inactivation rate (k_inact_) to inhibition constant (K_i_) ratio (k_inact_/K_i_) of 6,538 M^-1^s^-1^ (Figure 5D), thereby outperforming the initial screening hits BB07CA28 and BB07CA48. (Figure S9). These experiments also showed that the type of reducing agent present in the buffer (e.g. cysteine or TCEP) to keep the USP18 active site cysteine in a reduced state did not significantly affect the inhibitory potency.

**Figure 5.**
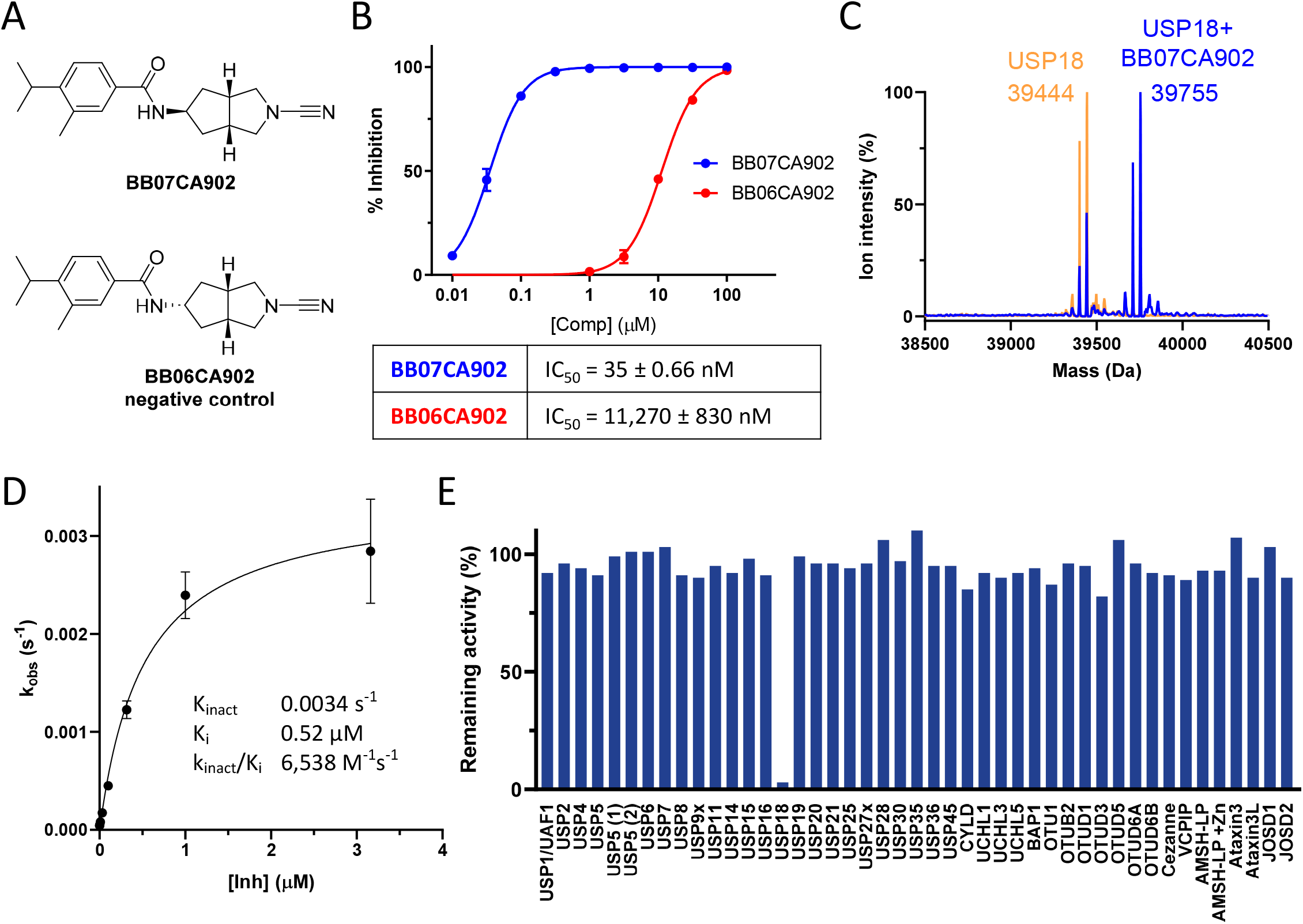
Biochemical characterization of improved USP18 inhibitor. A) Structures of improved USP18 inhibitor and inactive control compound. B) IC_50_ determination of USP18 inhibitor and its negative control. C) Deconvoluted mass spectra of USP18 before (orange) and after (blue) reaction with BB07CA902. D) *k*_*obs*_ plot to determine the kinetic parameters of covalent USP18 inhibition by BB07CA902. E) Commercial DUB inhibition screen with 0.25 µM BB07CA902. Data for USP18 inhibition was generated separately using a similar assay.

To gain further insights on the selectivity of BB07CA902 for USP18, the compound (0.25 µM concentration) was tested on a DUB panel comprising 41 different recombinant DUBs, including 22 from the structurally related USP family. Strikingly, none of these DUBs was inhibited by the compound, whereas it showed a near complete inhibition of USP18 at the same concentration (Figure 5E). In addition, the unrelated cysteine protease papain was only inhibited at an IC_50_ value of 27 µM, which was similar for both screening hits (Figure S7). Thus, BB07CA902 is a highly potent and selective covalent USP18 inhibitor.

To assess the cellular target engagement and cell permeability of these inhibitors, we performed activity-based protein profiling. Live HEK293T cells transiently overexpressing Flag-mUSP18 were treated with increasing concentrations of screening hits BB07CA28 and BB07CA48, optimized inhibitor BB07CA902 and its inactive control BB06CA902. Following washing and cell lysis, cell extracts were incubated with a Rhodamine-tagged murine ISG15 C-terminal domain propargylamide activity-based probe (Rho-mISG15ct-PA), which covalently labels the active site cysteine of active deISGylating enzymes.^[40]^ The labelled enzymes were resolved by SDS-PAGE and imaged by in-gel fluorescence, where inhibition of USP18 is reflected by disappearance of the USP18-probe-labelled bands around 55 kDa (Figure 6A, Figure S11A). All three inhibitors engaged with cellular mUSP18 in a dose-dependent manner, with BB07CA902 and BB07CA28 showing efficient inhibition already at 1 µM concentration. In line with the IC_50_ results, BB07CA28 was less potent, showing evident inhibition between 1-10 µM, while inhibition by control compound BB06CA902 was only observed at the highest dose of 100 µM. The in-cell selectivity of these inhibitors for USP18 over DUBs was confirmed in a similar ABPP experiment using a Rhodamine-tagged ubiquitin propargylamide probe (Rho-Ub-PA),^[41]^ which labels the active site cysteine of all active DUBs. None of the compounds showed any DUB inhibition up to 10 µM. Only at 100 µM, a few DUBs showed a diminished probe-labelled DUB band (Figure S10A).

**Figure 6.**
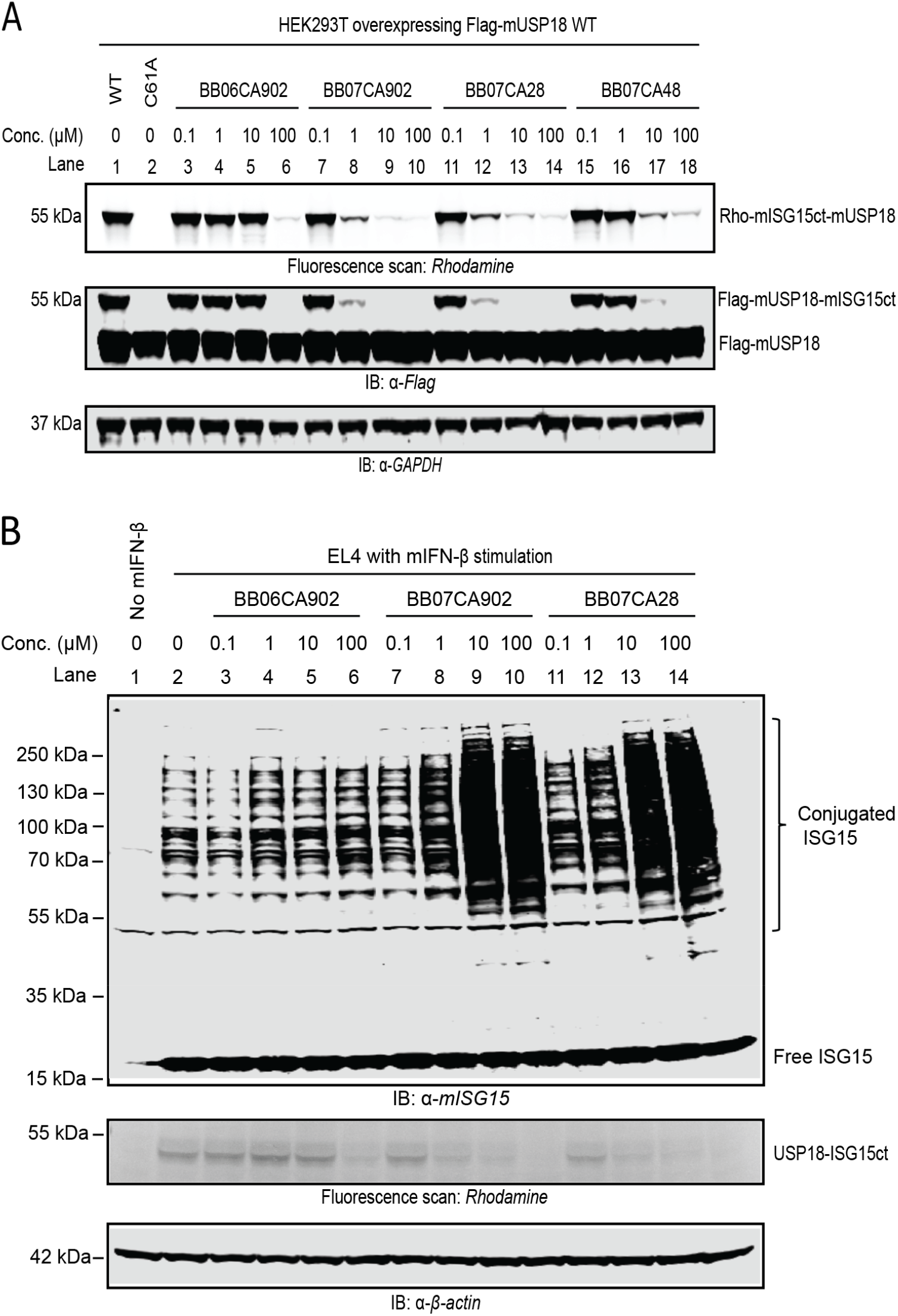
Inhibition of murine USP18 activity in living cells. A) Gel-based ABPP potency assessment using a Rho-mISG15ct-PA probe in HEK293T cells overexpressing Flag-mUSP18. At 24 hours following transfection, cells were incubated with DMSO or increasing concentrations of indicated inhibitors for 4 hours, lysed, and treated with Rho-mISG15ct-PA probe. Top panel: fluorescence scan. Middle panel: anti-Flag western blot. Bottom panel: anti-GAPDH western blot confirms equal loading. B) Accumulation of ISGylation in mouse IFN-β stimulated EL4 cells. At 24 hours following mouse IFN-β treatment, cells were incubated with DMSO or increasing concentrations of indicated inhibitors for 4 hours, lysed, and treated with Rho-mISG15ct-PA probe. Top panel: anti-mISG15 western blot. Middle panel: fluorescence scan. Bottom panel: anti-β-actin western blot confirms equal loading.

Ultimately, we were interested whether USP18 inhibitor treatment could result in a relevant phenotype in cells. USP18 is the major deISGylase to remove ISG15 from target proteins, and genetic inactivation of USP18 isopeptidase activity leads to enhanced ISGylation *in vivo* and in cells.^[25]^ We therefore examined whether the USP18 inhibitors enhance cellular ISGylation. Accordingly, EL4 cells were stimulated with IFN-β to induce the ISG15 machinery, prior to treatment with the inhibitors. After cell lysis and SDS-PAGE, ISGylated proteins were stained by immunoblotting with an ISG15 antibody (Figure 6B, top panel). Strikingly, treatment with BB07CA902 or BB07CA48 significantly elevated cellular ISGylation in a dose-dependent manner, and a maximum accumulation of ISGylation was reached at 10 µM, while the inactive control BB06CA902 hardly affected ISGylation, even at 100 µM. To confirm inhibition of endogenous USP18 in these experiments, samples were post-lysis treated with Rho-mISG15ct-PA probe. Fluorescence scanning after SDS-PAGE indeed showed a dose-dependent decrease of the endogenous USP18 probe-labelled bands upon treatment with both inhibitors (Figure 6B, middle panel, Figure S11B). Notably, inhibitor treatment of IFN-β stimulated EL4 cells did not lead to attenuated DUB activity as revealed by post-lysis incubation of the samples with Rho-Ub-PA DUB probe (Figure S10B). Altogether, these results show that selective inhibition of endogenous USP18 by BB07CA902 and BB07CA28 causes increased cellular ISGylation in IFN-stimulated EL4 cells.

## Conclusion

One of the prerequisites for a successful screening campaign is the availability of an appropriate compound library. Specifically, the use of a targeted compound library can increase the chances of obtaining suitable hit compounds, which would be especially beneficial for DUBs and UbL proteases, as they are known to be highly challenging to target. Here, we developed a high-throughput synthesis methodology that enabled the highly efficient preparation of a 7,536-member DUB-targeted cyanimide compound library for direct application in high-throughput screening. Key to our approach is the use of acoustic liquid transfer, using an Echo acoustic liquid handler, to rapidly combine small, microliter volumes of different reaction components in a 1,536-well plate. There are some major advantages of this ‘in-plate synthesis’ that makes it superior towards conventional methods. Each well in the 1,536-well plate represents a single reaction, which allows for a very rapid production of large compound sets. The volume per reaction is kept small (2.5 µL), resulting in very small required amounts of reactants and reagents and a concomitant cost reduction. As the compounds are directly prepared in Echo-compatible multi-well plates, further handling can readily be achieved by the Echo acoustic liquid dispenser, which we showcased by conducting LC-MS quality control and by screening the entire library against a panel of 12 Ub/UbL proteases.

In this study, we chose the amidation reaction for our high-throughput synthesis because this reaction is compatible with the cyanimide moiety in the target compounds and can be conducted in DMSO, the default solvent for acoustic dispensing and high-throughput screening. But in-plate synthesis is not limited to amidation, and other chemistries have been reported, including Ugi multicomponent reactions and catalysed reactions of diazo compounds, either in combination with Echo acoustic dispensing^[10-15]^ or using other means of liquid transfer.^[42,43]^ Although the in-plate strategy allows for a rapid production of thousands of compounds, a compromise had to be made on the compound purity as the scale (e.g. many compounds in small amounts) does not allow for purification. We tackled this problem by optimising reaction conditions, allowing for a high conversion with little side-product formation, and by using reagents that do not interfere with the screening assay. Despite these efforts, a complete formation of all library compounds is hard to accomplish, and indeed, our MS analysis revealed a ∼85% success rate over several plates. This underscores the importance of an appropriate validation of any screening hit to eliminate false positives. Nevertheless, the confirmation after re-synthesis of both USP18 hits (BB07CA28 and BB07CA48), as well as our results for USP30 inhibitor BB01CA066/compound **1**, taken along as control, prove the validity of our strategy.

In this study, we identified two screening hits for USP18, the main protease for Ub-like protein ISG15. Optimization of these hits resulted in the development of the nanomolar active, covalent inhibitor BB07CA902, which we showed to selectively target USP18 amongst a panel of 41 recombinant DUBs, including 22 from the structurally related USP family. This USP18 selectivity was also observed in cells, and we showed a significant increase in cellular protein ISGylation by inhibiting USP18. ISGylation plays a critical role in antiviral response by modifying either viral or host proteins.^[44]^ Genetic inactivation of USP18 enzymatic activity in cells or (USP18^C61A/C61A^ knock-in) mice leads to enhanced ISGylation upon IFN-β stimulation, and elevated ISGylation is accompanied by increased viral resistance against vaccinia virus, influenza B virus, and coxsackievirus infections.^[25,26]^ Moreover, the level of ISG15 and ISGylation is found to be enhanced in many cancer cells; however, the role of ISGylation in anti-tumor or pro-tumor function in cell-based studies is controversial.^[45,46]^ An *in vivo* study using immunocompetent mice provided evidence for a tumor-suppressor function of the ISG15-ISGylation network in breast cancer. Polyomavirus mT-induced breast tumor growth was found to be enhanced in ISG15 E1 enzyme Ube1l-deleted (KO) mice, and suppressed in USP18^C61A/C61A^ mice.^[47]^ These studies indicate that USP18 is a promising target in antiviral and potential anti-tumor treatment. Our USP18 inhibitors can be a good starting point to test USP18-targeted therapeutic hypotheses in mouse models, and serve as a valuable research tool to further investigate USP18’s function, regulation, and biochemical mechanisms.

In summary, we developed a highly efficient method for the high-throughput synthesis of large compound libraries, making use of acoustic dispensing, which can be directly used for high-throughput screening. The as such created 7,536-member DUB-targeted cyanimide library can easily be expanded, simply by increasing the number of amines and carboxylic acids, but already yielded potent and selective inhibitors for USP18, which were shown to increase ISGylation in cells. We believe that our high-throughput synthesis-to-screening method will open new opportunities for the swift preparation and screening of targeted compound libraries, which will accelerate future drug development programs.

## Supporting Information

Supporting Information containing Supplementary Figures, experimental procedures, compound synthesis and characterization data is provided as a separate file (PDF).

## Acknowledgements

Research reported in this publication was supported by the Innovative Medicines Initiative 2 (IMI2) Joint Undertaking under grant agreement no. 875510 (EUbOPEN project) and by Oncode Accelerator, a Dutch National Growth Fund project under grant number NGFOP2201 (P.P.G.). J.G. was supported by the European Cooperation in Science and Technology (COST) with a short-term scientific mission grant (STSM, Action CA15138). A.S. was supported by the Institute of Chemical Immunology (grant no. ICI00026).

## References

[1] R. Macarron, M. N. Banks, D. Bojanic, D. J. Burns, D. A. Cirovic, T. Garyantes, D. V. Green, R. P. Hertzberg, W. P. Janzen, J. W. Paslay, U. Schopfer, G. S. Sittampalam, Nat. Rev. Drug Discov. 2011, 10, 188–195.

[2] S. K. Ashenden, Meth. Enz. 2018, 610, 73–96.

[3] D. A. Erlanson, S. W. Fesik, R. E. Hubbard, W. Jahnke, H. Jhoti, Nat. Rev. Drug Discov. 2016, 15, 605–619.

[4] J. L. Medina-Franco, K. Martinez-Mayorga, N. Meurice, Exp. Opin. Drug Discov. 2014, 9, 151–165.

[5] S. L. Schreiber, Science 2000, 287, 1964–1969.

[6] R. A. Goodnow, Jr., C. E. Dumelin, A. D. Keefe, Nat. Rev. Drug Discov. 2017, 16, 131–147.

[7] R. Liu, X. Li, K. S. Lam, Curr. Opin. Chem. Biol. 2017, 38, 117–126.

[8] G. Schneider, Nat. Rev. Drug Discov. 2018, 17, 97–113.

[9] R. Ellson, M. Mutz, B. Browning, L. Lee, M. F. Miller, R. Papen, JALA: J. Assoc. Lab. Autom. 2003, 8, 29–34.

[10] M. P. Plesniak, E. K. Taylor, F. Eisele, C. Kourra, I. N. Michaelides, A. Oram, J. Wernevik, Z. S. Valencia, H. Rowbottom, N. Mann, L. Fredlund, V. Pivnytska, A. Novén, M. Pirmoradian, T. Lundbäck, R. I. Storer, M. Pettersson, G. M. De Donatis, M. Rehnström, ACS Med. Chem. Lett. 2023, 14, 1882–1890.

[11] G. Sangouard, A. Zorzi, Y. Wu, E. Ehret, M. Schüttel, S. Kale, C. Díaz-Perlas, J. Vesin, J. Bortoli Chapalay, G. Turcatti, C. Heinis, Angew. Chem. Int. Ed. Engl. 2021, 60, 21702–21707.

[12] S. Shaabani, R. Xu, M. Ahmadianmoghaddam, L. Gao, M. Stahorsky, J. Olechno, R. Ellson, M. Kossenjans, V. Helan, A. Dömling, Green Chem. 2019, 21, 225–232.

[13] C. E. Arcadia, E. Kennedy, J. Geiser, A. Dombroski, K. Oakley, S. L. Chen, L. Sprague, M. Ozmen, J. Sello, P. M. Weber, S. Reda, C. Rose, E. Kim, B. M. Rubenstein, J. K. Rosenstein, Nat. Commun. 2020, 11, 691.

[14] F. Sutanto, S. Shaabani, C. G. Neochoritis, T. Zarganes-Tzitzikas, P. Patil, E. Ghonchepour, A. Dömling, Sci. Adv. 2021, 7.

[15] Y. Wang, S. Shaabani, M. Ahmadianmoghaddam, L. Gao, R. Xu, K. Kurpiewska, J. Kalinowska-Tluscik, J. Olechno, R. Ellson, M. Kossenjans, V. Helan, M. Groves, A. Dömling, ACS Cent. Sci. 2019, 5, 451–457.

[16] K. N. Swatek, D. Komander, Cell Res. 2016, 26, 399–422.

[17] M. J. Clague, C. Heride, S. Urbé, Trends Cell Biol. 2015, 25, 417–426.

[18] T. Sun, Z. Liu, Q. Yang, Mol. Canc. 2020, 19, 146.

[19] M. F. Schmidt, Z. Y. Gan, D. Komander, G. Dewson, Cell Death Differ. 2021, 28, 570–590.

[20] L. L. Zheng, L. T. Wang, Y. W. Pang, L. P. Sun, L. Shi, Eur. J. Med. Chem. 2024, 266, 116161.

[21] D. X. Zhang, D. E. Zhang, J. Interferon Cytokine Res. 2011, 31, 119–130.

[22] A. Basters, P. P. Geurink, F. El Oualid, L. Ketscher, M. S. Casutt, E. Krause, H. Ovaa, K. P. Knobeloch, G. Fritz, FEBS J. 2014, 281, 1918–1928.

[23] O. A. Malakhova, K. I. Kim, J. K. Luo, W. G. Zou, K. G. S. Kumar, S. Y. Fuchs, K. Shuai, D. E. Zhang, EMBO J. 2006, 25, 2358–2367.

[24] K. I. Arimoto, S. Lochte, S. A. Stoner, C. Burkart, Y. Zhang, S. Miyauchi, S. Wilmes, J. B. Fan, J. J. Heinisch, Z. Li, M. Yan, S. Pellegrini, F. Colland, J. Piehler, D. E. Zhang, Nat. Struct. Mol. Biol. 2017, 24, 279–289.

[25] L. Ketscher, R. Hannss, D. J. Morales, A. Basters, S. Guerra, T. Goldmann, A. Hausmann, M. Prinz, R. Naumann, A. Pekosz, O. Utermohlen, D. J. Lenschow, K. P. Knobeloch, Proc. Natl. Acad. Sci. USA 2015, 112, 1577–1582.

[26] M. Kespohl, C. Bredow, K. Klingel, M. Voß, A. Paeschke, M. Zickler, W. Poller, Z. Kaya, J. Eckstein, H. Fechner, J. Spranger, M. Fähling, E. K. Wirth, L. Radoshevich, F. Thery, F. Impens, N. Berndt, K.-P. Knobeloch, A. Beling, Sci. Adv. 2020, 6, eaay1109.

[27] D. D. Nekrasov, Russ. J. Org. Chem. 2004, 40, 1387–1402.

[28] D. Laine, M. Palovich, B. McCleland, E. Petitjean, I. Delhom, H. B. Xie, J. H. Deng, G. L. Lin, R. Davis, A. Jolit, N. Nevins, B. G. Zhao, J. Villa, J. Schneck, P. McDevitt, R. Midgett, C. Kmett, S. Umbrecht, B. Peck, A. B. Davis, D. Bettoun, ACS Med. Chem. Lett. 2011, 2, 142–147.

[29] A. Jones, M. Kemp, M. Stockley, K. Gibson, G. Whitlock, WO2016/156816 A1 2016.

[30] A. Jones, M. I. Kemp, M. L. Stockley, K. R. Gibson, G. A. Whitlock, A. Madin, WO2016/046530 A1 2016.

[31] N. Panyain, A. Godinat, T. Lanyon-Hogg, S. Lachiondo-Ortega, E. J. Will, C. Soudy, M. Mondal, K. Mason, S. Elkhalifa, L. M. Smith, J. A. Harrigan, E. W. Tate, J. Am. Chem. Soc. 2020, 142, 12020–12026.

[32] R. Kooij, S. H. Liu, A. Sapmaz, B. T. Xin, G. M. C. Janssen, P. A. van Veelen, H. Ovaa, P. ten Dijke, P. P. Geurink, J. Am. Chem. Soc. 2020, 142, 16825–16841.

[33] C. Grethe, M. Schmidt, G. M. Kipka, R. O’Dea, K. Gallant, P. Janning, M. Gersch, Nat. Commun. 2022, 13, 5950.

[34] M. Schmidt, C. Grethe, S. Recknagel, G. M. Kipka, N. Klink, M. Gersch, Angew. Chem. Int. Ed. Engl. 2024, 63, e202318849.

[35] A. Jones, M. I. Kemp, M. L. Stockley, M. D. Woodrow, WO2017/158381 A1, 2017.

[36] M. I. Kemp, M. L. Stockley, M. D. Woodrow, A. Jones, WO2017/149313 A1, 2017.

[37] J. Yang, E. L. Weisberg, X. X. Liu, R. S. Magin, W. C. Chan, B. Hu, N. J. Schauer, S. Z. Zhang, I. Lamberto, L. Doherty, C. C. Meng, M. Sattler, L. Cabal-Hierro, E. Winer, R. Stone, J. A. Marto, J. D. Griffin, S. J. Buhrlage, Leukemia 2022, 36, 210–220.

[38] D. Conole, F. Y. Cao, C. W. A. Ende, L. Xue, S. Kantesaria, D. Kang, J. Jin, D. Owen, L. Lohr, M. Schenone, J. D. Majmudar, E. W. Tate, Angew. Chem. Int. Ed. Engl. 2023, 62, e202311190.

[39] W. C. Chan, X. X. Liu, R. S. Magin, N. M. Girardi, S. B. Ficarro, W. Y. Hu, M. T. Guzman, C. A. Starnbach, A. Felix, G. Adelmant, A. C. Varca, B. Hu, A. S. Bratt, E. DaSilva, N. J. Schauer, I. J. Maisonet, E. K. Dolen, A. X. Ayala, J. A. Marto, S. J. Buhrlage, Nat. Commun. 2023, 14.

[40] A. Basters, P. P. Geurink, A. Rocker, K. F. Witting, R. Tadayon, S. Hess, M. S. Semrau, P. Storici, H. Ovaa, K. P. Knobeloch, G. Fritz, Nat. Struct. Mol. Biol. 2017, 24, 270–278.

[41] R. Ekkebus, S. I. van Kasteren, Y. Kulathu, A. Scholten, I. Berlin, P. P. Geurink, A. de Jong, S. Goerdayal, J. Neefjes, A. J. Heck, D. Komander, H. Ovaa, J. Am. Chem. Soc. 2013, 135, 2867–2870.

[42] G. Karageorgis, S. Warriner, A. Nelson, Nat. Chem. 2014, 6, 872–876.

[43] A. Osipyan, S. Shaabani, R. Warmerdam, S. V. Shishkina, H. Boltz, A. Dömling, Angew. Chem. Int. Ed. Engl. 2020, 59, 12423–12427.

[44] Y.-C. Perng, D. J. Lenschow, Nat. Rev. Microbiol. 2018, 16, 423–439.

[45] H. G. Han, H. W. Moon, Y. J. Jeon, Cancer Lett. 2018, 438, 52–62.

[46] C. Zuo, X. Sheng, M. Ma, M. Xia, L. Ouyang, Oncotarget 2016, 7, 74393–74409.

[47] J. B. Fan, S. Miyauchi, H. Z. Xu, D. Liu, L. J. Y. Kim, C. Burkart, H. Cheng, K. I. Arimoto, M. Yan, Y. Zhou, B. Győrffy, K. P. Knobeloch, J. N. Rich, H. Cang, X. D. Fu, D. E. Zhang, Cancer Discov. 2020, 10, 382–393.

