## Supporting Information for "High-Throughput Synthesis and Screening of a Cyanimide Library Identifies Selective Inhibitors of ISG15-specific Protease USP18"

**Contents**

Figure S1. $IC_{50}$ determination of compounds **1-5**……………………………………………..S3

Figure S2. Full spectrum LC-MS analysis of the model reaction shown…………………...S4

Figure S3. LCMS analysis of the first generation compound library……………………..…S5

Figure S4. Plate outline of all 1,536 well synthesis plates…………………………………...S6

Figure S5. LCMS analysis of plate NCN-4……………………………………………………S7

Figure S6. LCMS analysis of plate NCN-5……………………………………………………S8

Figure S7. $IC_{50}$ determination of hit compounds for USP30 and papain…………………...S9

Figure S8. Determination of $k_{inact}$ and $K_i$ values for hit compounds on mUSP18…………..S10

Figure S9. LC-MS analysis of USP18-BB07CA902 covalent complex…………………...S11

Figure S10. Gel-based ABPP selectivity assessment using Rho-Ub-PA…………………...S12

Figure S11. Full image scan of Rho-mISG15ct-PA probe labelled enzymes……………….S13

Synthesis……………………………………………………………………………………...S14

Echo synthesis and analysis…………………………………………………………………..S28

Biochemistry methods………………………………………………………………………S32

References……………………………………………………………………………………S37

NMR and LC-MS spectra……………………………………………………………………S38

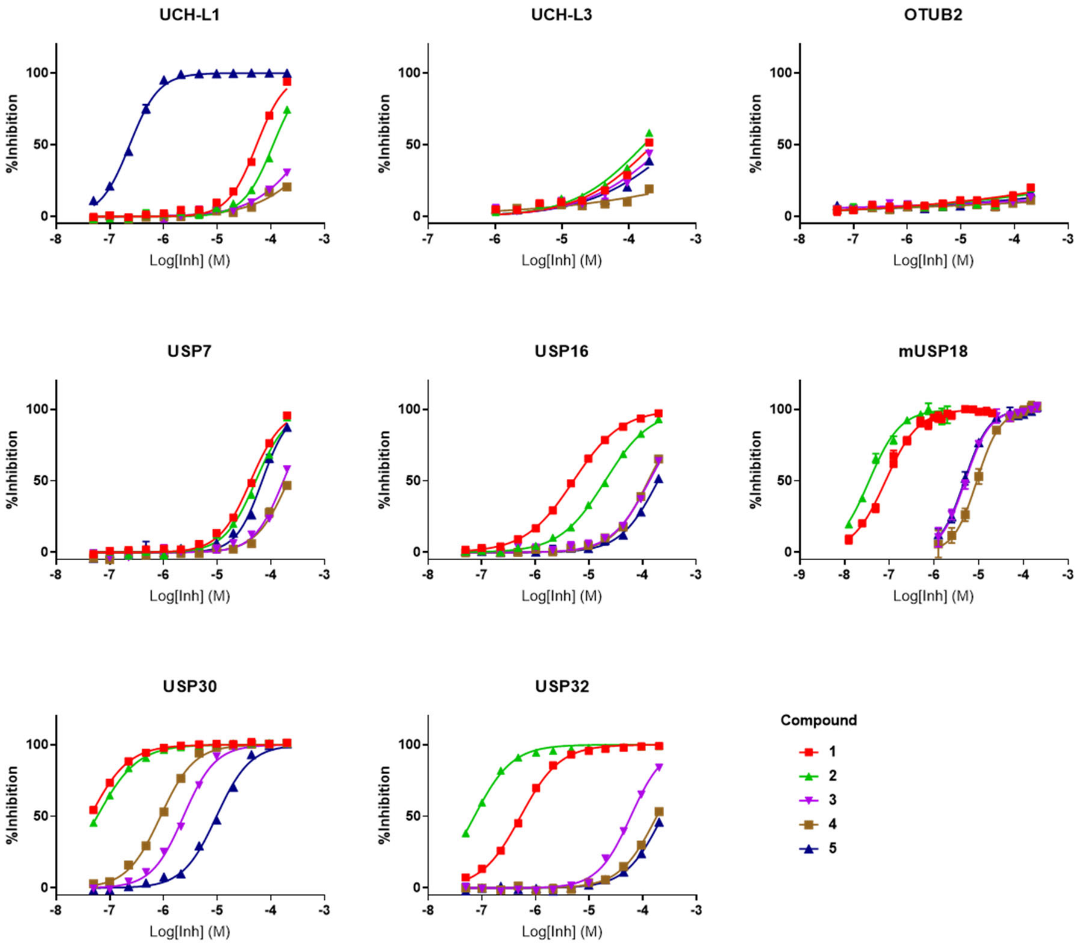


**Figure S1. IC$_{50}$ determination of compounds 1-5 for different Ub(like) proteases.** IC$_{50}$ values are shown in Figure 1.

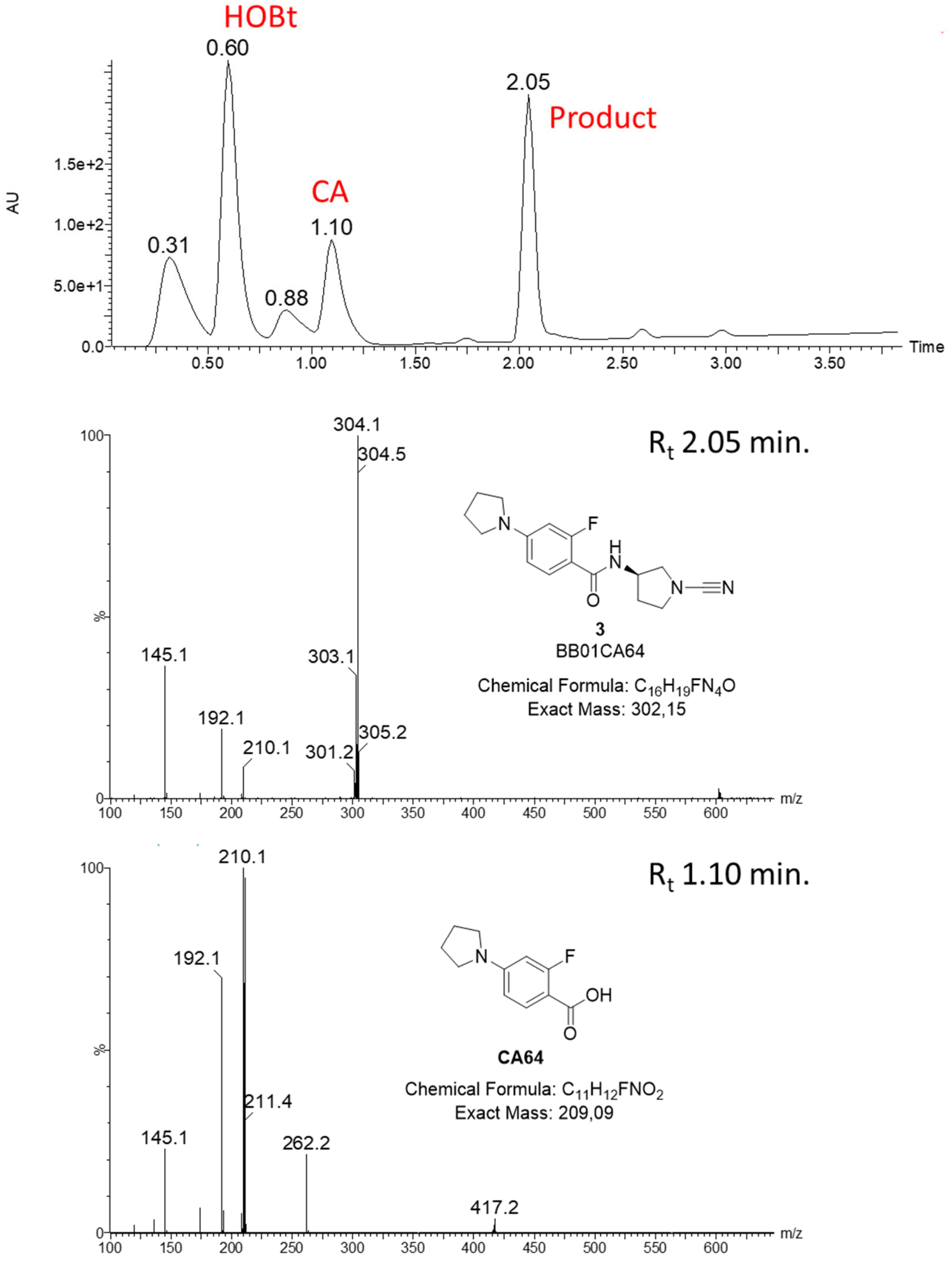


**Figure S2. Full spectrum LC-MS analysis of the model reaction shown in Figure 2C,D.**

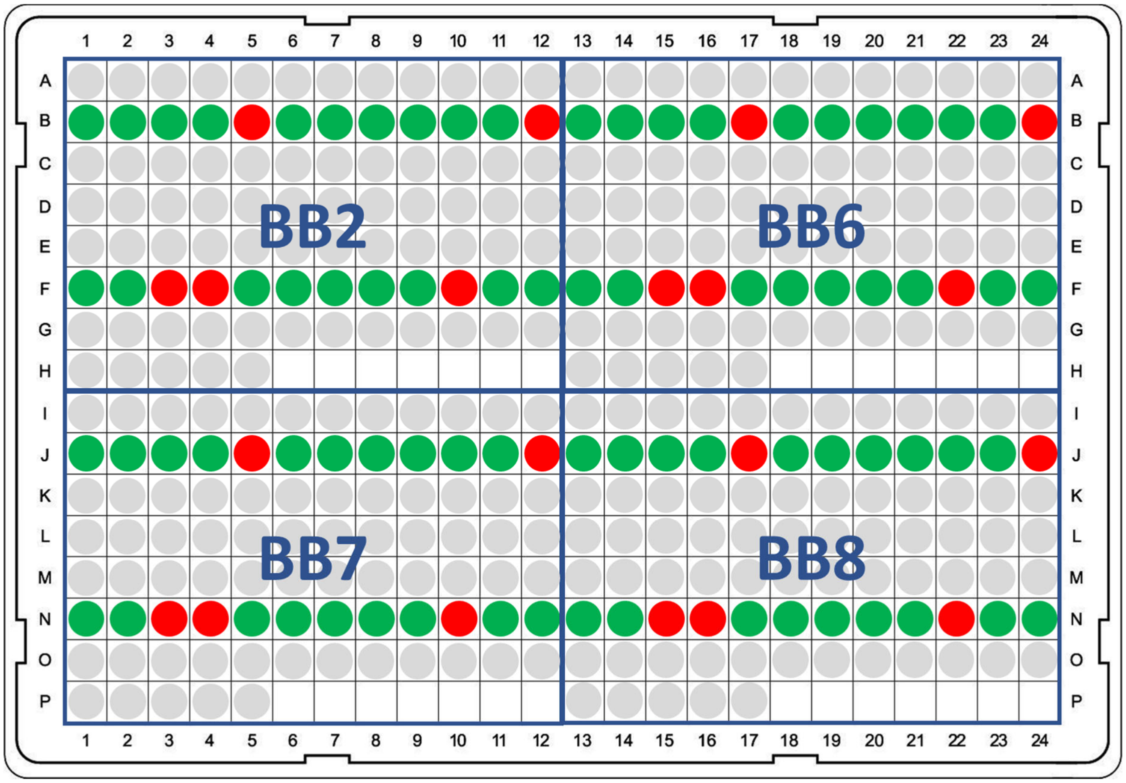


**Figure S3. Plate layout and LCMS analysis of the first generation compound library in 384-well format.** All compounds from the indicated wells were analyzed. Colors represent product quality. Green: >75% product; yellow 25-75% product; red: 0-25% product; grey: not analyzed; white: empty wells.

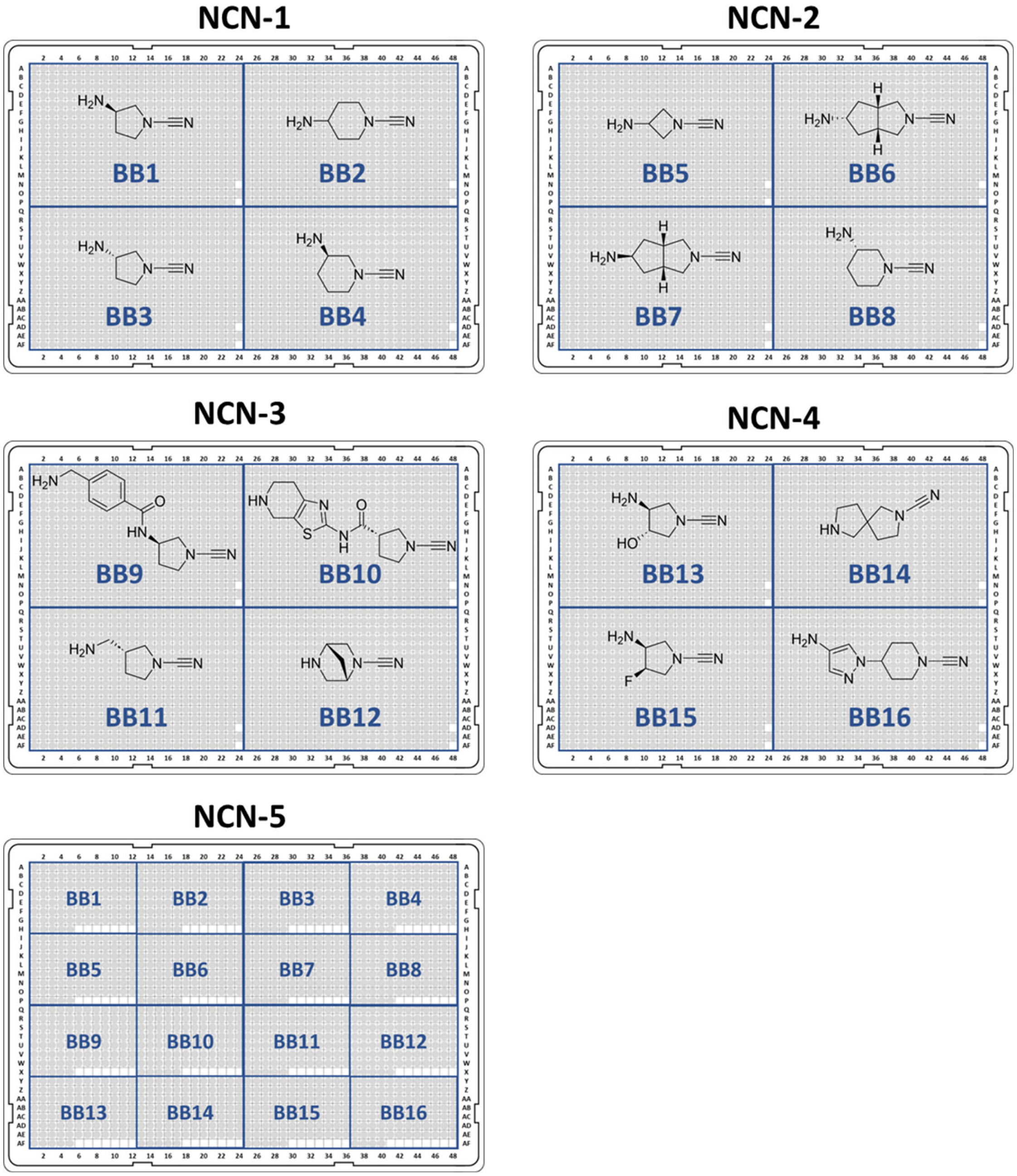


**Figure S4. Plate outline of all 1,536-well synthesis plates.** Plate numbers are indicated and correspond to the supplementary data file. Locations of amine BB's are indicated. Plates NCN-1-4 were synthesized with 382 Enamine CA's, plate NCN-5 was synthesized with 89 in-house selected CA's. Grey-colored wells contain compounds. White-colored wells are empty.

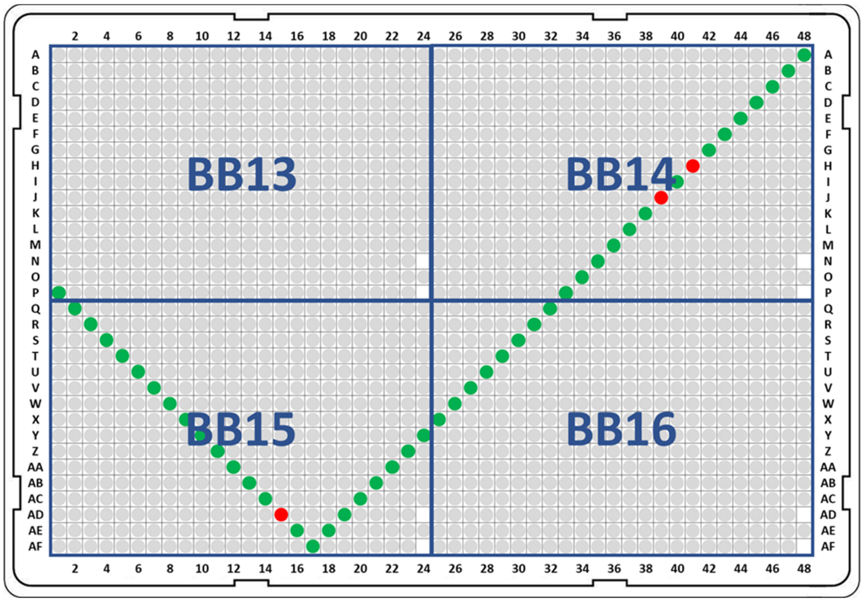


**Figure S5. LCMS analysis of the second generation compound library in 1,536-well format – plate NCN-4.** A representative number of compounds covering all rows and columns in the plate were analyzed as indicated. Colors represent product quality. Green: >75% product; yellow 25-75% product; red: 0-25% product; grey: not analyzed.

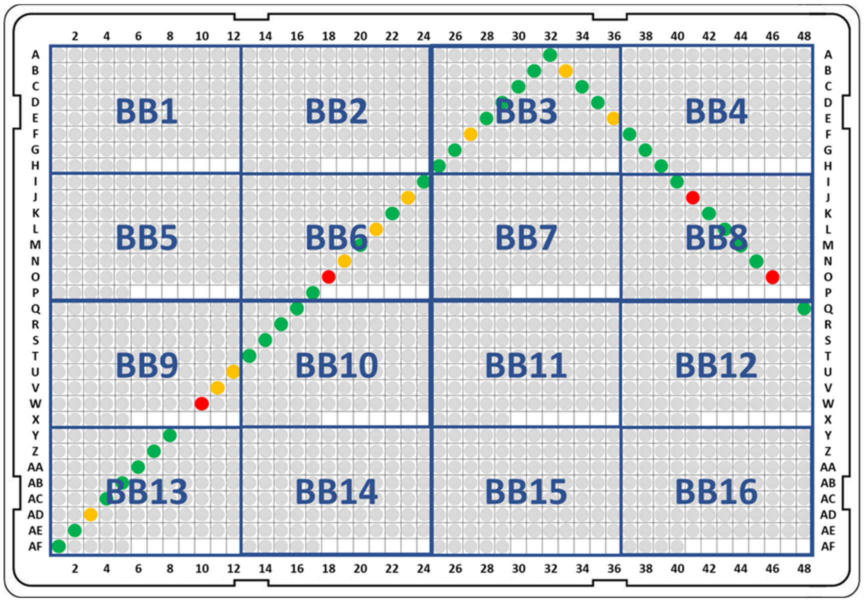


**Figure S6. LCMS analysis of the second generation compound library in 1,536-well format – plate NCN-5.** A representative number of compounds covering all rows and columns in the plate were analyzed as indicated. Colors represent product quality. Green: >75% product; yellow 25-75% product; red: 0-25% product; grey: not analyzed.

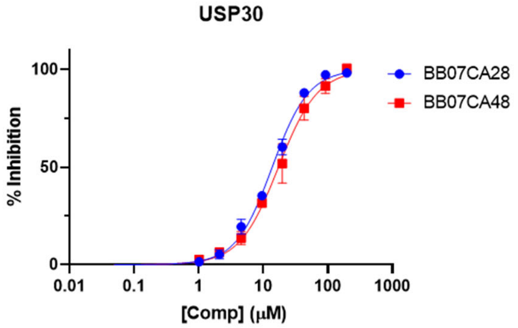


| Compound | $IC_{50}$ (μM) |
|---|---|
| BB07CA28 | 13.7 ± 0.74 |
| BB07CA48 | 17.0 ± 1.0 |

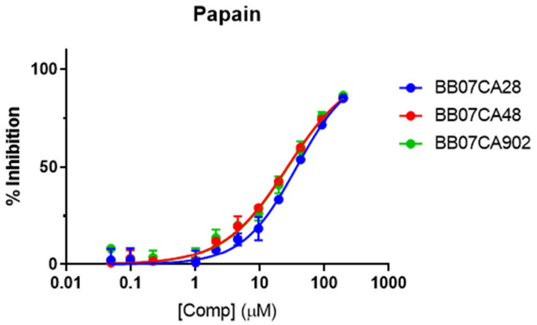


| Compound | $IC_{50}$ (μM) |
|---|---|
| BB07CA28 | 37.2 ± 2.18 |
| BB07CA48 | 26.6 ± 1.23 |
| BB07CA902 | 27.3 ± 1.93 |

**Figure S7. $IC_{50}$ determination of hit compounds for USP30 and papain.**

**A**

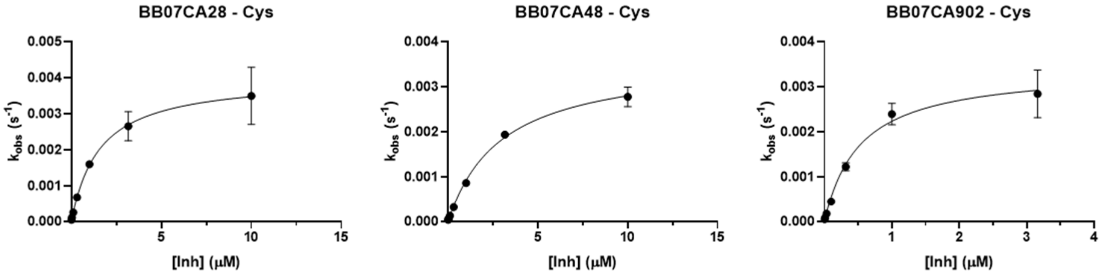


| Compound | $k_{inact}$ ($s^{-1}$) | $K_i$ (μM) | $k_{inact}/K_i$ ($M^{-1}s^{-1}$) |
|---|---|---|---|
| BB07CA28 | 0.0040 ± 0.00050 | 1.54 ± 0.061 | 2597 |
| BB07CA48 | 0.0036 ± 0.00099 | 2.94 ± 0.21 | 1224 |
| BB07CA902 | 0.0034 ± 0.00014 | 0.52 ± 0.066 | 6538 |

**B**

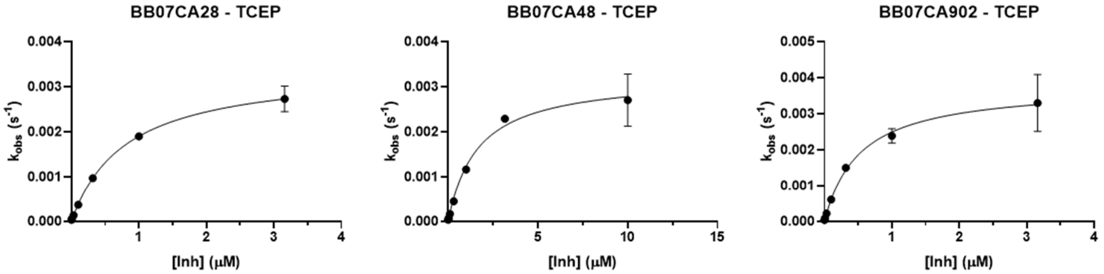


| Compound | $k_{inact}$ ($s^{-1}$) | $K_i$ (μM) | $k_{inact}/K_i$ ($M^{-1}s^{-1}$) |
|---|---|---|---|
| BB07CA28 | 0.0034 ± 0.00045 | 0.80 ± 0.029 | 4250 |
| BB07CA48 | 0.0032 ± 0.00013 | 1.63 ± 0.19 | 1963 |
| BB07CA902 | 0.0038 ± 0.00092 | 0.52 ± 0.039 | 7308 |

**Figure S8. Determination of $k_{inact}$ and $K_i$ values for hit compounds on mUSP18.** The assay was performed in the presence of A) 2 mM cysteine or B) 1 mM TCEP as reducing agent.

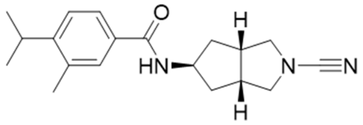


**BB07CA902**

Chemical Formula: $C_{19}H_{25}N_3O$

Exact Mass: 311,2

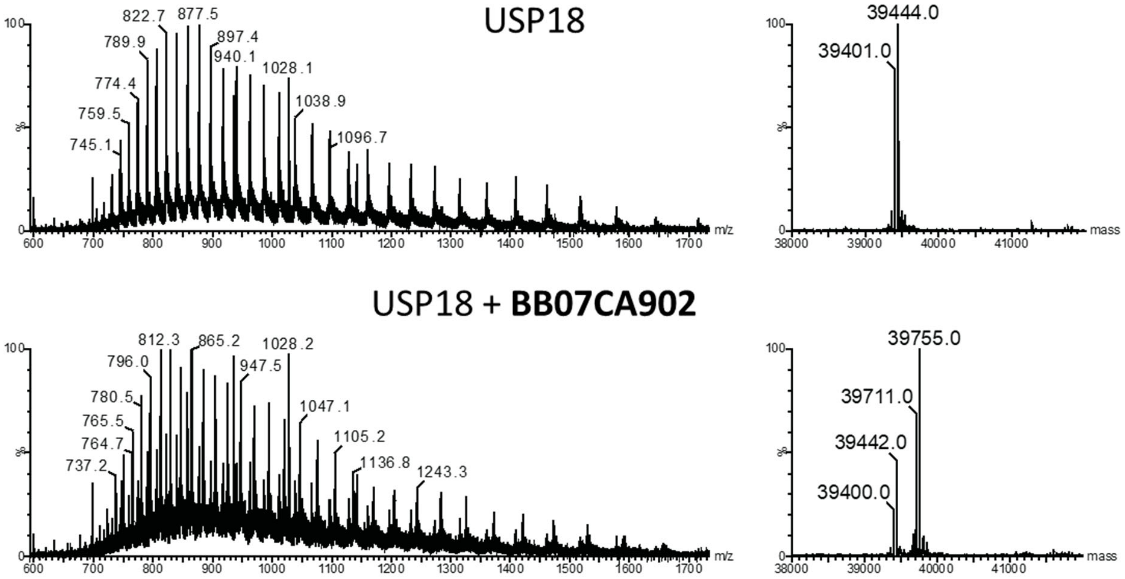


**Figure S9. LC-MS analysis confirms formation of a covalent complex of USP18 with one molecule of BB07CA902.** Ion traces (left) and deconvoluted mass (right) of USP18 before (top) and after (bottom) incubation with BB07CA902. The mass difference of 311 Da corresponds to the addition of exactly one molecule of BB07CA902. This figure is connected to Figure 5D.

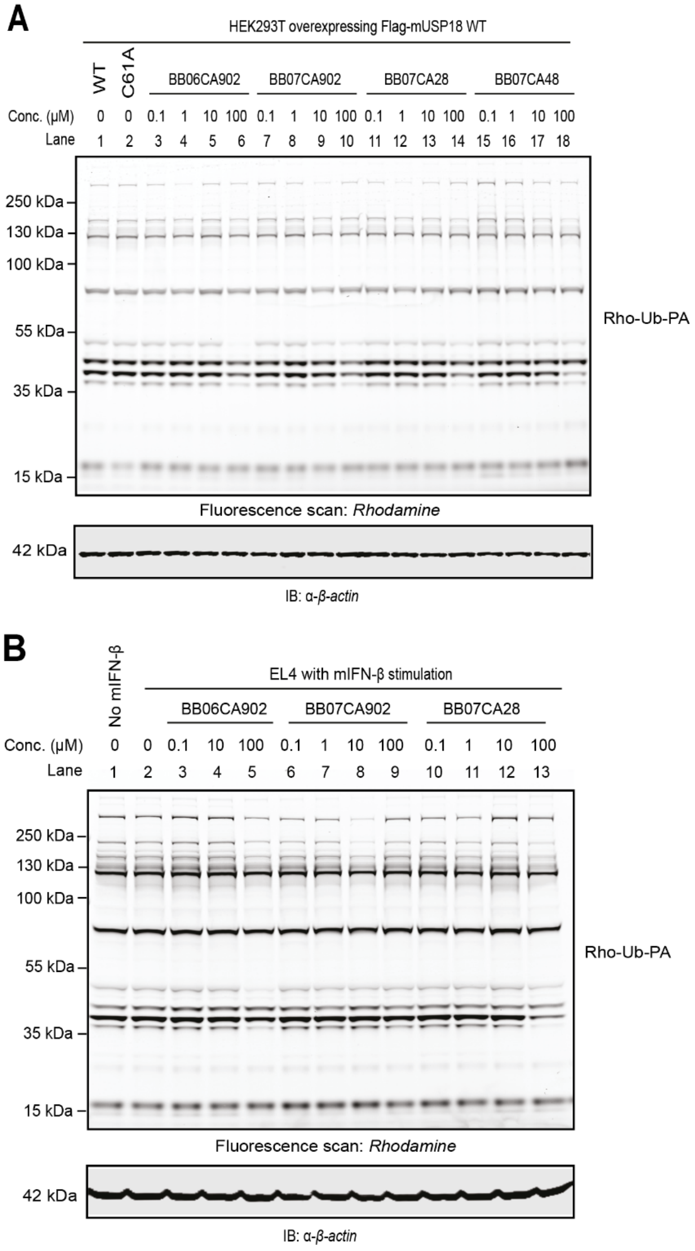


**Figure S10. Gel-based ABPP selectivity assessment using Rho-Ub-PA probe.** A) ABPP in HEK293 cells overexpressing Flag-mUSP18. At 24 hours following transfection, cells were incubated with DMSO or increasing concentrations of indicated inhibitors for 4 hours, lysed, and treated with Rho-Ub-PA. Top: fluorescence scan of Rho-Ub-PA probe labelled human DUBs. Bottom: anti-β-actin western blot confirming equal loading of each sample. B) ABPP in mIFN-β stimulated mouse EL4 cells. At 24 hours following transfection, cells were incubated with DMSO or increasing concentrations of indicated inhibitors for 4 hours, lysed and treated with Rho-Ub-PA. Top: fluorescence scan of Rho-Ub-PA probe labelled mouse DUBs. Bottom: anti-β-actin western blot confirming equal loading of each sample.

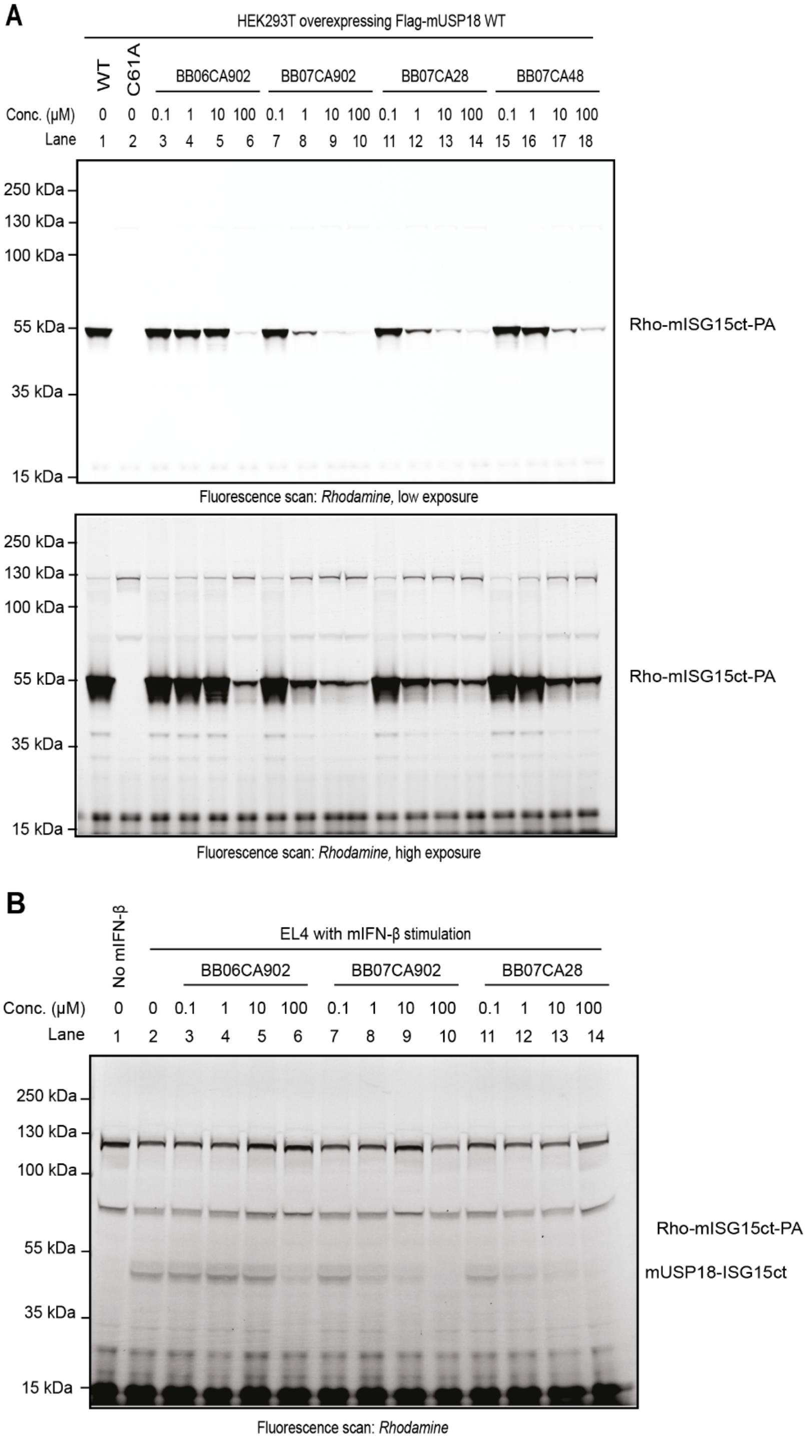


**Figure S11. Full image scan of Rho-mISG15ct-PA probe labelled enzymes.** A) Gel-based ABPP assessment using a Rho-mISG15ct-PA probe in HEK293 cells overexpressing Flag-mUSP18, corresponding to top panel in Figure 6A. Top panel: fluorescence scan in low exposure; bottom panel: fluorescence scan in high exposure. B) Gel-based ABPP assessment using a Rho-mISG15ct-PA probe in mIFN-β stimulated mouse EL4 cells, corresponding to middle panel in Figure 6B.

#### Synthesis

##### General

**Reagents, solvents and solutions.** General reagents were purchased from Sigma-Aldrich, Acros and Combi-Blocks, Inc. and were used as received. Solvents were purchased from VWR, Biosolve or Sigma-Aldrich. Thin Layer Chromatography (TLC) was performed on Merck aluminum sheets (pre-coated with silica gel 60 $F_{254}$). Compounds were visualized by UV adsorption (254 nm) and by using a solution of $KMnO_4$ (7.5 g $L^{-1}$) and $K_2CO_3$ (50 g $L^{-1}$) in $H_2O$ or a solution of ninhydrin (15 g $L^{-1}$) in 3% AcOH/EtOH v/v.

**Purification Techniques.** Compounds (unless stated otherwise) were purified on a Büchi Sepacore automatic flash chromatography system X10/X50. The Büchi Sepacore system was equipped with two Büchi pump modules C-605, a Büchi control unit C-620, Büchi fraction collector C-660 and a Büchi UV Photometer C-640. The silica columns were purchased at GraceResolv™ and were packed with a grade of Davisil® silica.

**Instrumentation for Compound Characterization**. NMR spectra ($^1H$, $^{13}C$) were recorded on a Bruker Ultrashield 300 MHz spectrometer at 298 K. Resonances are indicated with symbols ‘d’ (doublet), ‘s’ (singlet), ‘t’ (triplet) and ‘m’ (multiplet). Chemical shifts (δ) are given in ppm relative to $CDCl_3$, DMSO-$d_6$ or $CD_3OD$ as an internal standard and coupling constants ($J$) are quoted in hertz (Hz). LC-MS measurements were performed on an LC-MS system equipped with a Waters 2795 Separation Module (Alliance HT), a Waters 2996 Photodiode Array Detector (190–750 nm), an Xbridge C18 column (2.1 × 100 mm, 3.5 μm) and an LCT ESI-Orthogonal Acceleration Time of Flight Mass Spectrometer. Samples were run using 2 mobile phases: A = 1% $CH_3CN$ and 0.1% formic acid in $H_2O$ and B = 1% $H_2O$ and 0.1% formic acid in $CH_3CN$. Data processing was performed using Waters MassLynx Mass Spectrometry Software 4.1. LC-MS Program: Waters Xbridge C18 column (2.1 × 100 mm, 3.5 μm); flow rate = 0.4 mL $min^{-1}$, runtime = 13 min, column T = 40 °C, mass detection: 100–1500 Da. Gradient: 0–0.4 min: 5% B; 0.4–9.0 min: 5% → 95% B; 9.0–11.2 min: 95% B; 11.2–11.3 min: 95% → 5% B; 11.3–13.00 min: 5% B. Electrospray Ionization (ESI) high-resolution mass spectrometry was carried on a Waters XEVO-G2 XS Q-TOF mass spectrometer equipped with an electrospray ion source in positive mode (capillary voltage: 3.0 kV, desolvation gas flow: 900 L $h^{-1}$, temperature: 60 °C) with a resolution R = 22,000 using 200 pg μ$L^{-1}$ Leu-Enk ($m/z$ = 556.2771) as a “lock mass”. Samples were run using 2 mobile phases: A = 0.1% formic acid in $H_2O$ and B = 0.1% formic acid in $CH_3CN$ on a Waters Acquity UPLC BEH C18 column (2.1 × 50 mm, 1.7 μm); flow rate = 0.6 mL $min^{-1}$, runtime = 3.00 min, column T = 60 °C, mass detection: 50–1500 Da. Gradient: 0–0.15 min: 2% B; 0.15–1.85 min: 2% → 100% B; 1.85–2.05: 100% B; 2.05–2.10 min: 100% → 2% B; 2.10–3.00 min: 100% B. Data processing was performed using Waters MassLynx Mass Spectrometry Software 4.1.

**Synthesis of ISG15-C-terminal domain-Rhodamine-morpholine (ISG15$_{CT}$-Rho-MP)**

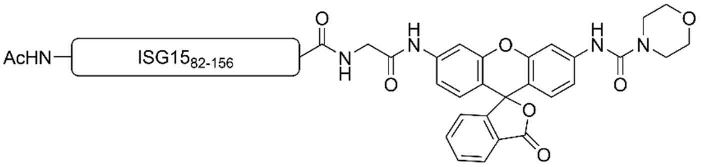


Protected Ac-ISG15$_{82-156}$ (20 µmol) was synthesized by Fmoc SPPS on trityl-resin as described.[1] It was dissolved in DMF (7 mL) and to this were added EDC.HCl (11.5 mg, 60 µmol 3.0 eq.), HOBt (8 mg, 60 µmol, 3.0 eq.) and Gly-Rho-morpholine[2] (30 mg, 60 µmol, 3.0 eq.) and the mixture was stirred at rt for 5 hours before being concentrated to dryness under reduced pressure. The resulting residue was dissolved in a mixture of 90.5/5/2/2.5 v/v/v/v TFA/$H_2O$/TIPS/phenol (7 mL) and stirred for 3 hours at rt. The protein was precipitated from ice-cold $Et_2O$/*n*-pentane (1/1; v/v; 30 mL). The solution was centrifuged (5 min. at 1500 rpm) and the supernatant was removed by decanting. The pellet was washed with $Et_2O$ (20 mL), the solution was vortexed, the suspension was centrifuged and $Et_2O$ was removed by decanting. The wash step was repeated twice. The pellet was dissolved in $H_2O/CH_3CN$/AcOH (65/25/10; v/v/v; 15 mL) and lyophilized. The protein was subsequently purified using RP-HPLC and the product was obtained as a white solid (55.4 mg, 6.1 µmol, 31%). LC-MS (10 min. run): $R_t$ (min.) 3.90; deconvoluted mass: 9008.0.

**(*R*)-3-(3-chlorophenyl)-*N*-(1-cyanopyrrolidin-3-yl)isoxazole-5-carboxamide (1)**

Synthesized as described in literature.[3]

$^1$H NMR (300 MHz, DMSO-$d_6$) δ 9.31 (d, *J* = 6.5 Hz, 1H), 8.00 (t, *J* = 1.9 Hz, 1H), 7.91 (dt, *J* = 7.0, 1.7 Hz, 1H), 7.65 – 7.54 (m, 2H), 4.56 – 4.42 (m, 1H), 3.65 (dd, *J* = 9.8, 6.4 Hz, 1H), 3.56 (dt, *J* = 9.1, 7.4 Hz, 1H), 3.46 (ddd, *J* = 9.0, 8.0, 5.4 Hz, 1H), 3.39 – 3.31 (m, 2H), 2.22 – 2.06 (m, 1H), 1.97 (ddt, *J* = 12.7, 7.5, 5.2 Hz, 1H). HR-MS calculated for $C_{15}H_{13}N_4O_2Cl$ $[M+H]^+$ 317.0805, found 317.0797.

**(*R*)-*N*-(1-cyanopyrrolidin-3-yl)-5-(2-methoxyphenyl)-1*H*-pyrazole-3-carboxamide (2)**

Synthesized as described in literature.[3]

$^1$H NMR (300 MHz, DMSO-$d_6$) δ 13.33 (s, 1H), 8.39 (s, 1H), 7.72 (s, 1H), 7.47 – 7.28 (m, 1H), 7.15 (d, *J* = 8.3 Hz, 1H), 7.04 (t, *J* = 7.4 Hz, 2H), 4.50 (td, *J* = 6.7, 5.3 Hz, 1H), 3.90 (s, 3H), 3.68 – 3.50 (m, 2H), 3.44 (ddd, *J* = 9.0, 7.6, 6.4 Hz, 1H), 3.38 – 3.29 (m, 2H), 2.18 –

2.05 (m, 1H), 2.05 – 1.92 (m, 1H). HRMS calculated for $C_{16}H_{17}N_5O_2$ $[M+H]^+$ 312.1461, found 312.1428.

**(*R*)-*N*-(1-cyanopyrrolidin-3-yl)-2-fluoro-4-(pyrrolidin-1-yl)benzamide (3)**

Synthesized as described in literature.[3]

$^1$H NMR (300 MHz, DMSO-$d_6$) δ 8.56 (d, *J* = 6.4 Hz, 1H), 7.21 – 7.12 (m, 1H), 6.52 – 6.34 (m, 2H), 4.35 (tdd, *J* = 6.2, 4.1, 2.0 Hz, 1H), 3.58 (dd, *J* = 9.8, 6.0 Hz, 1H), 3.53 – 3.39 (m, 2H), 3.33 (s, 1H), 3.28 (ddd, *J* = 9.8, 3.9, 0.8 Hz, 1H), 3.22 – 3.12 (m, 4H), 2.08 (dtd, *J* = 12.7, 8.0, 6.1 Hz, 1H), 1.87 (td, *J* = 6.7, 5.9, 2.5 Hz, 5H). HRMS calculated for $C_{16}H_{19}N_4OF$ $[M+H]^+$ 303.1621, found 303.1615.

**(*R*)-*N*-(1-cyanopyrrolidin-3-yl)-3,4-difluorobenzamide (4)**

Synthesized as described in literature.[3]

$^1$H NMR (300 MHz, DMSO-$d_6$) δ 8.71 (s, 1H), 7.92 (ddd, *J* = 11.7, 7.8, 2.2 Hz, 1H), 7.76 (dddd, *J* = 8.2, 4.6, 2.2, 1.3 Hz, 1H), 7.57 (dt, *J* = 10.5, 8.3 Hz, 1H), 4.46 (tt, *J* = 6.3, 4.5 Hz, 1H), 3.63 (dd, *J* = 9.8, 6.4 Hz, 1H), 3.55 (dt, *J* = 9.1, 7.4 Hz, 1H), 3.45 (ddd, *J* = 9.1, 8.0, 5.3 Hz, 1H), 3.31 (dd, *J* = 9.8, 4.1 Hz, 1H), 2.22 – 2.05 (m, 1H), 1.94 (ddt, *J* = 12.6, 7.4, 5.1 Hz, 1H). HRMS calculated for $C_{12}H_{11}N_3OF_2$ $[M+H]^+$ 252.0948, found 252.0952.

**(*S*)-1-cyano-*N*-(5-morpholinothiazol-2-yl)pyrrolidine-3-carboxamide (5)**

Synthesized as described in literature.[4]

$^1$H NMR (300 MHz, DMSO-$d_6$) δ 11.96 (s, 1H), 6.68 (s, 1H), 3.75 – 3.66 (m, 4H), 3.59 (dd, *J* = 9.5, 7.8 Hz, 1H), 3.52 – 3.35 (m, 3H), 3.31 – 3.20 (m, 1H), 3.02 – 2.94 (m, 4H), 2.22 – 2.09 (m, 1H), 2.09 – 1.95 (m, 1H). HRMS calculated for $C_{13}H_{17}N_5O_2S$ $[M+H]^+$ 308.1181, found 308.1170.

**General procedure for the synthesis of cyanimide building blocks (BB1-16)**

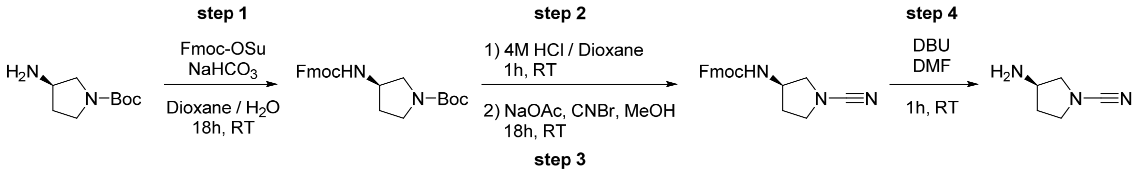


**Step 1: Fmoc protection.** Boc-protected diamine (500 mg, 1.0 eq.) was suspended in $H_2O$ (18 mL) and Fmoc-OSu (1.1 eq.) and $NaHCO_3$ (2.0 eq.) were added. Dioxane (9 mL) was added and the resulting reaction mixture was stirred at rt for 18h. TLC analysis indicated complete consumption of the starting material with formation of a less polar product. The

reaction mixture was diluted with $H_2O$ (22 mL) and EtOAc (40 mL) was added. The layers were separated and the organic layer was washed with $H_2O$ (1 x 20 mL) and BRINE (1 x 20 mL). The organic layer was dried over $Na_2SO_4$, filtered and concentrated *in vacuo* to obtain orthogonally protected diamine.

**Step 2: Boc deprotection.** Orthogonally protected diamine (1.0 eq.) was treated with 4M HCl / Dioxane (30 mL) and the resulting reaction mixture was stirred at rt for 1h. TLC analysis indicated complete consumption of the starting material with formation of a more polar product. The reaction mixture was concentrated *in vacuo* and the remaining residue was co-concentrated with DCM (3 x 30 mL) to yield mono-Fmoc protected diamine as the HCl salt.

**Step 3: Cyanimide synthesis.** Mono-Fmoc protected diamine (HCl salt) (1.0 eq) was dissolved in MeOH (15 mL) and NaOAc (5.0 eq.) and cyanogen bromide (4.0 eq.) were added. The resulting reaction mixture was stirred at rt for 18h. The reaction mixture was concentrated *in vacuo* and the crude residue was taken up in EtOAc (50 mL). The organic layer was washed with $H_2O$ (1 x 25 mL) and BRINE (1 x 25 mL). The organic layer was dried over $Na_2SO_4$, filtered and concentrated *in vacuo*. The product was isolated *via* Büchi flash chromatography (Heptane → EtOAc) to yield the mono-Fmoc protected cyanimide.

**Step 4: Fmoc deprotection.** Mono-Fmoc protected cyanimide (1.0 eq.) was dissolved in DMF (140 mM) and 1,8-Diazabicyclo[5.4.0]undec-7-ene (0.5 eq.) was added. The resulting reaction mixture was stirred at rt for 1h until TLC analysis indicated complete consumption of the starting material with formation of a more polar product. The product was isolated *via* Büchi flash chromatography (DCM → 5% MeOH, 0.7M $NH_3$) to yield the cyanimide building block.

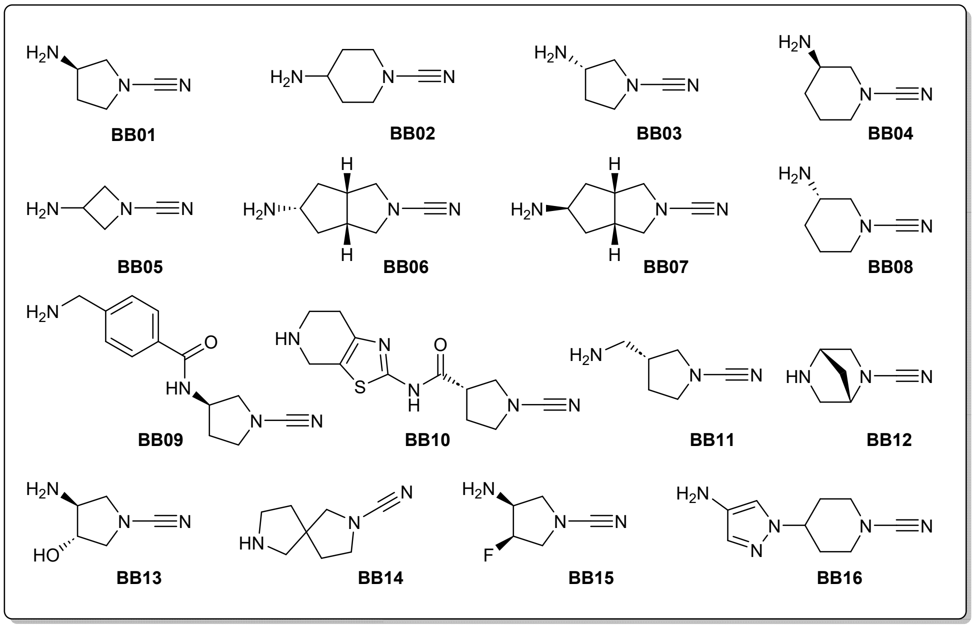


**BB1: (*R*)-3-Aminopyrrolidine-1-carbonitrile** was synthesized according to the general procedure for cyanimide building block synthesis starting from (*R*)-(+)-1-Boc-3-aminopyrrolidine. $^{1}$H NMR (300 MHz, $CDCl_3$) δ 3.63 – 3.51 (m, 2H), 3.47 (dd, *J* = 9.5, 5.7 Hz, 1H), 3.43 – 3.34 (ddd, *J* = 9.2, 8.1, 5.5 Hz, 1H), 3.03 (ddd, *J* = 9.5, 4.2, 0.8 Hz, 1H), 2.08 – 1.93 (m, 1H), 1.70 – 1.57 (m, 1H), 1.26 (s, 2H). $^{13}$C NMR (75 MHz, $CDCl_3$) δ 117.6, 58.3, 51.3, 49.0, 34.6. HR-MS calculated for $C_5H_9N_3$ $[M+H]^+$ 112.0875, found 112.0895.

**BB2: 4-Aminopiperidine-1-carbonitrile** was synthesized according to the general procedure for cyanimide building block synthesis starting from 1-Boc-4-aminopiperidine. $^{1}$H NMR (300 MHz, $CDCl_3$) δ 3.38 (dt, *J* = 13.0, 4.0 Hz, 2H), 3.05 – 2.91 (m, 2H), 2.83 – 2.67 (m, 1H), 1.79 (dd, *J* = 13.2, 3.6 Hz, 2H), 1.49 – 1.31 (m, 4H). $^{13}$C NMR (75 MHz, $CDCl_3$) δ 118.3, 48.3, 47.1, 34.2. HR-MS calculated for $C_6H_{11}N_3$ $[M+H]^+$ 126.1031, found 126.1048.

**BB3: (*S*)-3-Aminopyrrolidine-1-carbonitrile** was synthesized according to the general procedure for cyanimide building block synthesis starting from (*S*)-(-)-1-Boc-3-aminopyrrolidine. $^{1}$H NMR (300 MHz, $CDCl_3$) δ 3.68 – 3.35 (m, 4H), 3.05 (dt, *J* = 9.4, 4.4 Hz, 1H), 2.03 (dt, *J* = 12.8, 6.1 Hz, 1H), 1.66 (tt, *J* = 7.5, 4.3 Hz, 1H), 1.29 (s, 2H). $^{13}$C NMR (75 MHz, $CDCl_3$) δ 117.7, 58.4, 51.4, 49.1, 34.7. HR-MS calculated for $C_5H_9N_3$ $[M+H]^+$ 112.0875, found 112.0872.

**BB4: (*R*)-3-Aminopiperidine-1-carbonitrile** was synthesized according to the general procedure for cyanimide building block synthesis starting from (*R*)-*tert*-butyl 3-

aminopiperidine-1-carboxylate. $^1$H NMR (300 MHz, $CDCl_3$) δ 3.28 – 3.03 (m, 2H), 2.84 – 2.72 (m, 2H), 2.58 – 2.48 (m, 1H), 1.82 – 1.41 (m, 3H), 1.22 (s, 2H), 1.12 – 0.91 (m, 1H). $^{13}$C NMR (75 MHz, $CDCl_3$) δ 118.2, 56.8, 49.2, 46.4, 32.5, 22.8. HR-MS calculated for $C_6H_{11}N_3$ $[M+H]^+$ 126.1031, found 126.1033.

**BB5: 3-Aminoazetidine-1-carbonitrile** was synthesized according to the general procedure for cyanimide building block synthesis starting from 1-Boc-3-aminoazetidine. $^1$H NMR (300 MHz, $CDCl_3$) δ 4.29 – 4.18 (m, 2H), 3.90 – 3.75 (m, 3H), 1.63 (s, 2H). $^{13}$C NMR (75 MHz, $CDCl_3$) δ 117.8, 65.1, 45.1. HR-MS calculated for $C_4H_7N_3$ $[M+H]^+$ 98.0718, found 98.0724.

**BB6: (3a*R*,5*r*,6a*S*)-5-aminohexahydrocyclopenta[*c*]pyrrole-2(1*H*)-carbonitrile.**

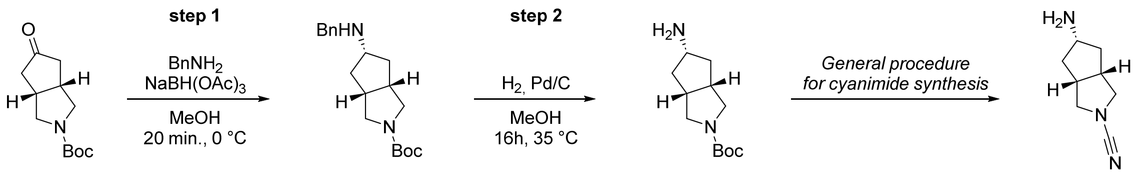


**Step 1: *tert*-Butyl (3a*R*,5*r*,6a*S*)-5-(benzylamino)hexahydrocyclopenta[*c*]pyrrole-2(1*H*)-carboxylate (6).[5]** *tert*-Butyl (3a*R*,6a*S*)-5-oxohexahydrocyclopenta[*c*]pyrrole-2(1*H*)-carboxylate (5.0 g, 22 mmol, 1.0 eq.) was dissolved in dry DCE (100 mL). Benzylamine (2.9 mL, 26 mmol, 1.2 eq.) was added and the reaction mixture was stirred for 10 min. at RT. Acetic acid (1.0 g, 22 mmol, 1.0 eq.) was added and the reaction mixture was cooled to 0 °C. Sodium triacetoxyborohydride (2.2 g, 49 mmol, 2.2 eq.) was added and the reaction was allowed to warm to RT. After TLC analysis indicated complete conversion of starting material after 16h the mixture was concentrated *in vacuo*. The residue was taken up in EtOAc (200 mL) and the organic layer was washed with sat. $NaHCO_3$ (2 x 100 mL) and BRINE (1 x 100 mL). The organic layer was dried over $Na_2SO_4$, filtered and concentrated *in vacuo* to obtain the crude product as a green/black suspension. The title compound (**6**) was isolated via Büchi flash chromatography (DCM → 5% MeOH/DCM, 0.5% $Et_3N$) as a brown oil (yield: 6.23 g, 20 mmol, 88%). $^1$H NMR (300 MHz, $CDCl_3$) δ 7.34 – 7.22 (m, 5H), 3.76 (s, 2H), 3.53 – 3.39 (m, 2H), 3.38 – 3.21 (m, 2H), 3.21 – 3.06 (m, 1H), 2.62 – 2.49 (m, 2H), 2.28 – 2.15 (m, 2H), 1.45 (s, 9H), 1.35 – 1.16 (m, 2H). $^{13}$C NMR (75 MHz, $CDCl_3$) δ 154.6, 140.4, 128.4, 128.1, 126.9, 60.6, 58.5, 53.0, 52.8, 52.0, 40.8, 28.5.

**Step 2: *tert*-Butyl (3a*R*,5*r*,6a*S*)-5-aminohexahydrocyclopenta[*c*]pyrrole-2(1*H*)-carboxylate (7).[5]** Compound **6** (6.1 g, 19 mmol, 1.0 eq.) was dissolved in MeOH (125 mL).

The reaction mixture was flushed with argon and Pd/C (10 wt %) (410 mg, 3.9 mmol, 0.2 eq.) was added. The reaction mixture was flushed with $H_2$ and stirred for 16h at 35 °C until TLC analysis indicated complete consumption of the starting material with formation of a more polar product. The mixture was filtered through Celite and concentrated *in vacuo* to obtain the title compound (**7**) as a yellow oil (yield: 4.4 g, 19 mmol, quant.). $^1$H NMR (300 MHz, MeOD) δ 3.63 – 3.42 (m, 3H), 3.34 (d, *J* = 1.9 Hz, 2H), 2.80 – 2.64 (m, 2H), 2.40 (h, *J* = 13.1, 6.7 Hz, 2H), 1.45 (s, 11H). $^{13}$C NMR (75 MHz, MeOD) δ 155.0, 79.6, 52.0, 51.1, 36.0, 27.4.

**BB6: (3a*R*,5*r*,6a*S*)-5-Aminohexahydrocyclopenta[*c*]pyrrole-2(1*H*)-carbonitrile** was synthesized according to the general procedure for cyanimide building block synthesis starting from compound **7**. $^1$H NMR (300 MHz, $CDCl_3$) δ 3.38 – 3.27 (m, 2H), 3.25 – 3.14 (m, 3H), 2.65 – 2.48 (m, 2H), 2.23 – 2.08 (p, 2H), 1.58 – 1.43 (s, 2H), 1.17 – 1.04 (q, 2H). $^{13}$C NMR (75 MHz, $CDCl_3$) δ 117.5, 57.0, 54.6, 42.6, 41.6. HR-MS calculated for $C_8H_{13}N_3$ $[M+H]^+$ 152.1188, found 152.1207.

**BB7: (3aR,5s,6aS)-5-aminohexahydrocyclopenta[c]pyrrole-2(1H)-carbonitrile.**

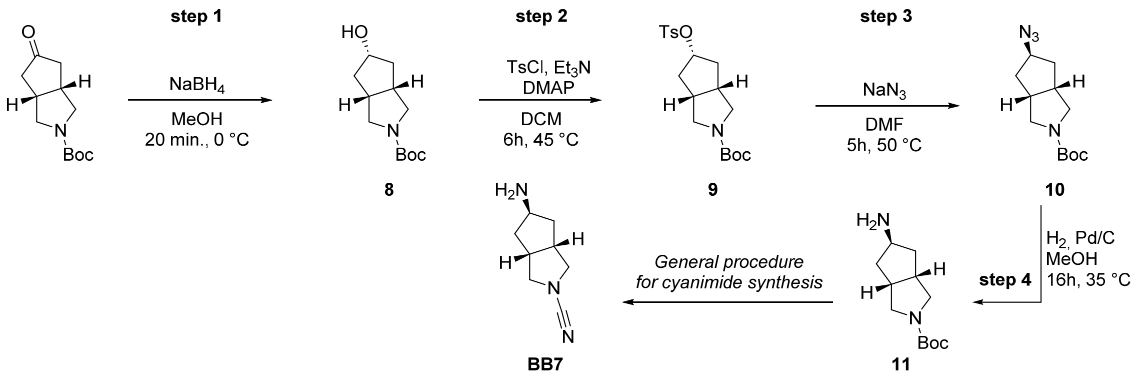


**Step 1: *tert*-Butyl (3a*R*,5*r*,6a*S*)-5-hydroxyhexahydrocyclopenta[*c*]pyrrole-2(1*H*)-carboxylate (8).** *tert*-Butyl (3a*R*,6a*S*)-5-oxohexahydrocyclopenta[*c*]pyrrole-2(1*H*)-carboxylate (5.0 g, 21 mmol, 1.0 eq.) was dissolved in MeOH (100 mL) and cooled to 0 °C. Sodium borohydride (1.6 g, 42 mmol, 2.0 eq.) was added and the reaction mixture was stirred at 0 °C for 20 min. until TLC analysis indicated complete consumption of the starting material. The reaction was quenched by addition of water (5 mL) and the reaction mixture was concentrated *in vacuo*. The residue was taken up in $H_2O$ (50 mL), EtOAc (200 mL) was added and the layers were separated. The organic layer was washed with 1M HCl (1 x 50 mL) and BRINE (1 x 50 mL). The organic layer was dried over $Na_2SO_4$, filtered and

concentrated *in vacuo* to obtain the title compound (**8**) as a colorless oil (yield: 4.6 g, 20 mmol, 97%). $^1$H NMR (300 MHz, $CDCl_3$) δ 4.26 (p, *J* = 6.6, 1.1 Hz, 1H), 3.47 (dd, *J* = 11.2, 7.6 Hz, 2H), 3.31 (dd, 2H), 2.79 (s, 1H), 2.66 – 2.50 (m, 2H), 2.14 (p, 2H), 1.53 – 1.39 (m, 11H). $^{13}$C NMR (75 MHz, $CDCl_3$) δ 154.7, 79.2, 74.5, 52.2, 41.0, 40.8, 28.5.

**Step 2: *tert*-Butyl (3a*R*,5*r*,6a*S*)-5-(tosyloxy)hexahydrocyclopenta[*c*]pyrrole-2(1*H*)-carboxylate (9).** Compound **8** (5.0 g, 21 mmol, 1.0 eq.) was dissolved in DCM (100 mL). *p*-Toluenesulfonyl chloride (4.5 g, 20 mmol, 1.2 eq.), 4-(Dimethylamino)pyridine (484 mg, 21 mmol, 0.2 eq.) and $Et_3N$ (8.3 g, 60 mmol, 3.0 eq.) were added and the reaction mixture was stirred for 16h at 45 °C. DCM (50 mL) was added after TLC analysis indicated complete consumption of the starting material. The organic layer was washed with 1M HCl (1 x 150 mL), sat. $NaHCO_3$ (1 x 150 mL) and BRINE (1 x 150 mL). The organic layer was dried over $Na_2SO_4$, filtered and concentrated *in vacuo* to obtain the title compound (**9**) as a brown oil (yield: 7.4 g, 19 mmol, 98%). $^1$H NMR (300 MHz, $CDCl_3$) δ 7.74 – 7.66 (m, 2H), 7.32 – 7.23 (m, 2H), 4.83 (tt, *J* = 6.9, 5.9 Hz, 1H), 3.49 – 3.35 (m, 2H), 3.25 – 3.11 (m, 2H), 2.48 (s, 2H), 2.37 (s, 3H), 2.06 (s, 2H), 1.60 (m, 2H), 1.39 (s, 9H). $^{13}$C NMR (75 MHz, $CDCl_3$) δ 154.4, 144.7, 134.0, 129.9, 127.7, 83.7, 79.3, 51.3, 37.8, 29.7, 28.5, 21.6.

**Step 3: *tert*-Butyl (3a*R*,5*s*,6a*S*)-5-azidohexahydrocyclopenta[*c*]pyrrole-2(1*H*)-carboxylate (10).** Compound **9** (7.4 g, 19 mmol, 1.0 eq.) was dissolved in DMF (100 mL). Sodium azide (6.3 g, 97 mmol, 5 eq.) was added and the reaction mixture was stirred for 5h at 50 °C. After TLC analysis indicated complete consumption of the starting material the mixture was concentrated *in vacuo*. The residue was taken up in EtOAc (200 mL) and the organic layer was washed with $H_2O$ (2 x 200 mL) and 1M LiCl (1 x 200 mL). The organic layer was dried over $Na_2SO_4$, filtered and concentrated *in vacuo* to obtain the crude product as a brown suspension. The title compound was isolated via Büchi flash chromatography (Heptane -> 20% EtOAc/Heptane) to obtain the title compound (**10**) as a colorless oil (yield: 4.7 g, 18 mmol, 95%). $^1$H NMR (300 MHz, $CDCl_3$) δ 4.07 – 4.01 (m, 1H), 3.48 – 3.35 (m, 2H), 3.21 – 3.06 (m, 2H), 2.83 – 2.63 (m, 2H), 1.96 – 1.85 (m, 2H), 1.72 – 1.58 (m, 2H), 1.38 (s, 9H). $^{13}$C NMR (75 MHz, $CDCl_3$) δ 154.6, 79.3, 63.4, 51.7, 38.3, 37.6, 29.7, 28.5.

**Step 4: *tert*-Butyl (3a*R*,5*s*,6a*S*)-5-aminohexahydrocyclopenta[*c*]pyrrole-2(1*H*)-carboxylate (11).** Compound **10** (4.7 g, 18 mmol, 1.0 eq.) was dissolved in MeOH (220 mL). The reaction mixture was flushed with argon and Pd/C (10 wt%) (410 mg, 3.9 mmol, 0.1 eq.) was added. The reaction was flushed with $H_2$ and stirred for 16h at 35 °C until TLC analysis indicated complete consumption of the starting material with formation of a more polar product. The mixture was filtered through Celite and concentrated *in vacuo* to obtain the title compound (**11**) as a white solid (yield: 4.2 g, 18 mmol, quant.). $^1$H NMR (300 MHz, $CDCl_3$) δ 3.56 – 3.37 (m, 3H), 3.05 (t, J = 6.3 Hz, 2H), 2.83 – 2.67 (m, 3H), 2.23 – 2.07 (m, 1H), 1.85 (dt, J = 14.6, 5.9 Hz, 1H), 1.70 (dp, J = 13.4, 3.3 Hz, 1H), 1.57 (ddd, J = 13.2, 8.1, 6.4 Hz, 1H), 1.38 (s, 9H). $^{13}$C NMR (75 MHz, $CDCl_3$) δ 154.5, 79.1, 52.5, 43.3, 41.8, 28.5.

**BB7: (3a*R*,5*s*,6a*S*)-5-Aminohexahydrocyclopenta[*c*]pyrrole-2(1*H*)-carbonitrile** was synthesized according to the general procedure for cyanimide building block synthesis starting from compound **11**.[5] $^1$H NMR (300 MHz, $CDCl_3$) δ 3.50 (p, *J* = 6.0 Hz, 1H), 3.45 – 3.36 (m, 2H), 3.05 (dd, *J* = 9.8, 3.4 Hz, 2H), 2.85 – 2.75 (m, 2H), 1.70 – 1.52 (m, 4H), 1.38 – 1.32 (m, 2H). $^{13}$C NMR (75 MHz, $CDCl_3$) δ 117.51, 56.99, 54.53, 42.57, 41.60. HR-MS calculated for $C_8H_{13}N_3$ $[M+H]^+$ 152.1188, found 152.1185.

**BB8: (*S*)-3-Aminopiperidine-1-carbonitrile** was synthesized according to the general procedure for cyanimide building block synthesis starting from (*S*)-*tert*-butyl 3-aminopiperidine-1-carboxylate. $^1$H NMR (300 MHz, $CDCl_3$) δ 3.23 (dd, *J* = 29.1, 14.5 Hz, 2H), 2.97 – 2.78 (m, 2H), 2.61 (t, 1H), 1.91 – 1.77 (m, 1H), 1.75 – 1.35 (m, 4H), 1.21 – 1.04 (m, 1H). $^{13}$C NMR (75 MHz, $CDCl_3$) δ 118.2, 56.8, 49.3, 46.5, 32.6, 22.8. HR-MS calculated for $C_6H_{11}N_3$ $[M+H]^+$ 126.1031, found 126.1034.

**BB9: (*R*)-4-(aminomethyl)-*N*-(1-cyanopyrrolidin-3-yl)benzamide.**

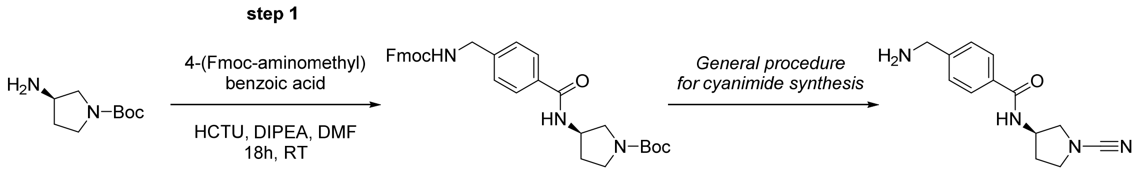

12 BB9

***tert*-Butyl (*R*)-3-(4-(((((9*H*-fluoren-9-yl)methoxy)carbonyl)amino)methyl)benzamido) pyrrolidine-1-carboxylate (12).** 4-(Fmoc-aminomethyl)benzoic acid (3.0 g, 8.05 mmol, 1.5 eq.) was dissolved in DMF (30 mL) and HCTU (3.33 g, 8.05 mmol, 1.5 eq.) and DiPEA (2.8 mL, 16.11 mmol, 3.0 eq.) were added. The resulting reaction mixture was stirred at rt for 10 min. (*R*)-(+)-1-Boc-3-aminopyrrolidine (1.0 g, 5.37 mmol, 1.0 eq) was added and the resulting reaction mixture was stirred at rt for 18h. The reaction mixture was concentrated *in vacuo* and the crude residue was taken up in EtOAc (100 mL). The organic layer was washed with 1M HCl (2 x 100 mL), sat. $NaHCO_3$ (2 x 100 mL) and BRINE (1 x 50 mL). The organic layer was dried over $Na_2SO_4$, filtered and concentrated *in vacuo*. The product was isolated *via* Büchi flash chromatography (Heptane → 70% EtOAc/Heptane) to yield compound **12** as a white solid (2.34 g, 4.32 mmol, 80%).

**BB9: (*R*)-4-(aminomethyl)-*N*-(1-cyanopyrrolidin-3-yl)benzamide** was synthesized according to the general procedure for cyanimide building block synthesis starting from compound **12**. $^1$H NMR (300 MHz, MeOD) δ 7.90 – 7.78 (m, 2H), 7.51 – 7.42 (m, 2H), 4.68 – 4.52 (m, 1H), 3.87 (s, 2H), 3.75 (dd, *J* = 9.9, 6.4 Hz, 1H), 3.70 – 3.49 (m, 2H), 3.46 – 3.37 (ddd, *J* = 9.9, 4.7, 0.7 Hz, 1H), 2.36 – 2.22 (m, 1H), 2.15 – 2.02 (m, 1H). $^{13}$C NMR (75 MHz, MeOD) δ 168.9, 146.1, 132.4, 127.4, 127.1, 116.9, 54.7, 50.1, 48.7, 44.8, 30.6. HR-MS calculated for $C_{13}H_{16}N_4O$ $[M+H]^+$ 245.1402, found 245.1409.

**BB10: (*S*)-1-Cyano-*N*-(4,5,6,7-tetrahydrothiazolo[5,4-*c*]pyridin-2-yl)pyrrolidine-3-carboxamide** was synthesized according to the general procedure for cyanimide building block synthesis starting from (9*H*-fluoren-9-yl)methyl (*S*)-2-(1-(*tert*-butoxycarbonyl)pyrrolidine-3-carboxamido)-6,7-dihydrothiazolo [5,4-*c*]pyridine-5(4*H*)-carboxylate. $^1$H NMR (300 MHz, $CDCl_3$) δ 4.00 (s, 2H), 3.74 – 3.57 (m, 3H), 3.47 (dt, *J* = 9.3, 7.4 Hz, 1H), 3.32 – 3.12 (m, 3H), 2.74 – 2.60 (m, 2H), 2.22 (td, *J* = 7.4, 5.2 Hz, 2H). $^{13}$C NMR (75 MHz, $CDCl_3$) δ 169.33, 155.97, 142.92, 122.53, 116.90, 52.79, 50.38, 44.29, 43.63, 43.26, 29.75, 27.77. HR-MS calculated for $C_{12}H_{15}N_5OS$ $[M+H]^+$ 278.1075, found 278.1086.

**BB11: (*R*)-3-(Aminomethyl)pyrrolidine-1-carbonitrile** was synthesized according to the general procedure for cyanimide building block synthesis starting from (*R*)-1-Boc-3-(aminomethyl)pyrrolidine. $^1$H NMR (300 MHz, $CDCl_3$) δ 3.56 – 3.31 (m, 3H), 3.13 (t, 1H), 2.80 – 2.63 (m, 2H), 2.27 (hept, *J* = 7.0 Hz, 1H), 2.11 – 1.94 (m, 1H), 1.75 – 1.55 (m, 1H), 1.26 (s, 2H). $^{13}$C NMR (75 MHz, $CDCl_3$) δ 117.7, 53.8, 50.1, 44.3, 42.3, 29.3. HR-MS calculated for $C_6H_{11}N_3$ $[M+H]^+$ 126.1031, found 126.1036.

**BB12: (1*S*,4*S*)-2,5-Diazabicyclo[2.2.1]heptane-2-carbonitrile** was synthesized according to the general procedure for cyanimide building block synthesis starting from *tert*-butyl (1*S*,4*S*)-2,5-diazabicyclo[2.2.1]heptane-2-carboxylate. $^1$H NMR (300 MHz, $CDCl_3$) δ 4.04 (s, 1H), 3.67 (s, 1H), 3.45 (d, *J* = 8.8 Hz, 1H), 3.20 (dd, *J* = 15.8, 9.7 Hz, 2H), 2.96 (d, *J* = 10.5 Hz, 1H), 1.90 – 1.65 (m, 3H). $^{13}$C NMR (75 MHz, $CDCl_3$) δ 117.4, 62.1, 61.1, 55.8, 51.2, 36.9. HR-MS calculated for $C_6H_9N_3$ $[M+H]^+$ 124.0875, found 124.0871.

**BB13: (3*S*,4*S*)-3-Amino-4-hydroxypyrrolidine-1-carbonitrile** was synthesized according to the general procedure for cyanimide building block synthesis starting from (3*S*,4*S*)-N-Boc-3-amino-4-hydroxypyrrolidine. $^{1}$H NMR (300 MHz, MeOD) δ 4.77 (s, 2H), 4.06 – 3.99 (m, 1H), 3.74 (m, 2H), 3.41 – 3.18 (m, 4H). $^{13}$C NMR (75 MHz, MeOD) δ 117.4, 75,7, 57.4, 55.7, 55.6. HR-MS calculated for $C_5H_9N_3O$ $[M+H]^+$ 128.0824, found 128.0831.

**BB14: 2,7-Diazaspiro[4.4]nonane-2-carbonitrile** was synthesized according to the general procedure for cyanimide building block synthesis starting from tert-butyl 2,7-diazaspiro[4.4]nonane-2-carboxylate. $^{1}$H NMR (300 MHz, $CDCl_3$) δ 3.60 – 3.41 (m, 3H), 3.38 – 3.25 (m, 2H), 3.05 (t, *J* = 7.1 Hz, 2H), 2.95 – 2.79 (m, 2H), 1.94 – 1.86 (m, 2H), 1.85 – 1.67 (m, 2H). $^{13}$C NMR (75 MHz, $CDCl_3$) δ 117.5, 60.2, 56.3, 50.0, 46.2, 36.4, 36.2. HR-MS calculated for $C_8H_{13}N_3$ $[M+H]^+$ 152.1188, found 152.1182.

**BB15: 4-(4-Amino-1H-pyrazol-1-yl)piperidine-1-carbonitrile** was synthesized according to the general procedure for cyanimide building block synthesis starting from *cis*-1-Boc-3-amino-4-fluoropyrrolidine. $^{1}$H NMR (300 MHz, $CDCl_3$) δ 4.99 – 4.94 (m, 0.5H), 4.79 (t, *J* = 2.8 Hz, 0.5H), 3.78 (d, *J* = 2.0 Hz, 1H), 3.73 – 3.64 (m, 2H), 3.65 – 3.45 (m, 1H), 3.25 – 3.16 (m, 1H). $^{13}$C NMR (75 MHz, $CDCl_3$) δ 116.45, 92.47 (d, *J* = 181.3 Hz), 55.36 (d, *J* = 23.1 Hz), 54.86 (d, *J* = 19.2 Hz), 53.77 (d, *J* = 1.8 Hz). HR-MS calculated for $C_5H_8FN_3$ $[M+H]^+$ 130.0780, found 130.0802.

**BB16: 4-(4-Amino-1*H*-pyrazol-1-yl)piperidine-1-carbonitrile** was synthesized according to the general procedure for cyanimide building block synthesis starting from 4-amino-1-(1-Boc-piperidin-4-yl)-1*H*-pyrazole. $^{1}$H NMR (300 MHz, $CDCl_3$) δ 7.18 (d, *J* = 0.9 Hz, 1H), 7.04 (d, *J* = 0.8 Hz, 1H), 4.12 (tt, *J* = 10.2, 5.1 Hz, 1H), 3.67 – 3.50 (m, 2H), 3.27 – 3.11 (m, 2H), 2.93 (s, 2H), 2.20 – 2.07 (m, 4H). HR-MS calculated for $C_9H_{13}N_5$ $[M+H]^+$ 192.1249, found 192.1253.

**BB07CA28 and BB07CA48**

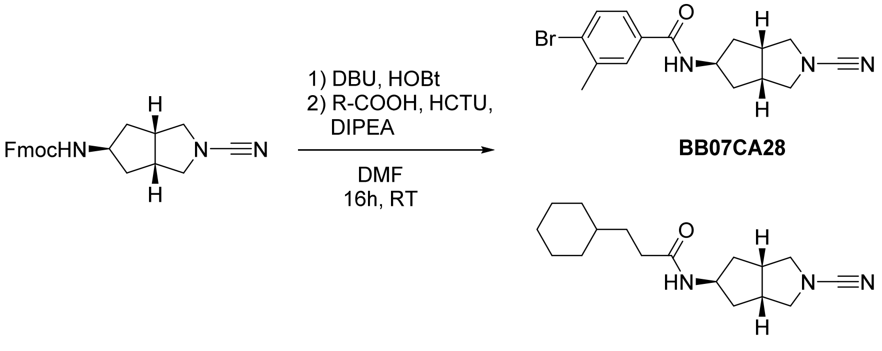


**BB07CA28: 4-Bromo-*N*-((3a*R*,5*s*,6a*S*)-2-cyanooctahydrocyclopenta[*c*]pyrrol-5-yl)-3-methylbenzamide.** Fmoc-protected **BB07** (50 mg, 133 µmol, 1.0 eq.) was dissolved in DMF (1 mL). DBU (0.5 eq., 57 µmol, 10 µL) was added and the mixture was stirred for 30 min. until TLC indicated complete removal of the Fmoc group. The reaction was quenched by addition of HOBt (1.5 eq., 201 µmol, 27 mg) and stirred for another 30 min. 4-Bromo-3-methylbenzoic acid (1.5 eq., 201 µmol, 31 mg) was dissolved in DMF (1 mL) and to this were added DiPEA (3.0 eq., 0.40 mmol, 70 µL) and HCTU (1.5 eq., 201 µmol, 10 mg). To reaction was stirred for 5 min. before being added to the deprotected amine. The whole reaction mixture was stirred for 16h after which it was concentrated under reduced pressure. The residue was taken up in EtOAc (20 mL) and washed with 1M HCl (2 x 10 mL), sat. $NaHCO_3$ (2 x 10 mL) and BRINE (1 x 10 mL). The organic layer was dried over $Na_2SO_4$, filtered and concentrated *in vacuo*. The product was purified by Büchi flash chromatography (Heptane $\rightarrow$ EtOAc) to obtain the title compound as a colorless solid (yield: 33 mg, 113 µmol, 85%). $^1$H NMR (300 MHz, $CDCl_3$) δ 7.56 – 7.52 (d, *J* = 2.2 Hz, 1H), 7.52 – 7.45 (d, *J* = 8.2 Hz, 1H), 7.33 – 7.26 (dd, *J* = 8.2, 2.3 Hz, 1H), 6.09 (d, *J* = 7.1 Hz, 1H), 4.45 (h, *J* = 7.1 Hz, 1H), 3.54 – 3.38 (m, 2H), 3.10 (dd, *J* = 10.0, 3.9 Hz, 2H), 2.81 (dq, *J* = 7.7, 3.9 Hz, 2H), 2.35 (s, 3H), 1.97 – 1.86 (ddd, *J* = 13.4, 6.5, 3.3 Hz, 2H), 1.84 – 1.72 (dt, *J* = 13.5, 8.0 Hz, 2H). $^{13}$C NMR (75 MHz, $CDCl_3$) δ 166.6, 138.4, 133.5, 132.5, 129.4, 128.5, 125.5 116.9, 57.3, 51.1, 41.2, 38.6, 22.9. HR-MS calculated for $C_{16}H_{18}BrN_3O$ $[M+H]^+$ 350.0711, found 350.0757.

**BB07CA48: *N*-((3a*R*,5*s*,6a*S*)-2-cyanooctahydrocyclopenta[*c*]pyrrol-5-yl)-3-cyclohexylpropanamide.** This compound was synthesized according to the procedure described for **BB07CA28**, but coupling 3-cyclohexylpropanoic acid on a the same scale. (yield: 31 mg, 106 µmol, 80%). $^1$H NMR (300 MHz, $CDCl_3$) δ 5.42 (d, *J* = 7.2 Hz, 1H), 4.29

(h, $J$ = 7.1 Hz, 1H), 3.46 (dd, $J$ = 10.0, 7.9 Hz, 2H), 3.07 (dd, $J$ = 10.0, 3.9 Hz, 2H), 2.82 – 2.65 (m, 2H), 2.14 – 2.02 (m, 2H), 1.90 – 1.76 (m, 2H), 1.75 – 1.50 (m, 7H), 1.49 – 1.37 (m, 2H), 1.27 – 1.01 (m, 4H), 0.91 – 0.72 (m, 2H). $^{13}$C NMR (75 MHz, $CDCl_3$) δ 173.2, 116.9, 57.3, 50.4, 41.2, 38.6, 37.4, 34.3, 33.1, 26.5, 26.2. HR-MS calculated for $C_{17}H_{27}N_3O$ $[M+H]^+$ 290.2232, found 290.2268.

**BB07CA902 and BB06CA902**

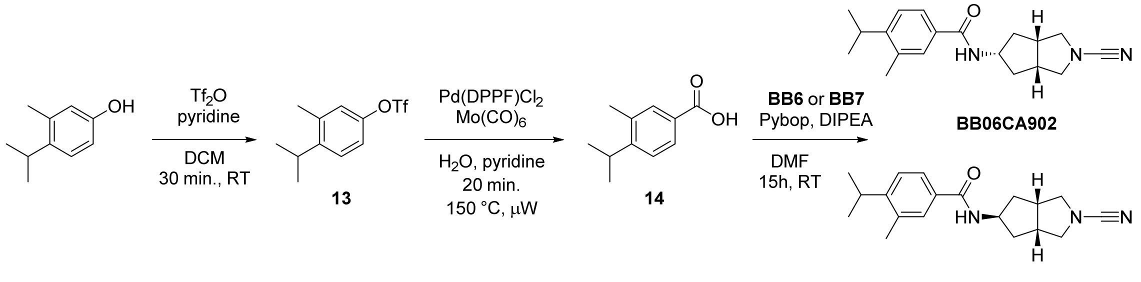


**4-Isopropyl-3-methylphenyl trifluoromethanesulfonate (13)**. A solution of 4-isopropyl-3-methylphenol (1.5 g, 10 mmol, 1.0 eq.) in dry DCM (50 mL) was flushed with argon and cooled to 0 °C. Dry pyridine (2.0 eq., 20 mmol, 1.6 mL) was added followed by a dropwise addition of trifluoromethanesulfonic anhydride (1.2 eq., 12 mmol, 2.1 mL) The reaction was slowly warmed to room temperature and TLC analysis indicated complete conversion of starting material after 30 min. The reaction was quenched by addition of water (10 mL), followed by 1M HCl (10 mL). The mixture was diluted with $Et_2O$ (100 mL), the layers were separated and the organic layer was washed with 1M HCl (1 x 50 mL), sat. $NaHCO_3$ (1 x 50 mL) and BRINE (1 x 50 mL). The organic layer was dried over $Na_2SO_4$, filtered and concentrated *in vacuo* to obtain the title compound as a colorless oil (yield: 2.79 g, 9.89 mmol, 99%). $^1$H NMR (300 MHz, $CDCl_3$) δ 7.28 (d, $J$ = 8.6 Hz, 1H), 7.06 (dd, $J$ = 8.5, 2.7 Hz, 1H), 7.02 (d, $J$ = 2.7 Hz, 1H), 3.13 (hept, $J$ = 6.8 Hz, 1H), 2.36 (s, 3H), 1.22 (d, $J$ = 6.9 Hz, 6H). $^{13}$C NMR (75 MHz, $CDCl_3$) δ 147.30, 137.87, 126.61, 122.60, 121.03, 118.81, 29.19, 23.23, 19.52.

**4-Isopropyl-3-methylbenzoic acid (14)**. This compound was synthesized using a reported procedure.[6] A microwave tube was charged with a stirring bar, triflate **13** (1.13 g, 4.0 mmol, 1 eq.), Pd(DPPF)$Cl_2$ (0.1 eq., 0.4 mmol, 293 mg), water (12 mL), pyridine (1.5 mL) and $Mo(CO)_6$ (0.5 eq., 2 mmol, 528 mg). The tube was capped and the mixture was heated to 150 °C in a microwave for 20 min. while stirring. After cooling the black reaction mixture the cap was removed and 6M aqueous HCl (10 mL) was added. The mixture was extracted with $Et_2O$ (2 x 50 mL) after which the combined organic layers were extracted with 2M NaOH (2 x

20 mL). The combined NaOH layers were acidified with concentrated HCl followed by extraction with EtOAc (2 x 50 mL). The combined EtOAc layers were dried over $Na_2SO_4$, filtered and concentrated *in vacuo* to obtain the title compound as a colorless solid (yield: 180 mg, 1.0 mmol, 25%). $^1$H NMR (300 MHz, $CDCl_3$) δ 10.40 (s, 1H), 7.93 (d, *J* = 8.1 Hz, 1H), 7.90 (s, 1H), 7.34 (d, *J* = 8.1 Hz, 1H), 3.20 (hept, *J* = 6.9 Hz, 1H), 2.40 (s, 3H), 1.25 (d, *J* = 6.9 Hz, 6H). $^{13}$C NMR (75 MHz, $CDCl_3$) δ 172.54, 153.54, 135.52, 132.04, 128.36, 126.62, 125.09, 29.74, 23.05, 19.39.

**BB07CA902: *N*-((3a*R*,5*s*,6a*S*)-2-cyanooctahydrocyclopenta[*c*]pyrrol-5-yl)-4-isopropyl-3-methylbenzamide.** Fmoc-protected **BB07** (14 mg, 37 µmol, 1.0 eq.) was dissolved in DMF (0.5 mL). DBU (0.5 eq., 18.5 µmol, 2.76 µL) was added and the mixture was stirred for 30 min. until TLC indicated complete removal of the Fmoc group. The reaction was quenched by addition of HOBt (1.5 eq., 55 µmol, 7.5 mg) and stirred for another 30 min. Carboxylic acid **14** (1.5 eq., 56 µmol, 10 mg) was dissolved in DCM (0.5 mL) and to this were added DiPEA (3.0 eq., 0.11 mmol, 20 µL) and Pybop (1.5 eq., 56 µmol, 29 mg). To reaction was stirred for 5 min. before being added to the deprotected amine. The whole reaction mixture was stirred for 15h after which it was concentrated under reduced pressure. The residue was taken up in EtOAc (20 mL) and washed with sat. $NaHCO_3$ (2 x 10 mL) and BRINE (1 x 10 mL). The organic layer was dried over $Na_2SO_4$, filtered and concentrated *in vacuo*. The product was purified by Büchi flash chromatography (Heptane → EtOAc) to obtain the title compound as a colorless solid (yield: 10 mg, 32 µmol, 87%). $^1$H NMR (300 MHz, $CDCl_3$) δ 7.54 – 7.47 (m, 2H), 7.28 (d, *J* = 4.5 Hz, 1H), 5.97 (d, *J* = 7.1 Hz, 1H), 4.55 (h, *J* = 7.1 Hz, 1H), 3.55 (dd, *J* = 10.0, 7.9 Hz, 2H), 3.24 – 3.09 (m, 3H), 2.86 (dt, *J* = 7.7, 4.0 Hz, 2H), 2.37 (s, 3H), 1.99 (ddd, *J* = 13.4, 6.4, 3.4 Hz, 2H), 1.90 – 1.78 (m, 2H), 1.22 (d, *J* = 6.9 Hz, 6H). $^{13}$C NMR (75 MHz, $CDCl_3$) δ 167.54, 151.03, 135.67, 131.68, 128.89, 125.11, 124.68, 117.05, 57.42, 51.14, 41.42, 38.82, 29.50, 23.13, 19.46. HR-MS calculated for $C_{19}H_{25}N_3O$ $[M+H]^+$ 312.2076, found 312.2100.

**BB06CA902: *N*-((3a*R*,5*r*,6a*S*)-2-cyanooctahydrocyclopenta[*c*]pyrrol-5-yl)-4-isopropyl-3-methylbenzamide.** This compound was synthesized according to the procedure described for **BB07CA902**, starting from **BB06** on a 25 mg, 67 µmol scale. (yield: 16 mg, 51 µmol, 76%). $^1$H NMR (300 MHz, $CDCl_3$) δ 7.54 (m, 2H), 7.29 (d, *J* = 8.1 Hz, 1H), 6.22 (d, *J* = 5.7 Hz, 1H), 4.50 – 4.32 (m, 1H), 3.42 (dd, *J* = 10.0, 7.0 Hz, 2H), 3.31 (dd, *J* = 10.0, 1.8 Hz, 2H), 3.16 (hept, *J* = 6.9 Hz, 1H), 2.73 (dd, *J* = 7.9, 4.7 Hz, 2H), 2.46 (dt, *J* = 13.8, 7.1 Hz, 2H), 2.38 (s, 3H), 1.38 (m, 2H), 1.22 (d, *J* = 6.8 Hz, 6H). $^{13}$C NMR (75 MHz, $CDCl_3$) δ 151.07, 135.72, 128.82, 125.19, 124.75, 57.47, 51.57, 41.20, 39.69, 29.52, 23.14, 19.47. HR-MS calculated for $C_{19}H_{25}N_3O$ $[M+H]^+$ 312.2076, found 312.2100.

#### Echo synthesis and analysis

##### Test reaction and optimizations

The coupling of carboxylic acid **CA64** with amine **BB1** was used as a model reaction to find the optimal conditions for in-plate synthesis. Reactions were conducted in 1.5 mL Eppendorf tubes using DMSO as solvent at a total volume of 300 µL. Stock solutions of **BB1** (200 mM in DMSO), **CA64** and all reagents (each 500 mM in DMSO) were made. Empty vials were first charged with DMSO, followed by addition of **CA64**, coupling reagents and base where appropriate. All tested reaction conditions are listed in the table below. After 30 min. **BB1** was added and samples were taken after 1h, 3h and 16h, and analyzed by TLC (15% MeOH/DCM). The choice for the optimal reaction conditions (condition 6.2: 5 eq. DIC/HOBt) was based on the degree of consumed **BB1** and the amount of undesired side product formation.

| Condition | Amine:Reagent:base (eq.) | Coupling reagent/Base |
|---|---|---|
| 1.1 | 1:2:3 | HBTU/NMM |
| 1.2 | 1:5:15 | HBTU/NMM |
| 1.3 | 1:10:30 | HBTU/NMM |
| 2.1 | 1:2:3 | HCTU/NMM |
| 2.2 | 1:5:15 | HCTU/NMM |
| 2.3 | 1:10:30 | HCTU/NMM |
| 3.1 | 1:2:3 | HATU/NMM |
| 3.2 | 1:5:15 | HATU/NMM |
| 3.3 | 1:10:30 | HATU/NMM |
| 4.1 | 1:2:3 | Pybop/NMM |
| 4.2 | 1:5:15 | Pybop/NMM |
| 4.3 | 1:10:30 | Pybop/NMM |
| 5.1 | 1:2 | EDC+HOBt |
| 5.2 | 1:5 | EDC+HOBt |
| 5.3 | 1:10 | EDC+HOBt |
| 6.1 | 1:2 | DIC+HOBt |
| 6.2 | 1:5 | DIC+HOBt |
| 6.3 | 1:10 | DIC+HOBt |
| 7.1 | 1:2:3 | EDC/DMAP |
| 7.2 | 1:5:15 | EDC/DMAP |

| | | |
|---|---|---|
| 7.3 | 1:10:30 | EDC/DMAP |
| 8.1 | 1:2:3 | DIC/DMAP |
| 8.2 | 1:5:15 | DIC/DMAP |
| 8.3 | 1:10:30 | DIC/DMAP |

NMM: *N*-methylmorpholine; HBTU: 2-(1*H*-benzotriazol-1-yl)-1,1,3,3-tetramethyluronium hexafluorophosphate; HCTU: *O*-(1*H*-6-Chlorobenzotriazole-1-yl)-1,1,3,3-tetramethyluronium hexafluorophosphate ; HATU: 1-[Bis(dimethylamino)methylene]-1*H*-1,2,3-triazolo[4,5-b]pyridinium 3-oxide hexafluorophosphate; Pybop: (Benzotriazol-1-yloxy)tripyrrolidinophosphonium hexafluorophosphate; DIC: *N*,*N*'-diisopropylcarbodiimide, EDC: 1-Ethyl-3-(3-dimethylaminopropyl)carbodiimide; HOBt: 1-hydroxybenzotriazool; DMAP: 4-dimethylaminopyridine. No base was used for conditions where HOBt was used.

**Carboxylic acid selection**

Carboxylic acids were chosen from all in-house available acids (89 compounds) and purchased from Enamine (382 compounds). For the acids obtained from Enamine, the following procedure was followed: A list containing all available bicyclic carboxylic acids with molecular weight below 400 Da was obtained. All compounds containing any of the moieties shown below were excluded. This resulted in a list of 1,304 compounds. The final selection of 382 compounds was made based on a high degree of structural variety, pricing and immediate availability. The compounds were received as 100 mM solutions in DMSO in 96-deepwell plates. Compounds were transferred to 384PP Echo-ready plates (Labcyte P-05525) using a multichannel pipette and stored at -20 °C.

Excluded moieties:

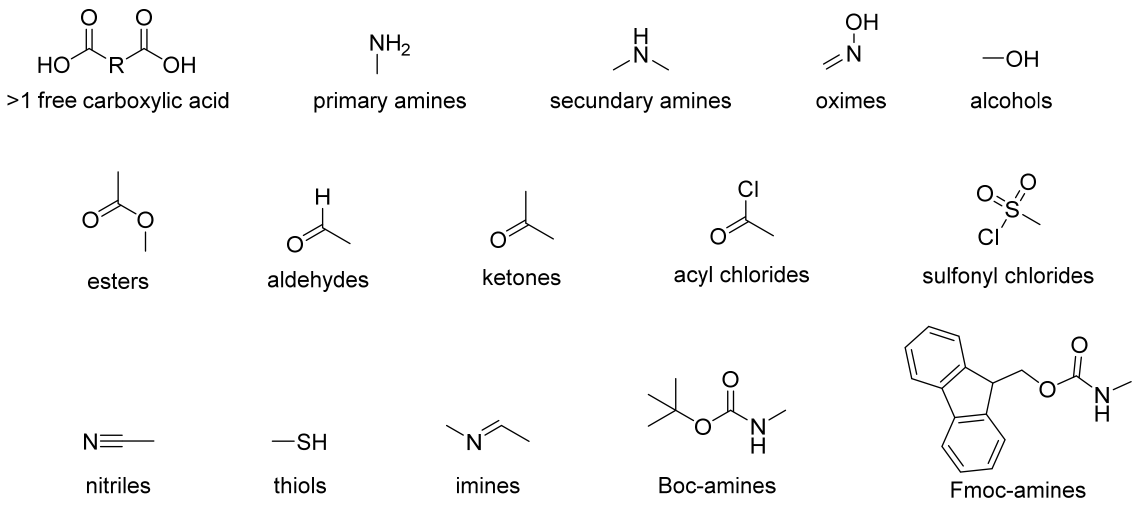

#### Miniaturized high-throughput Echo synthesis

**Synthesis in 384 well plates.** Synthesis was performed in 384LDV Echo-ready plates (Labcyte LP-0200). Each well had a total volume of 10 µL. First, 6 µL DMSO was dispensed in each well using a CombiNL Microplate dispenser (ThermoFisher). Next, 2.5 µL of the carboxylic acids **CA1-CA89** (200 mM in DMSO), 0.5 µL DIC (1M DMSO) and 0.5 µL HOBt (1M DMSO) were transferred using an Echo550 acoustic dispenser. After 30 min. incubation, 0.5 µL of the amines **BB1-BB4** and **BB6-BB8** (200 mM in DMSO) were transferred using an Echo550 acoustic dispenser. The plates were sealed and kept for 16h at room temperature and afterwards stored at -20 °C.

##### Synthesis in 1536 well plates (plates NCN1-5)

Synthesis was performed in 1536LDV Echo-ready plates (Labcyte LP-0400). Each well had a total volume of 2.5 µL. First, each well was charged with 1125 nL of a solution of DIC + HOBt (111.1 mM) in DMSO using a CombiNL Microplate dispenser (ThermoFisher). Next, 1250 nL of the carboxylic acids **CA1-CA469** (100 mM in DMSO) were transferred using an Echo550 acoustic dispenser. After 30 min. incubation, 125 nL of the amines **BB1-BB16** (200 mM in DMSO) were transferred using an Echo550 acoustic dispenser. The synthesis plates were sealed and kept for 16h at room temperature, after which 10 fold dilution daughter plates (1 mM compound concentration) were made, by dispensing 2700 nL DMSO into empty 1536LDV Echo-ready plates (using a CombiNL Microplate dispenser) followed by transfer of 300 nL from the synthesis plates (using an Echo550 acoustic dispenser). All plates were sealed and stored at -20 °C. In each plate, at least 8 wells were left empty for future controls. Plate lay-outs can be found in the supporting data files.

#### Mass spectrometry analysis of library compounds

Compounds (30 nL) from the indicated wells in the synthesis plates (see Figures) were transferred to a 96 well LC-MS plate using the Echo550 acoustic dispenser, diluted with 30 µL acetonitrile/water 1:1 v/v and analyzed on a Waters Xevo LC-MS system by injecting 1 µL sample. For the indicated 1,536 well plates, 48 compounds diagonally distributed ('hockey stick' or 'checkmark' pattern) over the plate were analyzed. By picking the compounds in such a way, at least one compound from each row and each column is analyzed, and as such serves as a good representation of the whole plate, without the need to check all 1,536 compounds in each plate. All MS spectra were analyzed and checked for the presence of the product, CA, BB and possible side products. Based on this, each compound well was annotated as ‘good’ (>90% conversion, almost no side products), ‘medium’ (50-90% conversion, or mass difficult to find due to bad ionization, some side products), or ‘bad’ (no to low conversion, many side products), and colored accordingly: green, orange and red, respectively. In some cases the ‘hockey stick’ pattern resulted in an empty well taken along in the analysis (for example N24, P24, N48, P48). These wells were purposely left empty to use as control wells in the high-throughput screen.

#### Biochemistry methods

##### Purified recombinant enzymes used in this study

| Enzyme | Domain | Tag | UniProt ID | Source | Final assay concentration |
|---|---|---|---|---|---|
| UCH-L1 | Full length (1-223) | - | P09936 | *In-house* [Larsen, 1996][7] | 1 nM |
| UCH-L3 | Full length (1-230) | - | P15374 | *In house*, [Larsen, 1996][7] | 0.01 nM |
| OTUB2 | Full length (1-234) | 6-His, clvd | Q96DC9 | *In-house*, [Nanao, 2004][8] | 25 nM |
| USP7 | Full length (1-1102) | GST-tagged, clvd | Q93009 | *In-house*, [Kim, 2019][9] | 1 nM |
| USP16 | Full length (22-823) | N-terminal His-tag | Q9Y5T5-3 | *In-house*, [Mons, 2021][10] | 2 nM |
| mUSP18 | Full length (46-368) | N-terminal His-tag | Q9UMW8 | [Basters, 2014][11] | 2 nM |
| USP30 | CD (57-517) | - | Q70CQ3 | Ubiquigent #64-0057-050 | 10 nM |
| USP32 | Full length (1-1604) | N-terminal His-tag | Q8NFA0 | *In-house*, [Sapmaz, 2019][12] | 0.8 nM |
| Papain (from papaya latex) | | - | P00784 | Sigma Aldrich #P3125 | 3 nM |

##### $IC_{50}$ and $k_{inact}/K_I$ determination

The assays were performed at room temperature in "nonbinding surface flat bottom low flange" black 384-well plates (Corning 3820) at room temperature in a buffer containing 50 mM Tris·HCl, 100 mM NaCl, pH 7.6, 2.0 mM cysteine, 1 mg/mL 3-[(3-cholamidopropyl)-dimethylammonio]propanesulfonic acid (CHAPS), and 0.5 mg/mL γ-globulins from bovine blood (BGG). Each well had a final volume of 20.4 µL. The compounds were dissolved in 10 mM DMSO stocks, and appropriate volumes were transferred to the empty plates using a Labcyte Echo550 acoustic dispenser. A DMSO back-fill was performed to obtain equal volumes of DMSO (400 µL) in each well. *N*-ethylmaleimide (NEM, 10 mM) was used a positive control (100% inhibition) and DMSO as negative control (0% inhibition). All buffer dispensing steps were performed on a Biotek MultiflowFX liquid dispenser. A 10 µL portion of buffer was added and the plate was vigorously shaken for 20s. Next, 5 µL of recombinant enzyme (4x final concentration) was added followed by incubation for 1h. A 5 µL portion of the substrate (Ub-RhoMP[2] for all DUBs, ISG15ct-RhoMP (*vide supra*) for USP18 and Z-FR-AMC [Bachem, #I-1160] for papain) was added (final concentration 400 nM for Ub-RhoMP and ISG15ct-RhoMP, 10 µM for Z-FR-AMC), and the increase in fluorescence over time was recorded using a BMG Labtech PHERAstar plate reader (excitation 487 nm, emission 535 nm for Rho; excitation 360 nm, emission 450 nm for AMC). The initial enzyme velocities were calculated from the slopes, normalized to the positive and negative controls, and plotted against the inhibitor concentrations using the built-in equation "[inhibitor] vs response − variable slope (four parameters), least-squares fit" with constraints "Bottom = 0" and "Top = 100" in GraphPad Prism 7 software to obtain the $IC_{50}$ values.

In case of the $k_{inact}/K_I$ determinations the order of mUSP18 and substrate addition was reversed and no incubation time was used. All data fitting and calculations were done using GraphPad Prism 7 software. The fluorescence intensities were plotted against time (in seconds) after a baseline correction using the DMSO control for each inhibitor concentration. The data were fitted to the equation:

$$FI = \frac{v_i}{k_{obs}}(1 - e^{-k_{obs}t})$$

Where FI is the fluorescence intensity (in AU), $v_i$ is the initial enzyme velocity (in AU/sec), kobs is the observed rate constant (in $sec^{-1}$) and t is the time (in sec). Thus obtained $k_{obs}$ values were plotted against the inhibitor concentration ([inh]) and the data were fitted to the equation below to determine the values of $k_{inact}$ and $K_I$.

$$k_{obs} = \frac{k_{inact}[inh]}{[inh] + K_I} + y_0$$

**High-throughput screening**

The screens were performed at room temperature in "flat bottom, non-treated, black polystyrene" 1536-well plates (Corning 3724) at room temperature in a buffer containing 50 mM Tris·HCl, 100 mM NaCl, pH 7.6, 2.0 mM cysteine, 1 mg/mL 3-[(3-cholamidopropyl)-dimethylammonio]propanesulfonic acid (CHAPS), and 0.5 mg/mL γ-globulins from bovine blood (BGG). Each well had a final volume of 8 µL and final compound concentration was 1.25 µM. The compounds (10 nL each) were transferred from the 1 mM daughter plates to the empty assay plates using a Labcyte Echo550 acoustic dispenser. Four wells containing DMSO (10 nL) and four wells containing NEM (10 mM final concentration) were used in each plate as negative (0% inhibition) and positive (100% inhibition) controls, respectively. All buffer dispensing steps were performed on a Biotek MultiflowFX liquid dispenser. A 6 µL enzyme solution (1.33x final concentration) was added to each well, followed by incubation for 30 min. Next, 2 µL of a substrate solution (4x final concentration) was added to each well (Ub-RhoMP[2] for all DUBs and ISG15ct-RhoMP (*vide supra*) for USP18). After 1h or 2h incubation the fluorescence intensity was recorded using a BMG Labtech PHERAstar plate reader (excitation 487 nm, emission 535 nm). The percentage inhibition was calculated for each well using the following formula:

$$\%inhibition = 100 - 100 \times \frac{FI_x - FI_{pos}}{FI_{neg} - FI_{pos}}$$

Where $FI_x$ is the fluorescence intensity of each well, $FI_{pos}$ is the average fluorescence intensity of the positive (NEM) control wells, and $FI_{neg}$ is the average fluorescence intensity of the negative (DMSO) control wells.

**Selectivity determination (Ubiquigent)**

DUBprofiler assays were performed by Ubiquigent. NCN07CA902 was tested at 0.25 µM using 100 nM ubiquitin-rhodamine110 as substrate.

**Covalent Complex Formation Mass Spectrometry Analysis**

Samples of 1.4 µM mUSP18 in 70 µL buffer containing 50 mM Tris·HCl, 100 mM NaCl at pH 7.6 and 2.0 mM cysteine were prepared. These samples were treated with 1 µL of DMSO or 1 µL of a 10 mM PG157BB7 stock solution in DMSO (140 µM final concentration) and incubated for 30 min at room temperature. Samples were then diluted 3-fold with water and analyzed by mass spectrometry by injecting 1 µL into a Waters XEVO-G2 XS Q-TOF mass spectrometer equipped with an electrospray ion source in positive mode (capillary voltage 1.2 kV, desolvation gas flow 900 L/h, T = 60 °C) with a resolution of R = 26,000. Samples were run using two mobile phases: (A) 0.1% formic acid in water and (B) 0.1% formic acid in $CH_3CN$ on a Waters Acquity UPLC protein BEH C4 column [300 Å, 1.7 µm (2.1 × 50 $mm^2$), flow rate = 0.5 mL/min, run time = 14.00 min, column T = 60 °C, and mass detection 200−2500 Da]. Gradient: 2−100% B. Data processing was performed using Waters MassLynx mass spectrometry software 4.1, and ion peaks were deconvoluted using the built-in MaxEnt1 function.

#### Cell lines and reagents

Human HEK293T (Cat# ATCC® CRL-3216™) and mouse EL4 (Cat# ATCC® TIB-39™) cells were cultured in under standard conditions in Dulbecco's modified Eagle's medium (DMEM) (Gibco) supplemented with 8% FCS (Biowest) and 1% penicillin/streptomycin at 37 °C and 5% $CO_2$. All cell lines were tested for mycoplasma contamination using MycoAlert™ Mycoplasma Detection Kit (Lonza, Catalog #: LT07-318) on a monthly basis. Interferon β from mouse (mIFN-β) was purchased from Sigma-Aldrich (Cat# I9032).

#### Murine USP18 overexpression and in-cell inhibition assays

For DNA transfections, HEK293T cells were seeded into 6-well plate to achieve 50–60% confluence the following day and then transfected with plasmids Flag-mUSP18 wildtype (Flag-mUSP18 wt) and Flag-mUSP18 catalytic cysteine-to-alanine mutant (Flag-mUSP18 C61A)[1] using PEI (polyethylenimine, Polysciences Inc., Cat# 23966) as follows: 200 µL DMEM medium without supplements was mixed with DNA and PEI (1 mg/mL) with a ratio of 1:3 (e.g.: 1 µg DNA : 3 µL PEI), incubated at RT for 20 min., and added drop-wise to the cells. At 24 hours following transfection, cells were incubated with indicated compounds at the indicated final concentrations at 37 °C for 4 hours, then washed twice with PBS and harvested for further analysis.

#### Generation of ISGylation and in-cell inhibition assay

Mouse EL4 cells were stimulated with 300 U/mL mouse IFN-β. At 24 hours following stimulation, cells were incubated with indicated compounds at the indicated final concentrations at 37 °C for 4 hours, then washed twice with PBS and harvested for further analysis.

#### Activity-based probe labelling

The in-cell inhibition of DUBs and mUSP18 was assessed using activity-based probe labelling assays. In brief, cell pellets were resuspended in HR buffer (50 mM Tris-HCl, 5 mM $MgCl_2$, 250 mM sucrose, 2 mM TCEP and a Protease inhibitor tablet (Roche), pH 7.4), and lysed by sonication (Bioruptor, Diagenode, high intensity for 10 minutes with an ON/OFF cycle of 30 seconds) at 4°C. 40 µg clarified cell lysate was incubated with Rhodamine-mISG15ct-propargylamide probe[1] (final concentration 1 µM) or Rhodamine-Ubiquitin-propargylamide probe[13] (final concentration 1 µM) at 37 °C for 30 min. Reactions were stopped by the addition of LDS (lithium dodecyl sulfate) sample buffer (Invitrogen Life Technologies, Carlsbad, CA, USA) containing 2.5% β-mercaptoethanol, followed by boiling for 7 minutes.

#### SDS-PAGE, in-gel fluorescence scanning, and immunoblotting

Samples were resolved on precast Bis-Tris NuPAGE Gels (Invitrogen, including 4-12%, and 10% for different samples) using MOPS buffer (Invitrogen Life Technologies, Carlsbad, CA, USA). For fluorescence scan, labeled enzymes were visualized by in-gel fluorescence using a Typhoon FLA 9500 imaging system (GE Healthcare Life Sciences) (Rhodamine channel for probe, Cy5 channel for protein marker).

For immunoblotting, proteins were transferred to a nitrocellulose membrane (Protan BA85, 0.45 µm, GE Healthcare) at 300 mA for 3 hours. The membranes were blocked in 5% milk (skim milk powder, LP0031, Oxiod) in 1× PBS (P1379, Sigma-Aldrich), incubated with a primary antibody diluted in 5% milk in 0.1% PBS-Tween 20 (PBST) for overnight at cold room, washed three times for 5 min in 0.1% PBST, incubated with the secondary antibody diluted in 5% milk in 0.1% PBST for 1 hour, and washed three times again in 0.1% PBST. The signal was visualized using a LICOR Odyssey system. The following primary antibodies were used: mouse anti-Flag (M2, Sigma-Aldrich), rabbit anti-mISG15[14], rabbit anti-mUSP18[15], mouse anti-GAPDH (1D4, Enzo Lifesciences) or mouse anti-β-actin (clone AC-15, Sigma-Aldrich). The following fluorescent secondary antibodies (from LICOR) were used: anti-mouse-680, anti-rabbit-800.

#### References

[1] A. Basters, P. P. Geurink, A. Rocker, K. F. Witting, R. Tadayon, S. Hess, M. S. Semrau, P. Storici, H. Ovaa, K. P. Knobeloch, G. Fritz, *Nat. Struct. Mol. Biol.* **2017**, *24*, 270-278.

[2] R. Kooij, S. H. Liu, A. Sapmaz, B. T. Xin, G. M. C. Janssen, P. A. van Veelen, H. Ovaa, P. ten Dijke, P. P. Geurink, *J. Am. Chem. Soc.* **2020**, *142*, 16825-16841.

[3] A. Jones, M. Kemp, M. Stockley, K. Gibson, G. Whitlock, *WO2016/156816 A1* **2016**.

[4] A. Jones, M. I. Kemp, M. L. Stockley, K. R. Gibson, G. A. Whitlock, A. Madin, *WO2016/046530 A1* **2016**.

[5] R. C. Lemoine, A. C. Petersen, L. Setti, T. Baldinger, J. Wanner, A. Jekle, G. Heilek, A. deRosier, C. H. Ji, D. M. Rotstein, *Bioorg. Med. Chem. Lett.* **2010**, *20*, 1674-1676.

[6] G. Lesma, A. Sacchetti, A. Silvani, *Synthesis* **2006**, 594-596.

[7] C. N. Larsen, J. S. Price, K. D. Wilkinson, *Biochem.* **1996**, *35*, 6735-6744.

[8] M. H. Nanao, S. O. Tcherniuk, J. Chroboczek, O. Dideberg, A. Dessen, M. Y. Balakirev, *EMBO Rep.* **2004**, *5*, 783-788.

[9] R. Q. Kim, P. P. Geurink, M. P. C. Mulder, A. Fish, R. Ekkebus, F. El Oualid, W. J. van Dijk, D. van Dalen, H. Ovaa, H. van Ingen, T. K. Sixma, *Nat. Commun.* **2019**, *10*, 231.

[10] E. Mons, R. Q. Kim, B. R. van Doodewaerd, P. A. van Veelen, M. P. C. Mulder, H. Ovaa, *J. Am. Chem. Soc.* **2021**, *143*, 6423-6433.

[11] A. Basters, P. P. Geurink, F. El Oualid, L. Ketscher, M. S. Casutt, E. Krause, H. Ovaa, K. P. Knobeloch, G. Fritz, *FEBS J.* **2014**, *281*, 1918-1928.

[12] A. Sapmaz, I. Berlin, E. Bos, R. H. Wijdeven, H. Janssen, R. Konietzny, J. J. Akkermans, A. E. Erson-Bensan, R. I. Koning, B. M. Kessler, J. Neefjes, H. Ovaa, *Nat. Commun.* **2019**, *10*.

[13] R. Ekkebus, S. I. van Kasteren, Y. Kulathu, A. Scholten, I. Berlin, P. P. Geurink, A. de Jong, S. Goerdayal, J. Neefjes, A. J. Heck, D. Komander, H. Ovaa, *J. Am. Chem. Soc.* **2013**, *135*, 2867-2870.

[14] A. Osiak, O. Utermöhlen, S. Niendorf, I. Horak, K. P. Knobeloch, *Mol. Cell. Biol.* **2005**, *25*, 6338-6345.

[15] L. Ketscher, R. Hannss, D. J. Morales, A. Basters, S. Guerra, T. Goldmann, A. Hausmann, M. Prinz, R. Naumann, A. Pekosz, O. Utermohlen, D. J. Lenschow, K. P. Knobeloch, *Proc. Natl. Acad. Sci. USA* **2015**, *112*, 1577-1582.

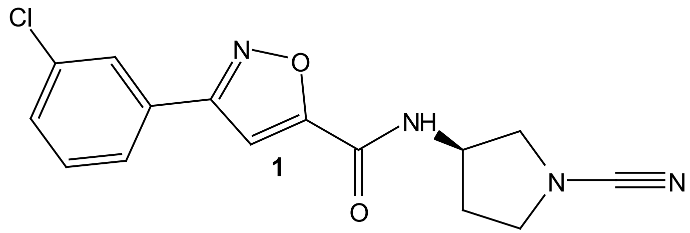
Cl
N
O
NH
N
N
O
1

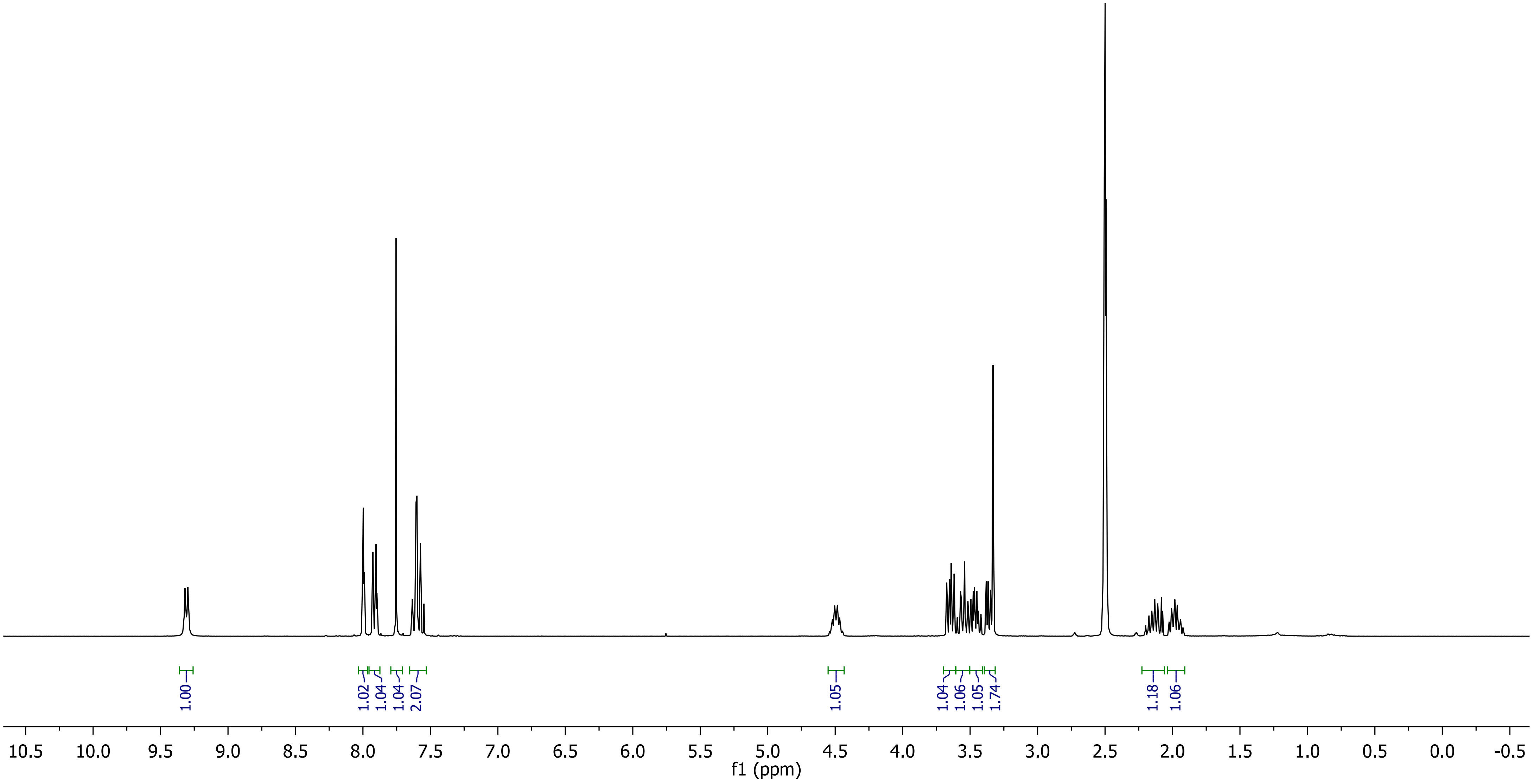
1.00
1.02
1.04
1.04
2.07
1.05
1.04
1.06
1.05
1.74
1.18
1.06
10.5 10.0 9.5 9.0 8.5 8.0 7.5 7.0 6.5 6.0 5.5 5.0 4.5 4.0 3.5 3.0 2.5 2.0 1.5 1.0 0.5 0.0 -0.5
f1 (ppm)

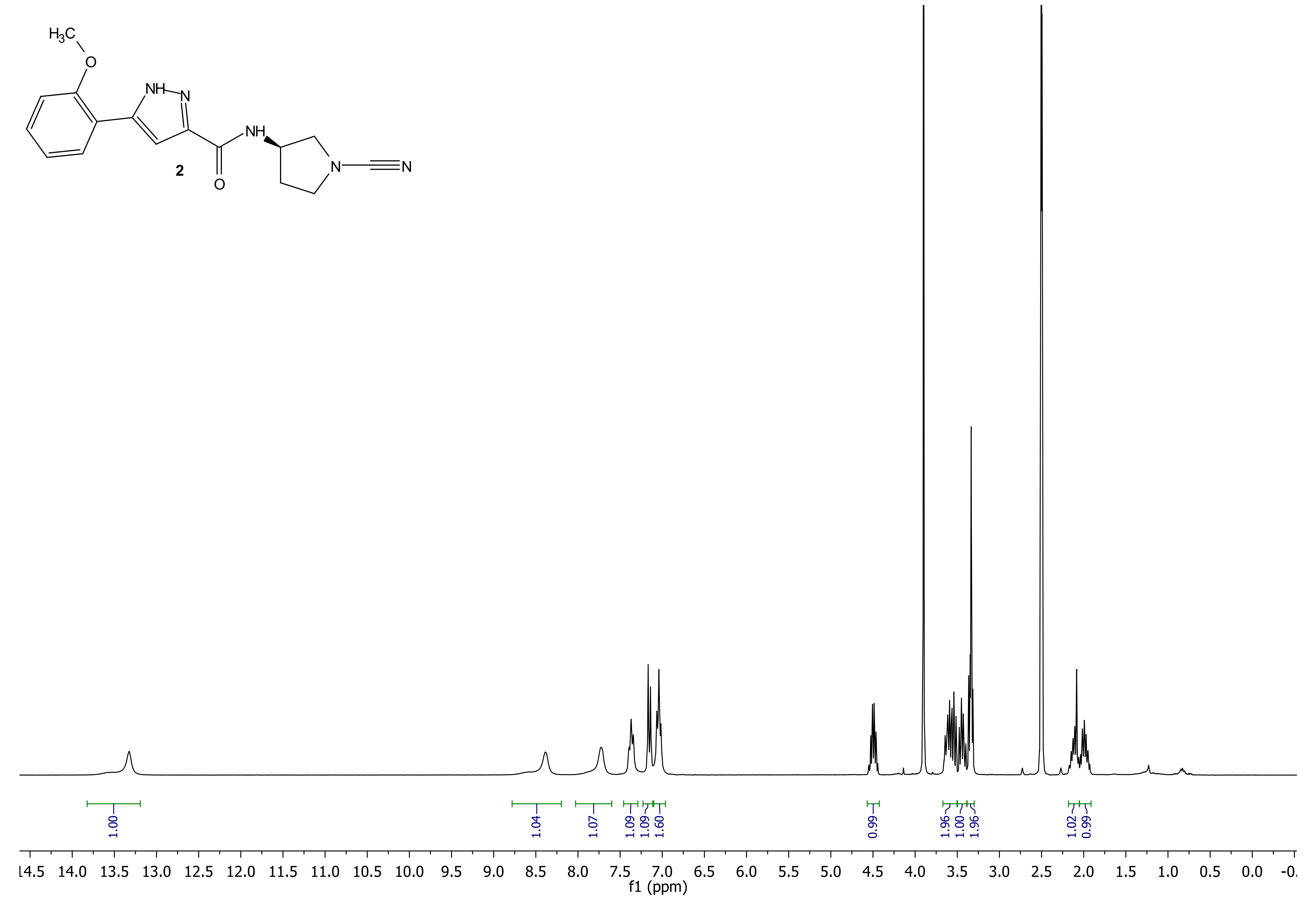

H3C
O
NH
N
NH
N
N
O
2
1.00
1.04
1.07
1.09
1.09
1.60
0.99
1.96
1.00
1.96
1.02
0.99
14.5 14.0 13.5 13.0 12.5 12.0 11.5 11.0 10.5 10.0 9.5 9.0 8.5 8.0 7.5 7.0 6.5 6.0 5.5 5.0 4.5 4.0 3.5 3.0 2.5 2.0 1.5 1.0 0.5 0.0
f1 (ppm)

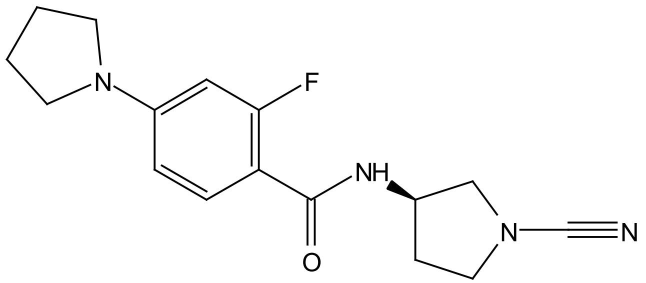
F
NH
N
N
O

**3**

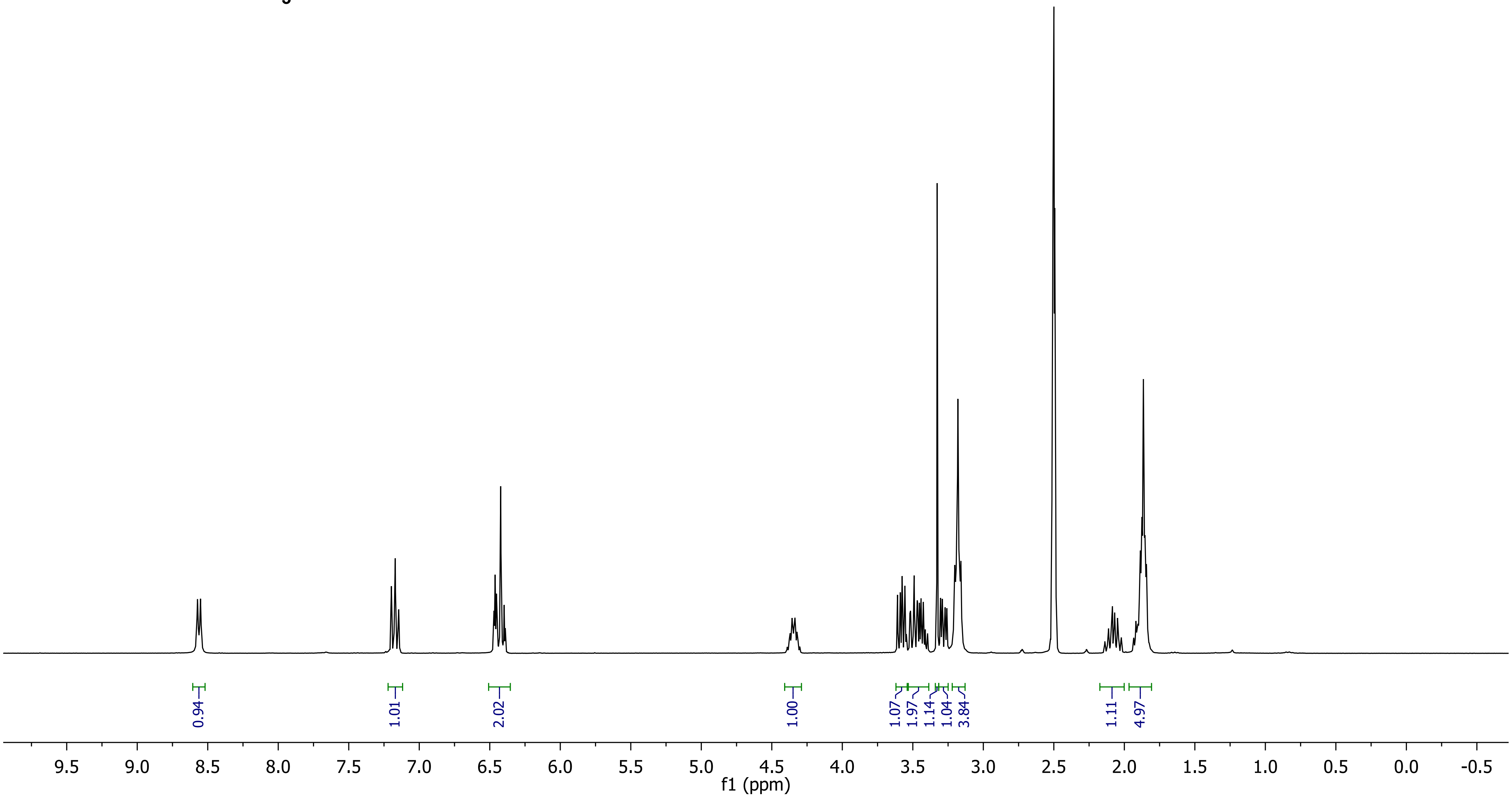
0.94
1.01
2.02
1.00
1.07
1.97
1.14
1.04
3.84
1.11
4.97
9.5 9.0 8.5 8.0 7.5 7.0 6.5 6.0 5.5 5.0 4.5 4.0 3.5 3.0 2.5 2.0 1.5 1.0 0.5 0.0 -0.5
f1 (ppm)

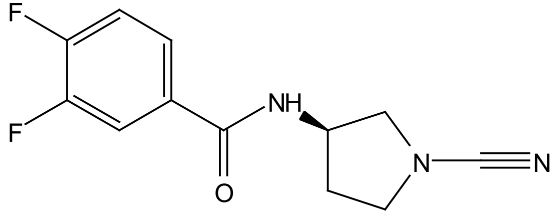
F
F
NH
O
N
N

**4**

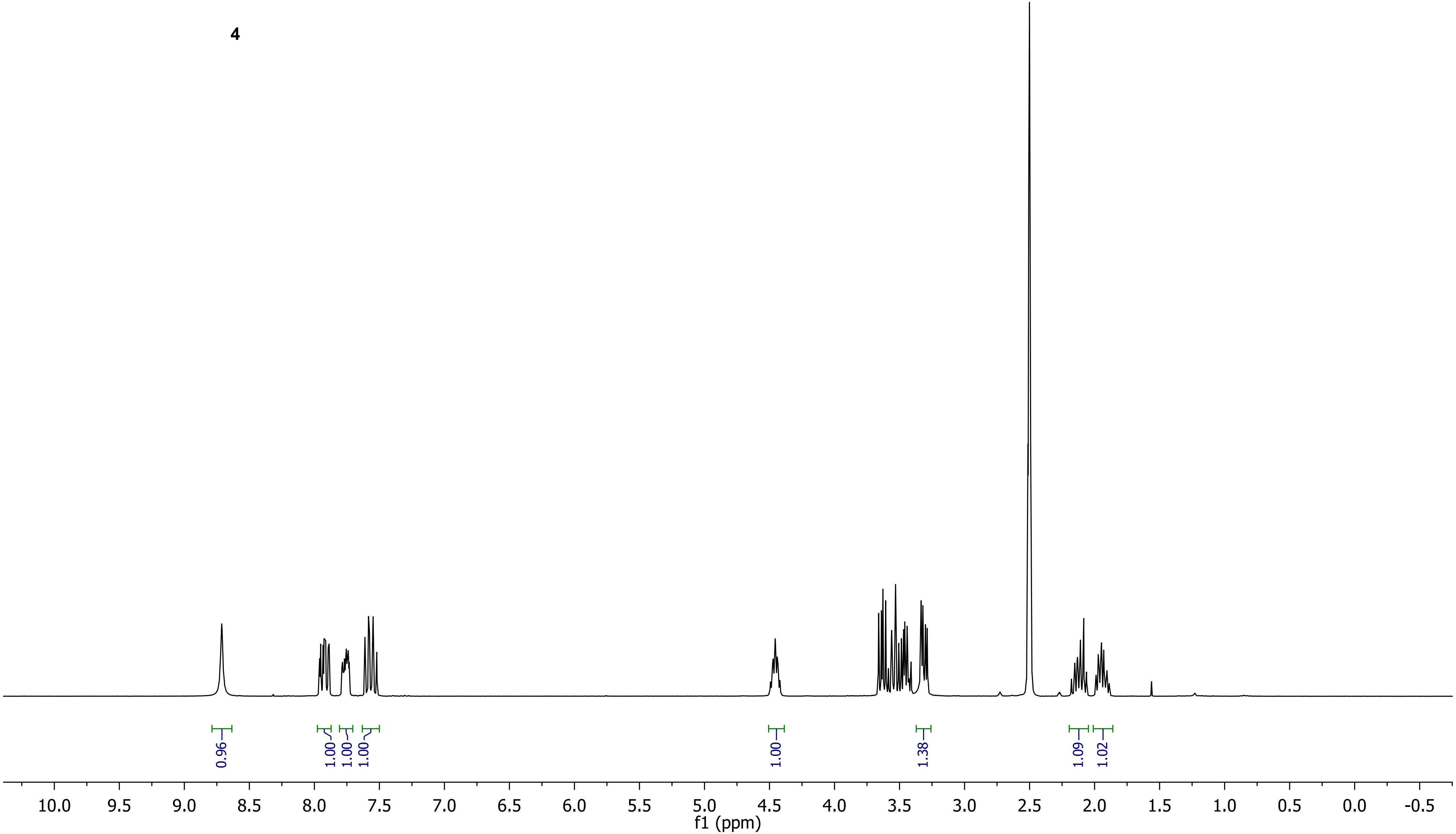
0.96
1.00
1.00
1.00
1.00
1.38
1.09
1.02
10.0 9.5 9.0 8.5 8.0 7.5 7.0 6.5 6.0 5.5 5.0 4.5 4.0 3.5 3.0 2.5 2.0 1.5 1.0 0.5 0.0 -0.5
f1 (ppm)

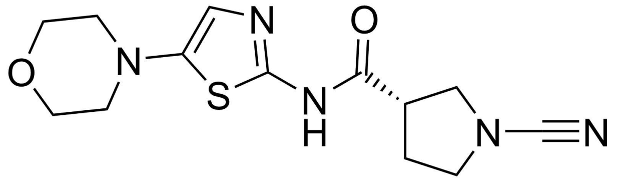

**5**

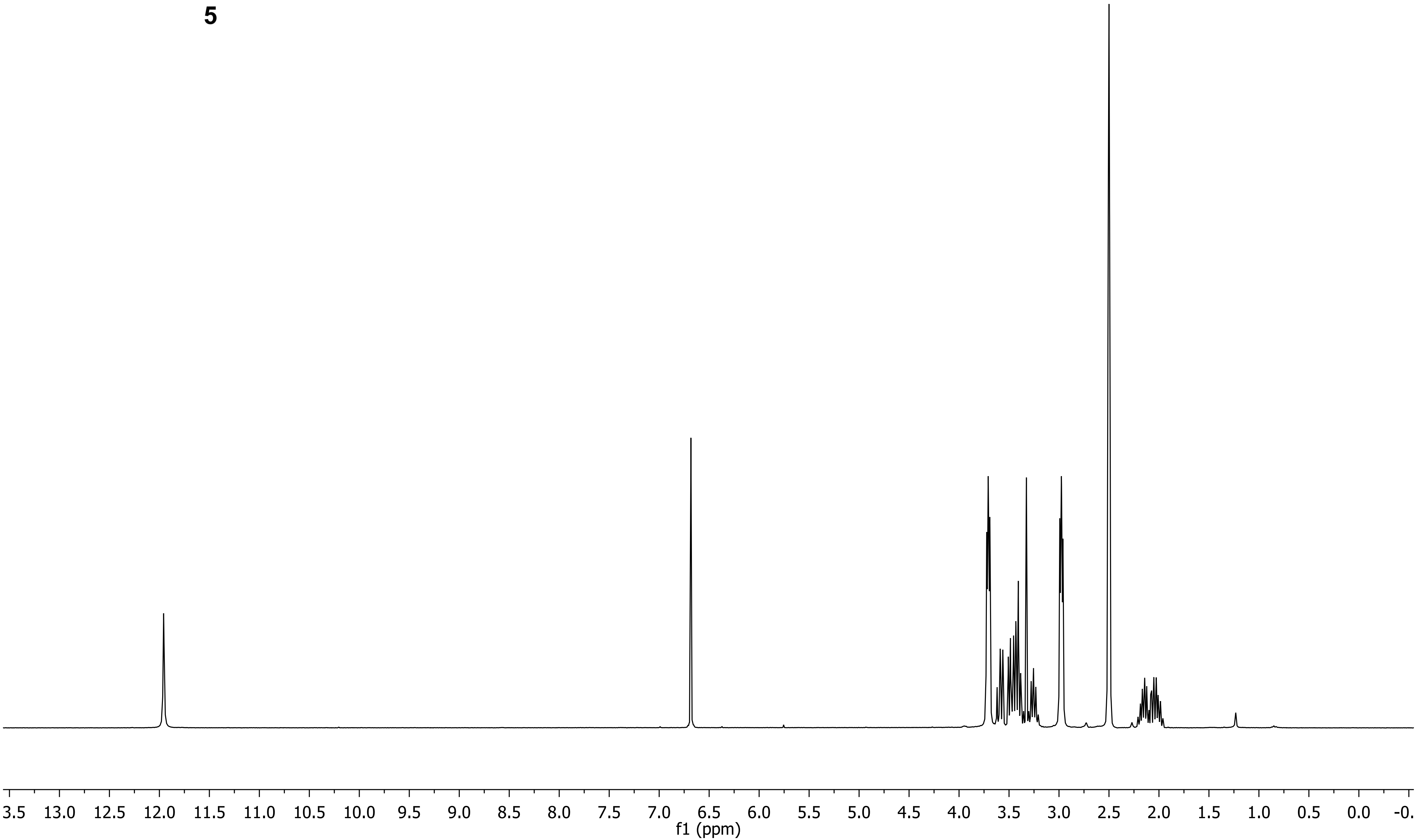
3.5 13.0 12.5 12.0 11.5 11.0 10.5 10.0 9.5 9.0 8.5 8.0 7.5 7.0 6.5 6.0 5.5 5.0 4.5 4.0 3.5 3.0 2.5 2.0 1.5 1.0 0.5 0.0 -0.
f1 (ppm)

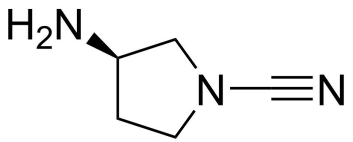
H2N
N
N

**BB1**

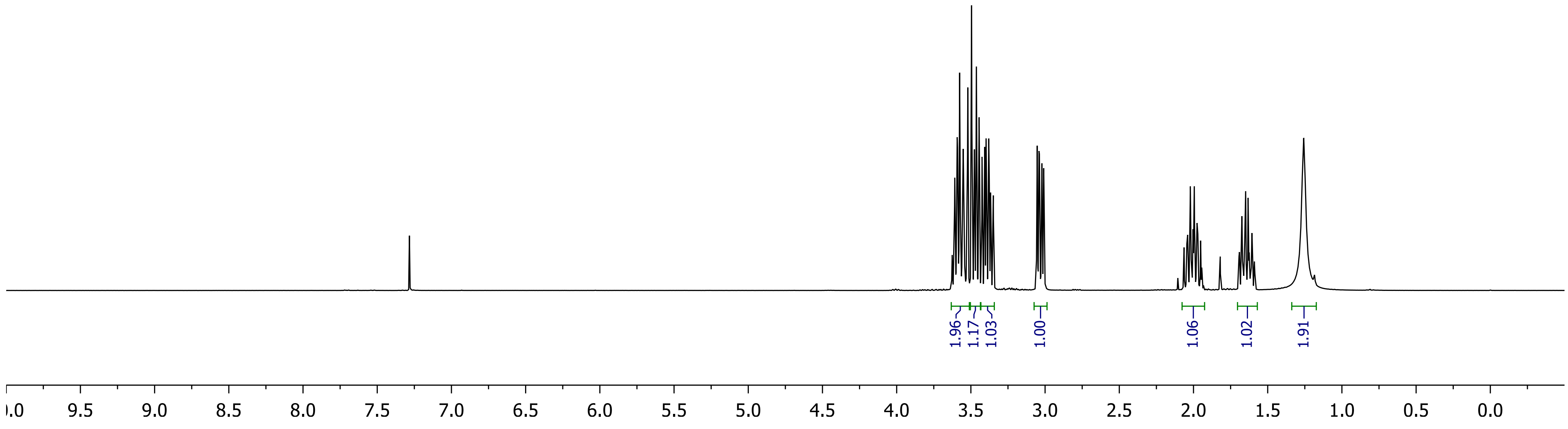
1.96
1.17
1.03
1.00
1.06
1.02
1.91
9.5
9.0
8.5
8.0
7.5
7.0
6.5
6.0
5.5
5.0
4.5
4.0
3.5
3.0
2.5
2.0
1.5
1.0
0.5
0.0

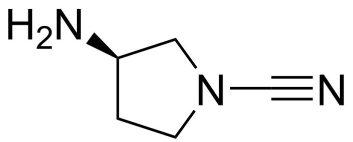
H2N
N
N

**BB1**

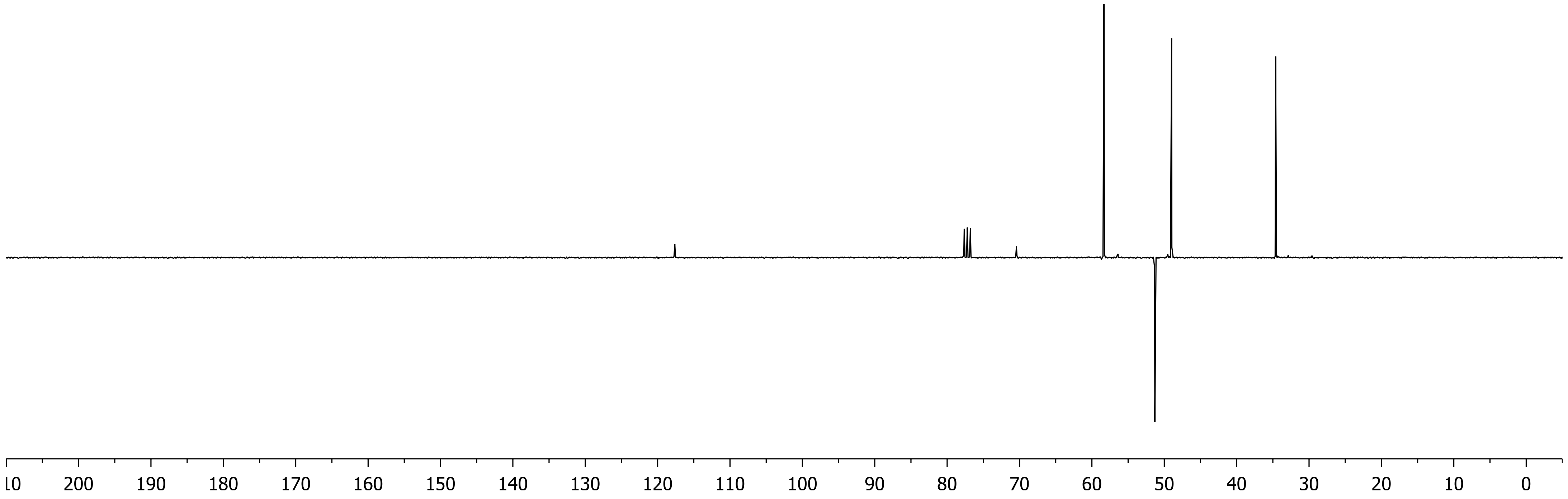
10
200
190
180
170
160
150
140
130
120
110
100
90
80
70
60
50
40
30
20
10
0

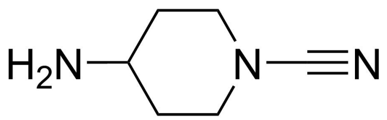
H2N
N
N

**BB2**

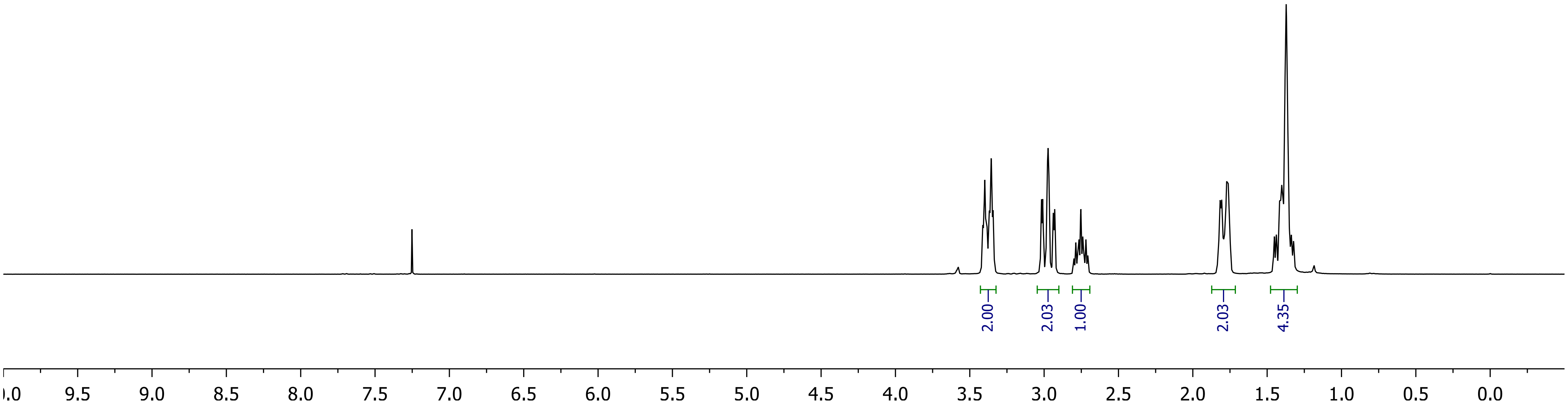
2.00
2.03
1.00
2.03
4.35
9.5
9.0
8.5
8.0
7.5
7.0
6.5
6.0
5.5
5.0
4.5
4.0
3.5
3.0
2.5
2.0
1.5
1.0
0.5
0.0

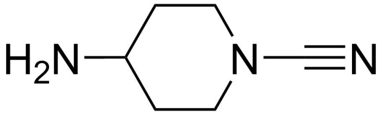

H2N
N
N


**BB2**

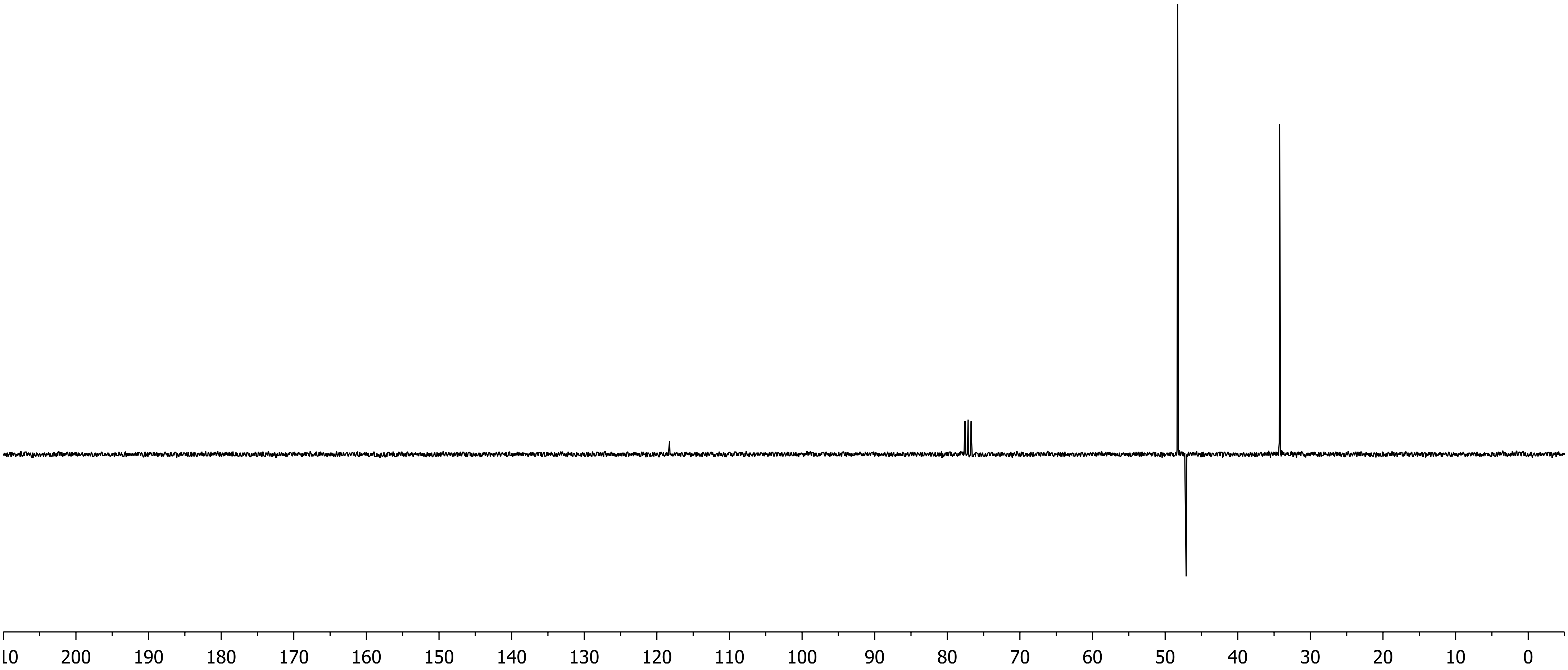

200
190
180
170
160
150
140
130
120
110
100
90
80
70
60
50
40
30
20
10
0

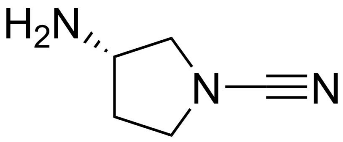
H2N
N
N

**BB3**

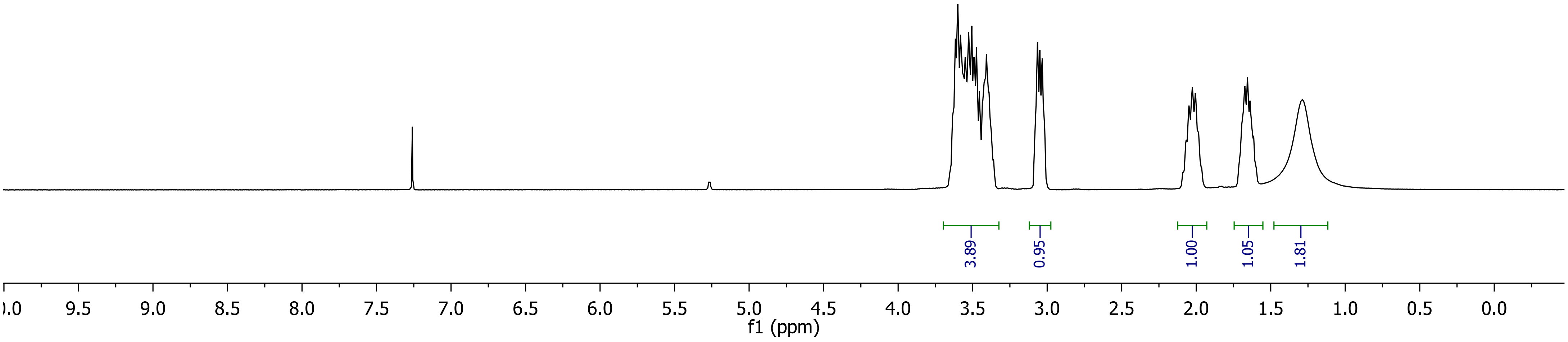
3.89
0.95
1.00
1.05
1.81
9.5
9.0
8.5
8.0
7.5
7.0
6.5
6.0
5.5
5.0
4.5
4.0
3.5
3.0
2.5
2.0
1.5
1.0
0.5
0.0
f1 (ppm)

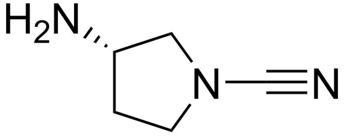

**BB3**

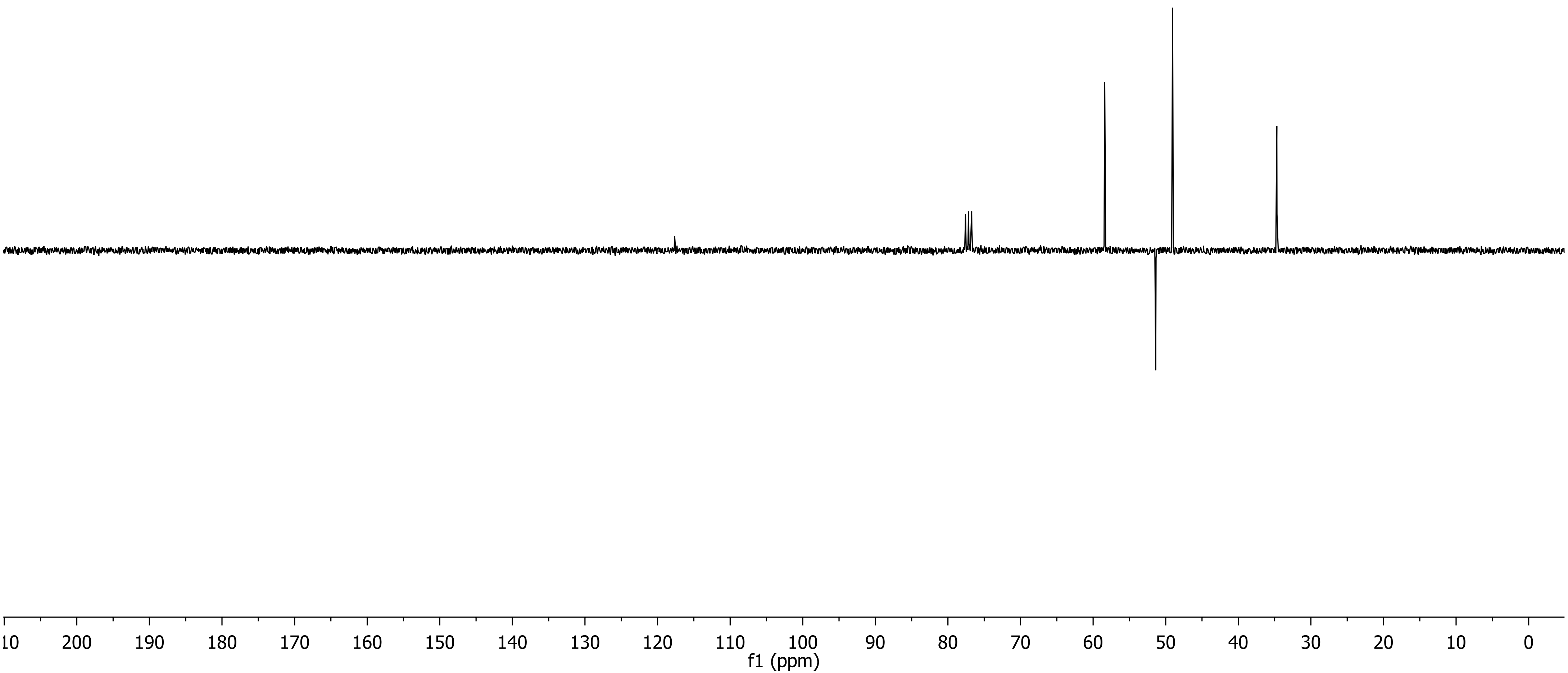
10
200
190
180
170
160
150
140
130
120
110
100
90
80
70
60
50
40
30
20
10
0
f1 (ppm)

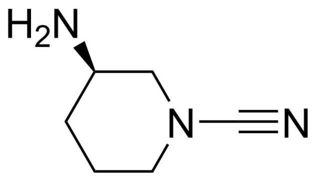

H2N
N
N


**BB4**

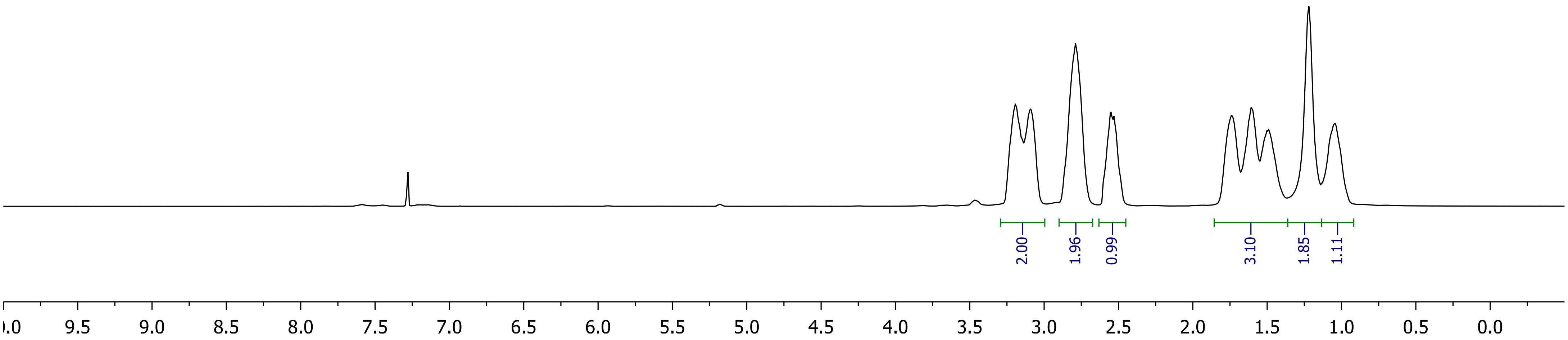

2.00
1.96
0.99
3.10
1.85
1.11
9.5
9.0
8.5
8.0
7.5
7.0
6.5
6.0
5.5
5.0
4.5
4.0
3.5
3.0
2.5
2.0
1.5
1.0
0.5
0.0

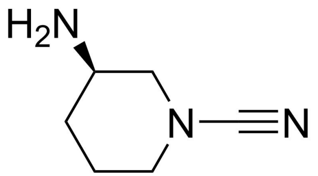

**BB4**

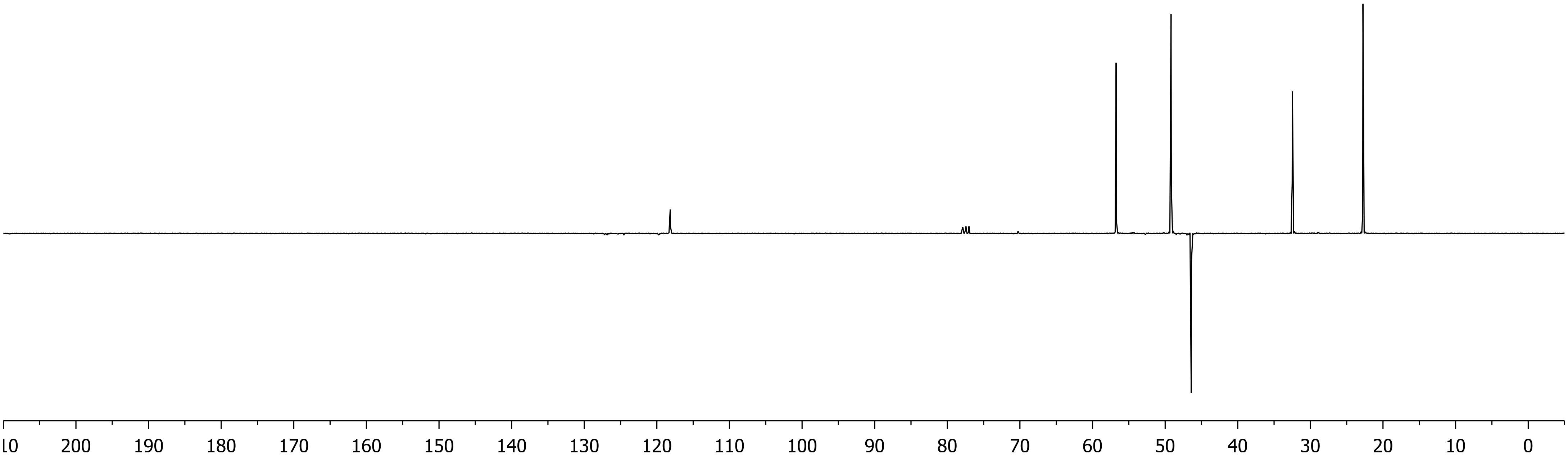

10
200
190
180
170
160
150
140
130
120
110
100
90
80
70
60
50
40
30
20
10
0

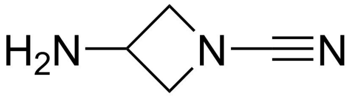

**BB5**

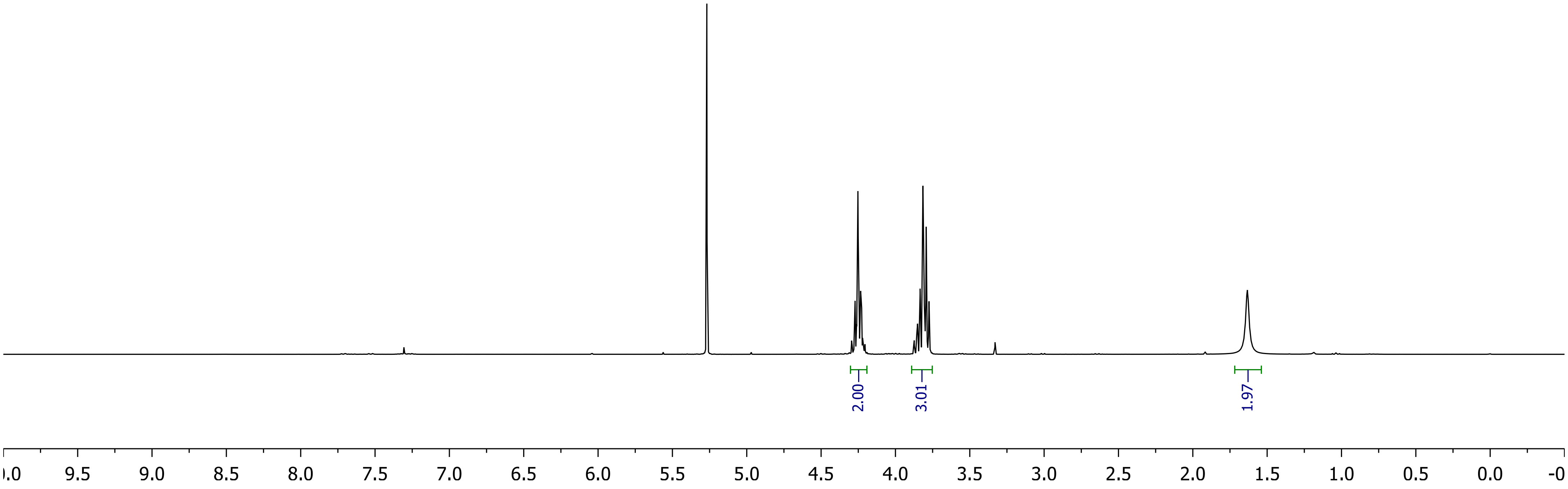

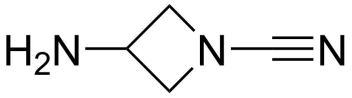
H2N
N
N

**BB5**

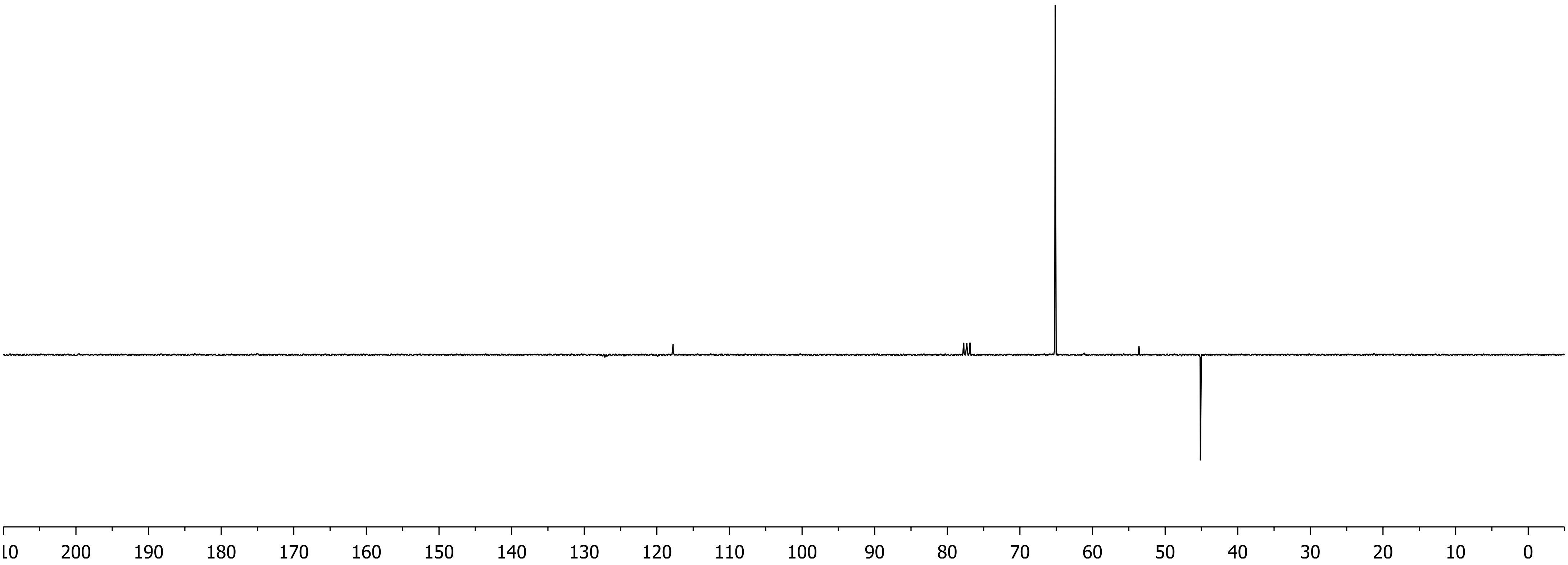
200
190
180
170
160
150
140
130
120
110
100
90
80
70
60
50
40
30
20
10
0

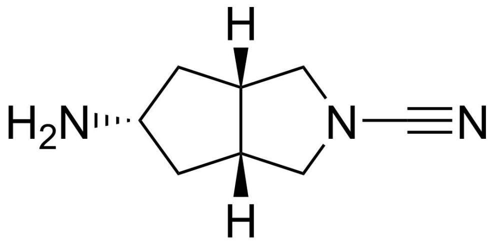

**BB6**

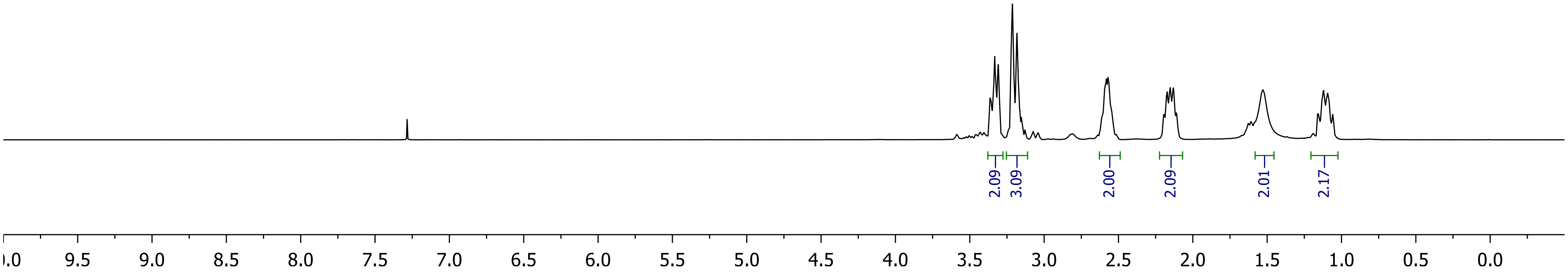
2.09
3.09
2.00
2.09
2.01
2.17
9.5
9.0
8.5
8.0
7.5
7.0
6.5
6.0
5.5
5.0
4.5
4.0
3.5
3.0
2.5
2.0
1.5
1.0
0.5
0.0

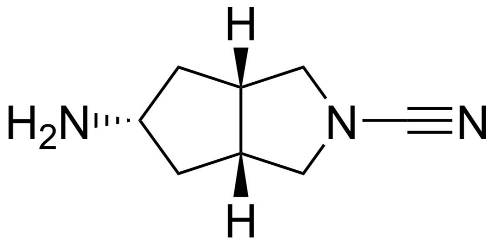
H
H2N
N
N
H

**BB6**

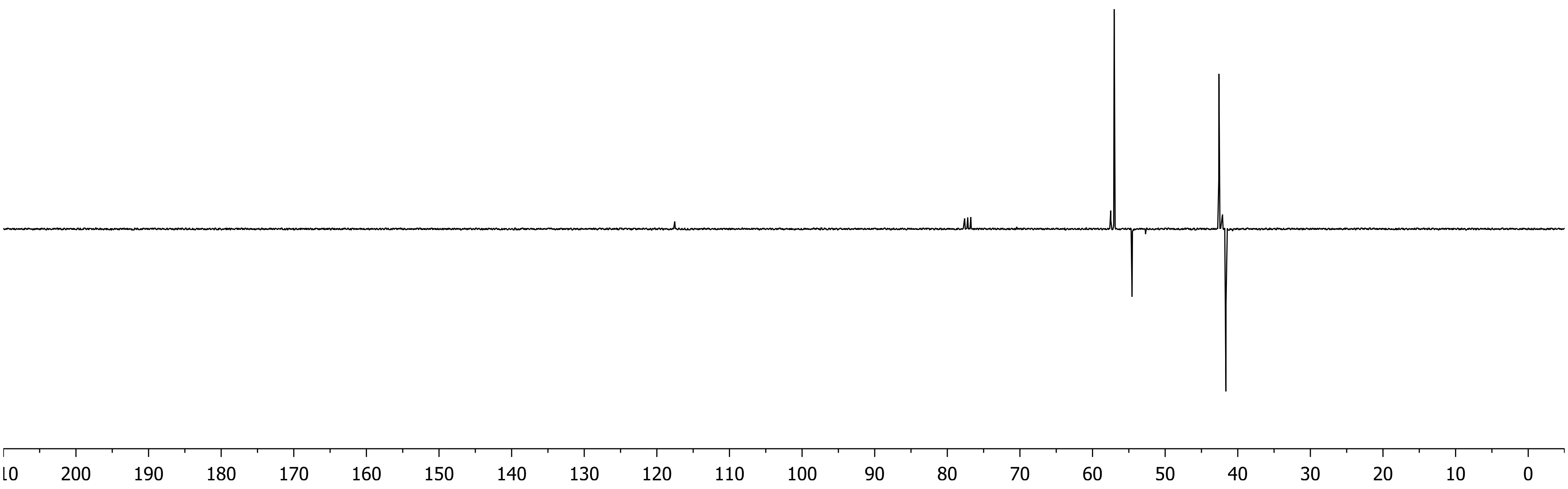
10
200
190
180
170
160
150
140
130
120
110
100
90
80
70
60
50
40
30
20
10
0

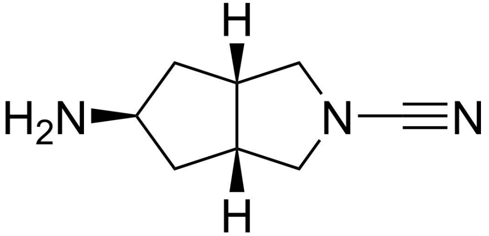

**BB7**

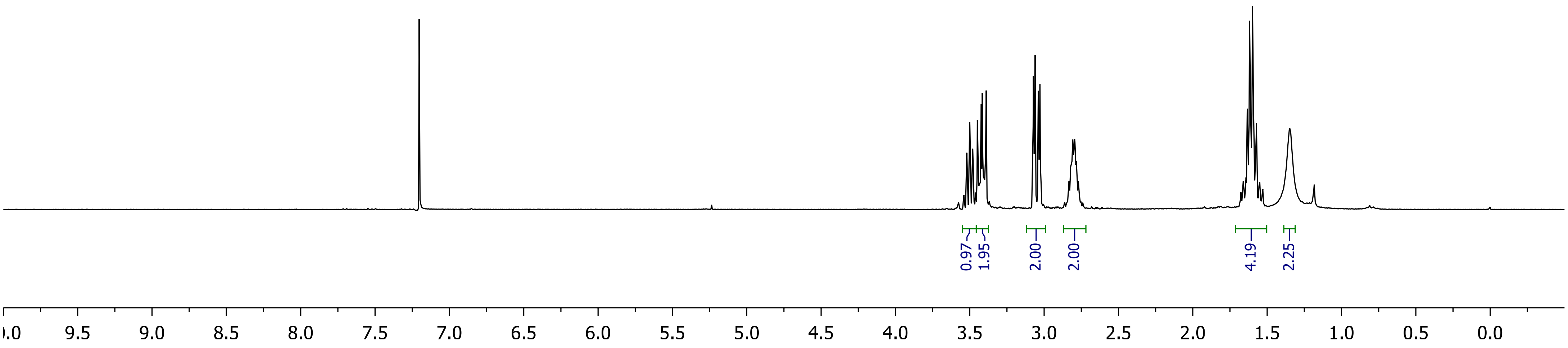

0.97
1.95
2.00
2.00
4.19
2.25
9.5
9.0
8.5
8.0
7.5
7.0
6.5
6.0
5.5
5.0
4.5
4.0
3.5
3.0
2.5
2.0
1.5
1.0
0.5
0.0

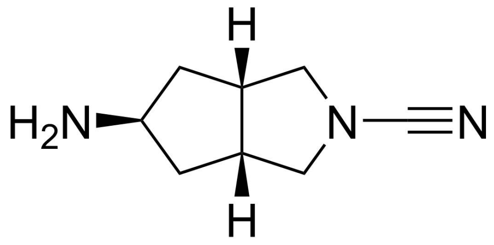
H
H₂N
N
N
H

**BB7**

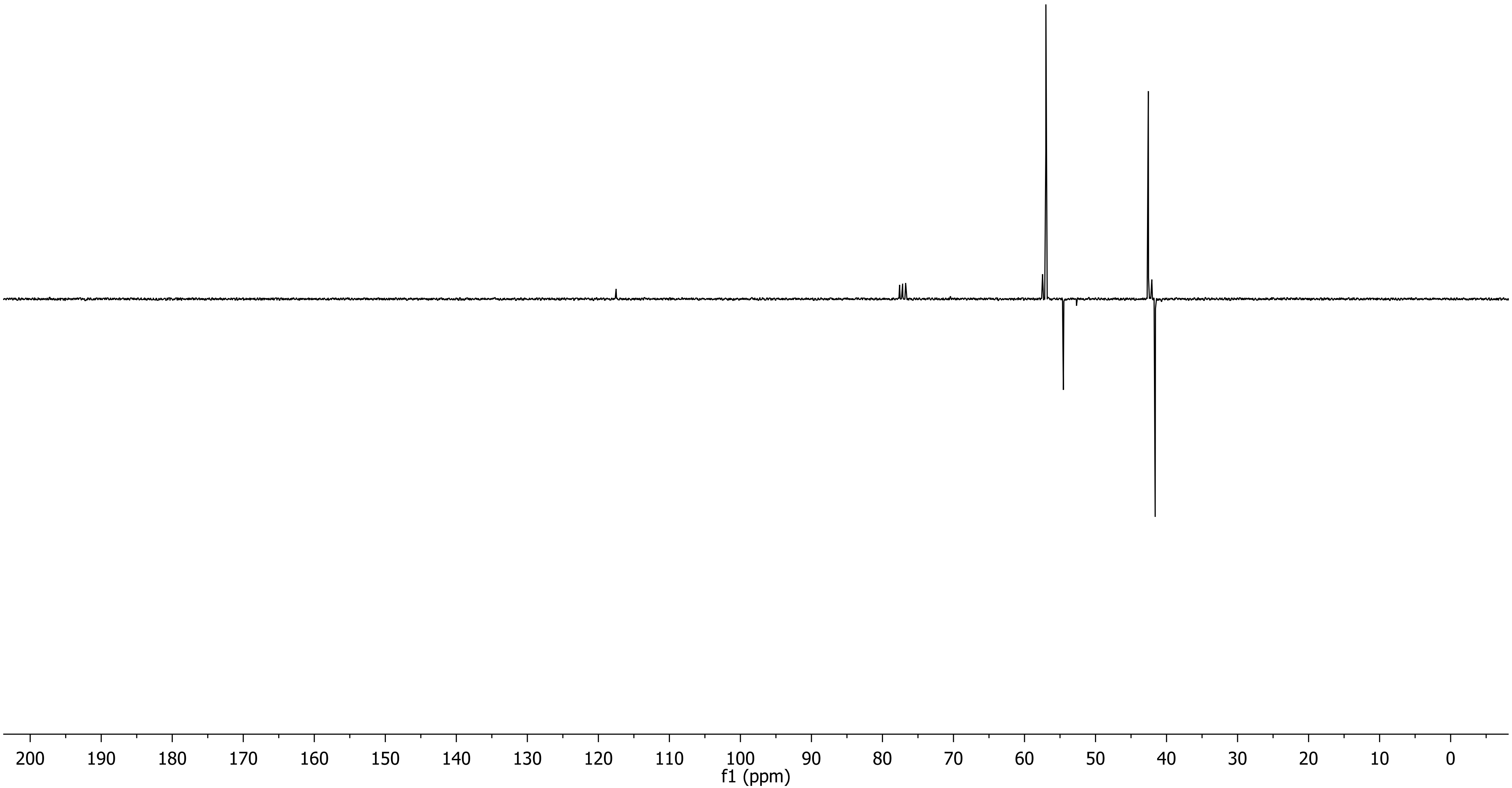
200
190
180
170
160
150
140
130
120
110
100
90
80
70
60
50
40
30
20
10
0
f1 (ppm)

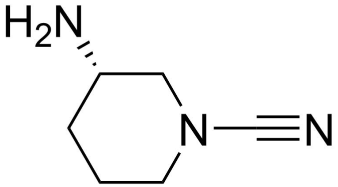

**BB8**

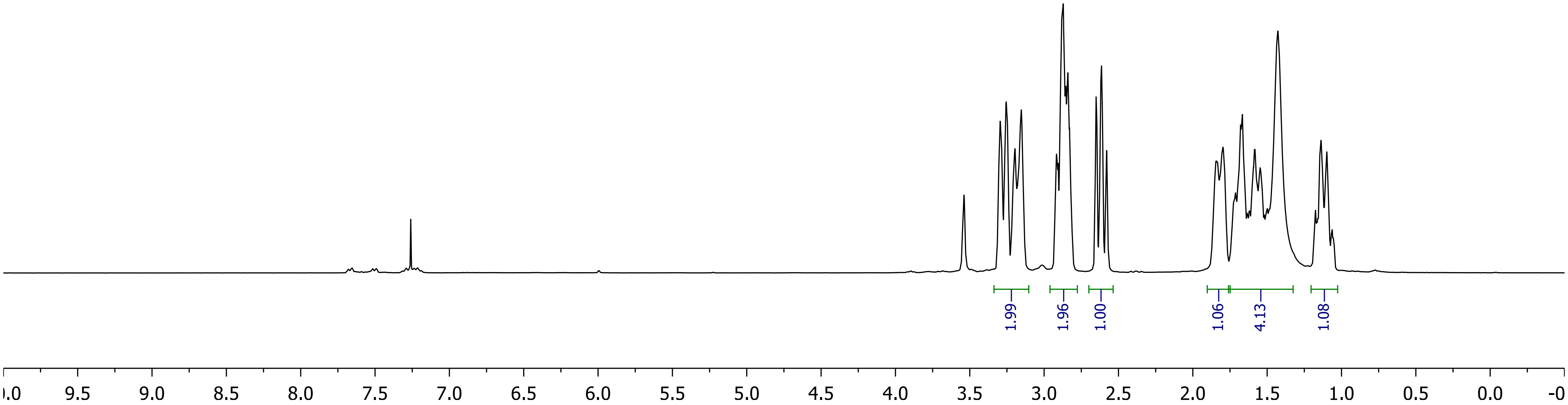
1.99
1.96
1.00
1.06
4.13
1.08
9.5
9.0
8.5
8.0
7.5
7.0
6.5
6.0
5.5
5.0
4.5
4.0
3.5
3.0
2.5
2.0
1.5
1.0
0.5
0.0

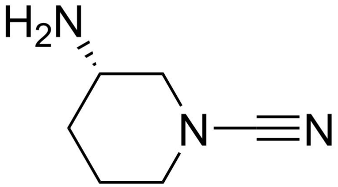
H2N
N
N

**BB8**

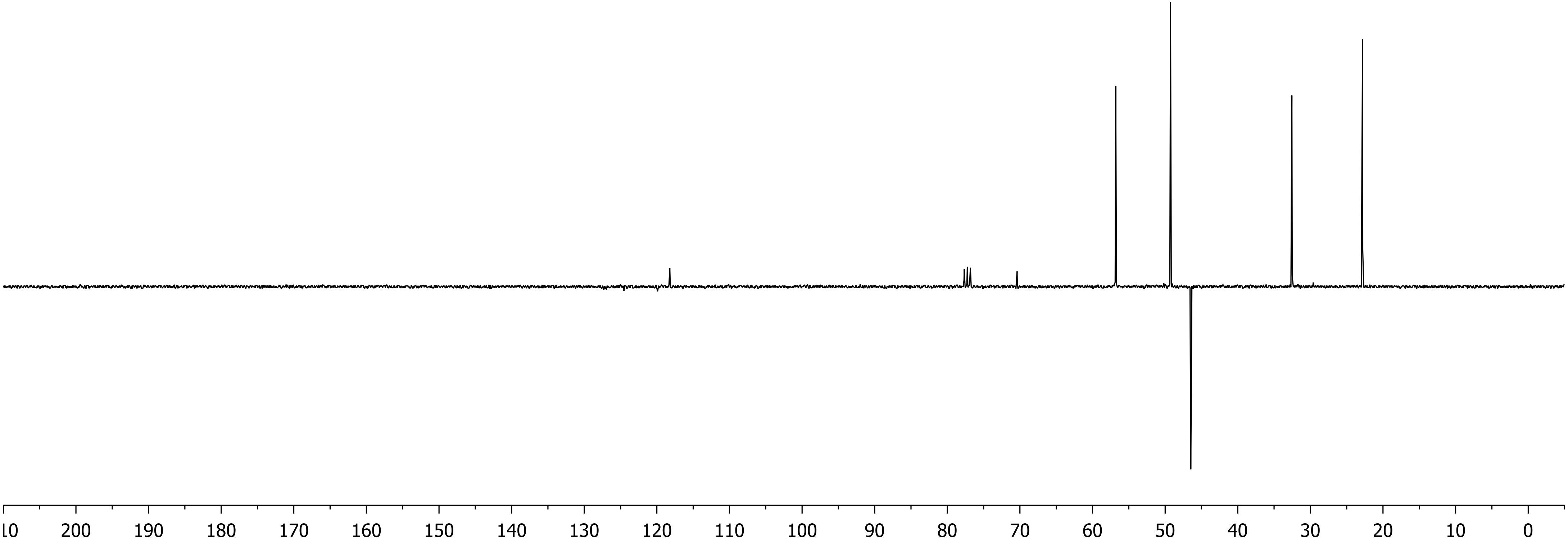
200
190
180
170
160
150
140
130
120
110
100
90
80
70
60
50
40
30
20
10
0

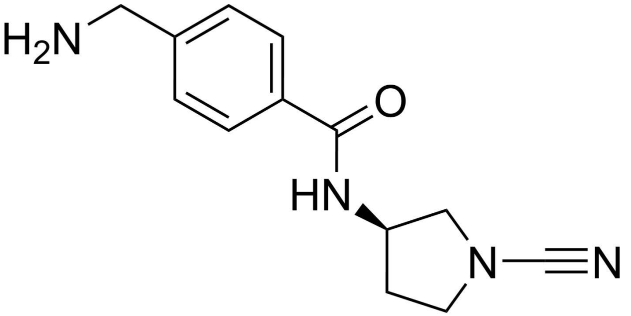
H2N
O
HN
N
N

**BB9**

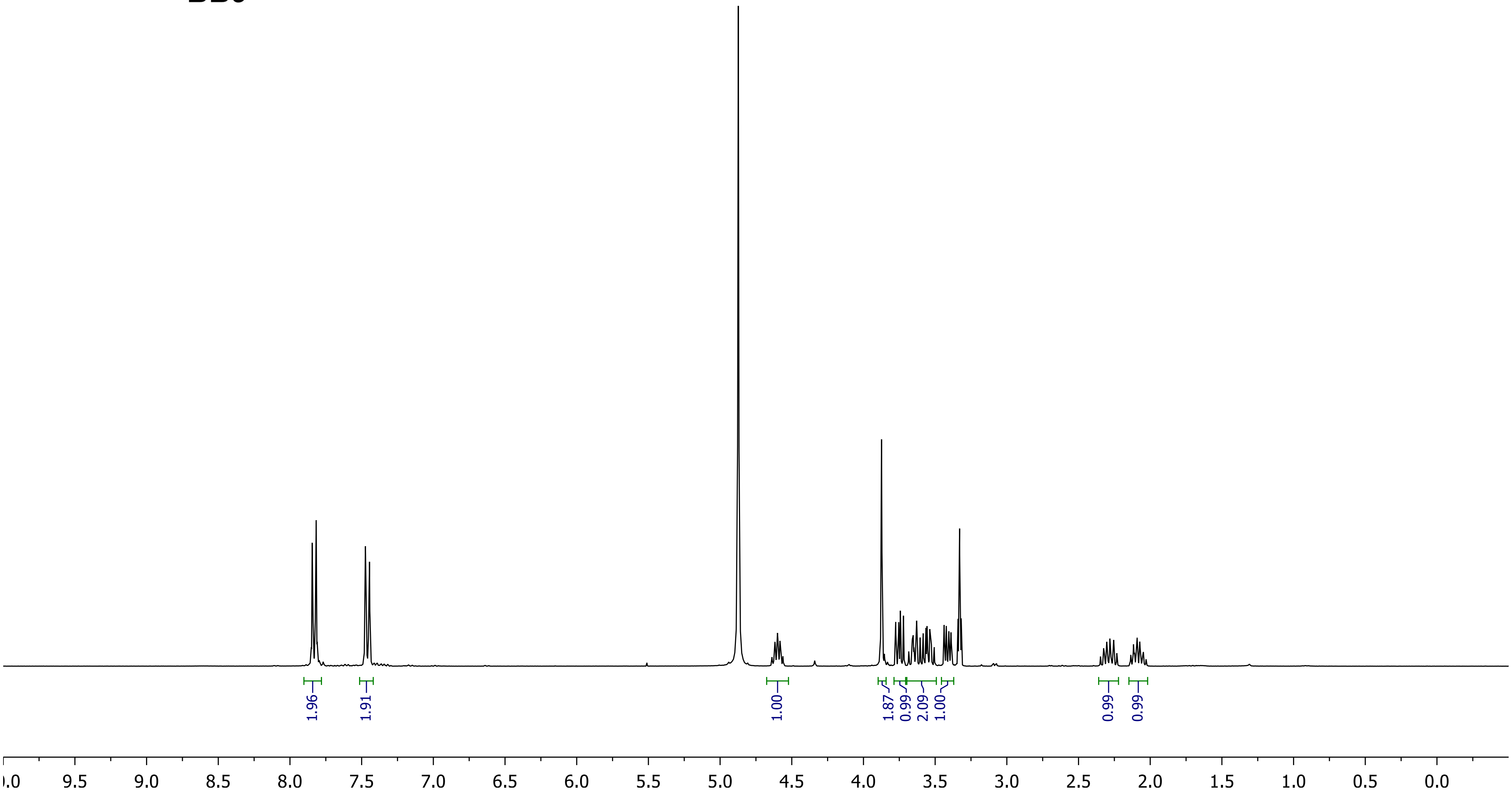
1.96
1.91
1.00
1.87
0.99
2.09
1.00
0.99
0.99
9.5
9.0
8.5
8.0
7.5
7.0
6.5
6.0
5.5
5.0
4.5
4.0
3.5
3.0
2.5
2.0
1.5
1.0
0.5
0.0

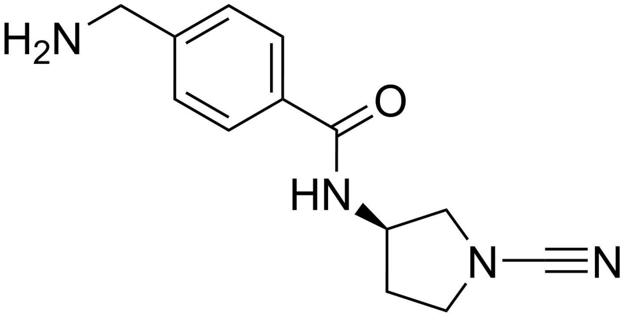
H2N
O
HN
N
N

**BB9**

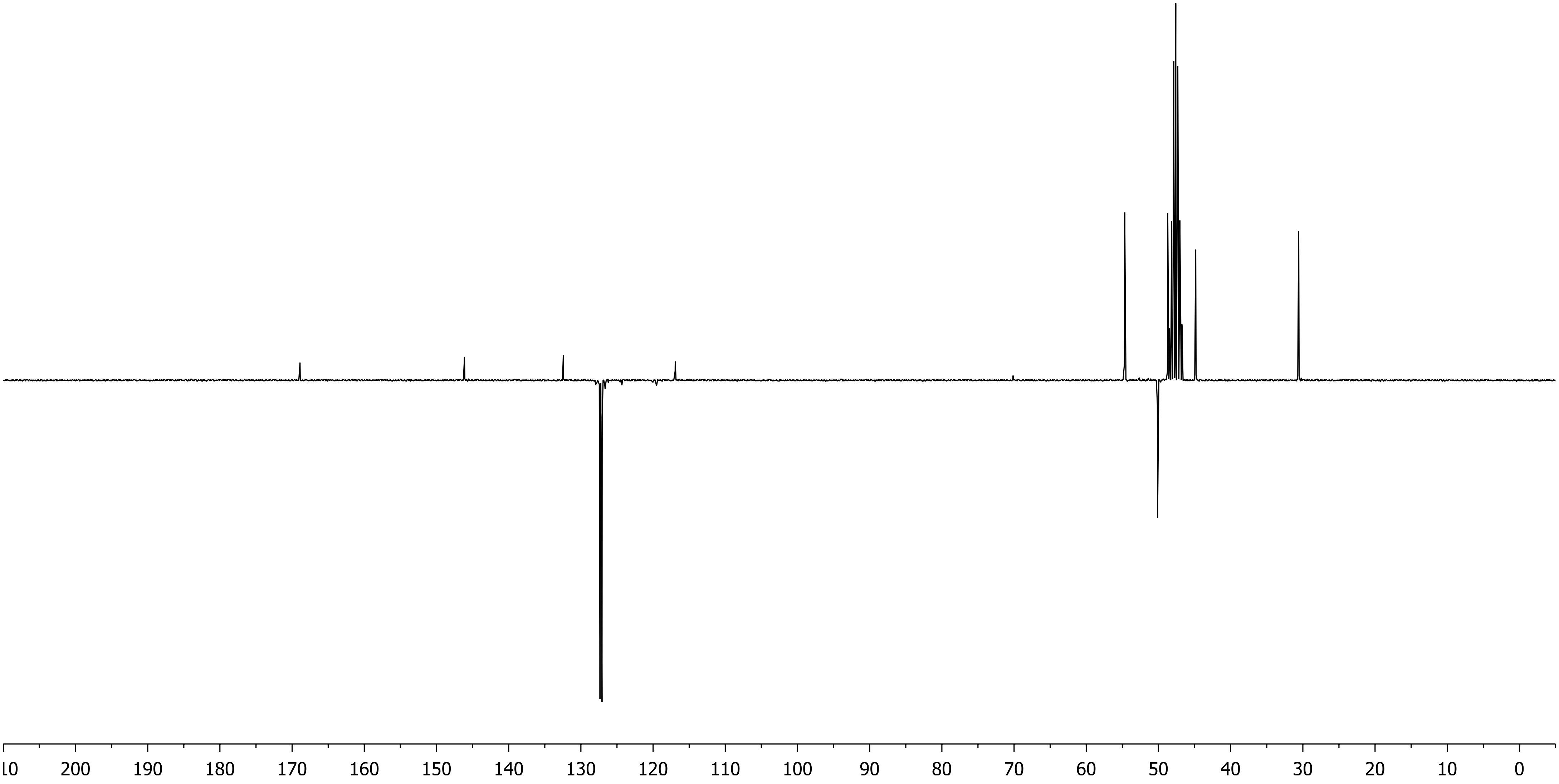
200
190
180
170
160
150
140
130
120
110
100
90
80
70
60
50
40
30
20
10
0

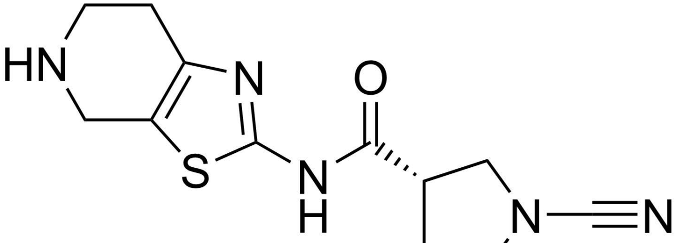
HN
N
S
N
H
O
N
N

**BB10**

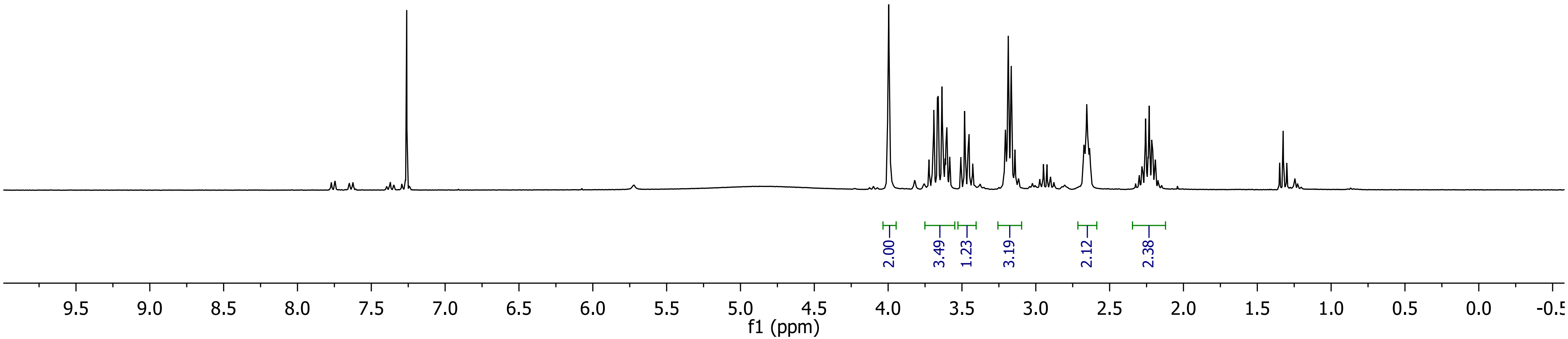
2.00
3.49
1.23
3.19
2.12
2.38
9.5
9.0
8.5
8.0
7.5
7.0
6.5
6.0
5.5
5.0
4.5
4.0
3.5
3.0
2.5
2.0
1.5
1.0
0.5
0.0
f1 (ppm)

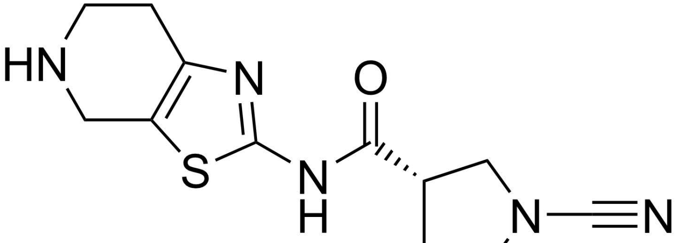
HN
N
O
S
N
H
N
N

**BB10**

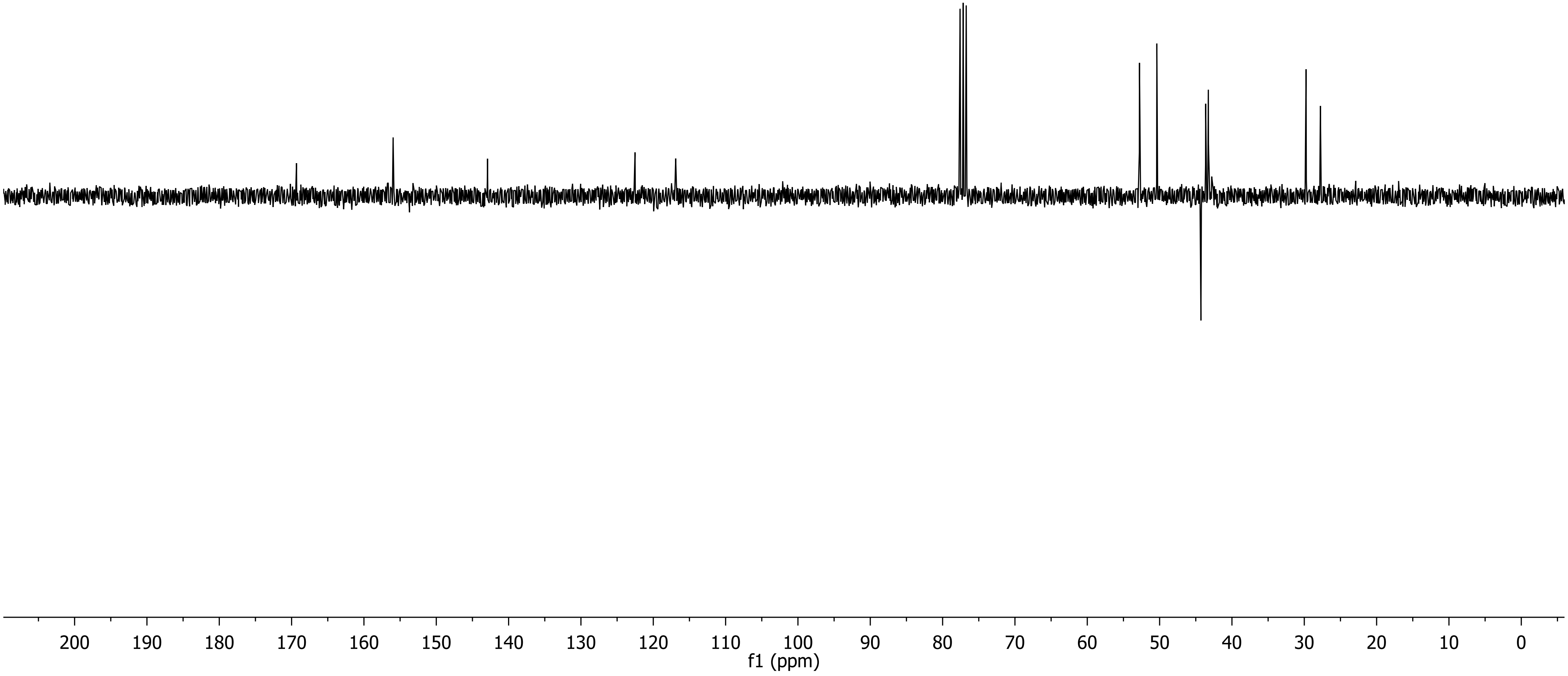
200 190 180 170 160 150 140 130 120 110 100 90 80 70 60 50 40 30 20 10 0
f1 (ppm)

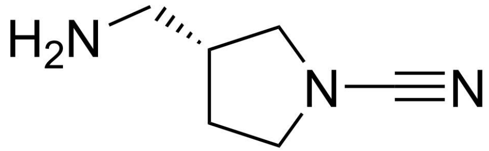
H$_2$N
N
N

**BB11**

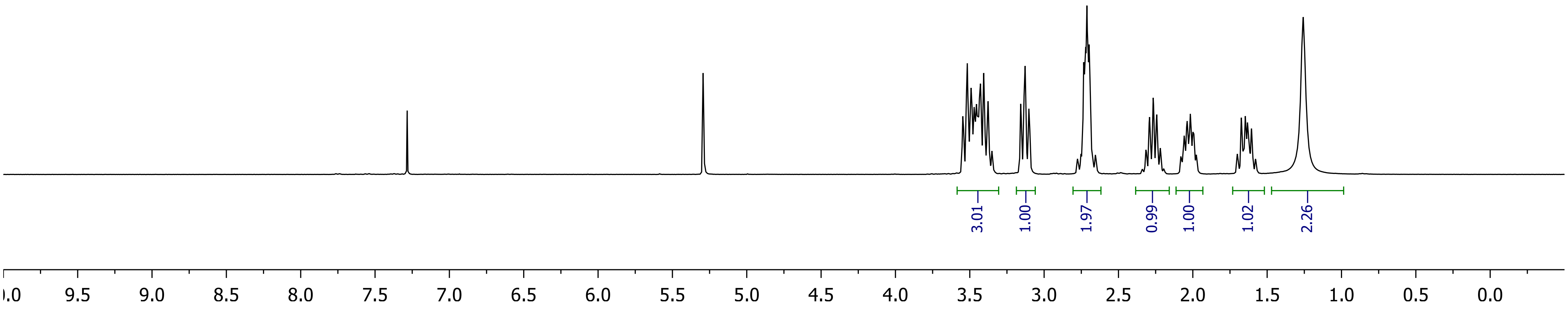
3.01
1.00
1.97
0.99
1.00
1.02
2.26
9.5 9.0 8.5 8.0 7.5 7.0 6.5 6.0 5.5 5.0 4.5 4.0 3.5 3.0 2.5 2.0 1.5 1.0 0.5 0.0

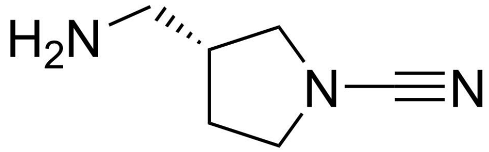

**BB11**

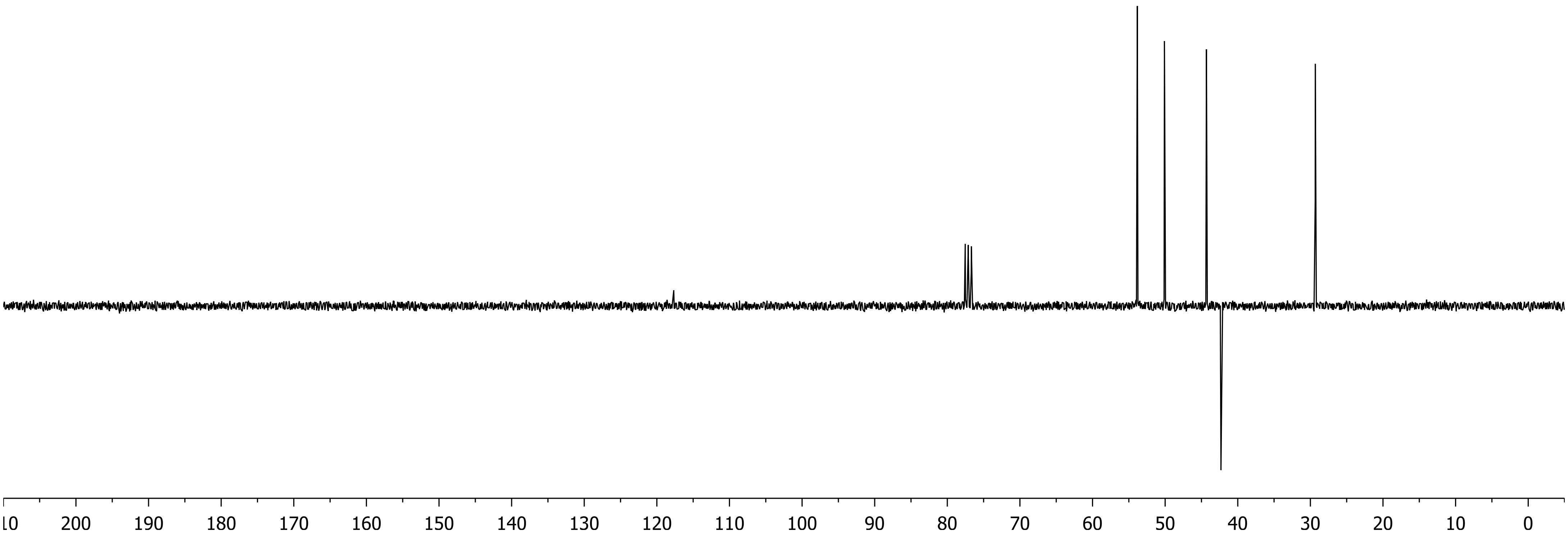

10
200
190
180
170
160
150
140
130
120
110
100
90
80
70
60
50
40
30
20
10
0

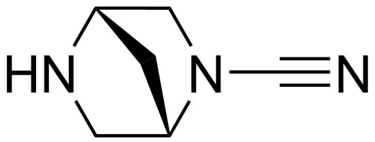

**BB12**

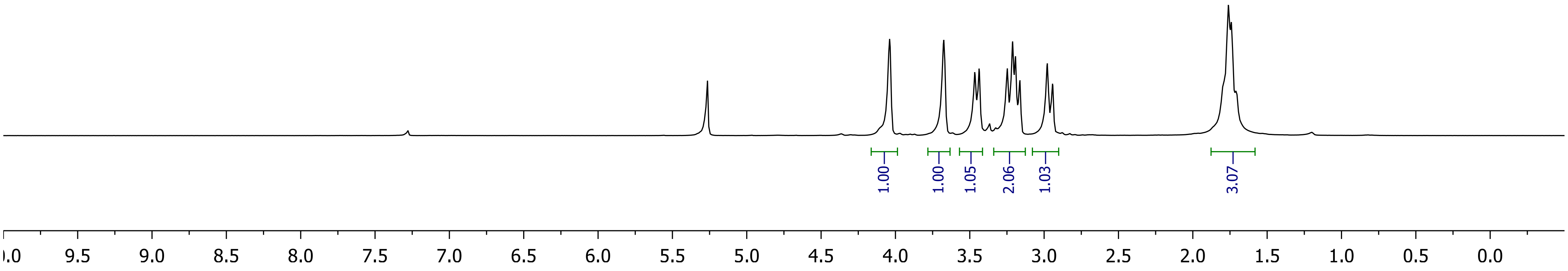

1.00
1.00
1.05
2.06
1.03
3.07
.0
9.5
9.0
8.5
8.0
7.5
7.0
6.5
6.0
5.5
5.0
4.5
4.0
3.5
3.0
2.5
2.0
1.5
1.0
0.5
0.0

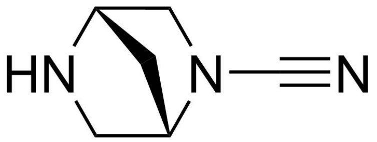

**BB12**

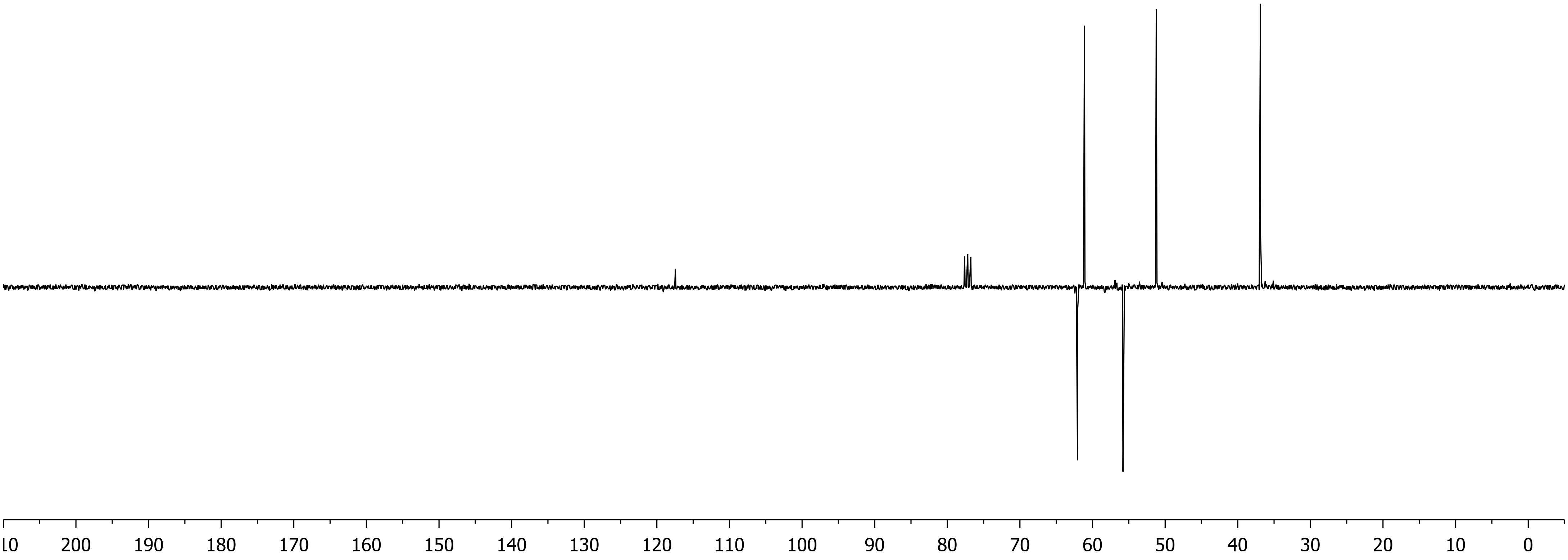

10
200
190
180
170
160
150
140
130
120
110
100
90
80
70
60
50
40
30
20
10
0

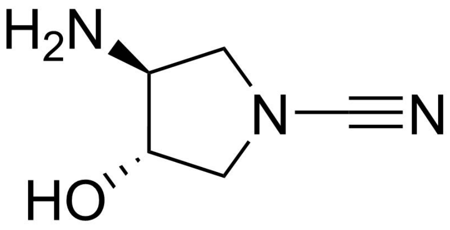

**BB13**

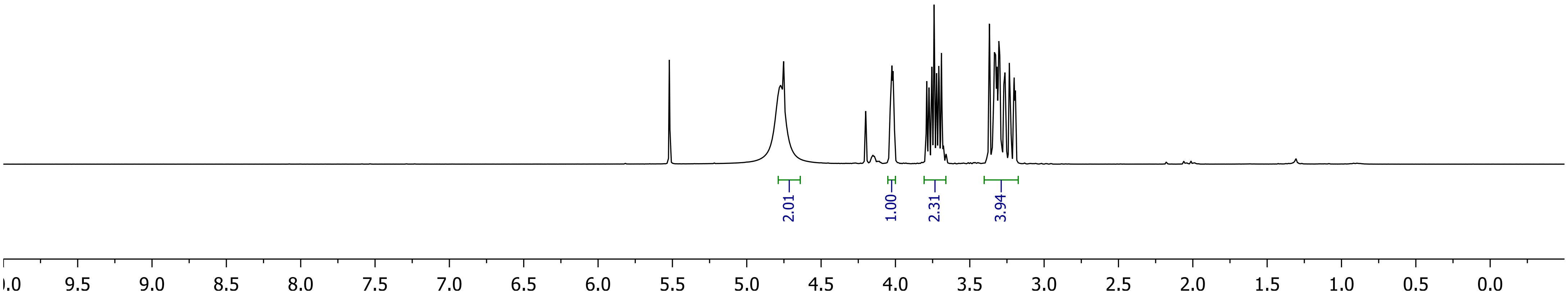
2.01
1.00
2.31
3.94
0.0
9.5
9.0
8.5
8.0
7.5
7.0
6.5
6.0
5.5
5.0
4.5
4.0
3.5
3.0
2.5
2.0
1.5
1.0
0.5
0.0

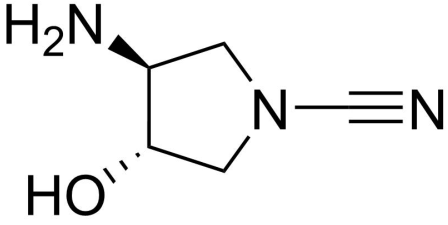

**BB13**

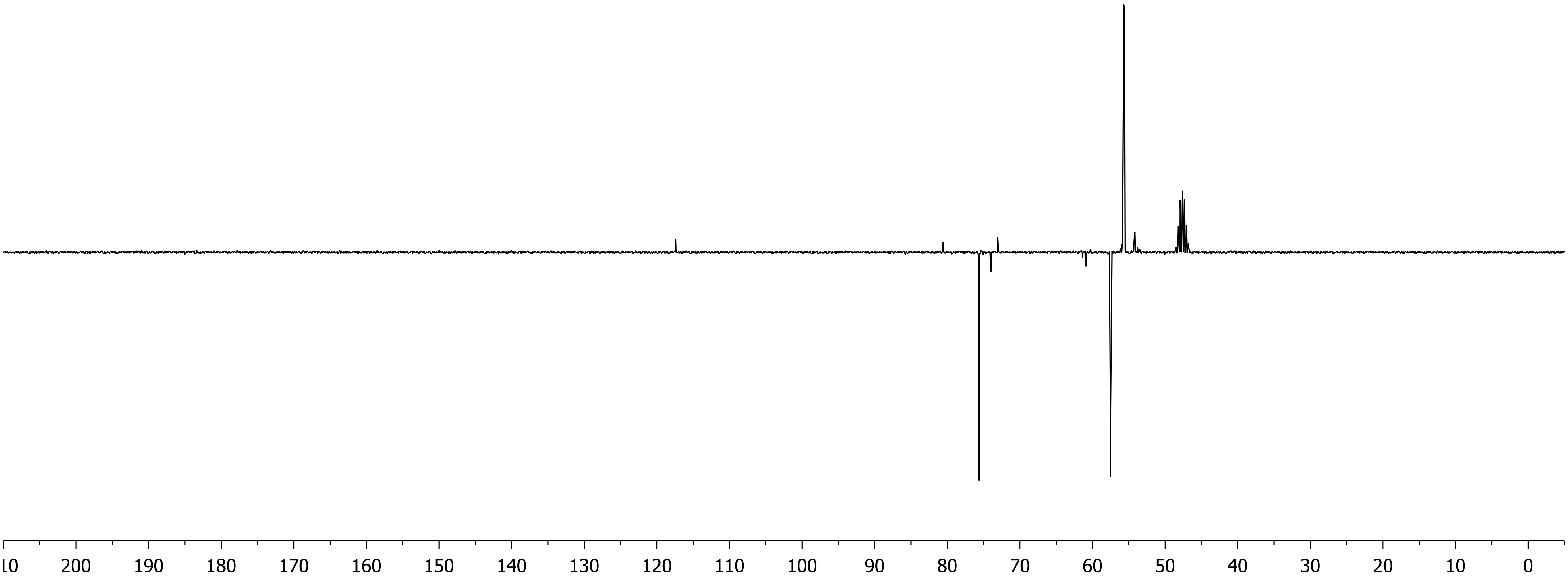

10 200 190 180 170 160 150 140 130 120 110 100 90 80 70 60 50 40 30 20 10 0

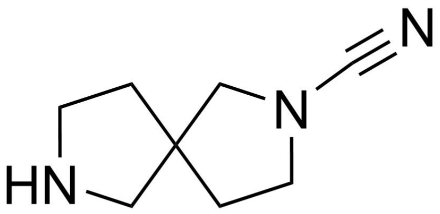
N
N
HN

**BB14**

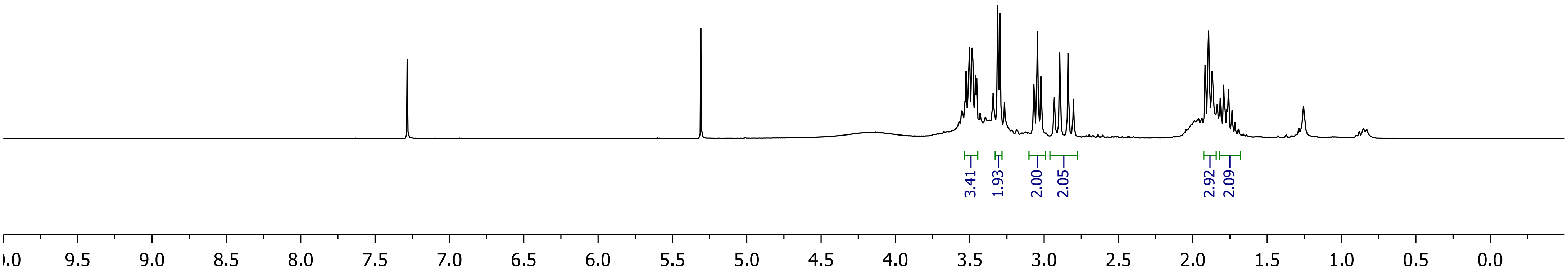
3.41
1.93
2.00
2.05
2.92
2.09
9.5 9.0 8.5 8.0 7.5 7.0 6.5 6.0 5.5 5.0 4.5 4.0 3.5 3.0 2.5 2.0 1.5 1.0 0.5 0.0

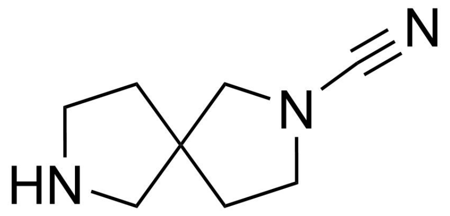

**BB14**

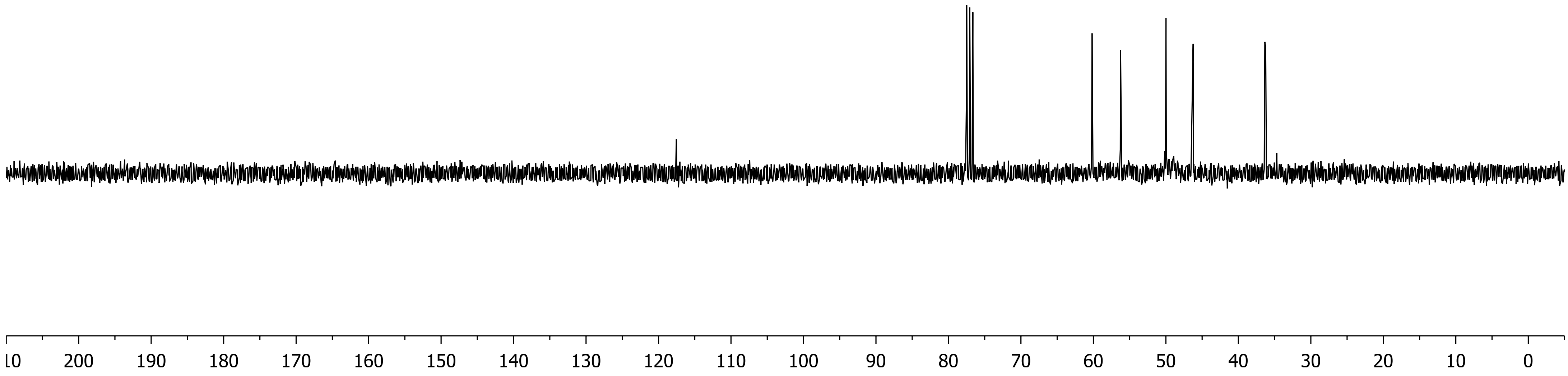
200
190
180
170
160
150
140
130
120
110
100
90
80
70
60
50
40
30
20
10
0

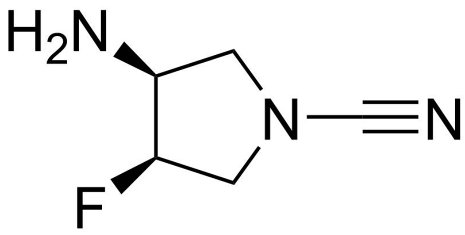

H2N
N
N
F


**BB15**

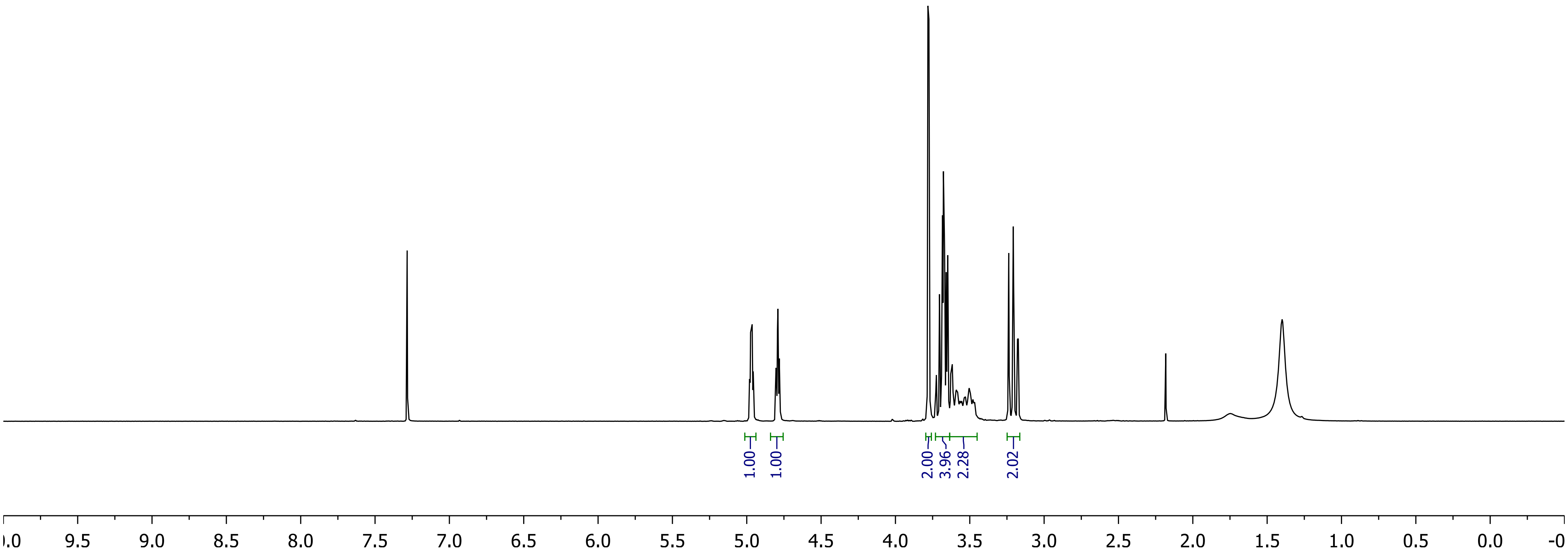

1.00
1.00
2.00
3.96
2.28
2.02
9.5
9.0
8.5
8.0
7.5
7.0
6.5
6.0
5.5
5.0
4.5
4.0
3.5
3.0
2.5
2.0
1.5
1.0
0.5
0.0

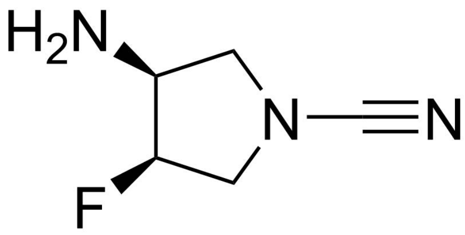

**BB15**

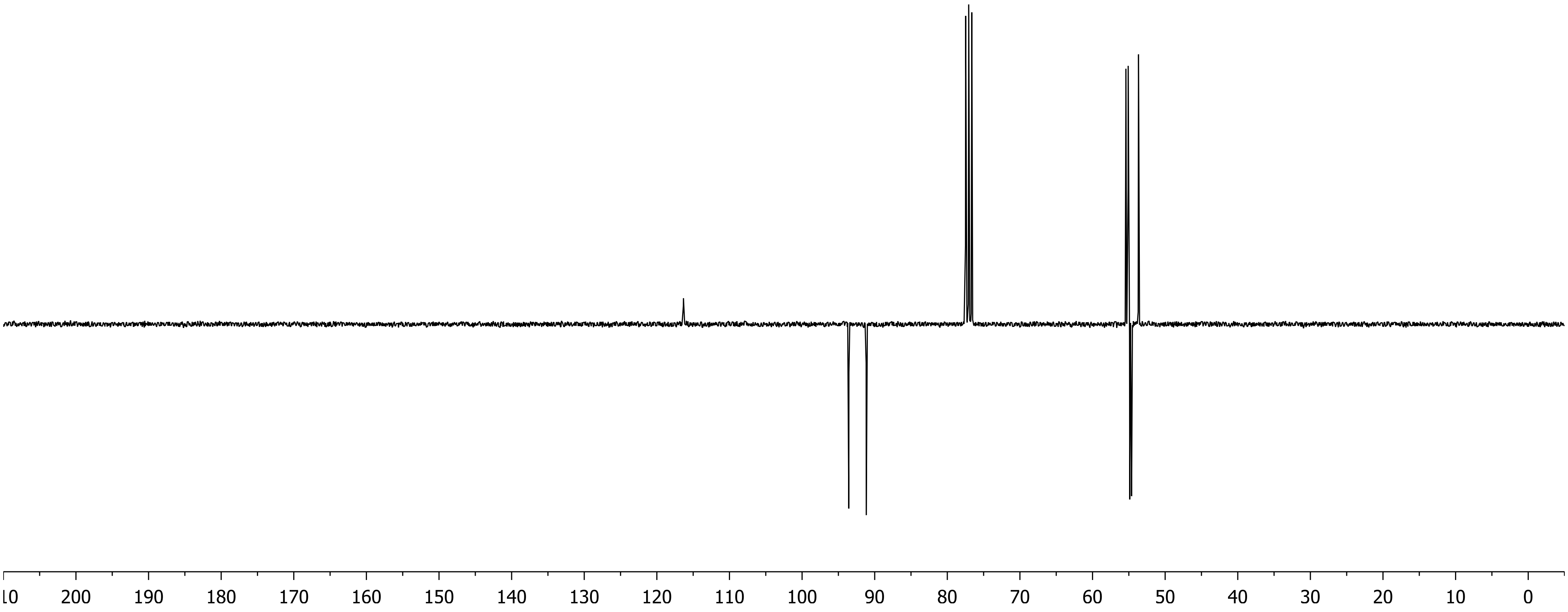
200
190
180
170
160
150
140
130
120
110
100
90
80
70
60
50
40
30
20
10
0

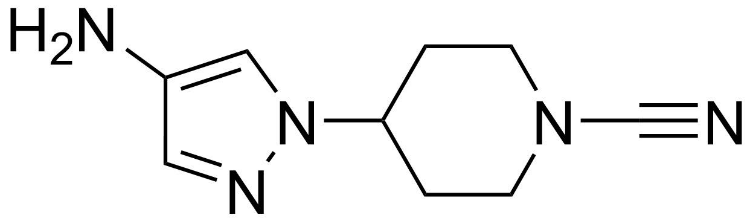
H2N
N
N
N
N

**BB16**

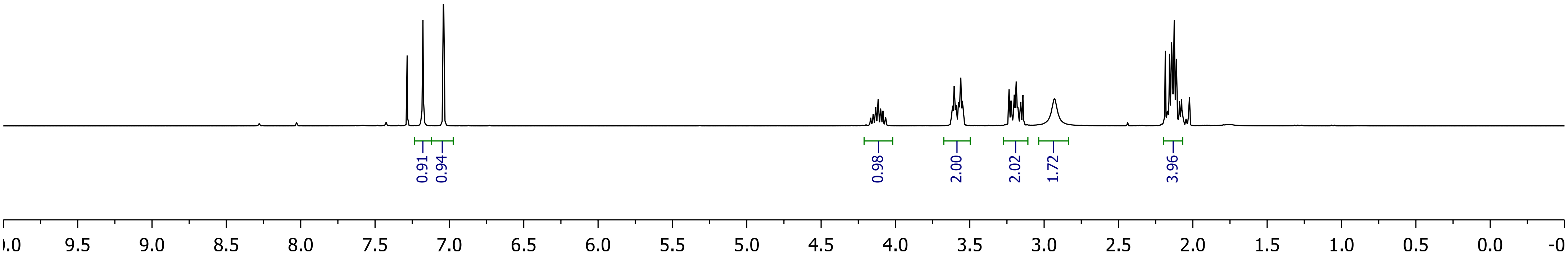
0.91
0.94
0.98
2.00
2.02
1.72
3.96
9.5
9.0
8.5
8.0
7.5
7.0
6.5
6.0
5.5
5.0
4.5
4.0
3.5
3.0
2.5
2.0
1.5
1.0
0.5
0.0

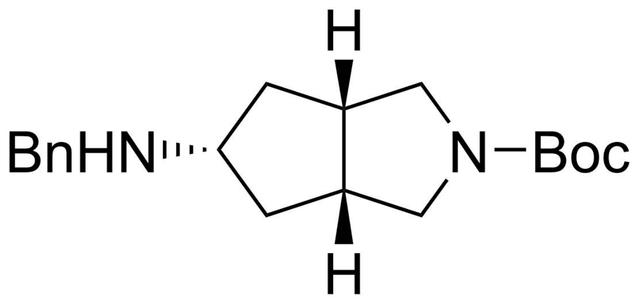
H
BnHN
N−Boc
H

**6**

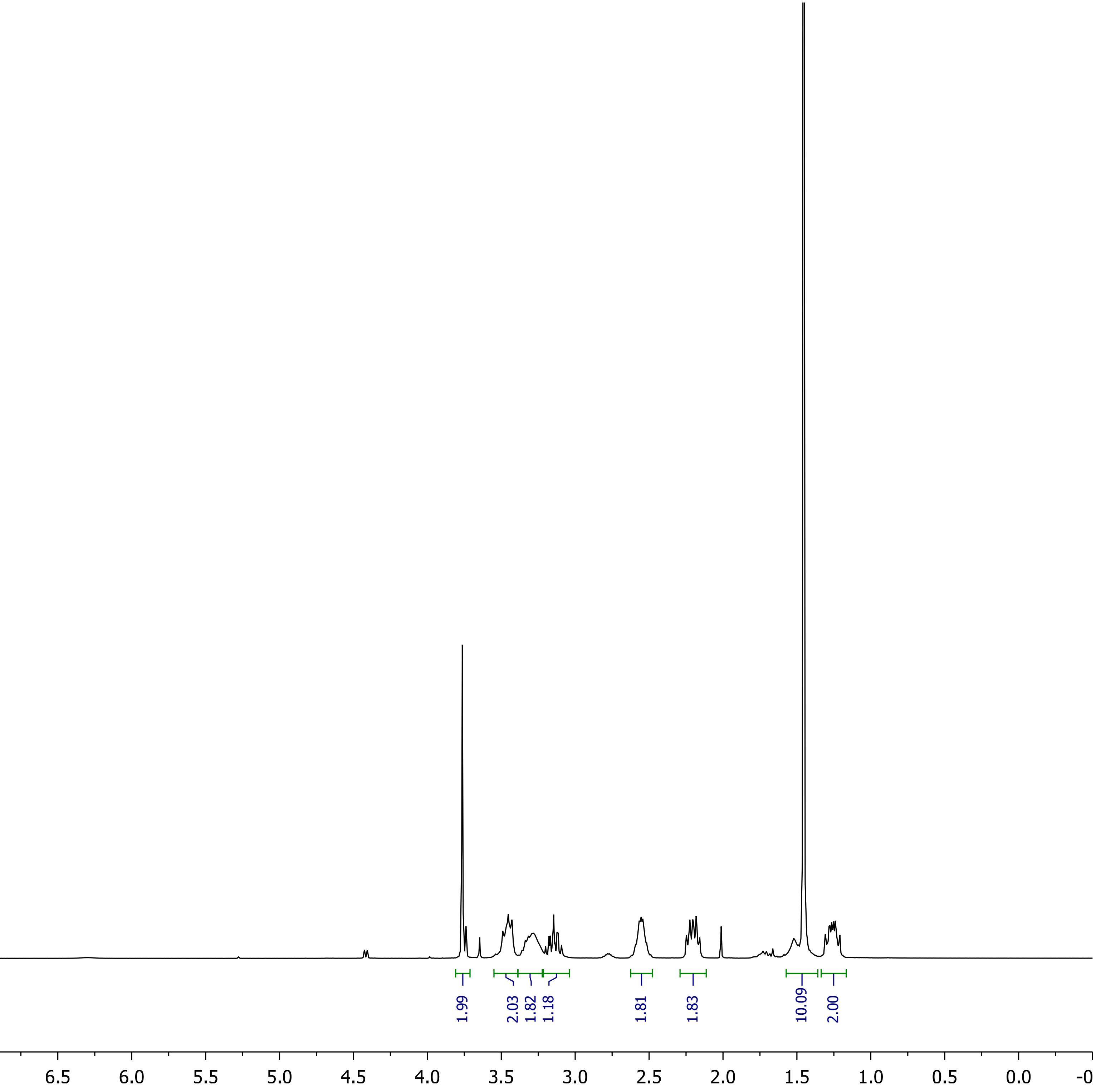
3.93
0.99
1.99
2.03
1.82
1.18
1.81
1.83
10.09
2.00
9.5 9.0 8.5 8.0 7.5 7.0 6.5 6.0 5.5 5.0 4.5 4.0 3.5 3.0 2.5 2.0 1.5 1.0 0.5 0.0

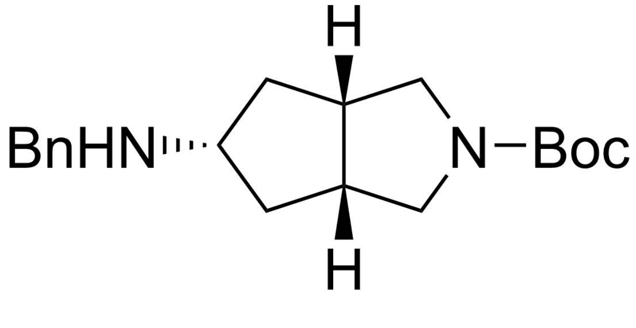
BnHN
H
N−Boc
H

**6**

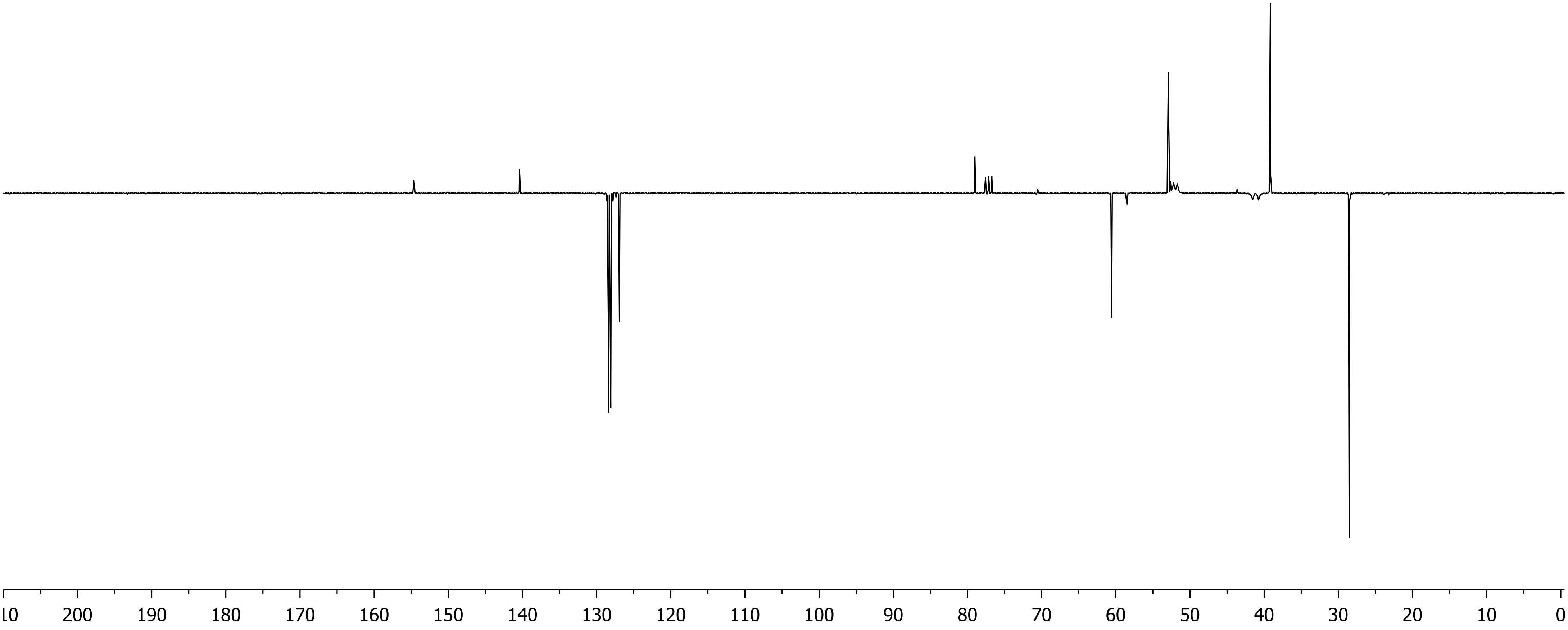
200 190 180 170 160 150 140 130 120 110 100 90 80 70 60 50 40 30 20 10 0

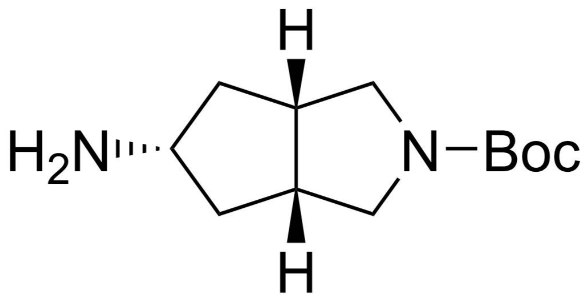
H
H₂N
N
Boc
H

**7**

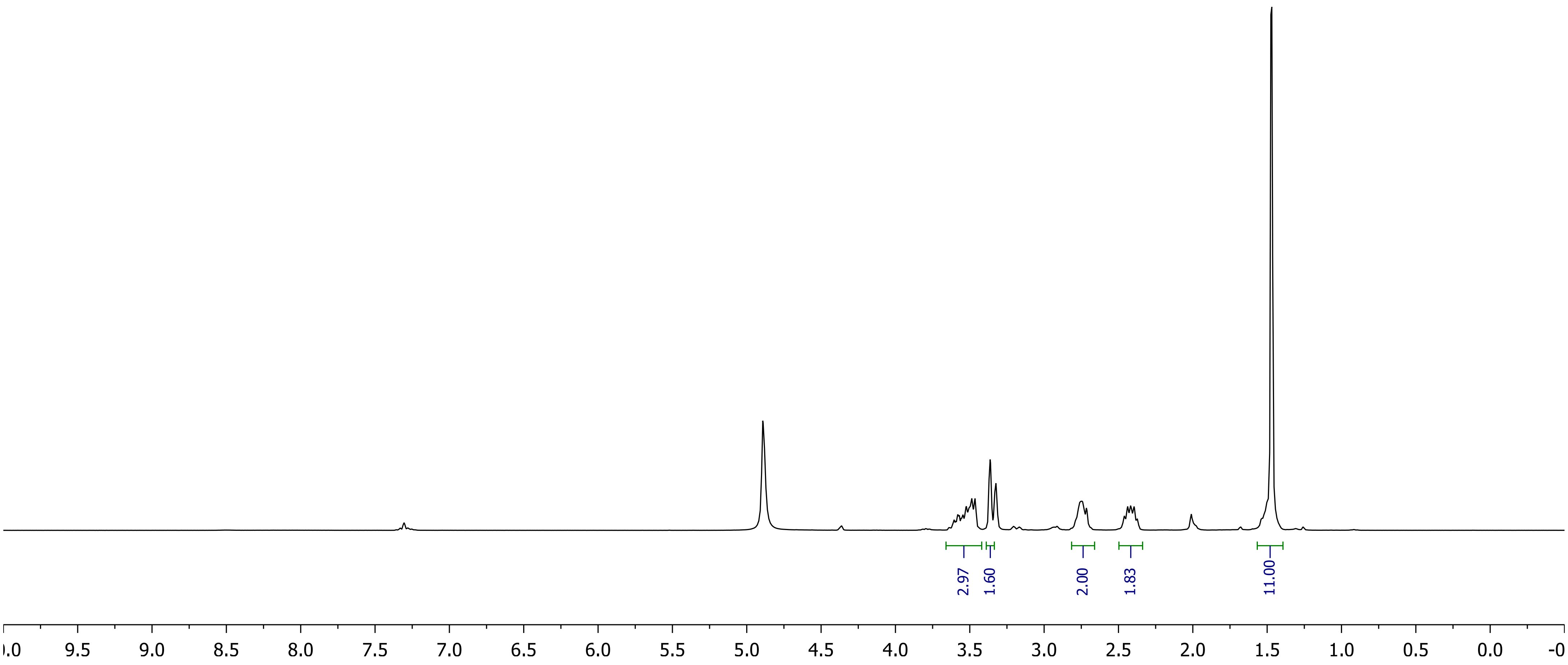
2.97
1.60
2.00
1.83
11.00
9.5
9.0
8.5
8.0
7.5
7.0
6.5
6.0
5.5
5.0
4.5
4.0
3.5
3.0
2.5
2.0
1.5
1.0
0.5
0.0

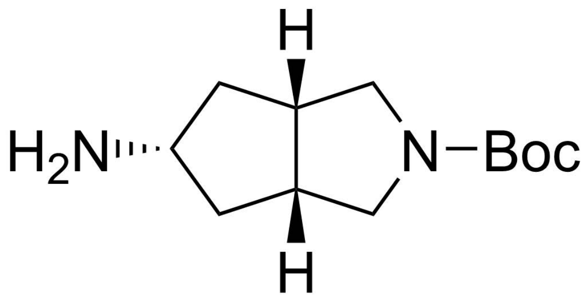
H
H2N
N–Boc
H

**7**

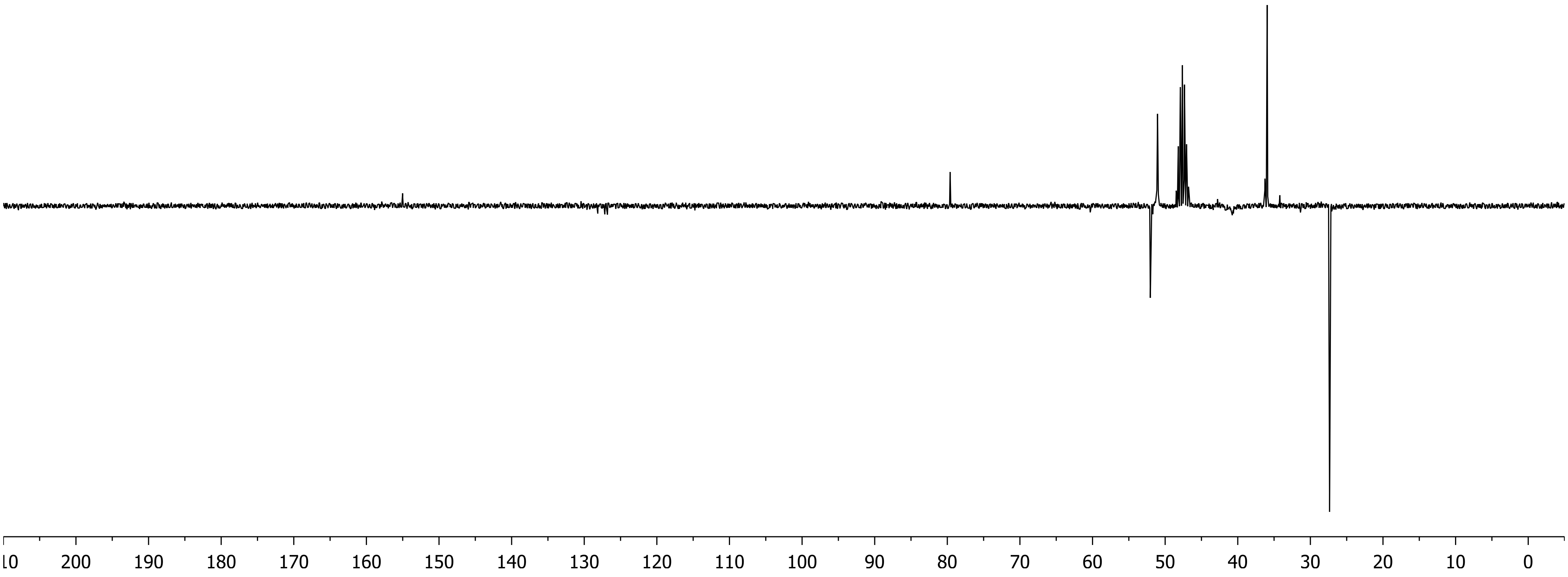
200 190 180 170 160 150 140 130 120 110 100 90 80 70 60 50 40 30 20 10 0

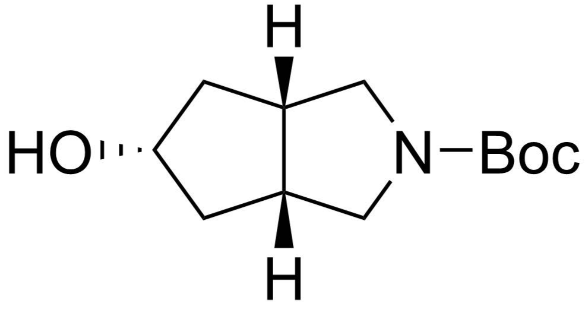
H
HO
N−Boc
H

**8**

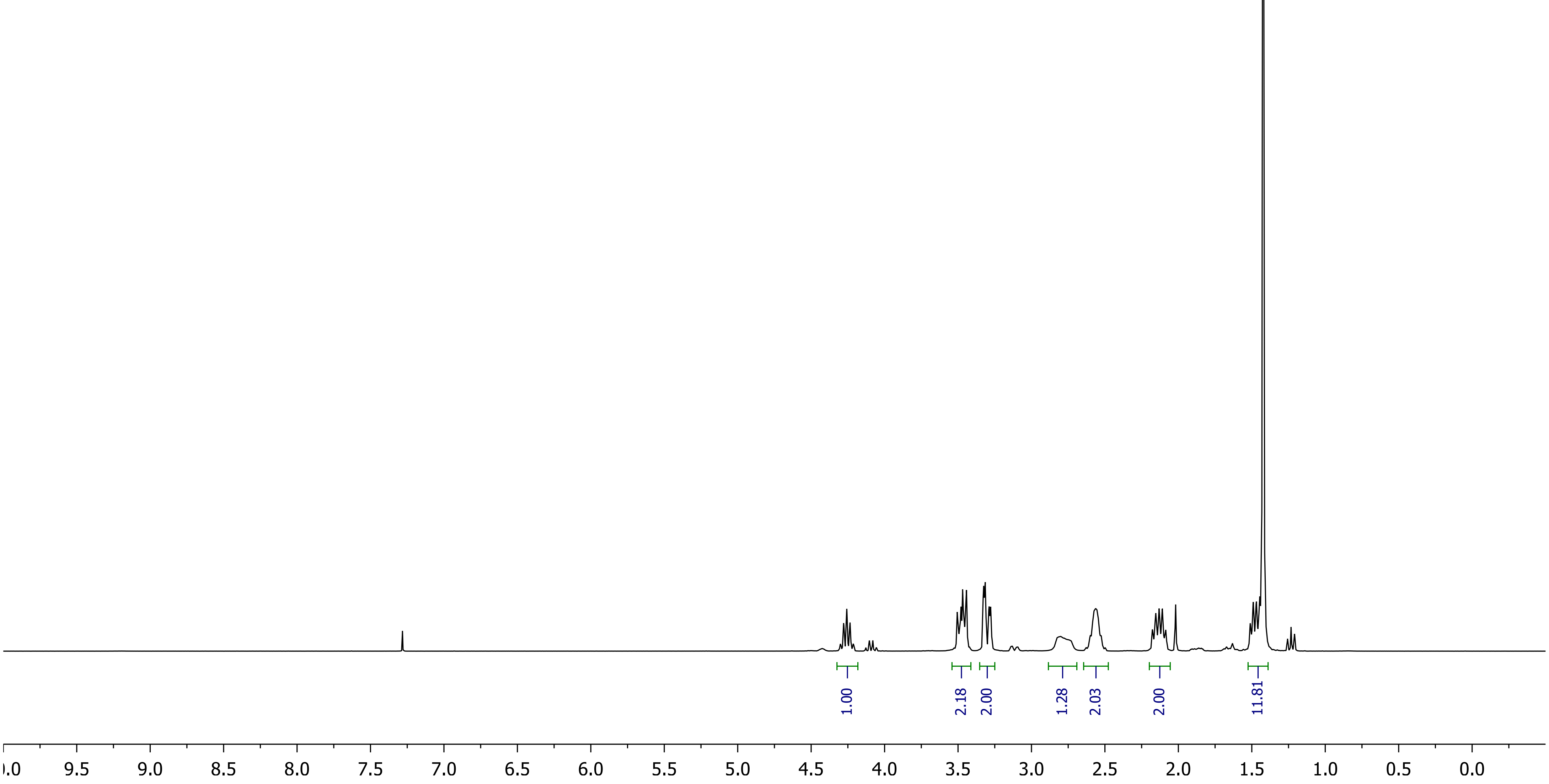
1.00
2.18
2.00
1.28
2.03
2.00
11.81
9.5
9.0
8.5
8.0
7.5
7.0
6.5
6.0
5.5
5.0
4.5
4.0
3.5
3.0
2.5
2.0
1.5
1.0
0.5
0.0

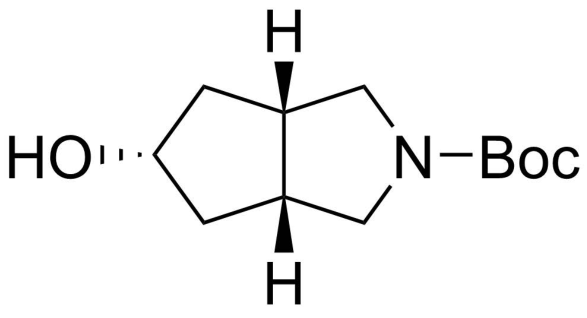

**8**

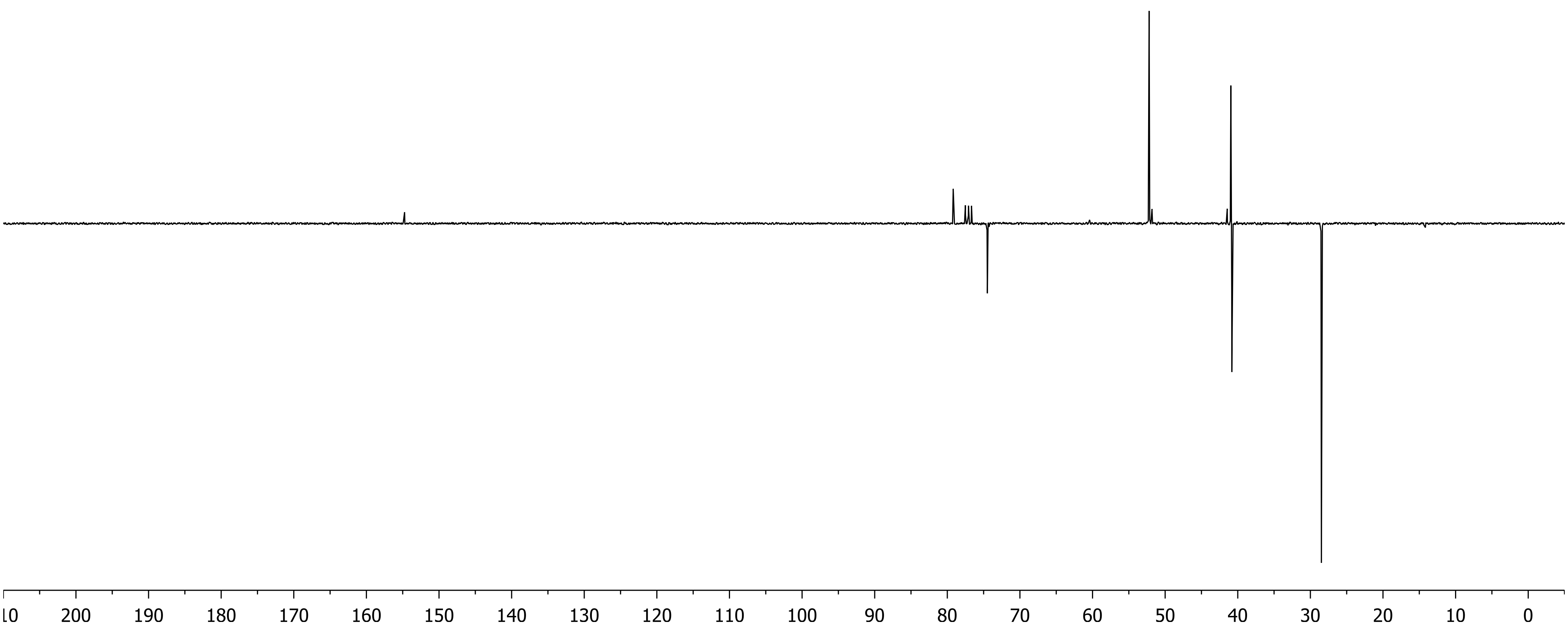
10
200
190
180
170
160
150
140
130
120
110
100
90
80
70
60
50
40
30
20
10
0

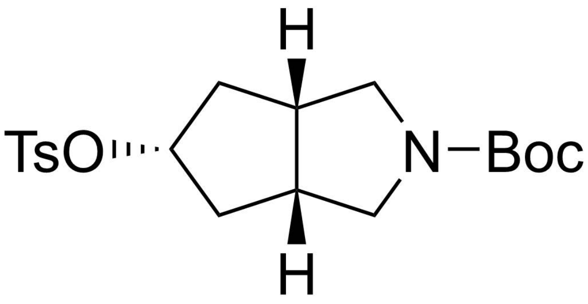

**9**

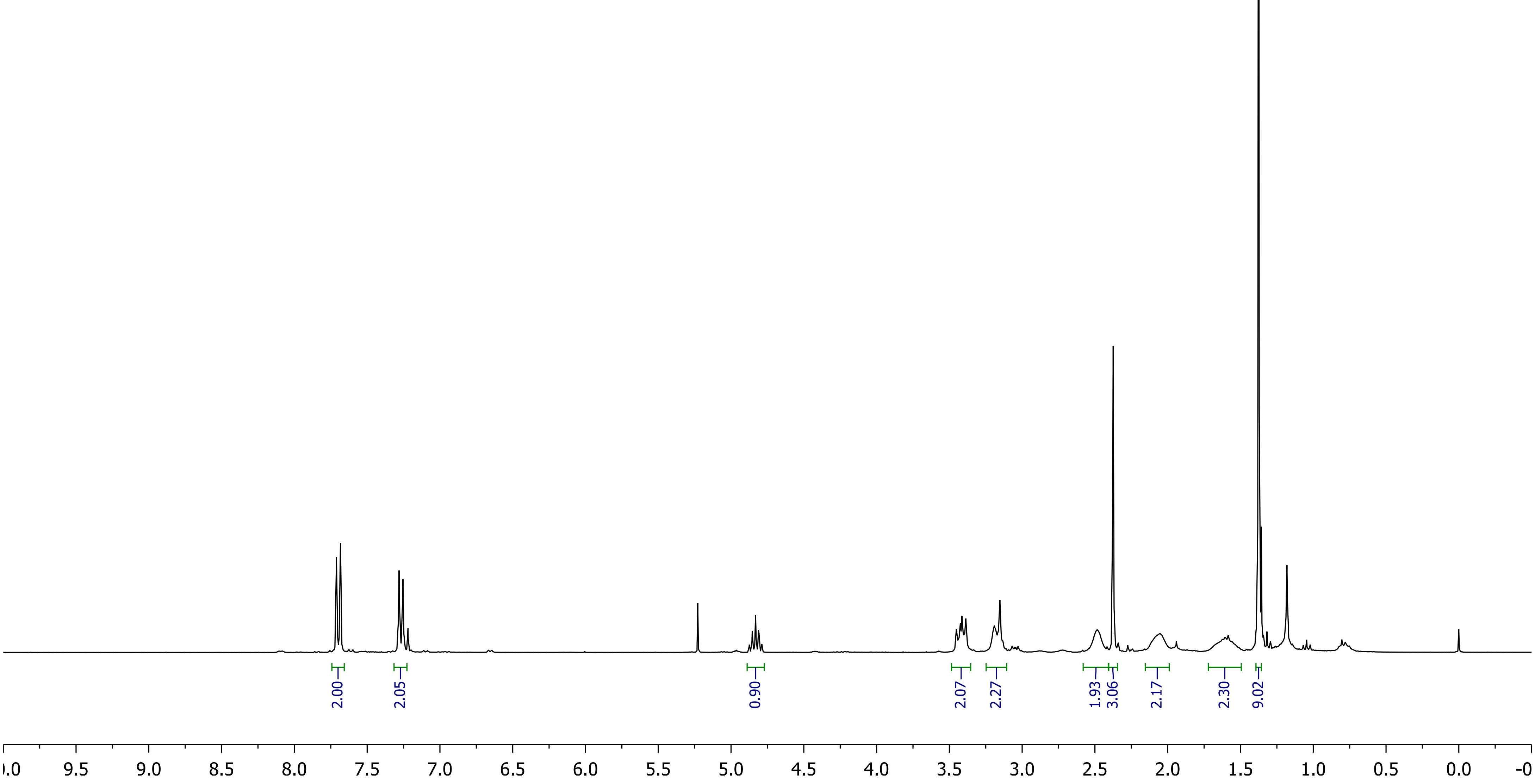
2.00
2.05
0.90
2.07
2.27
1.93
3.06
2.17
2.30
9.02
10.0
9.5
9.0
8.5
8.0
7.5
7.0
6.5
6.0
5.5
5.0
4.5
4.0
3.5
3.0
2.5
2.0
1.5
1.0
0.5
0.0
-0

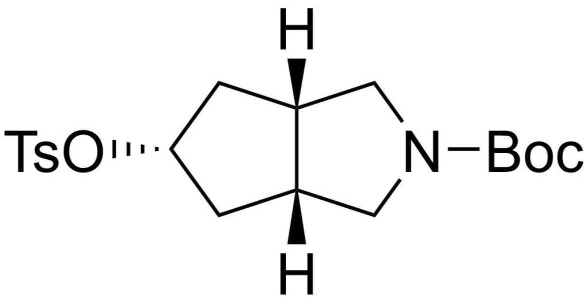

H
TsO
N
Boc
H


**9**

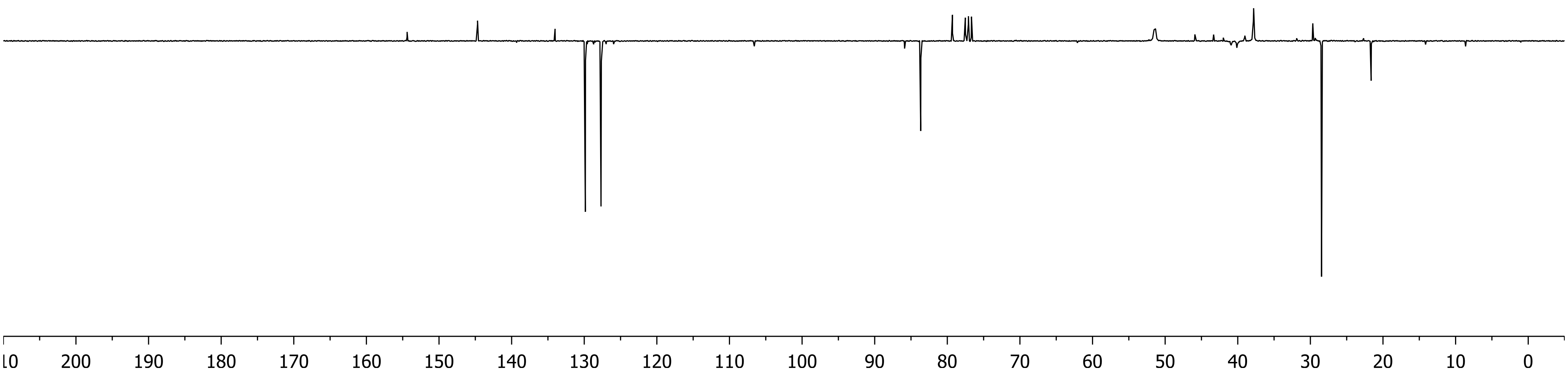

10
200
190
180
170
160
150
140
130
120
110
100
90
80
70
60
50
40
30
20
10
0

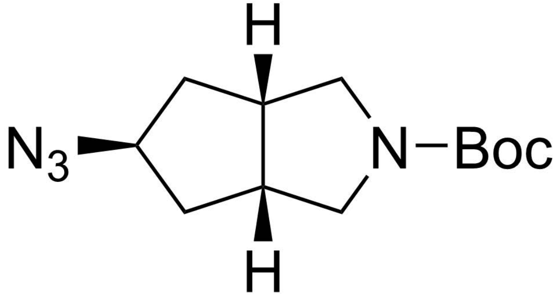
H
N3
N–Boc
H

**10**

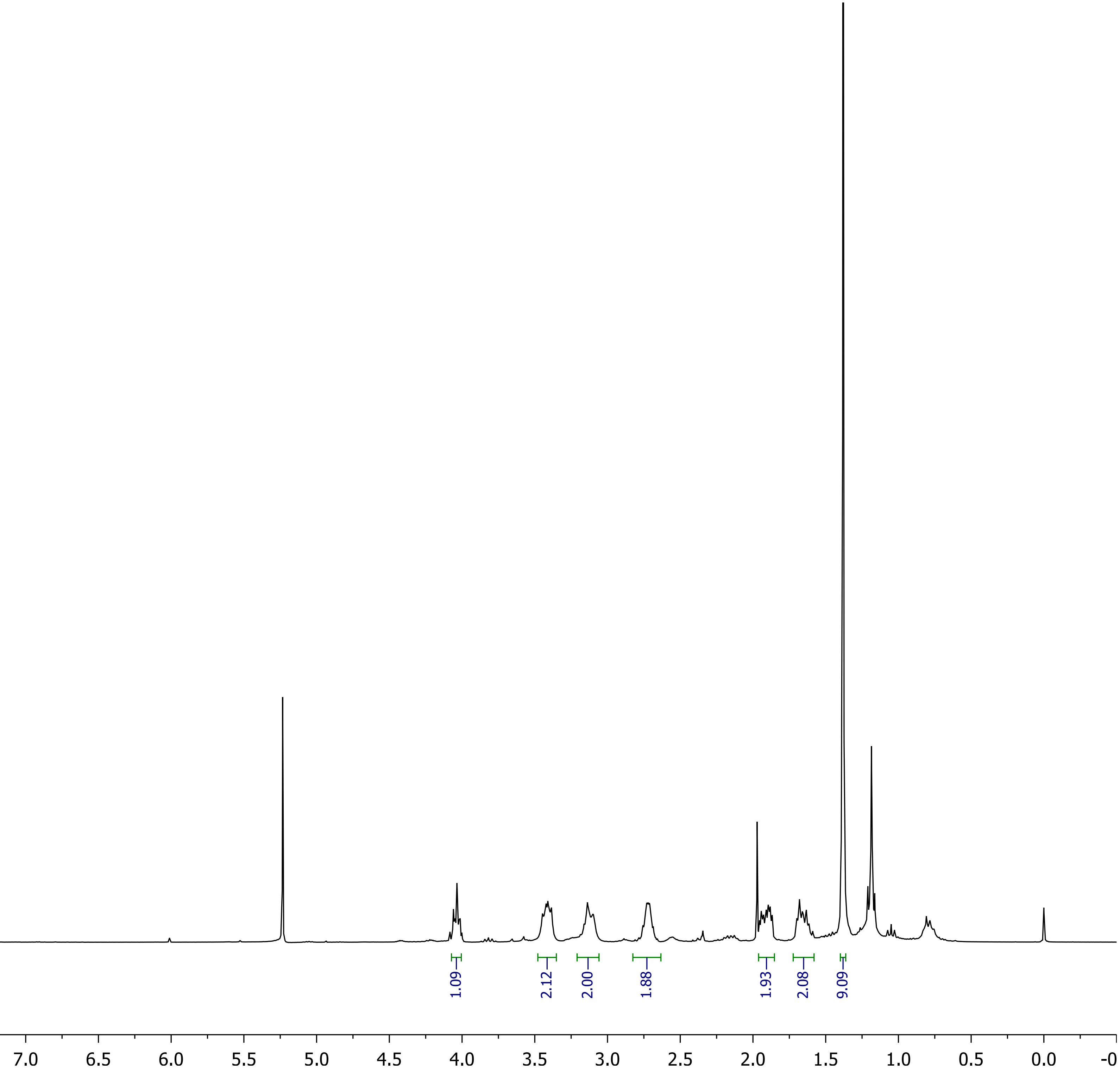
1.09
2.12
2.00
1.88
1.93
2.08
9.09
9.5
9.0
8.5
8.0
7.5
7.0
6.5
6.0
5.5
5.0
4.5
4.0
3.5
3.0
2.5
2.0
1.5
1.0
0.5
0.0

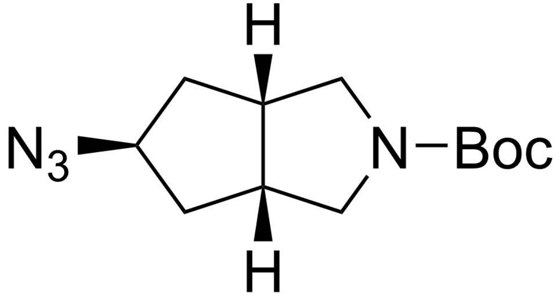

H
N3
N
Boc
H


**10**

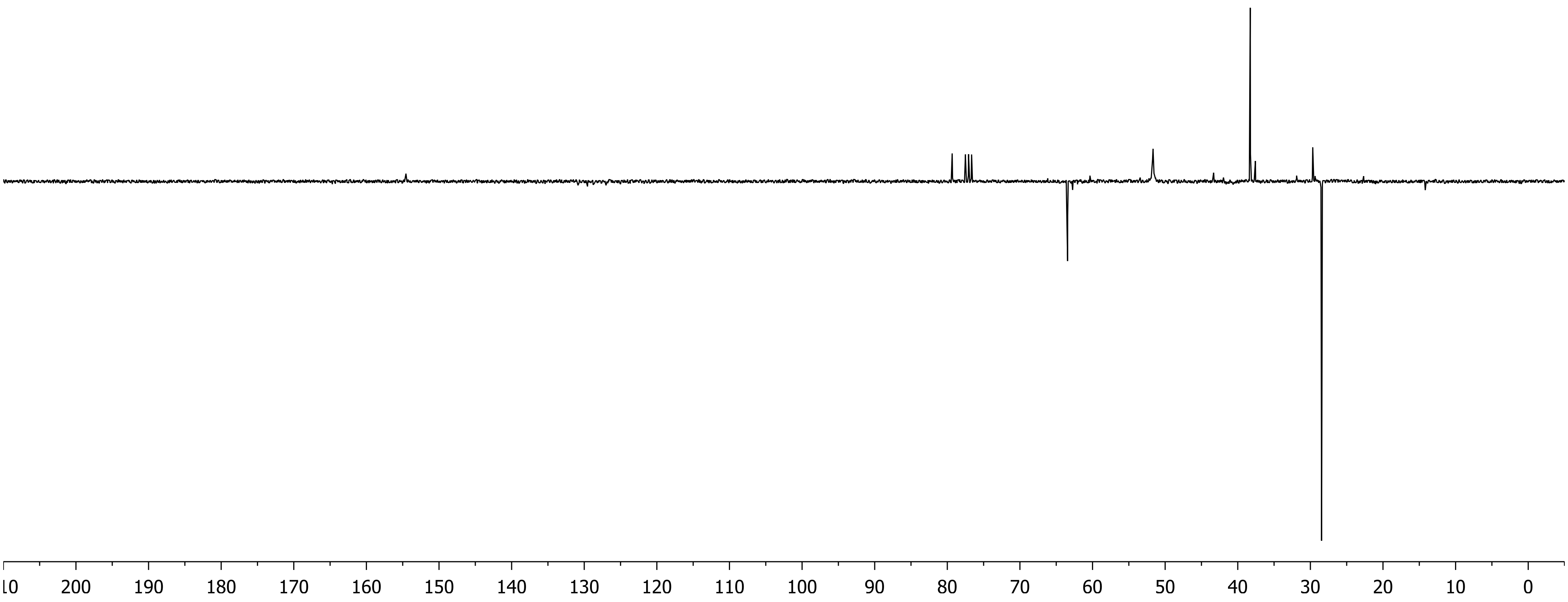

200
190
180
170
160
150
140
130
120
110
100
90
80
70
60
50
40
30
20
10
0

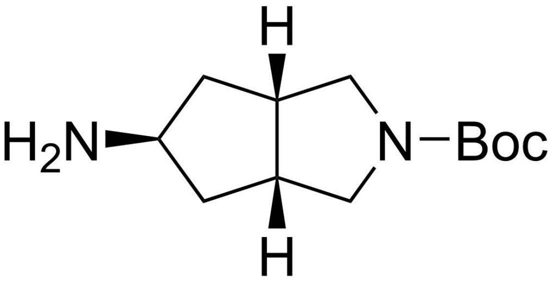
H
H₂N
N–Boc
H

**11**

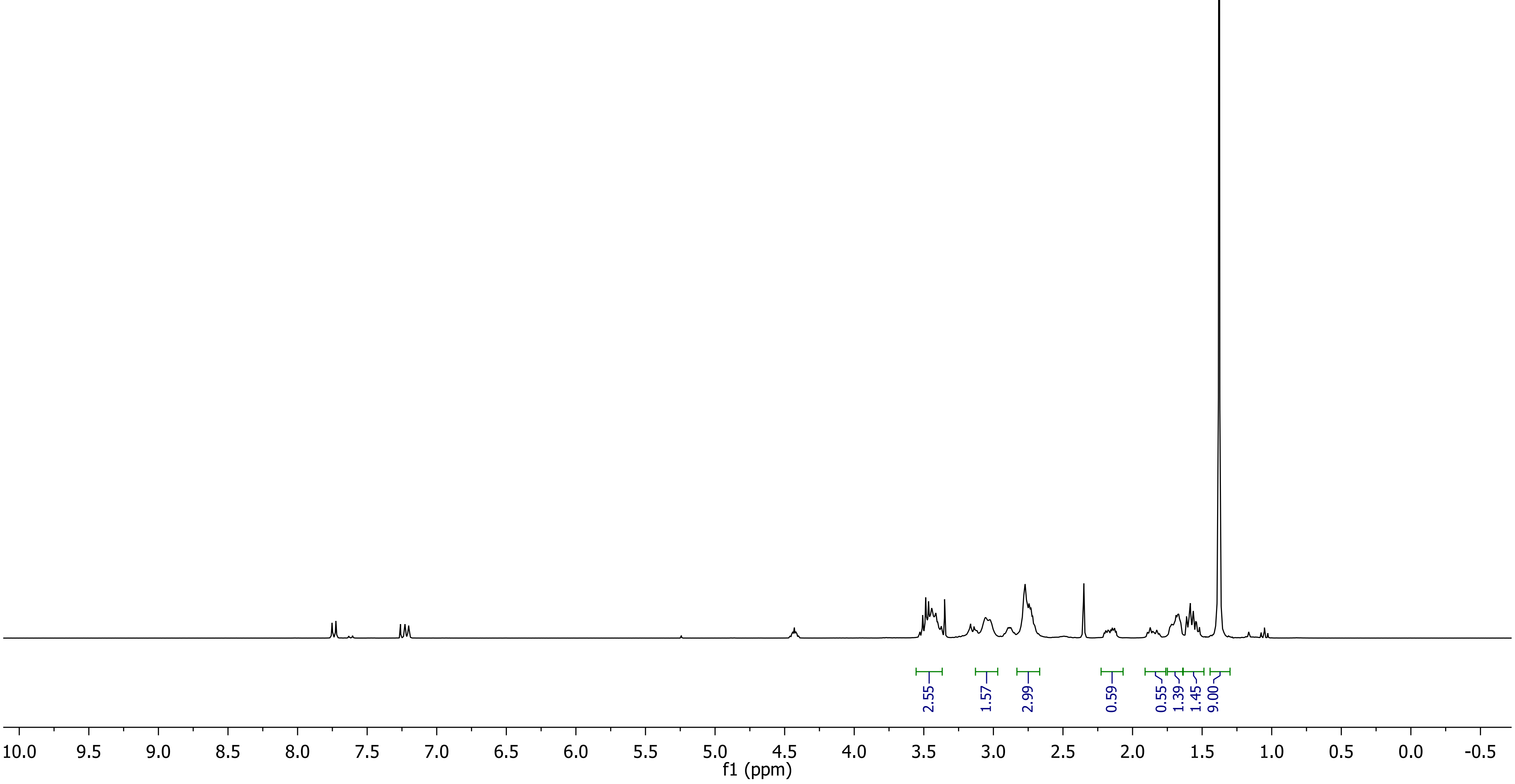
2.55
1.57
2.99
0.59
0.55
1.39
1.45
9.00
10.0 9.5 9.0 8.5 8.0 7.5 7.0 6.5 6.0 5.5 5.0 4.5 4.0 3.5 3.0 2.5 2.0 1.5 1.0 0.5 0.0 -0.5
f1 (ppm)

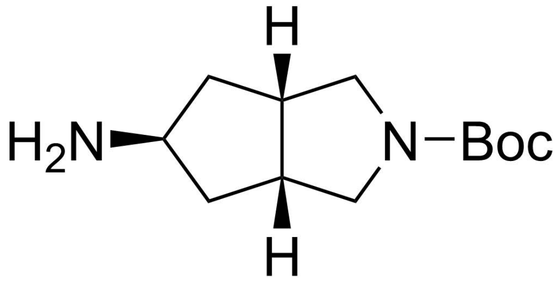
H
H
$H_2N$
N−Boc

**11**

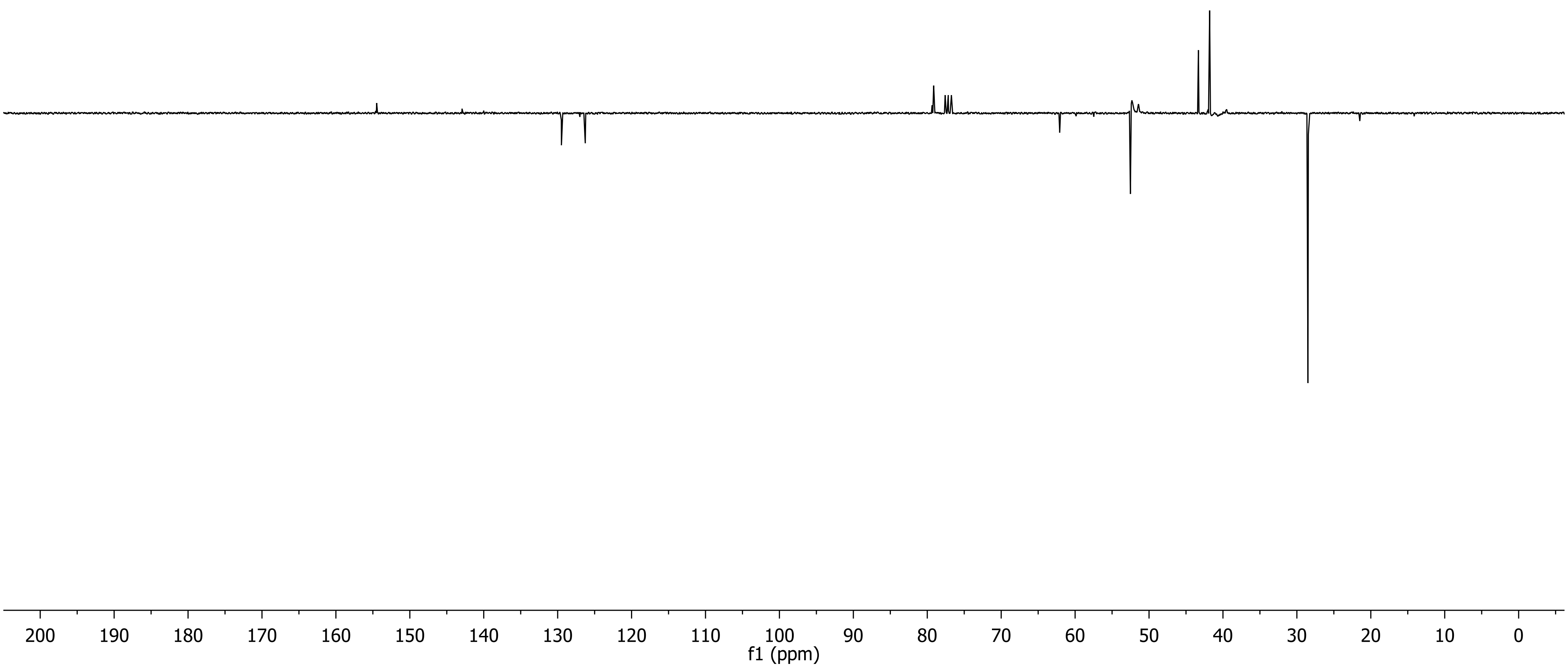
200 190 180 170 160 150 140 130 120 110 100 90 80 70 60 50 40 30 20 10 0
f1 (ppm)

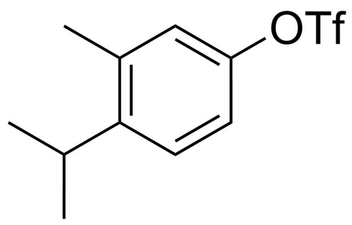
OTf

**13**

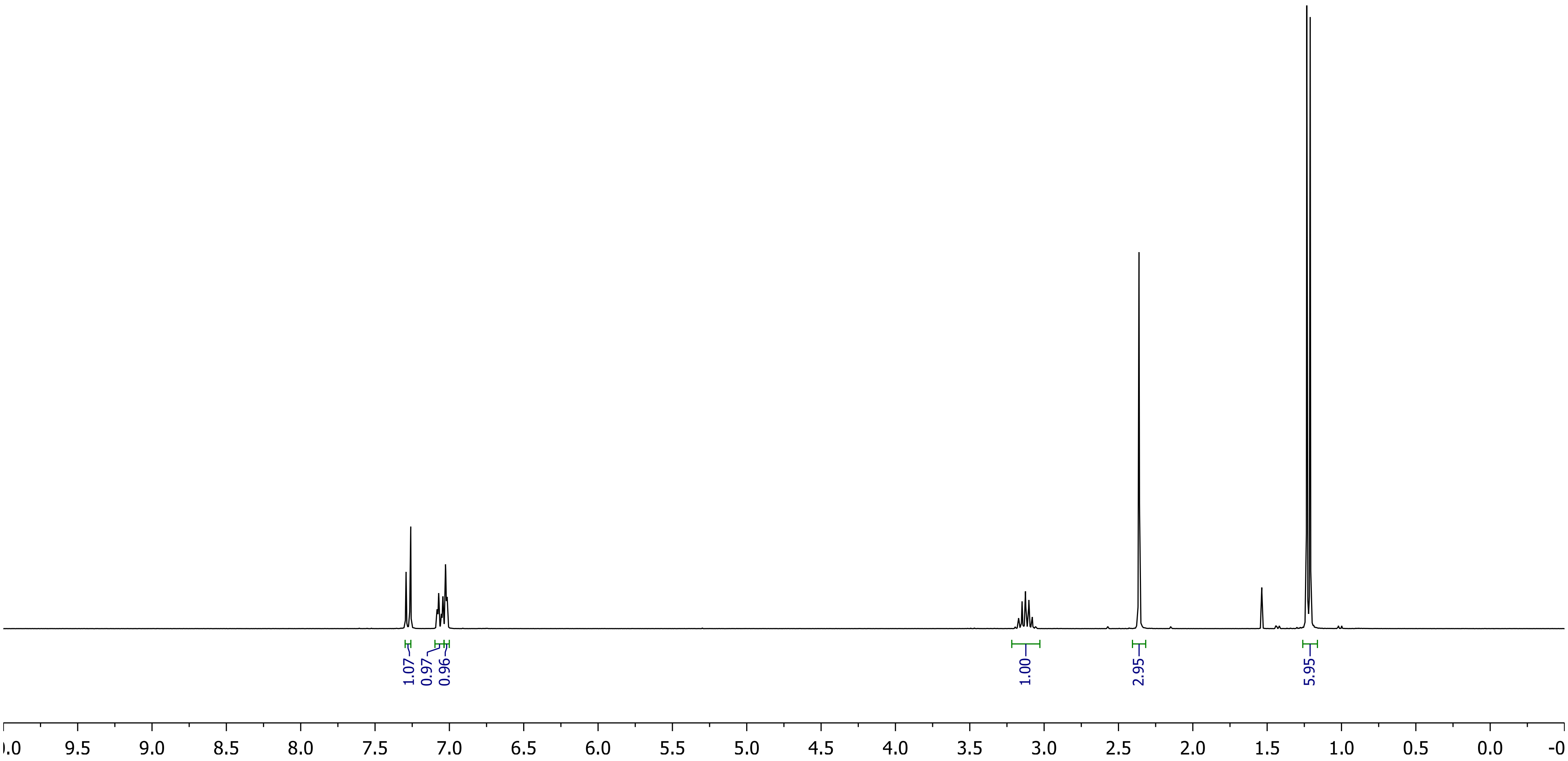
1.07
0.97
0.96
1.00
2.95
5.95
9.5 9.0 8.5 8.0 7.5 7.0 6.5 6.0 5.5 5.0 4.5 4.0 3.5 3.0 2.5 2.0 1.5 1.0 0.5 0.0

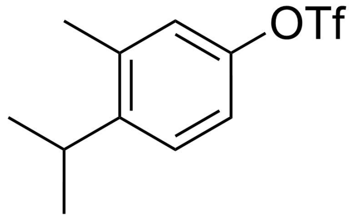
OTf

**13**

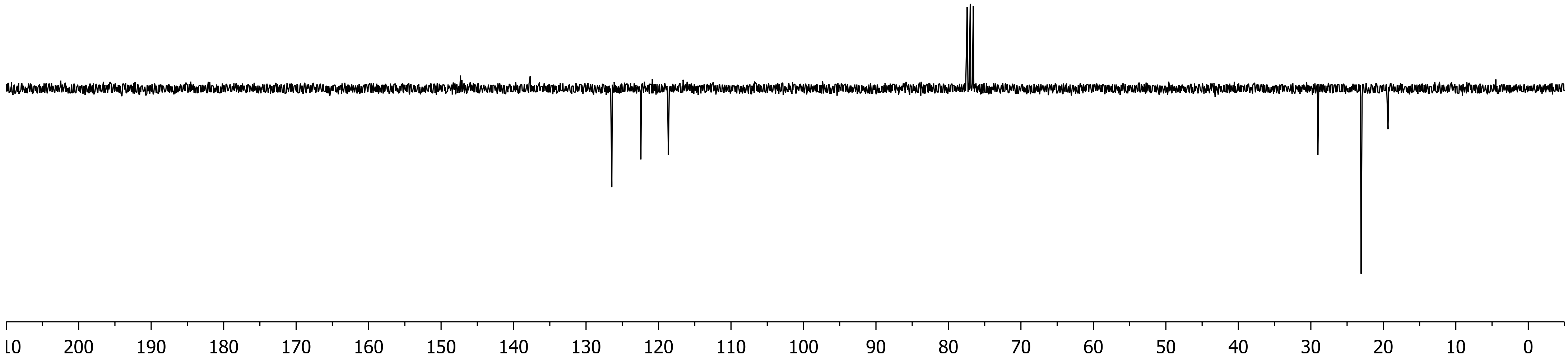
10
200
190
180
170
160
150
140
130
120
110
100
90
80
70
60
50
40
30
20
10
0

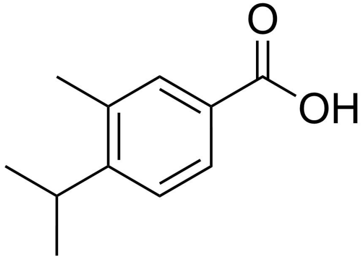
O
OH

**14**

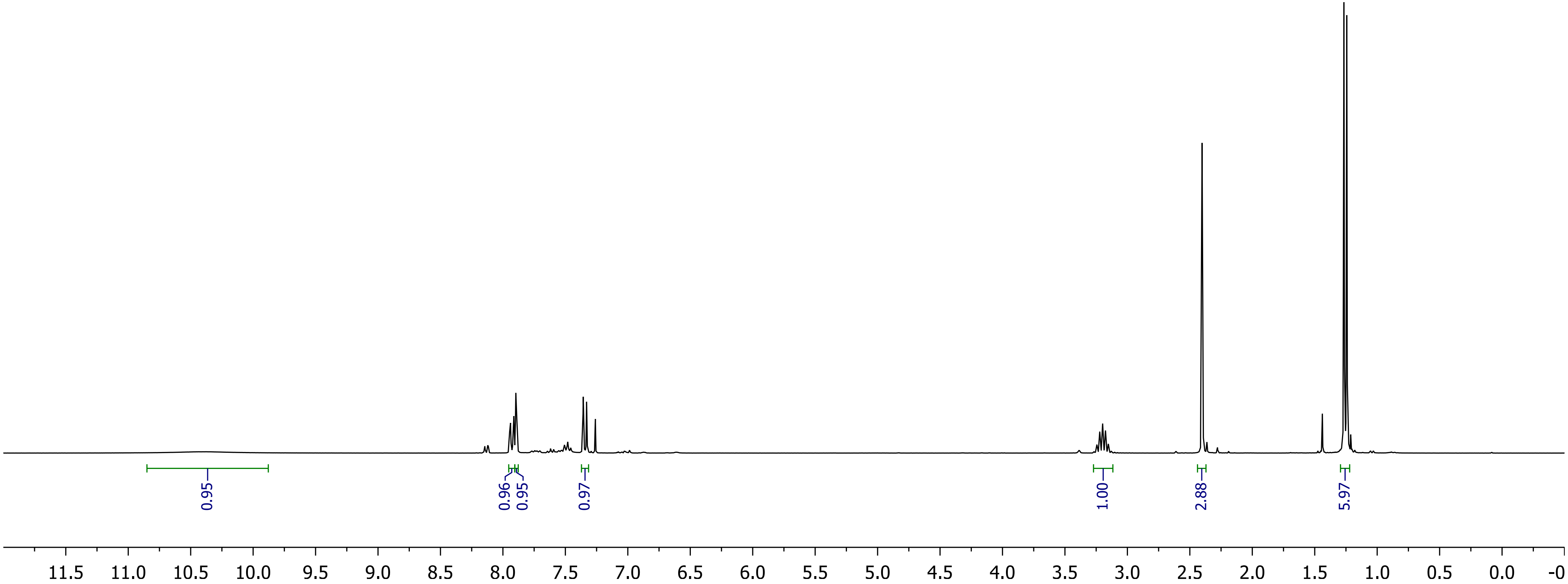
0.95
0.96
0.95
0.97
1.00
2.88
5.97
11.5 11.0 10.5 10.0 9.5 9.0 8.5 8.0 7.5 7.0 6.5 6.0 5.5 5.0 4.5 4.0 3.5 3.0 2.5 2.0 1.5 1.0 0.5 0.0 -0

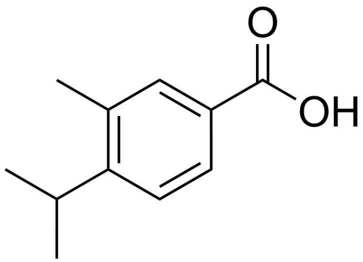

**14**

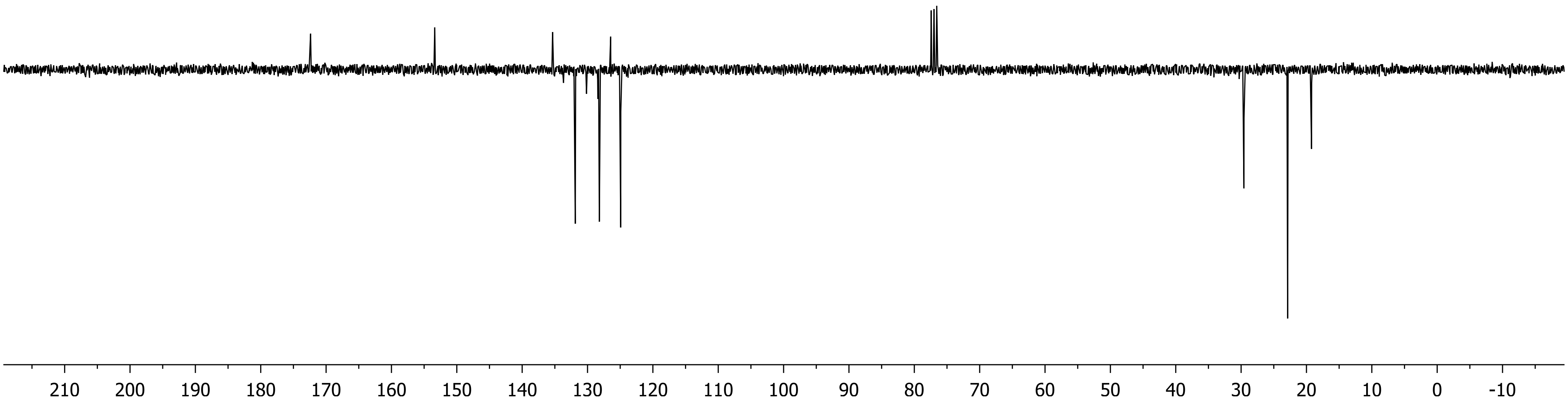

210
200
190
180
170
160
150
140
130
120
110
100
90
80
70
60
50
40
30
20
10
0
-10

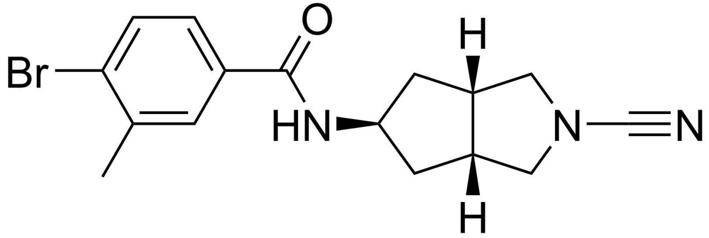

**BB07CA28**

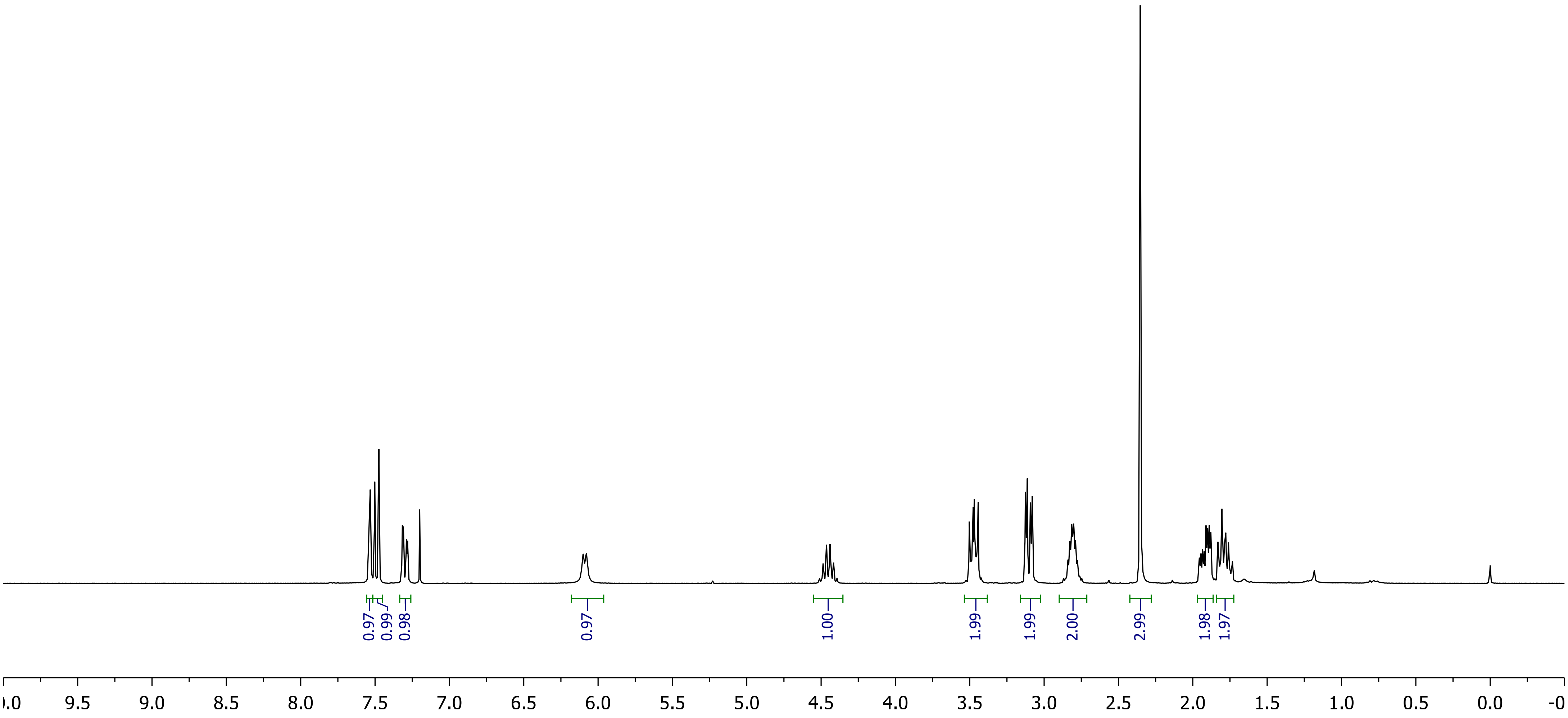
0.97
0.99
0.98
0.97
1.00
1.99
1.99
2.00
2.99
1.98
1.97
9.5
9.0
8.5
8.0
7.5
7.0
6.5
6.0
5.5
5.0
4.5
4.0
3.5
3.0
2.5
2.0
1.5
1.0
0.5
0.0

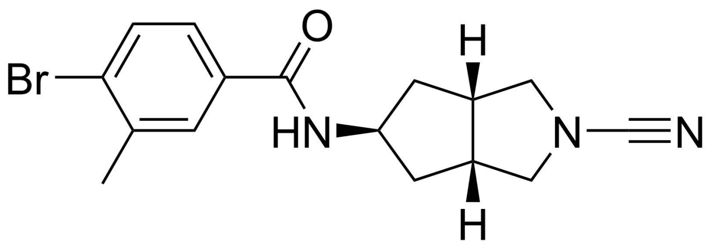

**BB07CA28**

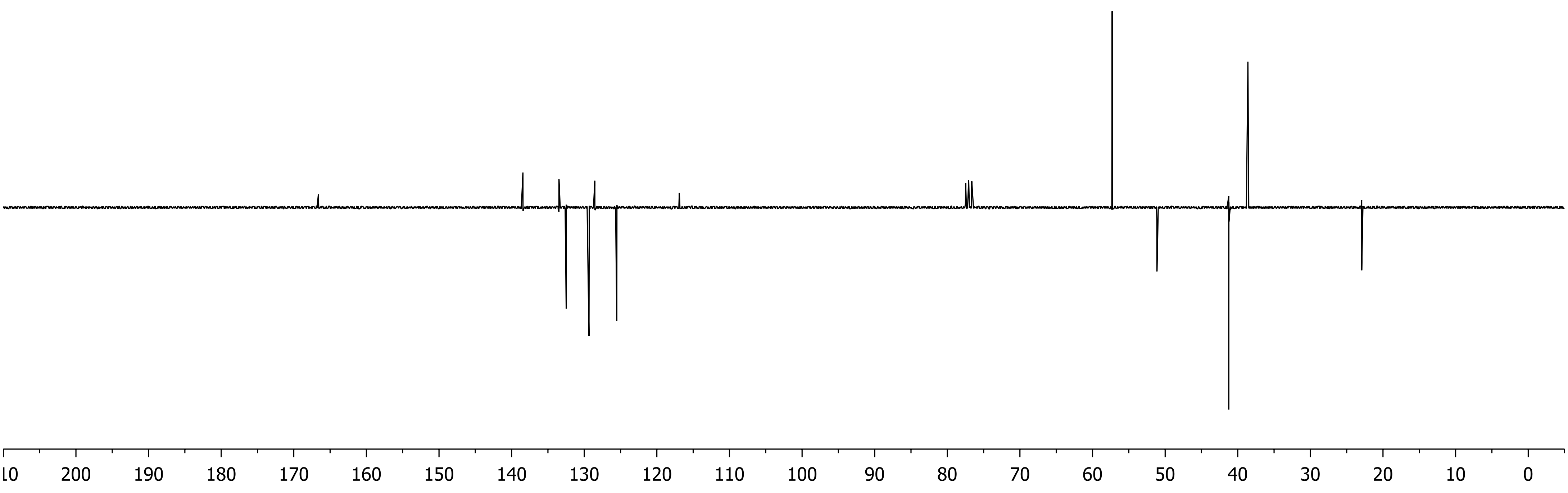

10
200
190
180
170
160
150
140
130
120
110
100
90
80
70
60
50
40
30
20
10
0

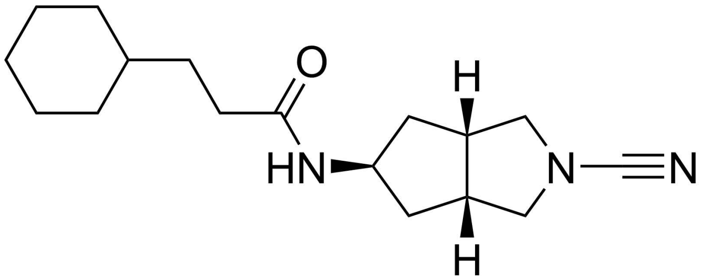

**BB07CA48**

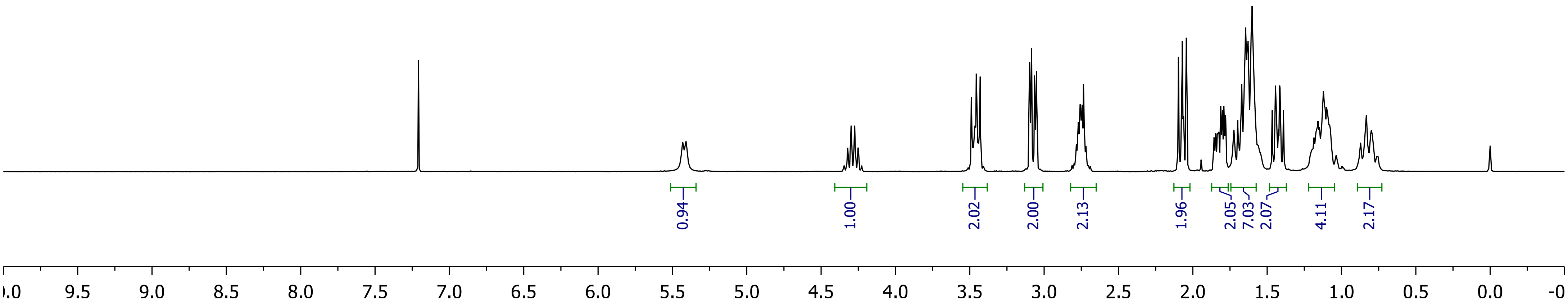
0.94
1.00
2.02
2.00
2.13
1.96
2.05
7.03
2.07
4.11
2.17
10.0 9.5 9.0 8.5 8.0 7.5 7.0 6.5 6.0 5.5 5.0 4.5 4.0 3.5 3.0 2.5 2.0 1.5 1.0 0.5 0.0 -0

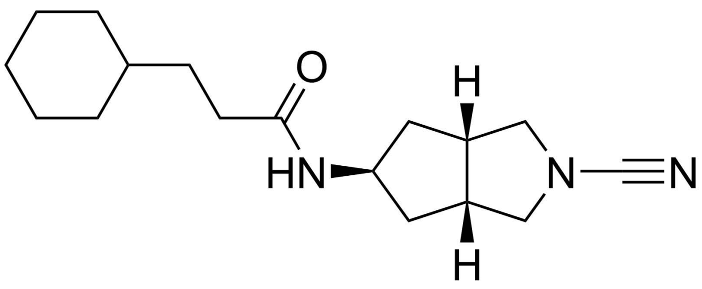
O
H
HN
N
N
H

**BB07CA48**

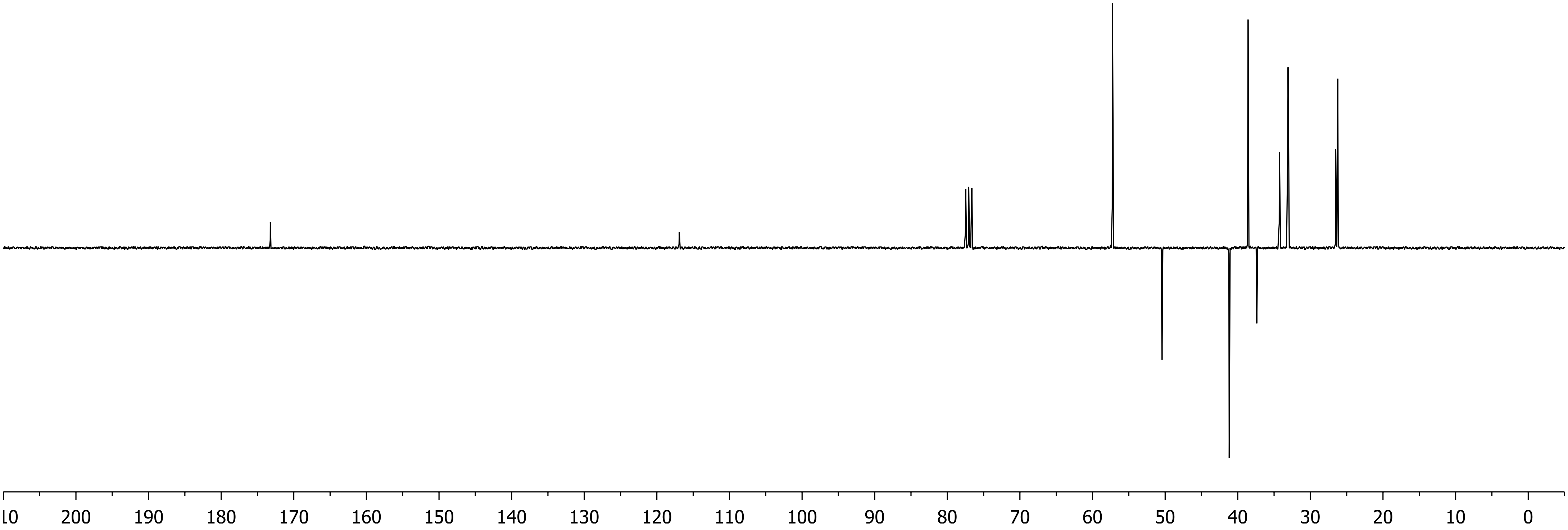
200 190 180 170 160 150 140 130 120 110 100 90 80 70 60 50 40 30 20 10 0

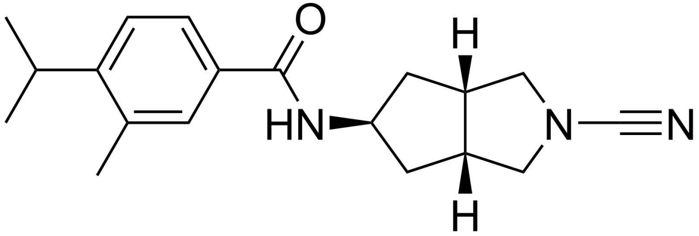

**BB07CA902**

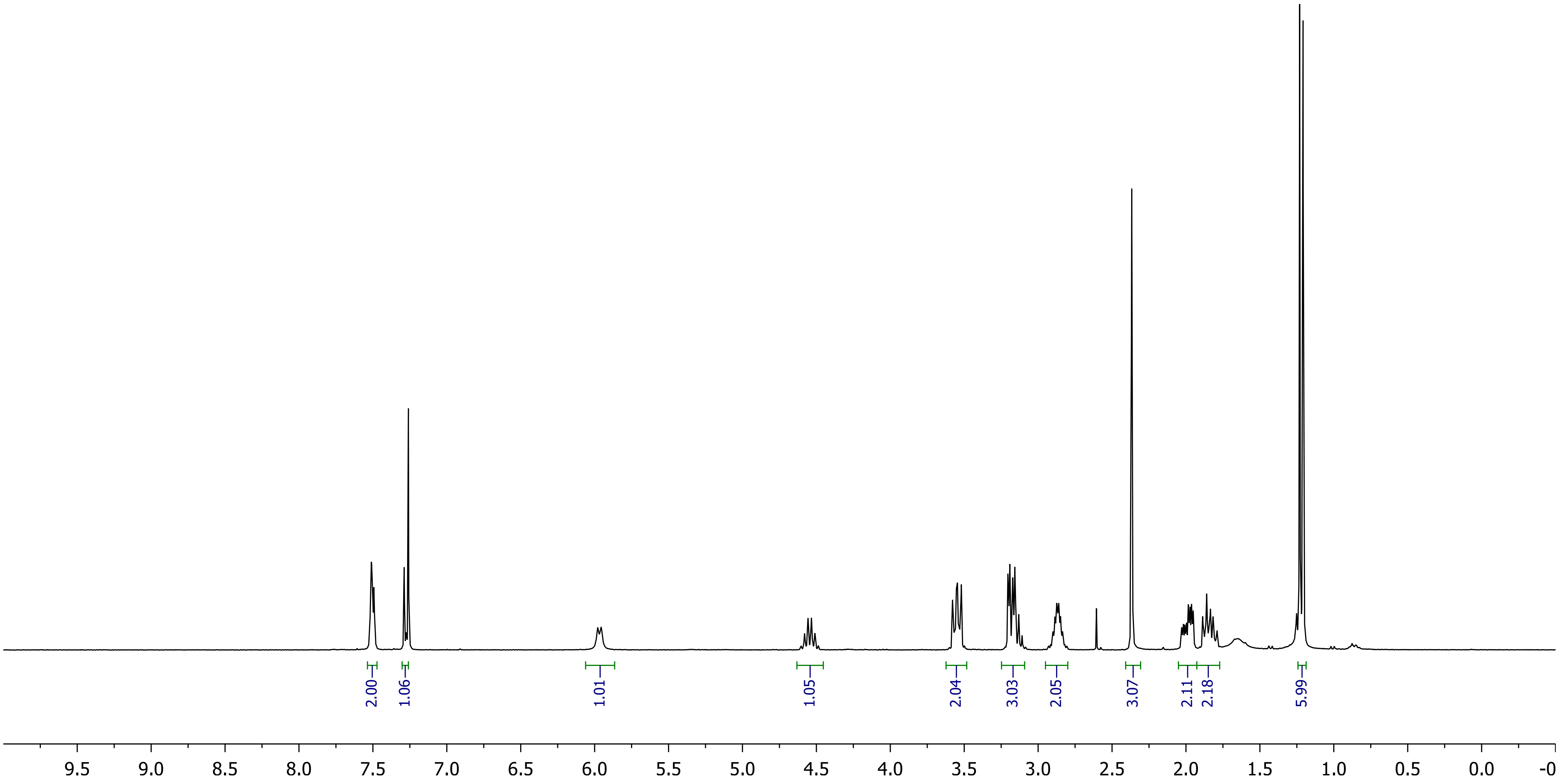
2.00
1.06
1.01
1.05
2.04
3.03
2.05
3.07
2.11
2.18
5.99
9.5 9.0 8.5 8.0 7.5 7.0 6.5 6.0 5.5 5.0 4.5 4.0 3.5 3.0 2.5 2.0 1.5 1.0 0.5 0.0 -0

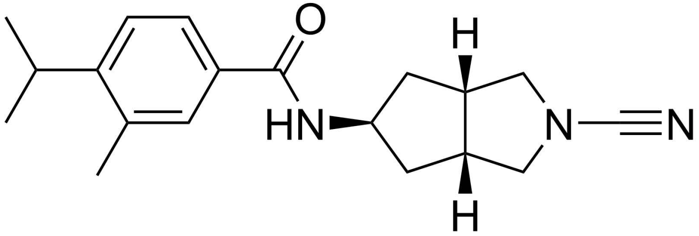

O
H
HN
N
N
H


**BB07CA902**

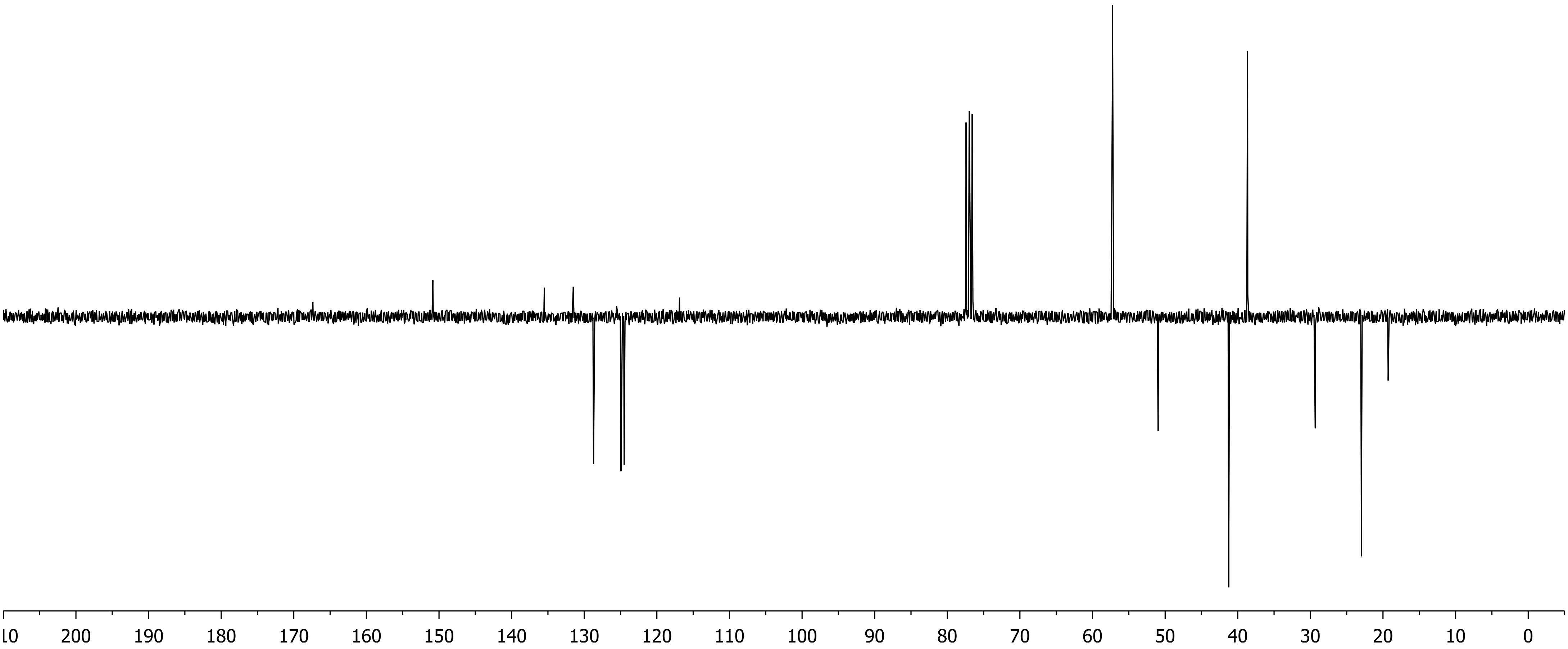

10
200
190
180
170
160
150
140
130
120
110
100
90
80
70
60
50
40
30
20
10
0

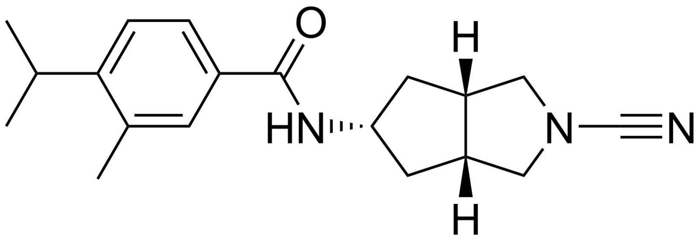

**BB06CA902**

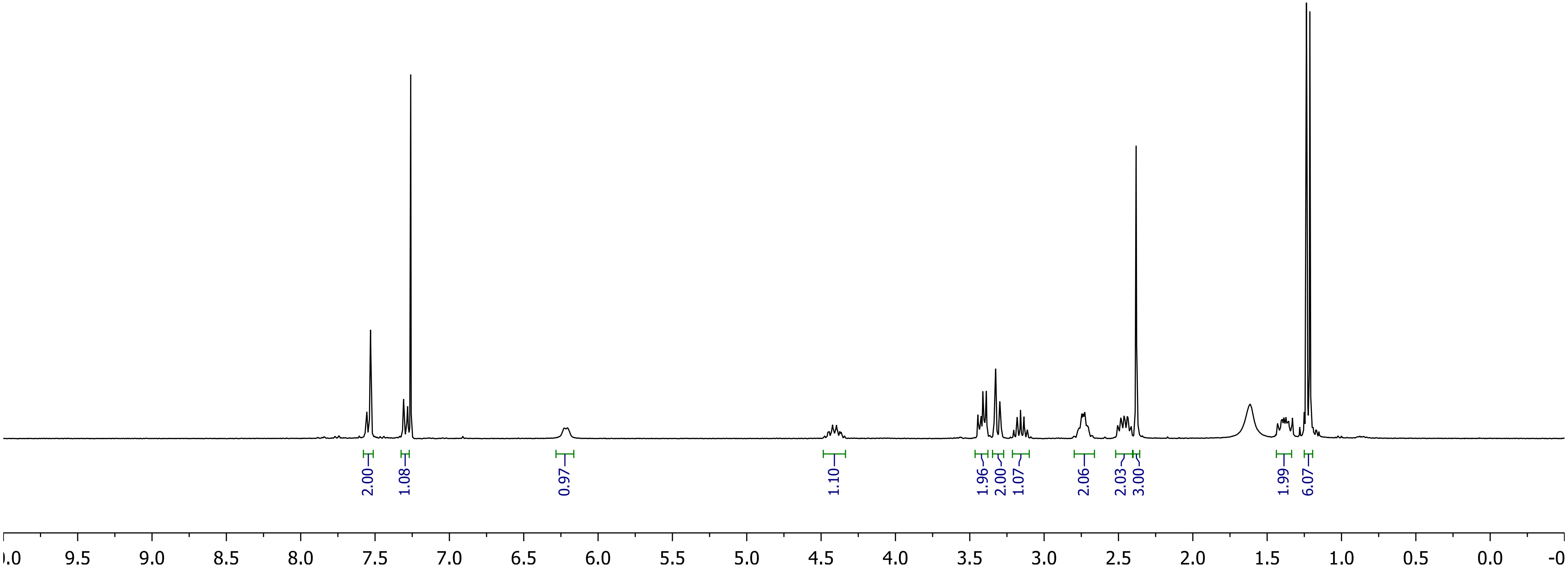

2.00
1.08
0.97
1.10
1.96
2.00
1.07
2.06
2.03
3.00
1.99
6.07
9.5
9.0
8.5
8.0
7.5
7.0
6.5
6.0
5.5
5.0
4.5
4.0
3.5
3.0
2.5
2.0
1.5
1.0
0.5
0.0

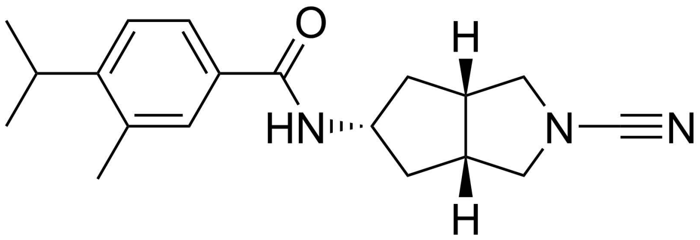
O
H
HN
N
N
H

**BB06CA902**

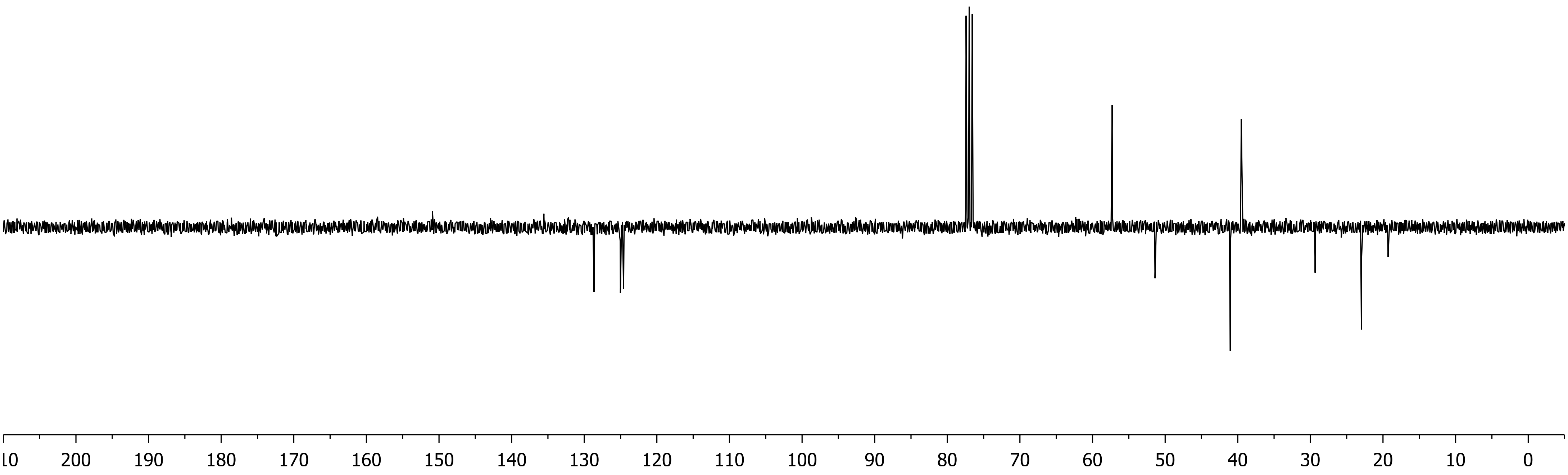
200
190
180
170
160
150
140
130
120
110
100
90
80
70
60
50
40
30
20
10
0

#### ISG15$_{CT}$-Rho-MP

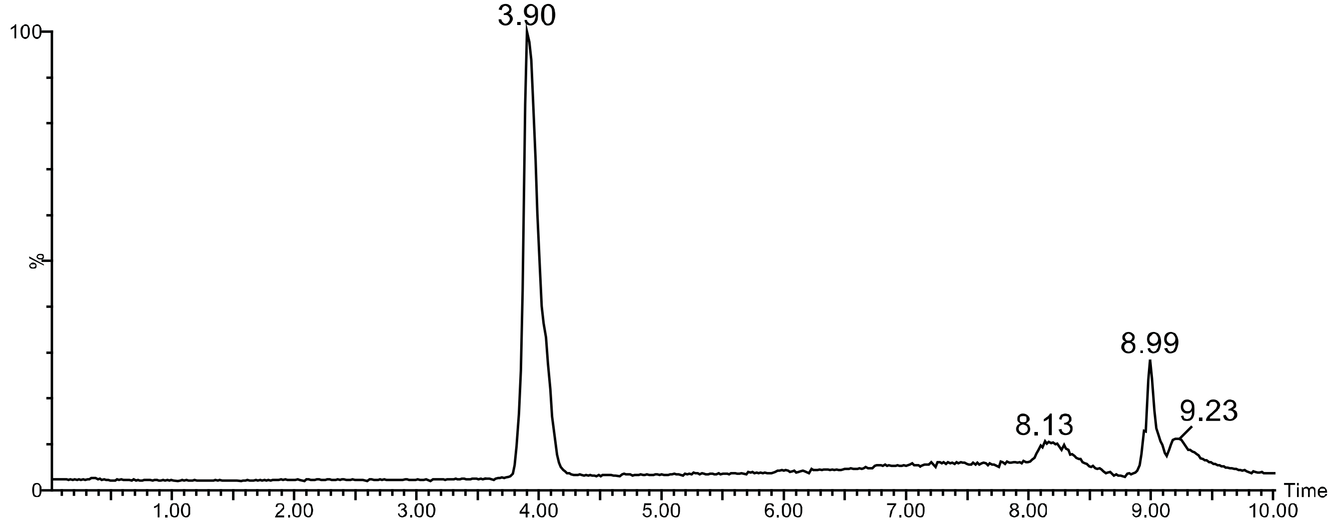


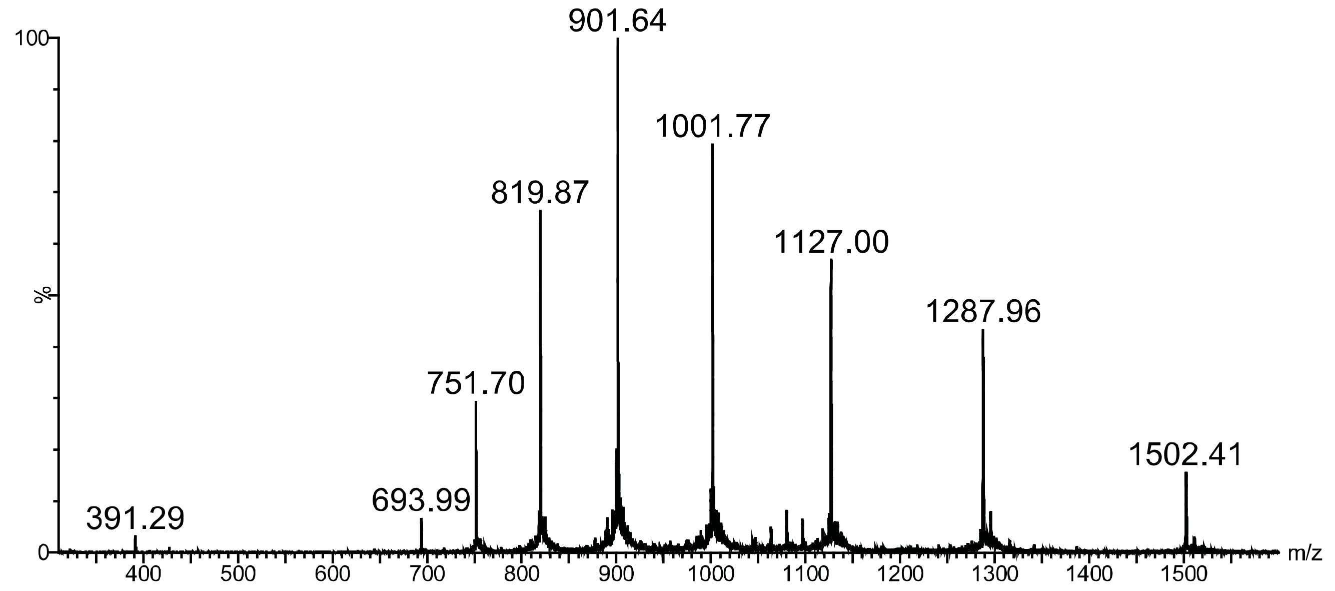


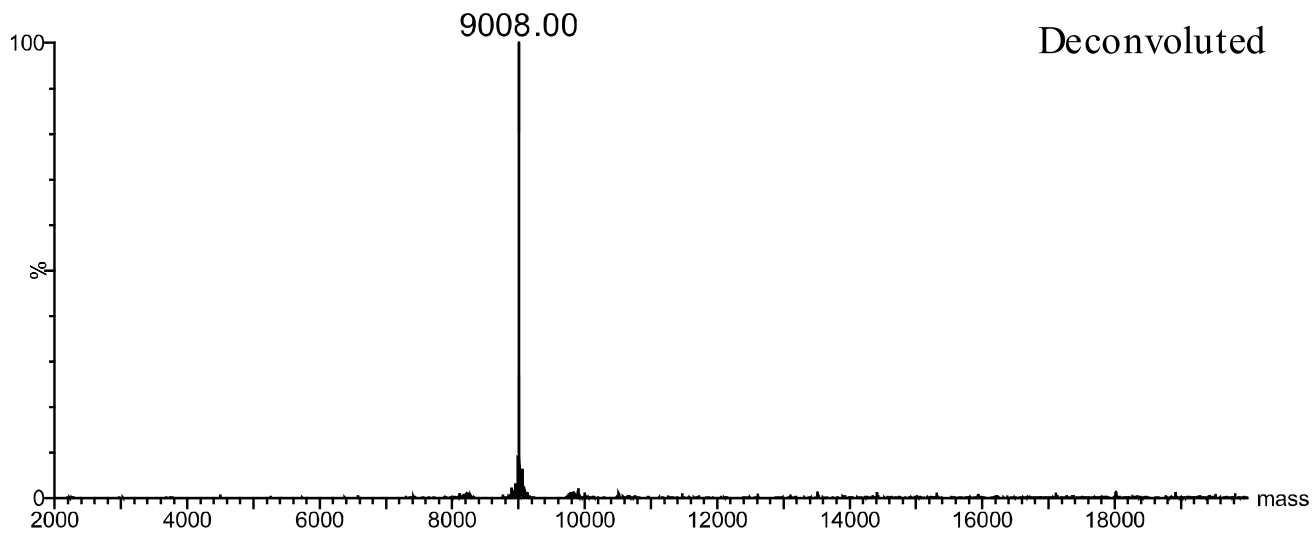

#### Compound 1

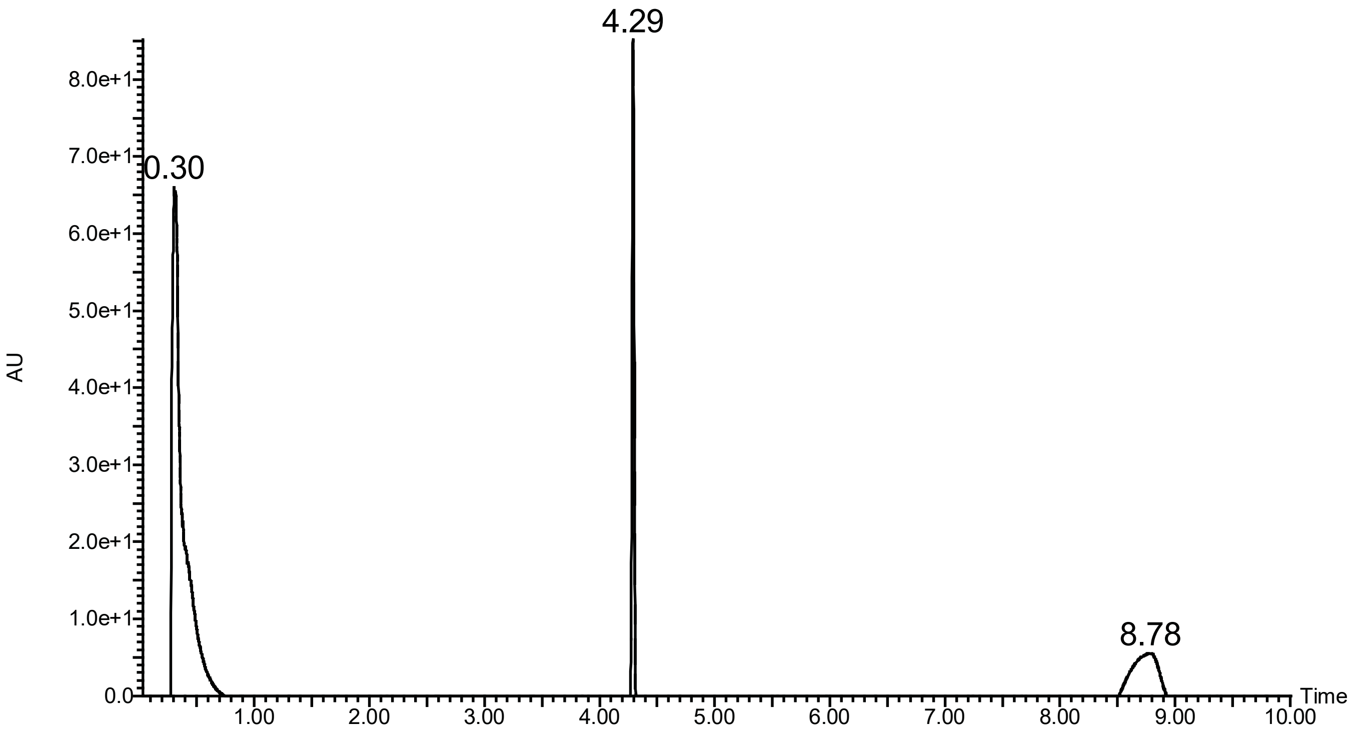

4.29
0.30
8.78
AU
8.0e+1
7.0e+1
6.0e+1
5.0e+1
4.0e+1
3.0e+1
2.0e+1
1.0e+1
0.0
1.00
2.00
3.00
4.00
5.00
6.00
7.00
8.00
9.00
10.00
Time


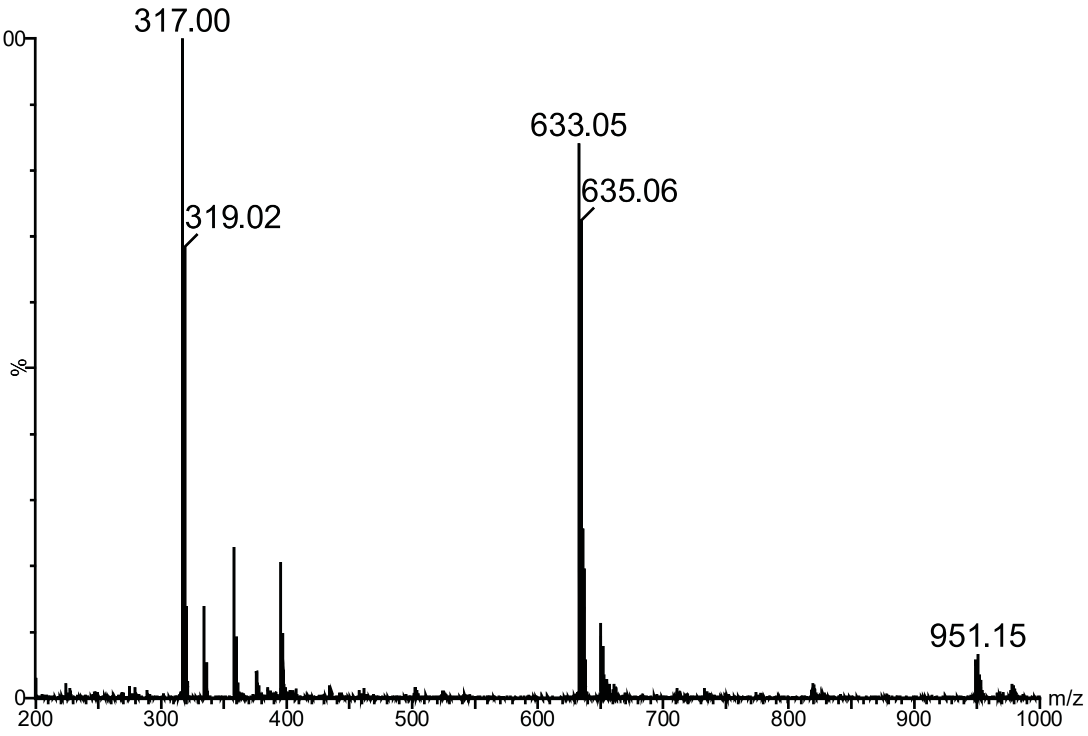

317.00
319.02
633.05
635.06
951.15
100
%
0
200
300
400
500
600
700
800
900
1000
m/z

### Compound 2

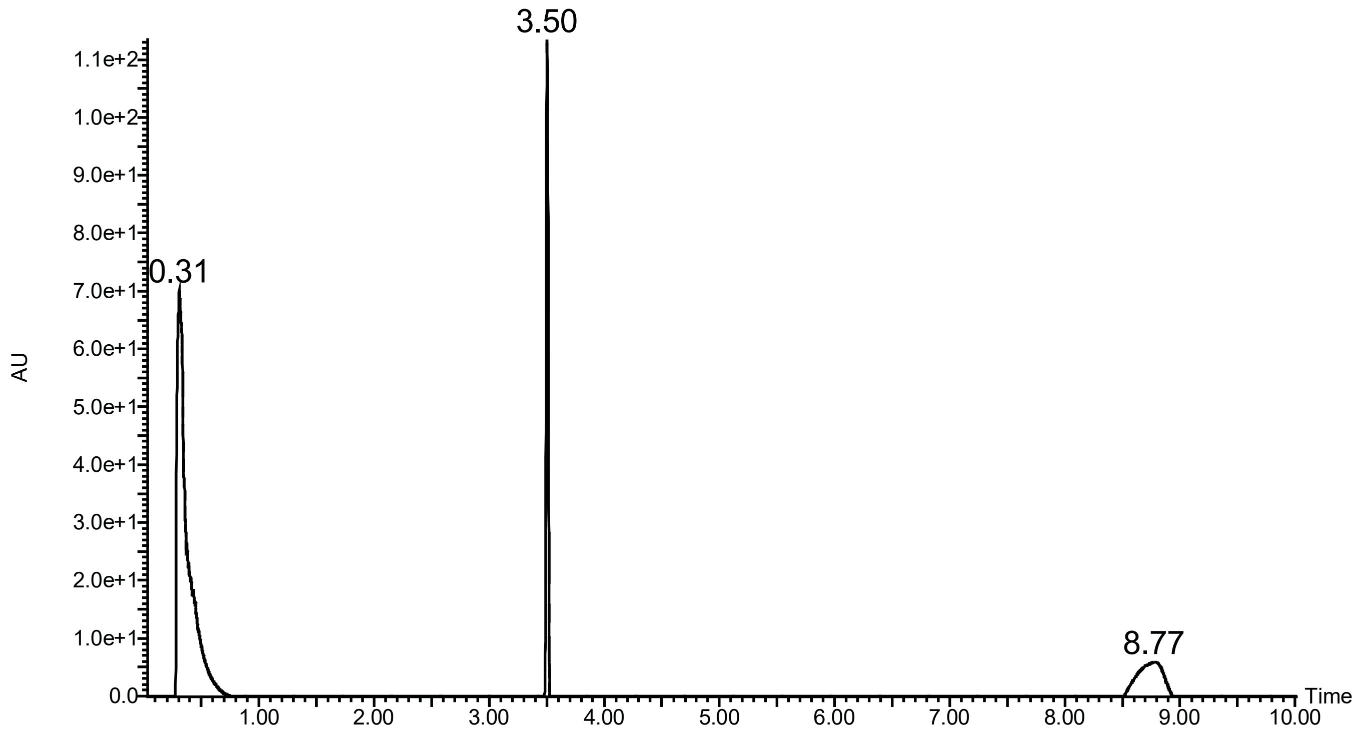

3.50
0.31
8.77
AU
1.1e+2
1.0e+2
9.0e+1
8.0e+1
7.0e+1
6.0e+1
5.0e+1
4.0e+1
3.0e+1
2.0e+1
1.0e+1
0.0
1.00
2.00
3.00
4.00
5.00
6.00
7.00
8.00
9.00
10.00
Time


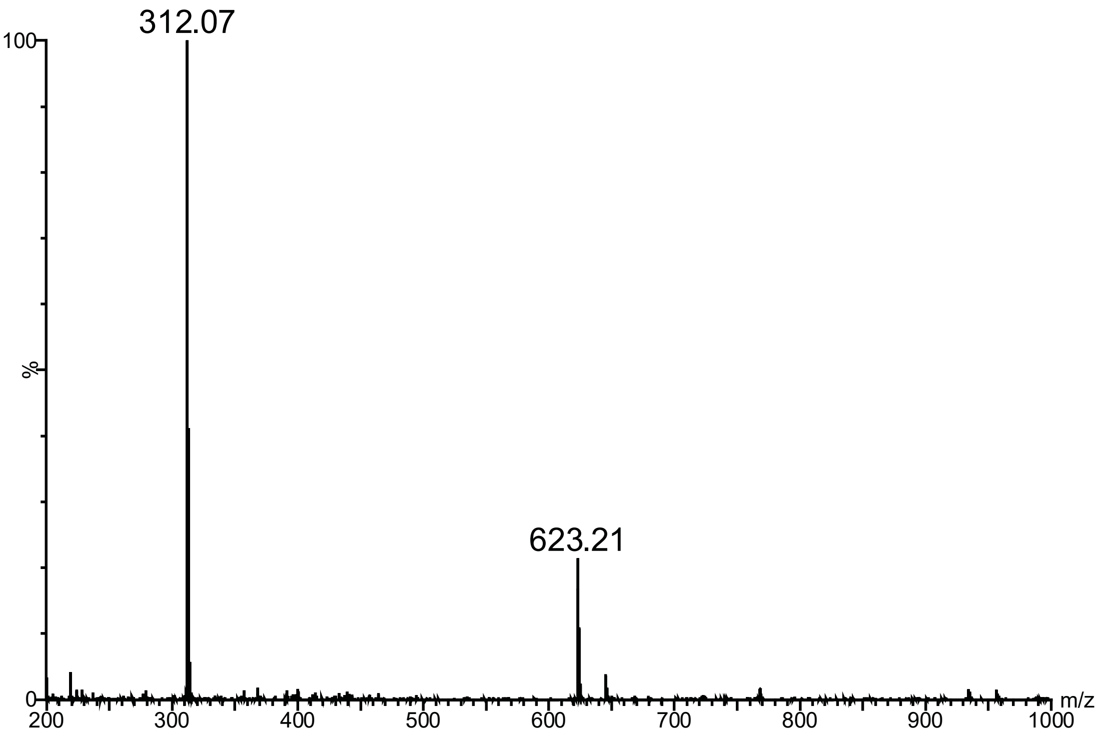

312.07
623.21
100
%
0
200
300
400
500
600
700
800
900
1000
m/z

### Compound 3

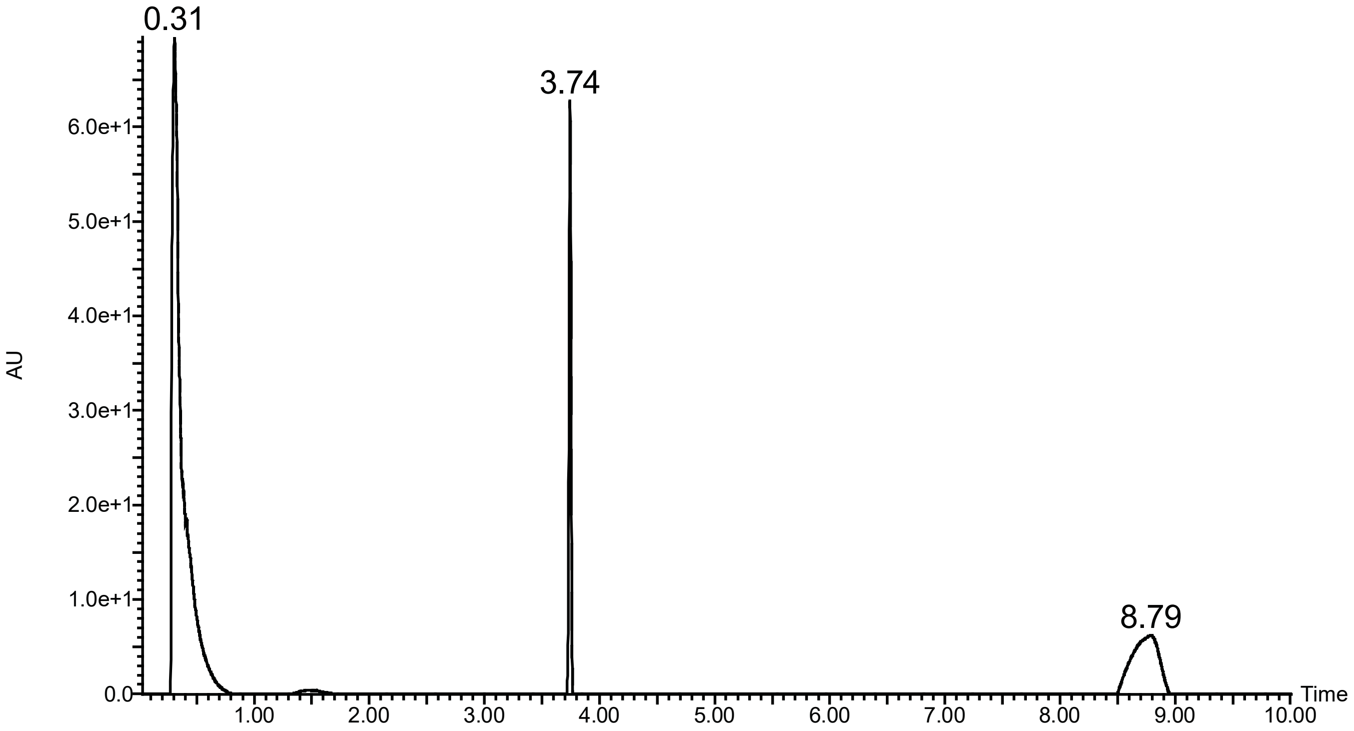
0.31
3.74
8.79
6.0e+1
5.0e+1
4.0e+1
3.0e+1
2.0e+1
1.0e+1
0.0
AU
1.00
2.00
3.00
4.00
5.00
6.00
7.00
8.00
9.00
10.00
Time

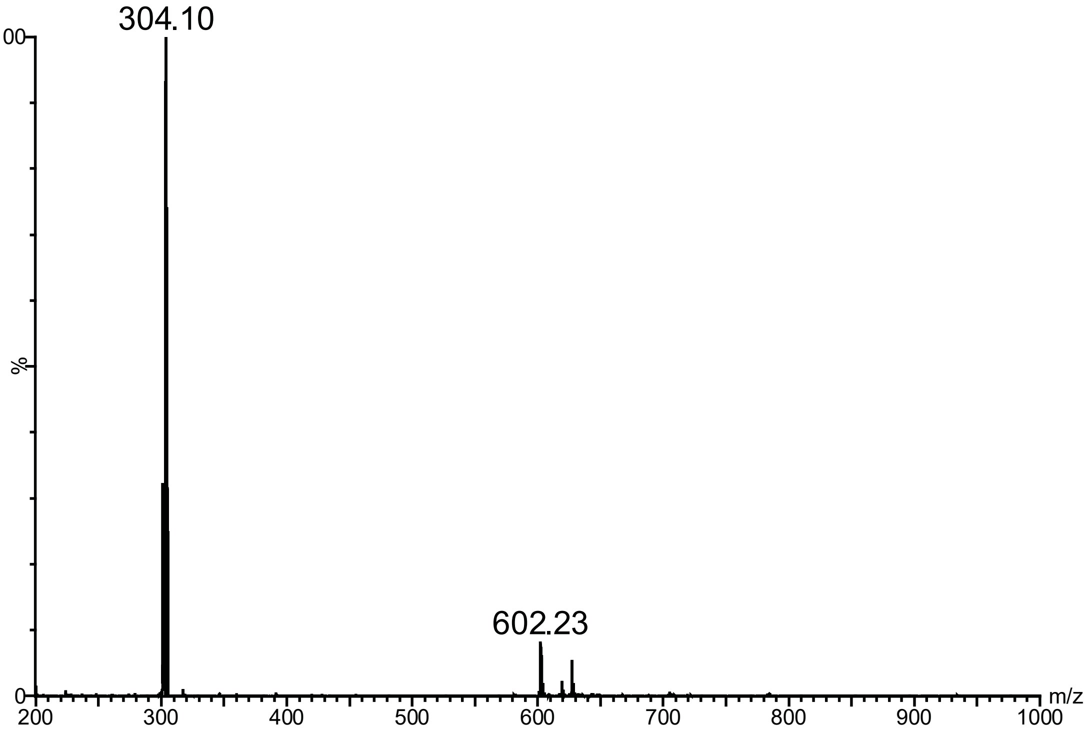
304.10
602.23
100
%
0
200
300
400
500
600
700
800
900
1000
m/z

### Compound 4

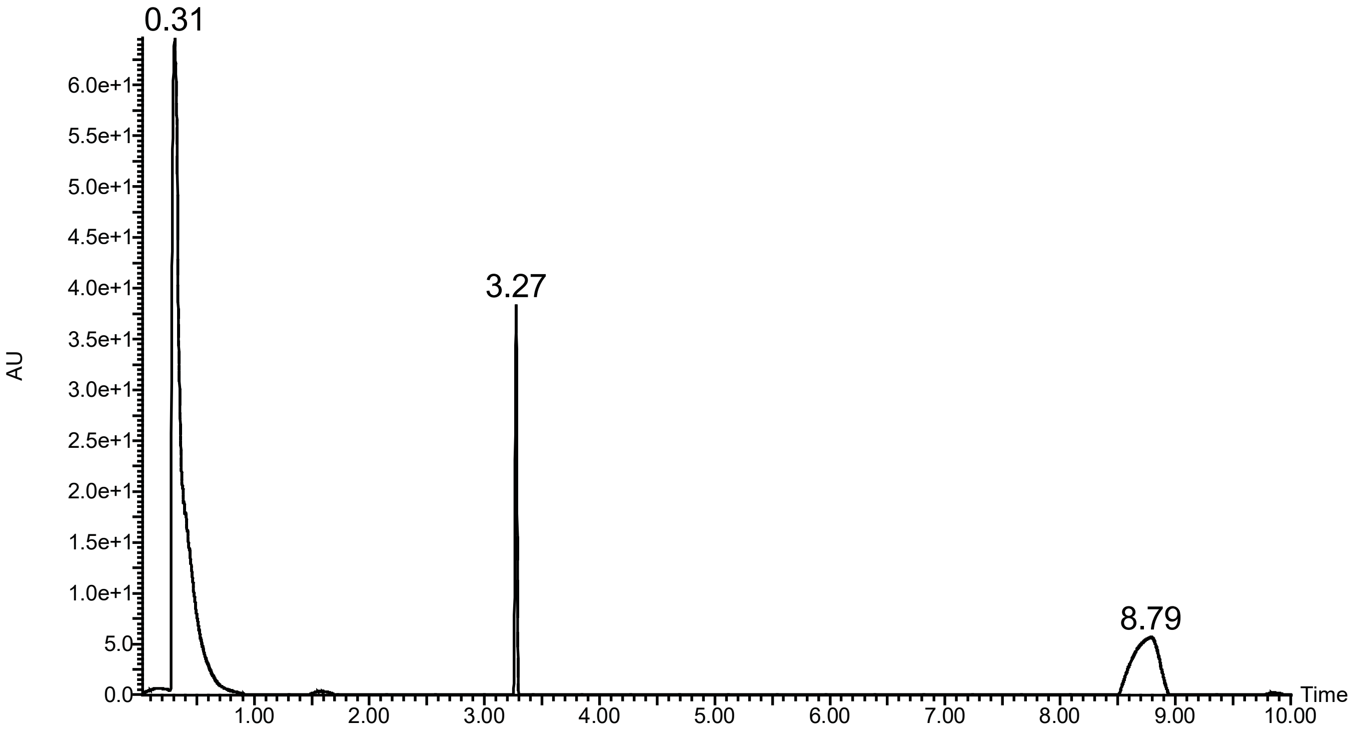

0.31
3.27
8.79
AU
6.0e+1
5.5e+1
5.0e+1
4.5e+1
4.0e+1
3.5e+1
3.0e+1
2.5e+1
2.0e+1
1.5e+1
1.0e+1
5.0
0.0
1.00
2.00
3.00
4.00
5.00
6.00
7.00
8.00
9.00
10.00
Time


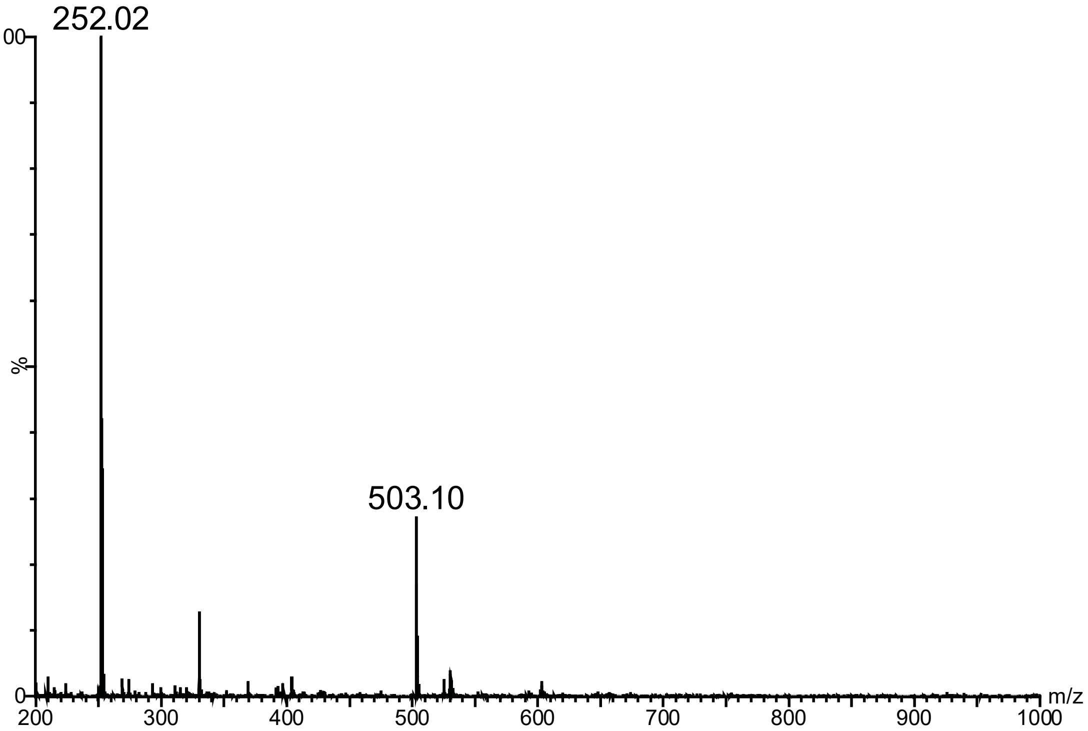

252.02
503.10
100
%
0
200
300
400
500
600
700
800
900
1000
m/z

### Compound 5

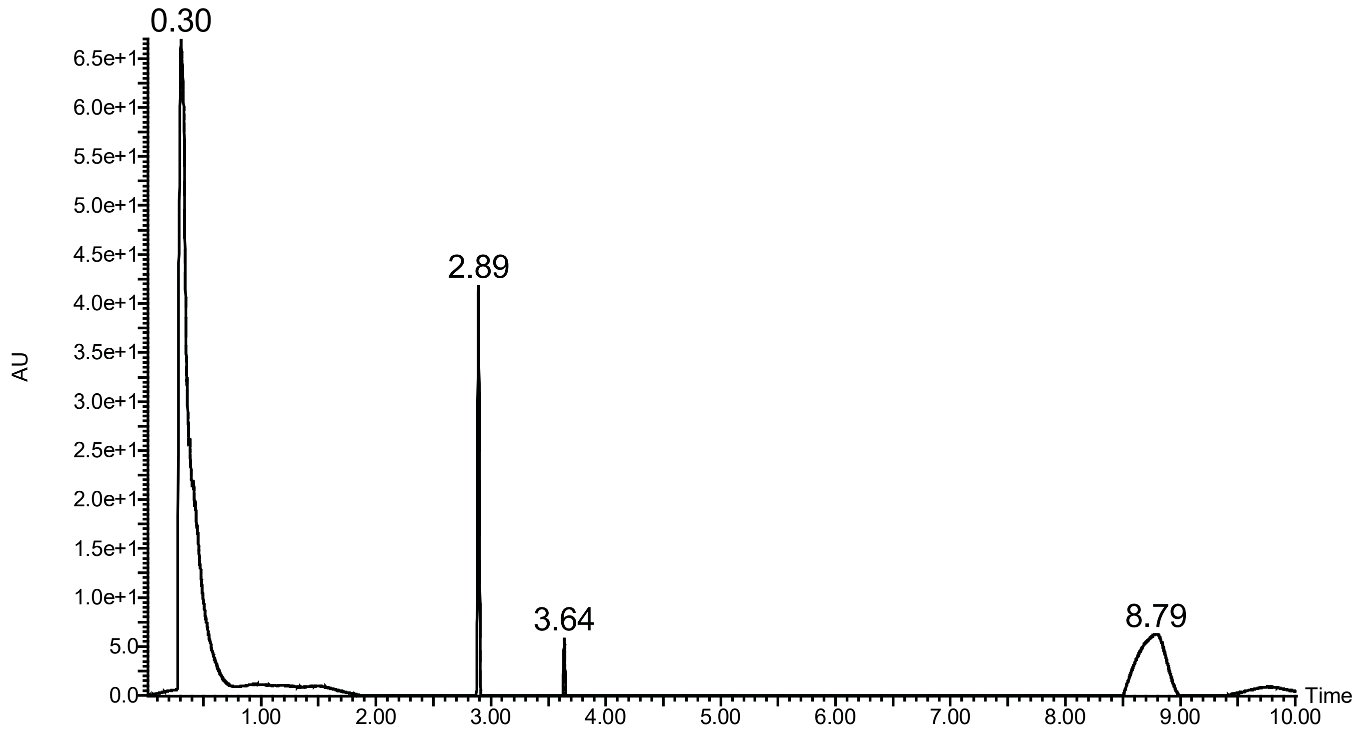

0.30
2.89
3.64
8.79
AU
Time
6.5e+1
6.0e+1
5.5e+1
5.0e+1
4.5e+1
4.0e+1
3.5e+1
3.0e+1
2.5e+1
2.0e+1
1.5e+1
1.0e+1
5.0
0.0
1.00
2.00
3.00
4.00
5.00
6.00
7.00
8.00
9.00
10.00


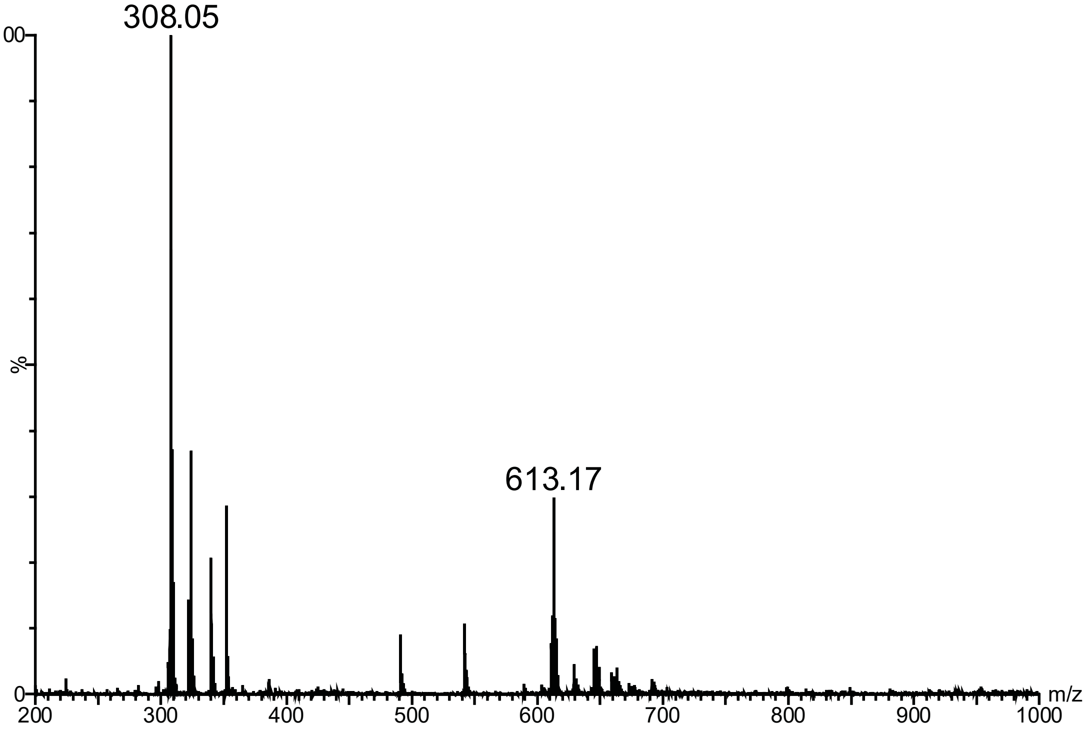

308.05
613.17
100
%
0
m/z
200
300
400
500
600
700
800
900
1000

**BB07CA28**

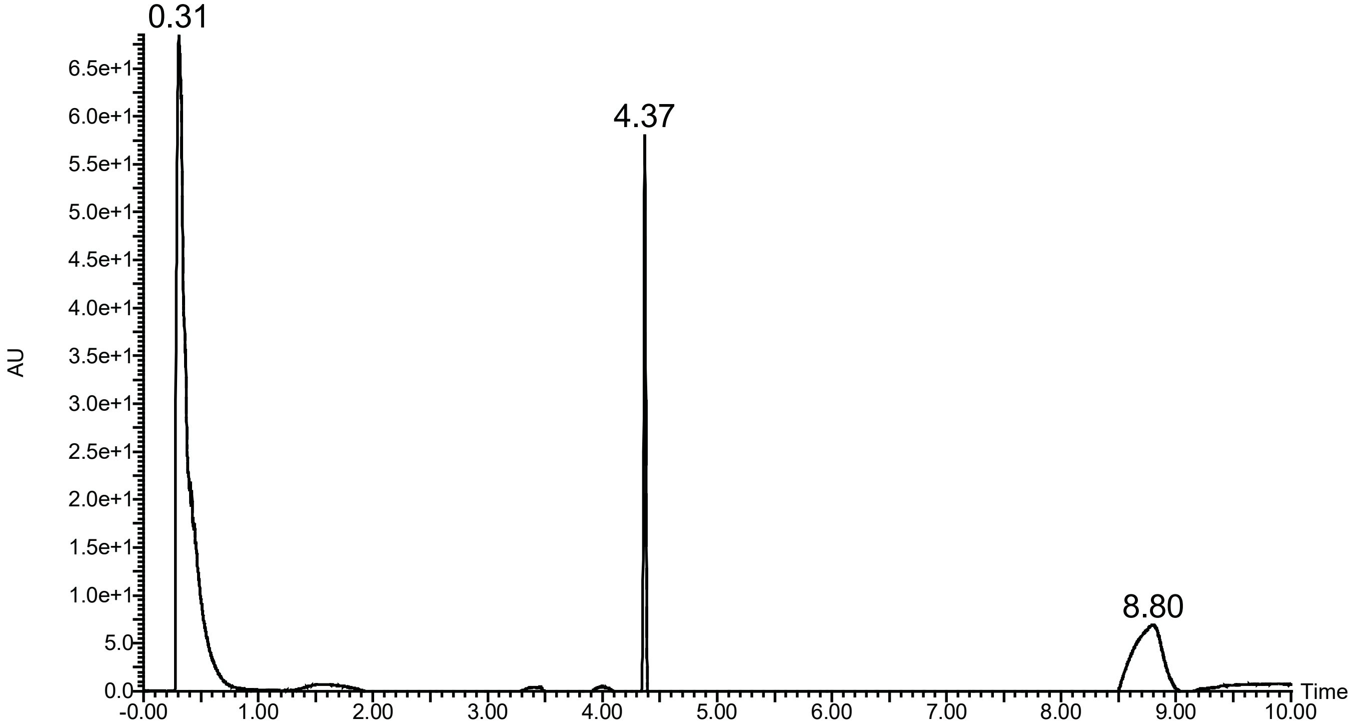
0.31
4.37
8.80
AU
6.5e+1
6.0e+1
5.5e+1
5.0e+1
4.5e+1
4.0e+1
3.5e+1
3.0e+1
2.5e+1
2.0e+1
1.5e+1
1.0e+1
5.0
0.0
Time
-0.00
1.00
2.00
3.00
4.00
5.00
6.00
7.00
8.00
9.00
10.00

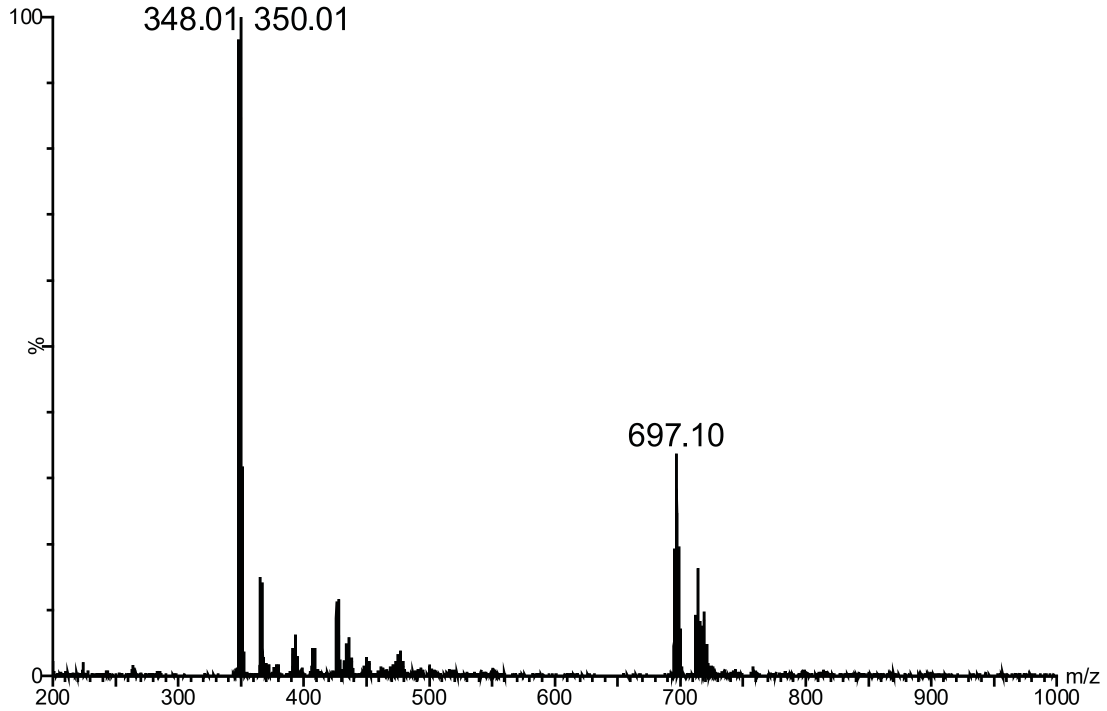
348.01
350.01
697.10
100
%
0
m/z
200
300
400
500
600
700
800
900
1000

**BB07CA48**

(not UV active)

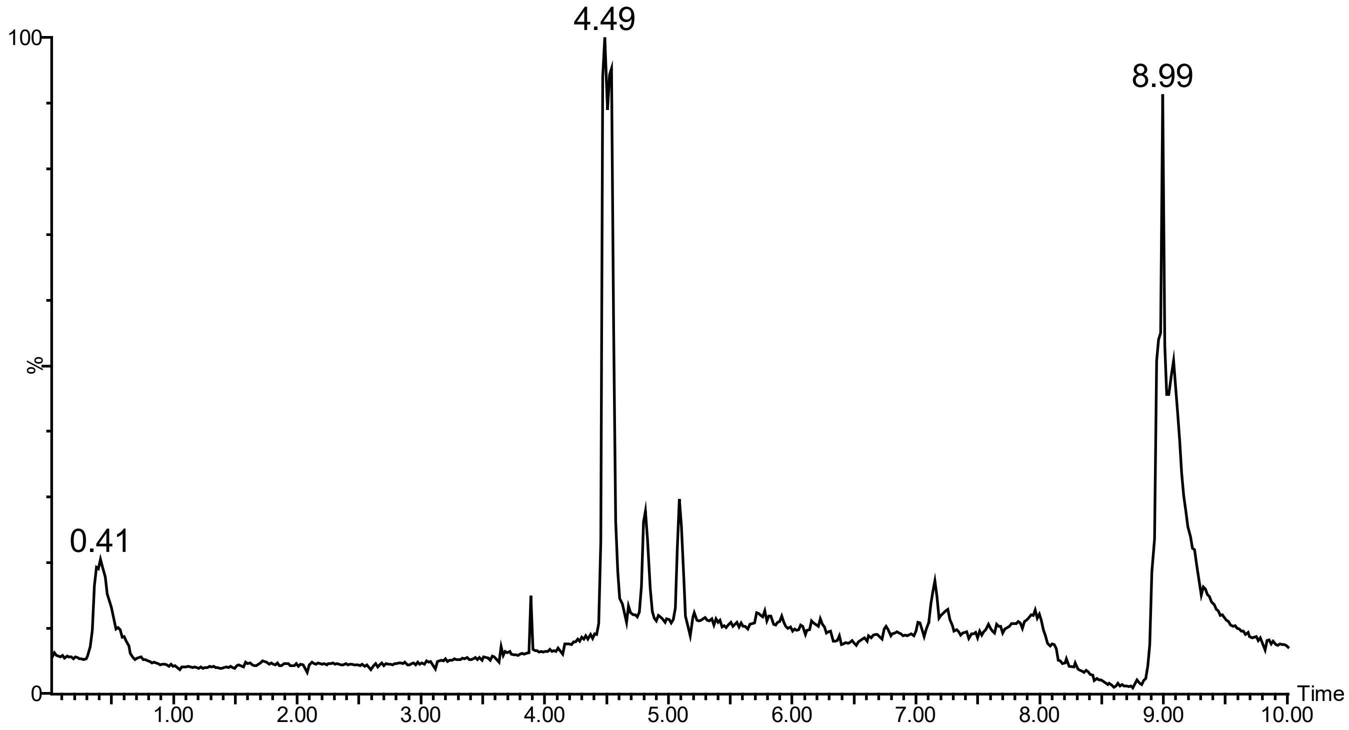


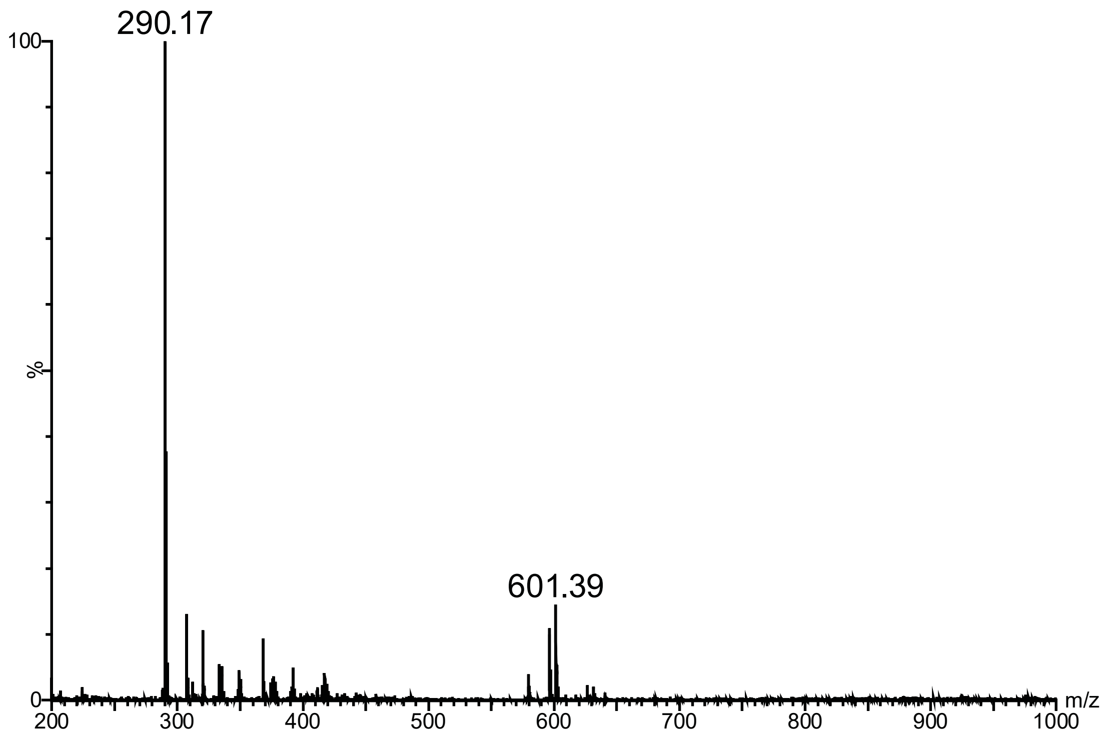

# BB06CA902

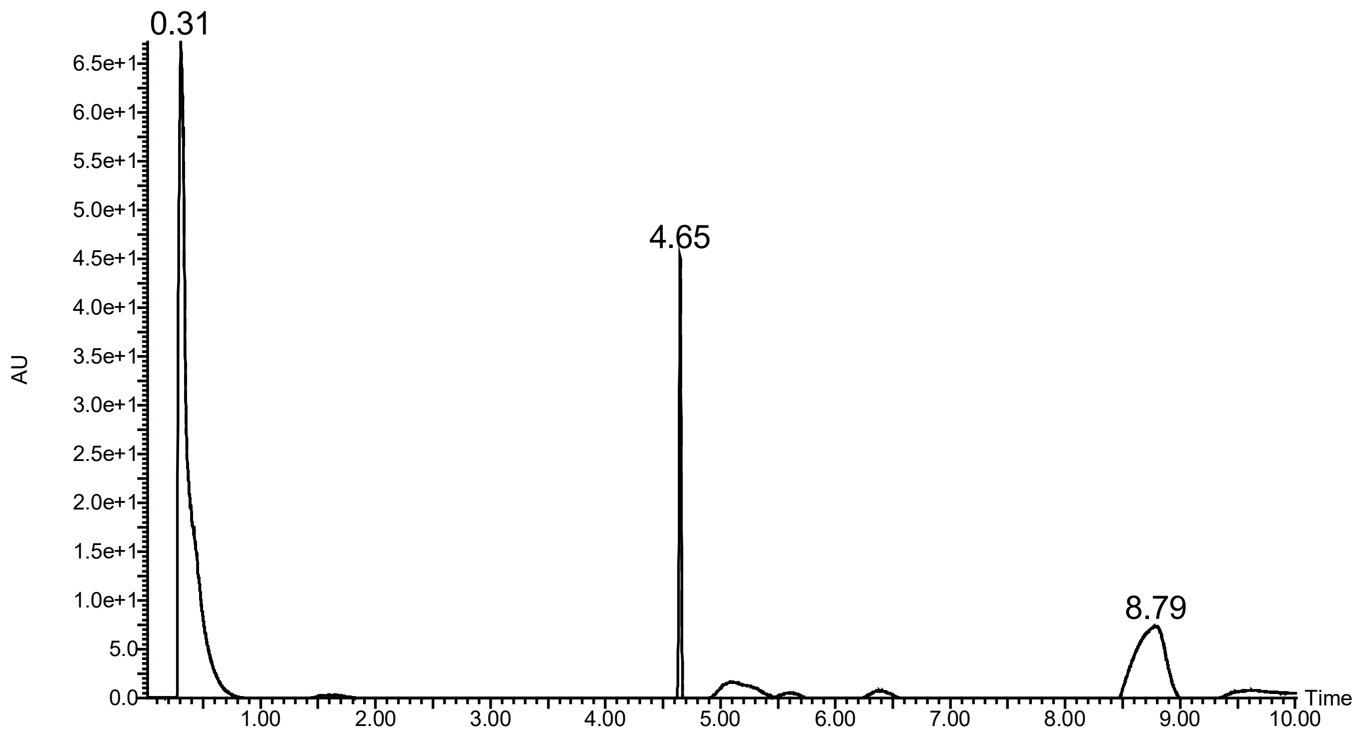

0.31
4.65
8.79
AU
Time
6.5e+1
6.0e+1
5.5e+1
5.0e+1
4.5e+1
4.0e+1
3.5e+1
3.0e+1
2.5e+1
2.0e+1
1.5e+1
1.0e+1
5.0
0.0
1.00
2.00
3.00
4.00
5.00
6.00
7.00
8.00
9.00
10.00


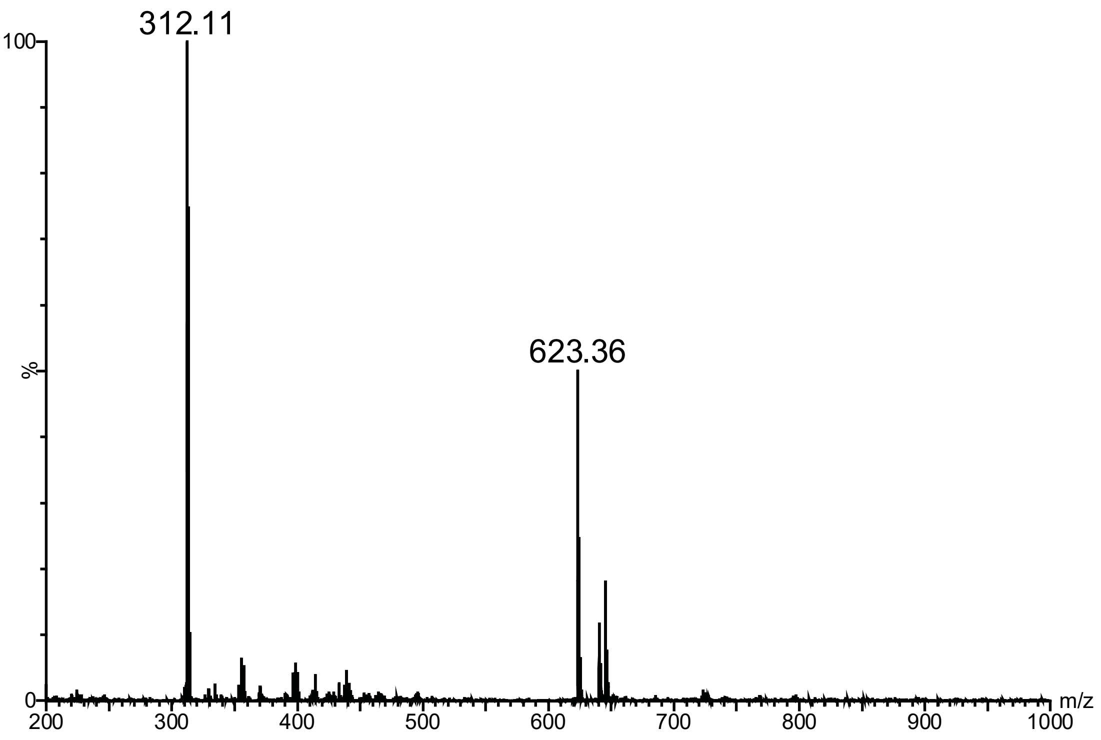

312.11
623.36
100
%
0
m/z
200
300
400
500
600
700
800
900
1000

# BB07CA902

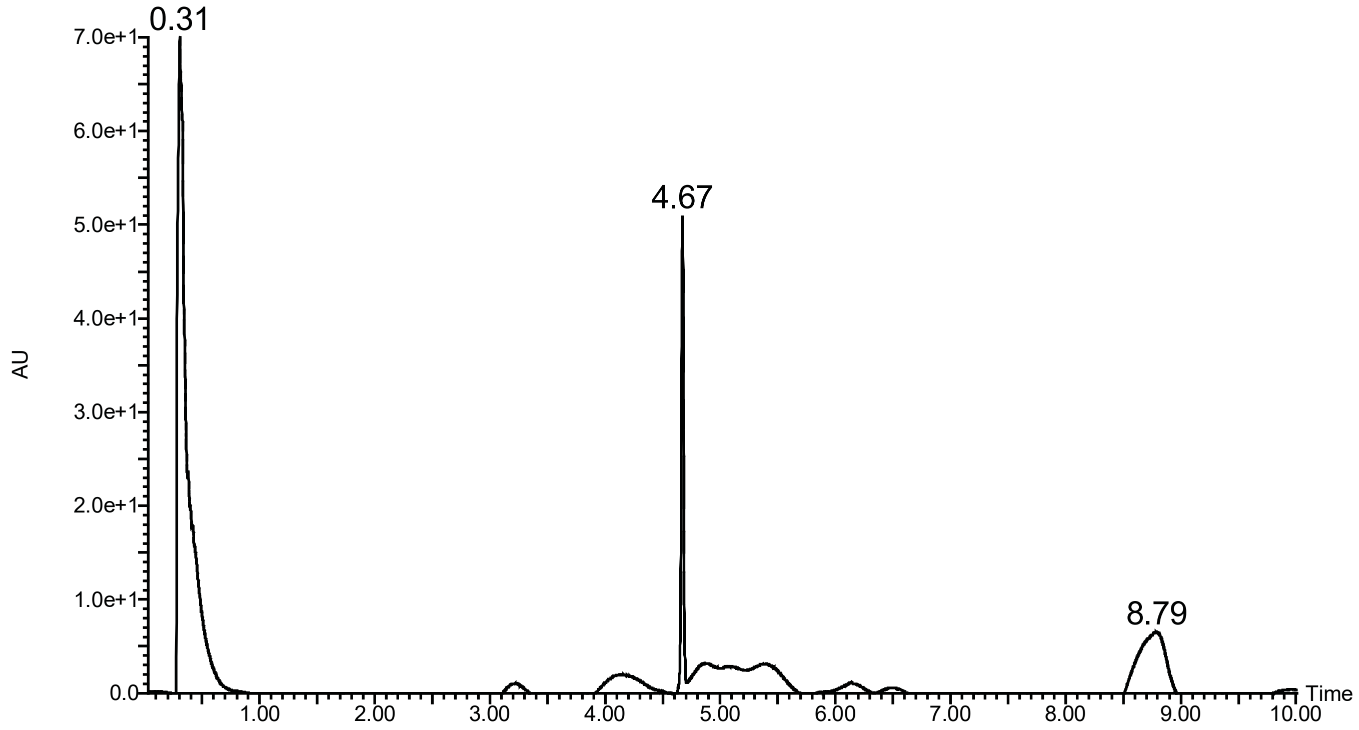

0.31
4.67
8.79
7.0e+1
6.0e+1
5.0e+1
4.0e+1
3.0e+1
2.0e+1
1.0e+1
0.0
AU
Time
1.00
2.00
3.00
4.00
5.00
6.00
7.00
8.00
9.00
10.00


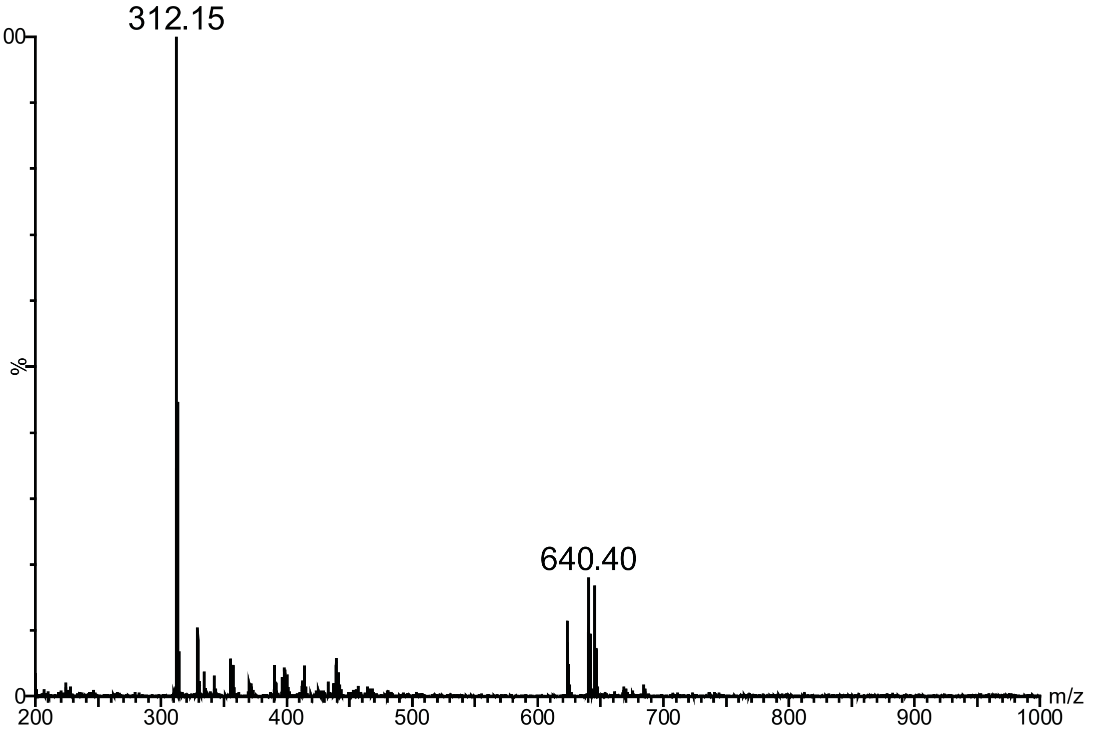

312.15
640.40
100
%
0
m/z
200
300
400
500
600
700
800
900
1000
